# Interactive machine learning for fast and robust cell profiling

**DOI:** 10.1101/2020.02.20.956268

**Authors:** Lisa Laux, Marie F.A. Cutiongco, Nikolaj Gadegaard, Bjørn Sand Jensen

**Author notes:** These authors contributed equally to this work (Lisa Laux, Marie F.A. Cutiongco).

## Abstract

Automatic profiling of cell morphology is a powerful tool for inferring cell function. However, this technique retains a high barrier to entry. In particular, configuring image processing parameters for optimal cell profiling is susceptible to cognitive biases and dependent on user experience. Here, we use interactive machine learning to identify the optimum cell profiling configuration that maximises quality of the cell profiling outcome. The process is guided by the user, from whom a rating of the quality of a cell profiling configuration is obtained. We use Bayesian optimisation, an established machine learning algorithm, to learn from this information and automatically recommend the next configuration to examine with the aim to maximize the quality of the processing or analysis. Compared to existing interactive machine learning tools that require domain expertise for per-class or per-pixel annotations, we rely on users explicit assessment of output quality of the cell profiling task at hand. We validated our interactive approach against the standard human trial-and-error scheme to optimise an object segmentation task using the standard software CellProfiler. Our toolkit enabled rapid optimisation of an object segmentation pipeline, increasing the quality of object segmentation over a pipeline optimised through trial-and-error. Users also attested to the ease of use and reduced cognitive load enabled by our machine learning strategy over the standard approach. We envision that our interactive machine learning approach can enhance the quality and efficiency of pipeline optimisation to democratise image-based cell profiling.

## Introduction

Image-based cell profiling is a powerful tool to capture the intricacies of cell phenotype. The resolution and rapidity stemming from image-based cell profiling has enabled study of mechanisms of and cellular response to disease [1], drugs [2], or materials [3]. Together with the explosion of automated and high-throughput microscopy techniques, image-based cell profiling is increasingly relied on as a biological toolkit. Central to image based profiling are software tools devoted to ease the burden of processing a large volume of images by making detection, segmentation and feature extraction automated [4].

To optimise a cell profiling process or pipeline for a particular image set, users configure the optimal values for various image processing parameters (e.g. image correction, object segmentation and feature extraction) in a trial and error process. The standard tool CellProfiler already reduces this task by carefully curating the most pertinent and widely-used parameters in cell profiling [5]. Yet selecting an optimum set of cell profiling pipeline parameters (or a pipeline ‘configuration’) from the available parameter space is still an onerous task and prone to biases. Optimising an image processing pipeline is biased against those with limited knowledge in biology, microscopy or image analysis. The high cognitive load of pipeline optimisation can inadvertently lead to decision-making bias that deteriorates the quality of the cell profiling result. Testing of pipelines on small datasets can also induce an availability bias, where positive results from small subsets are incorrectly assumed to generalise to the entire dataset. Furthermore, novice users may be susceptible to default bias, where default settings are selected over the true optimal ones. While incredibly informative and powerful for biology, cell profiling is hindered by the users’ capability to process images robustly and reproducibly.

Here, we present a new approach that integrates user input with machine learning to optimise the configuration of a cell profiling pipeline based on high-level quality assessments. We obtain from the user the quality score (QS), a metric to describe the performance of a pipeline configuration. We use a Bayesian optimisation (BO) [6, 7], a machine learning technique to rapidly learn the optimal pipeline configuration by maximising the QS in an iterative fashion. Effectively, we present a process that applies machine learning to divert the burden of pipeline optimisation from the user and automates and accelerates optimisation of pipeline hyperparameters. By using quality maximisation as the explicit target of our optimisation approach, user input bypasses labeling of images at the pixel or object level, which is currently required by prevailing image analysis toolkits [8–10]. Through our interactive machine learning approach, we reduce cognitive load and bias against new users and thus improve the rapidity and quality of cell profiling.

We created new modules on the standard biological toolbox CellProfiler (CP) to implement our interactive machine learning approach. The new modules can be easily integrated within the existing CP software infrastructure. We created two types of modules: evaluation modules to obtain QS from users; and, a BO module to define parameters that will be automatically optimised. Our approach in optimising pipeline configuration uses the evaluation and BO modules together to obtain QS and automatically change pipeline settings towards maximisation of the QS. We then tested our BO based approach to optimise a pipeline configuration for object segmentation. Users with varying levels of expertise obtained more accurate object segmentation using our approach compared to the conventional trial and error scheme. Users also attested to the ease of use of our interactive machine learning approach, with a majority electing to incorporate the process into their own pipeline optimisation process.

The rest of paper is organized as follows. First, we describe the conceptual framework behind our interactive machine learning approach to pipeline optimisation. Next, we present the results of user experiments comparing our approach to the conventional method of pipeline optimisation. Finally, we discuss the implications of our work for scientifically reliable, high quality, and rapid image-based cell profiling for all.

## Semi-automated pipeline optimisation using interactive machine learning

We propose to utilise a semi-automated, machine learning approach to optimise a cell profiling pipeline configuration (Fig. 1). Critical to this approach is the explicit definition of the level of performance of each cell profiling configuration. We thus define the QS as a metric of the quality of a pipeline configuration. We also created a highly customisable BO module that allows the user to define the image processing parameters to be optimised. The QS is then exploited by a BO algorithm to automatically change all user specified image processing parameters simultaneously. The BO process uses the evaluation and BO modules together to iteratively obtain the QS then automatically change pipeline parameters with the goal of QS maximisation. Our approach has been implemented as a collection of stand-alone CP modules which can be used as plugins to the existing software: *ManualEvaluation*, *AutomatedEvaluation* and *BayesianOptimisation* modules. The implementation, module plugins, CP pipelines, training and testing datasets, and results can be found on https://github.com/uofg-cellprofiler-modules/bayesopt4cellprofiler.

**Fig 1.**
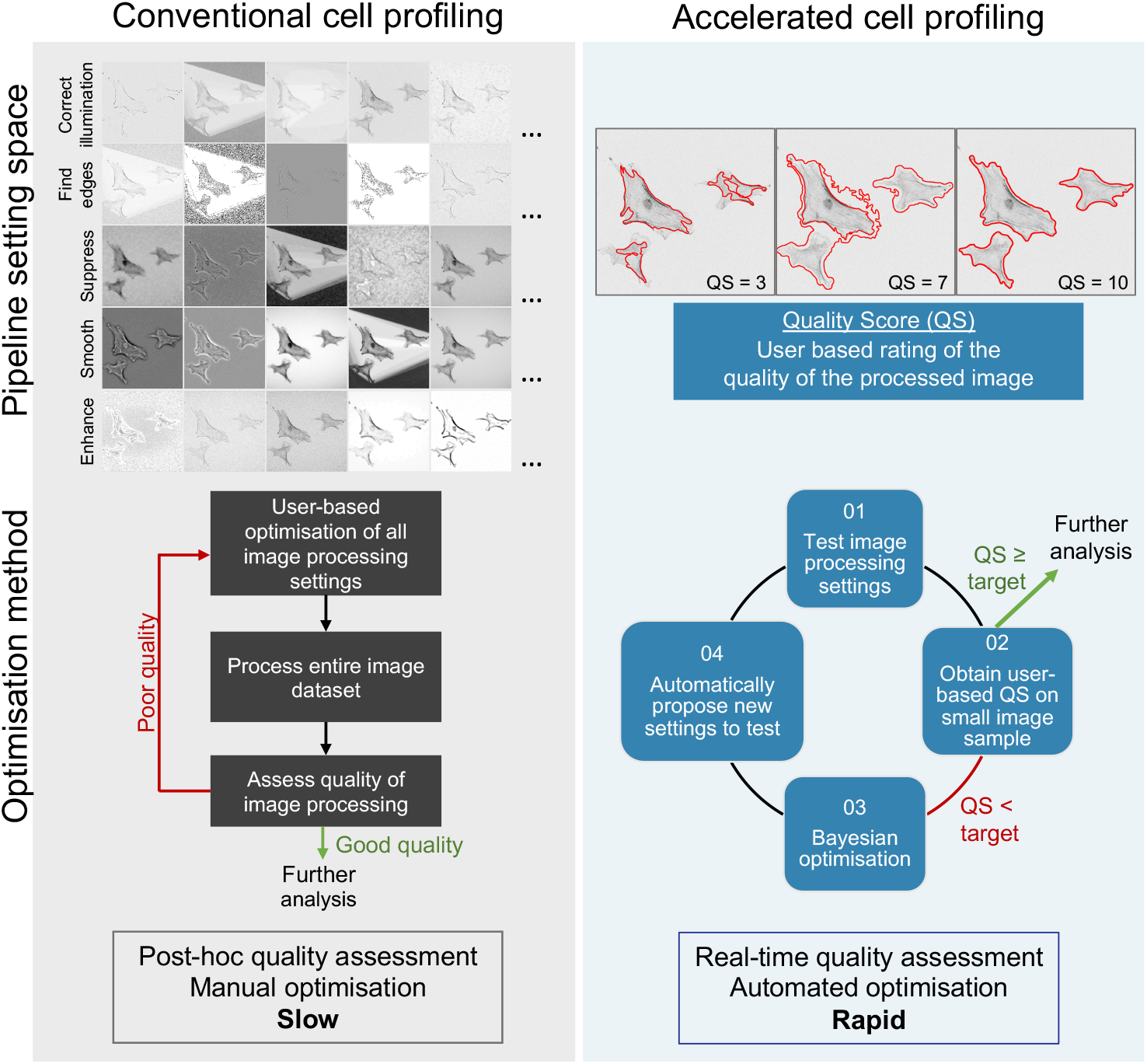
Optimising pipeline configuration through an interactive machine learning approach. The conventional approach to optimising a set of cell profiling parameters (or ‘configuration’) requires the user to change multiple settings in a trial and error manner. This is a slow and tedious process, with quality of the image processing pipelines usually only measured after analysis of the entire dataset. Our proposed approach combines machine learning with explicit definition of quality corresponding to a pipeline configuration obtained from the user in real time (or the quality score (QS)). The burden of choosing a pipeline configuration is then placed on a machine learning algorithm called Bayesian optimisation (BO), which learns the optimum pipeline settings that maximises the QS. Through this interactive machine learning approach, cell profiling can be rapidly optimised, reduce cognitive load on users and ensure high quality outcomes.

### Evaluation modules

The evaluation modules were created to obtain three key pieces of information at each iteration: the target object requiring optimisation, the minimum acceptable QS required by the user (referred to as the ‘target QS’), and the QS from the latest pipeline configuration (referred to as the ‘current QS’). Definition of the target and current QS depend on whether the user will provide a QS at each iteration (manual) or set a criteria that defines robust processing of the target object (automatic). To provide a concrete example, we discuss the application of our evaluation modules for object segmentation, a common bottleneck in pipeline optimisation and image analysis.

#### AutomatedEvaluation

The *AutomatedEvaluation* module automatically evaluates the quality of a pipeline configuration based on user-prescribed criteria that characterise an optimally segmented object (the target QS) (Fig. S2). Thus, *AutomatedEvaluation* requires prior knowledge of the optimally segmented object. For instance, an optimally segmented nucleus rarely contains any concavities, allowing us to define the target QS from high measurements of solidity. At least one target object with its characteristics (e.g. shape, texture, intensity) measured needs to be placed before *AutomatedEvaluation* in the pipeline. When multiple measurements of a segmented object are used, an aggregate is calculated to obtain a target QS. At each iteration of BO, *AutomatedEvaluation* calculates the current QS of the segmented object using the same measurements defined in the target QS. If the current QS falls below the target QS, the BO process continues. When the current QS meets or exceeds the target QS, the BO process stops and the segmented object resulting from the optimised pipeline configuration is displayed. If the user deems segmentation to be poor, the user will be prompted to redefine the target QS.

#### ManualEvaluation module

The *ManualEvaluation* module relies on the user’s subjective rating of a segmented object (Fig. S3). First, the user is required to define the minimum acceptable segmentation quality or target QS on a scale of 1 (poor quality) to 10 (excellent quality). During pipeline execution, *ManualEvaluation* temporarily interrupts the pipeline to display the segmented object from the current pipeline configuration. The user is required to rate the quality of the segmented object using the same rating scale of 1 to 10 to provide the current QS. The BO process will continue to iterate until the target QS is met or exceeded. Both *AutomatedEvaluation* and *ManualEvaluation* allows the user to customise objects and images to be displayed to the user at each iteration of BO.

### BayesianOptimisation module

The *BayesianOptimisation* module implements a BO algorithm to automatically optimise pipeline configuration by maximising the QS (Fig. S4). To do this, we created the highly customisable *BayesianOptimisation* module. *BayesianOptimisation* requires at least one evaluation module placed upstream from which the current QS can be obtained. *BayesianOptimisation* allows the combination of the two evaluation modules, with weighting of contribution to the joint current QS explicitly defined by the user. The *BayesianOptimisation* module also provides full customization of the image processing modules and settings to be optimised using the BO algorithm. Even settings within object identification modules (e.g. *IdentifySecondaryObject* (e.g. threshold correction factor or adaptive window value) can be optimised by the BO process. In principle, any parameters or settings with integer and float values in modules upstream of the *BayesianOptimization* module can be optimised by the BO process. *BayesianOptimisation* also gives the user control of the BO process, including setting the maximum number of iterations of BO.

Together, the evaluation and *BayesianOptimisation* modules aim to minimise the gap between the current QS and target QS by automatically changing pipeline configuration. A pop-up window shows the deviance of current from target QS at every iteration of the BO process (Fig. 2). The BO process iterates until the current QS matches the target QS (i.e. quality gap = 0) or the maximum number of iterations specified by the user has been attained.

**Fig 2.**
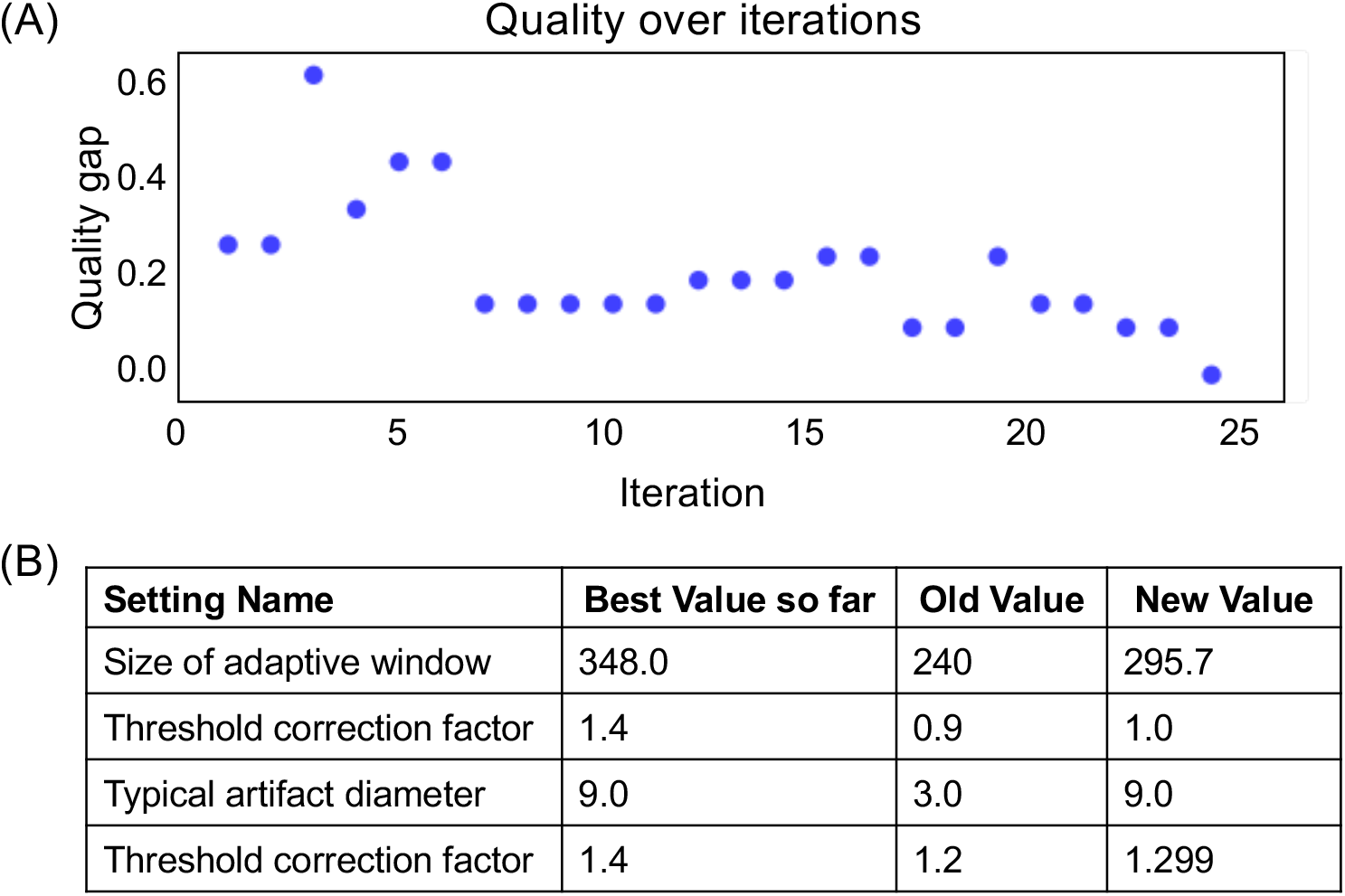
Visualisation of the BO process at every iteration. (A) Scatterplot showing the progress of reducing the quality gap between the current QS and target QS (y axis) across increasing number of BO iterations (x axis). The optimum pipeline configuration is achieved when the quality gap reaches 0 or when the current QS matches or exceeds the target QS. (B) Table showing the parameter settings (column 1) that minimise the deviance between current and target QS (column 2), was tested in the previous iteration (column 3), and is being tested in the current iteration (column 4). The table updates at every iteration of the BO process.

### Bayesian Optimisation algorithm

At the core of the *BayesianOptimisation* module is a custom version of a BO algorithm [6, 7, 11]. BO relies on a surrogate function/model that represents and provides calibrated predictive distributions for the quality score (QS), *y*, for a given pipeline configuration, *x*. We define the surrogate model, *f* (*x*), mapping from configuration to QS as a Bayesian regression model with a Gaussian likelihood, 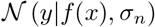, with a Gaussian process (GP) prior on *f* such that 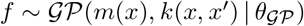 [12]. The GP is defined by the effective mean function, *m*(*x*) = 0, and chosen covariance function 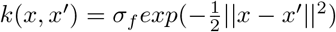 where the hyperparameters are collected in *θ* = {*σ_n_*, *σ_f_*, *σ*_*ℓ*_}. Given the GP and a training set, 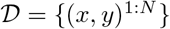, containing a certain pipeline configuration and its corresponding QS, the predictive distribution for any pipeline configuration, *x*\*, is directly available as *p*(*y*\*|*x*\*, *D*, *θ*). This allows us to estimate both the expected QS and its uncertainty for all unseen configurations. For simplicity, we have defined the model without priors on the hyperparameters and we do marginal likelihood optimisation of the hyperparameters (after an initial bootstrap phase). However, some BO algorithm hyperparameters and GP parameters (e.g. the length scale of the covariance function and the assumed noise level) can be customised in the *BayesianOptimisation* module.

The BO process exploits the predictive distribution at any point in the optimisation process to sequentially choose the next set of image processing parameters (i.e. the configuration) to evaluate. It does so by trading-off the desire to optimise the current QS with the implicit need to learn the surrogate model. To do so, here we applied Expected Improvement [6, 7]. At the end of each iteration, the current QS from the newly chosen pipeline configuration is subsequently included in the training set and the model re-estimated before repetition of the BO process. A summary of the BO process is given in (Fig. S1).

## User experiments

### Methods

User based experiments in pipeline optimisation for object segmentation were performed. These experiments were conducted to test our interactive machine learning approach against the trial and error (here referred to as ‘conventional’) method of optimising a pipeline configuration. Experiments involving human subjects were performed with approval from the Ethics committee of the College of Science and Engineering, University of Glasgow (case no. 300180170).

Participants were randomly assigned the objective of segmenting either cells or focal adhesions. Pipelines for both objectives were designed to have interdependent modules, where segmentation of cells and focal adhesions were dependent on nuclei and cell segmentation, respectively. Each participant was required to optimise 1 pipeline using the conventional approach, and 3 pipelines using our interactive machine learning approach. Participants were given 20 minutes to optimise each pipeline. In the conventional approach, participants were required to optimise settings across prescribed modules in a trial and error manner. Using the BO approach, participants were required to use *BayesianOptimisation* in conjunction with either *AutomatedEvaluation*, *ManualEvaluation* or both evaluation modules (called ‘Composite Evaluation’). A summary of the pipeline configuration automatically optimised by *BayesianOptimisation* is found in Table S1 and Table S2 for cell and focal adhesion segmentation, respectively. A summary of all tasks performed by each participant is summarised in Table 1. All tasks were conducted on the same computer running CellProfiler v3.1.8. CP pipelines and image sets used in both tasks are included in Supporting Information 1.

**Table 1.** Tasks in user based testing of the interactive machine learning approach for object segmentation. All users were asked to optimise cell or focal adhesion segmentation using an interactive machine learning or the conventional (trial and error) approach. All tasks used the same image sets for training and testing

| Task | Pipeline parameter setting | Quality evaluation |
| --- | --- | --- |
| Conventional | User based | None |
| Automated evaluation | *BayesianOptimisation* module | *AutomaticEvaluation* module |
| Manual evaluation |  | *ManualEvaluation* module |
| Composite evaluation |  | *AutomaticEvaluation* and *ManualEvaluation* modules |

Each pipeline optimisation task used identical image sets for training and testing. At the end of each task, 10 images were used to test the quality of the resulting pipeline configuration. At the end of each task, participants rated the QS of 10 images run through the resulting pipeline configuration. To provide a baseline measurement, participants also rated the QS of test images segmented using a pipeline optimised by a CP expert. Participants also rated the difficulty of each completed optimisation task using a Likert scale. The task sheet used to instruct and survey users is provided in Supporting Information 2.

To provide a quantitative measure of segmentation accuracy, we calculated the pixel-wise intersection over union (IoU) score of test images segmented by users (via CP), *M_user_*, against the ground truth (binary) segmentation mask, *M_groundtruth_*. Ground truth segmentation masks were obtained manually in ImageJ [13]. For each mask, we calculated the IoU score for individual images as follows (e.g. [14]):

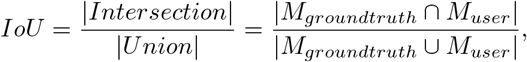

By normalising the number of pixels common to both masks by the total number of pixels in both masks, the IoU metric detects how well the predicted mask encompasses the ground truth without regard to actual (*x*, *y*) coordinates of the masks. An IoU score of 1.0 thus typically indicates excellent segmentation, and an IoU score less than 0.5 denotes poor segmentation.

IoU scores for cell segmentation were pooled from users assigned to both the cell and focal adhesion segmentation tasks. A one-way ANOVA with Tukey’s post-hoc test for pairwise comparison was used to test statistical significance in IoU between different optimisation approaches. Masks of and IoU scores measured from test images segmented by each user are given in Supporting Information 3.

A detailed description of methods (including cell preparation, image acquisition, and participant recruitment) are provided in Supporting Methods.

## Results

Here, we tested the performance of our interactive machine learning approach. We compared the quality of the segmentation, ease of use, and speed of optimisation between our approach and the conventional method of pipeline optimisation. First, we showed that our approach significantly enhanced segmentation over the conventional method. Immediately after each task, users rated the QS of images segmented by the interactive machine learning approach to be overwhelmingly higher than the conventional approach (Fig S6). Quantitative measurement of segmentation quality using the IoU score reinforce this (Fig 3). In particular, providing QS of cell segmentation in real time through the use of *ManualEvaluation* significantly improved the consistency and magnitude of the IoU score compared to the conventional approach. Use of either *ManualEvaluation* (mean *±* standard deviation 0.75 *±* 0.14) or the composite (0.69 *±* 0.24) evaluation mode yielded IoU scores for cell segmentation similar to those that can be achieved by a CP expert (0.74 *±* 0.20), indicating excellent segmentation quality.

**Fig 3.**
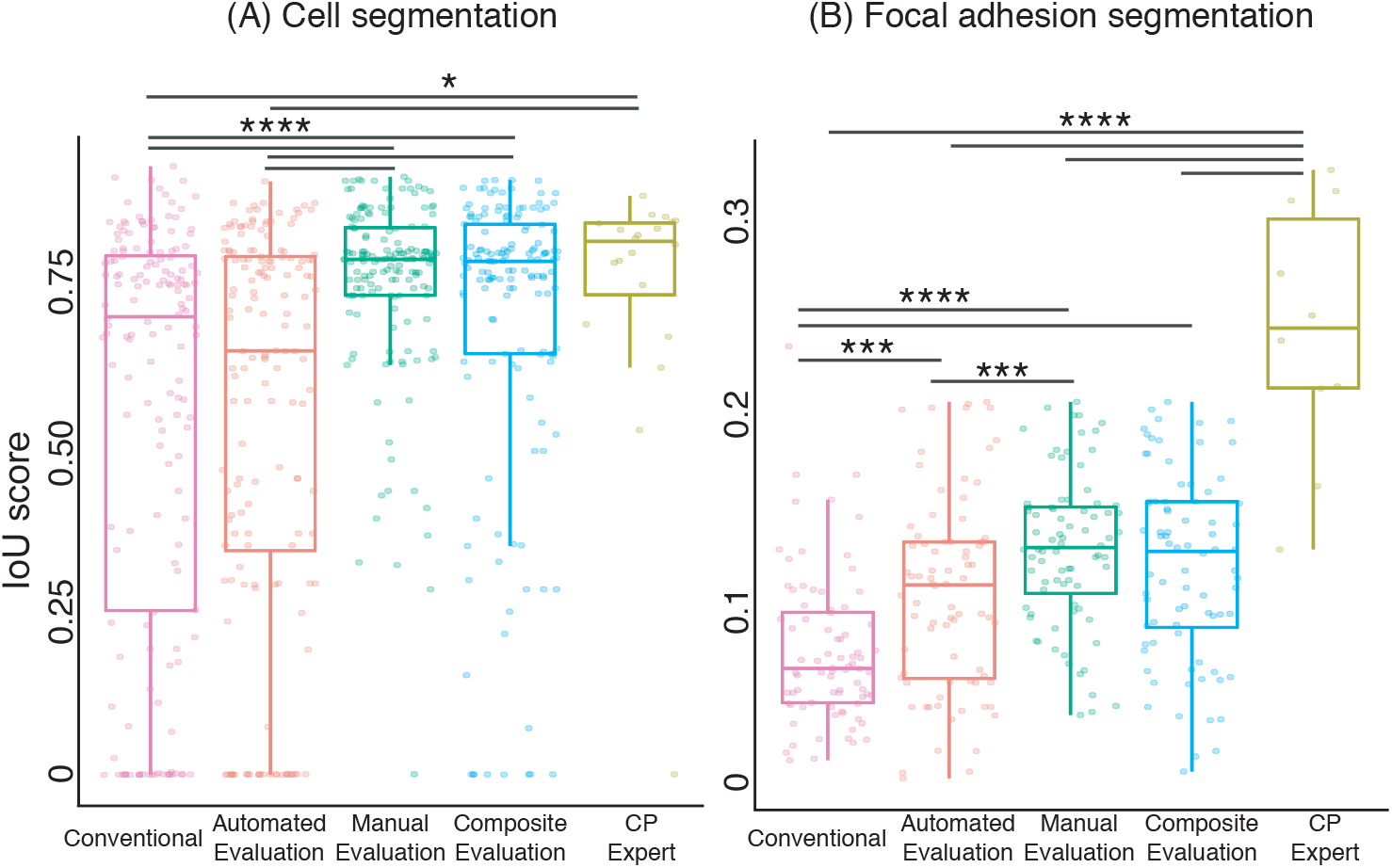
Improved object segmentation using an interactive machine learning approach. To optimise cell profiling pipeline configuration, *BayesianOptimisation* was tested together with *AutomatedEvaluation*, *ManualEvaluation* or both evaluation types (Composite Evaluation). Additionally, we measured segmentation of objects from a pipeline optimised by a CP expert. We calculated the intersection over union (IoU) score of each user-segmented against the corresponding ground truth image to measure quality of segmentation. An IoU score of 1 indicates user-based segmentation equivalent to the ground truth. Each point represents the IoU score calculated from one image segmented by one user. We denote statistical significance of * at p <0.05, ** at p <0.01, *** at p <0.005, **** at p <0.0001. (A) n = 160 image sets from 16 participants and n = 80 image sets from 8 participants.

In contrast to cell segmentation, all 3 modes of the interactive machine learning approach outperformed the conventional approach in segmenting focal adhesions. IoU scores for focal adhesion segmentation using the *AutomatedEvaluation*, *ManualEvaluation* and Ccmposite evaluation mode produced IoU score with mean *±* standard deviation of 0.11 *±* 0.05, 0.13 *±* 0.04, and 0.12 *±* 0.05, respectively. The conventional approach only yielded an IoU score of 0.08 *±* 0.04 (mean *±* standard deviation), while a CP expert reached a score of 0.24 *±* 0.07. We noted that none of the evaluation modes yielded segmentation quality similar to what a CP expert can achieve. Furthermore, none of the 5 segmentation approaches resulted in adequate segmentation of focal adhesions compared to the ground truth. The difficulty of focal adhesion segmentation is a well established issue and is caused by variability in intensity and size of these structures. The difficulty of this task is evidenced by the fact that even recent tools developed to improve focal adhesion segmentation are limited in throughput and requires high resolution microscopy (such as total internal reflection microscopy) [15, 16].

Though it failed to show a benefit for cell segmentation, *AutomatedEvaluation* improved focal adhesion segmentation. Presumably, measurements in the ratio scale that easily define focal adhesions (e.g. ellipticity and solidity) were easier to intuit and exploit compared to measurements in the interval scale (e.g. cell area). Under certain circumstances or for users with some experience, *AutomatedEvaluation* presents advantages for pipeline optimisation.

We noted that the use of *ManualEvaluation* (by itself or compositely with *AutomatedEvaluation*) was advantageous for object segmentation. Indeed, despite having different characteristics, both cells and focal adhesions were accurately segmented when using *ManualEvaluation*. Presenting visual evidence (Fig 4) allow users to evaluate the conformity of outlines to the edges of target objects. This is a critically simpler task than setting criteria to define optimal object segmentation, which may be unknown *a priori*, as required by *AutomatedEvaluation*.

**Fig 4.**
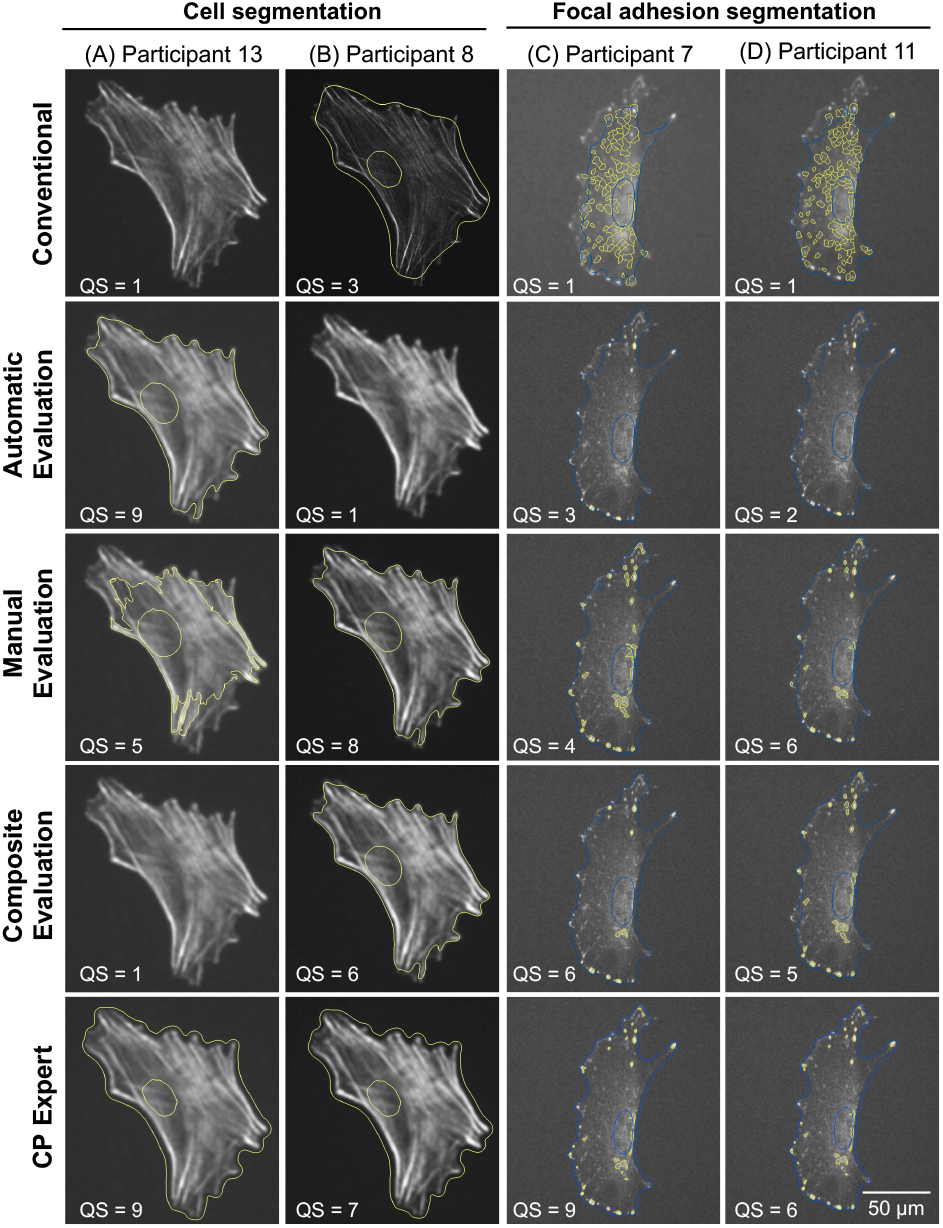
The interactive machine learning approach results in high quality segmentation. Representative images of segmented cells and focal adhesions selected from random users.

Next, we assessed the ease of use of the interactive machine learning approach (Fig 5). When asked to use *AutomatedEvaluation*, the number of users who found pipeline optimisation to be easy doubled in number. Feedback on *ManualEvaluation* was even more positive, as all participants considered pipeline optimisation to be easy when using this evaluation mode. Participants also overwhelmingly (15 out of 16 or 93.8%) elected to adopt our approach for future pipeline optimisation, indicating broad support for our interactive machine learning approach to optimise cell profiling. In line with poorer IoU scores compared to our approach, only a minority (3 out of 16 or 18.8%) of users found it easy to optimise pipeline configuration using the conventional method.

**Fig 5.**
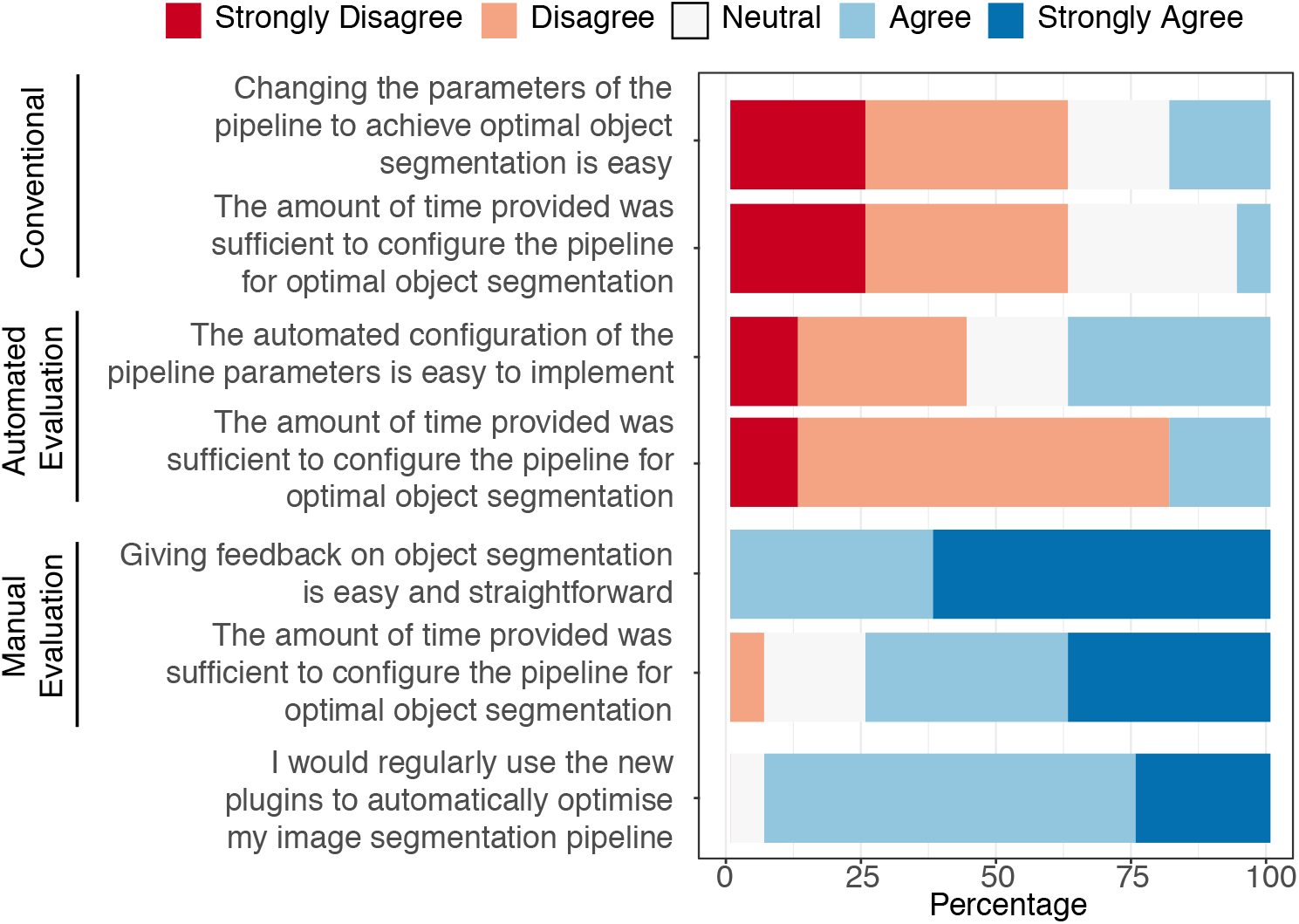
Ease of usage of the interactive machine learning approach for pipeline optimisation. Survey data were accumulated from all participants from both experiments (n = 16).

Finally, we demonstrated the efficiency of our approach over the conventional method for pipeline optimisation. Prior to user based experiments, we tested our approach against a method that randomly selected parameters of pipeline modules (Fig S5). The random selection process approximated the conventional trial and error method. On average, our approach required less iterations to optimise nucleus, cell and focal adhesion segmentation compared to the conventional approach. User based experiments supported these findings, where 10 out of 16 (62.5%) users required more than 20 minutes to sufficiently optimise a pipeline using the conventional method (Fig 5). A large number (13 out of 16 or 81.3%) of users found 20 minutes insufficient for pipeline optimisation using *AutomatedEvaluation*. Meanwhile, a majority of users (12 out of 16 or 75%) found that 20 minutes was sufficient to optimise a pipeline using *ManualEvaluation*, regardless of prior experience in cell profiling. We showed here that our approach empowers robust and rapid cell profiling without compromising on ease of use, cognitive burden, or bias against novice users.

## Discussion

Robust and reproducible image-based cell profiling depends on the optimal configuration of the image processing pipeline. The conventional method of optimising an image processing pipeline is effectively a trial and error process, and is thus time consuming, tedious and prohibitive to those with minimal experience in image analysis or biology. Here, we propose a semi-automated approach that relies on minimal user intervention and machine learning to accelerate pipeline optimisation, and enhance the quality of cell profiling.

A key component in our proposed approach is the iterative acquisition of the QS from the user. By obtaining a QS corresponding to a certain pipeline configuration, we were able to effectively incorporate learning into the process of pipeline parameter optimisation. This was performed using a BO algorithm. Importantly, the BO algorithm is ideal for optimising broad parameter spaces such as in synthetic gene design [17], hyperparameter tuning [7] or crystal structure prediction [18]. Here, we also showed that the BO algorithm optimised the broad combinatorial space for image processing parameters across multiple segmentation objectives. This is especially important for users with little to no experience in image analysis, where the BO algorithm can reduce default bias in pipeline optimisation.

The BO algorithm is also an effective remedy to memory bias, which increases in propensity with longer and more complex pipelines. Because the conventional method relies on a user to remember outcomes corresponding to a image processing configuration, the process is highly susceptible to memory and cognitive biases. Not only do these biases severely narrow the setting space being tested, they prevent users from obtaining the optimum processing pipeline that is crucial to accurate cell profiling. Diverting the user’s focus towards providing the QS is also an essential feature of our method that reduces cognitive load on users without compromising on the quality of pipeline outcomes.

Though intended for completely autonomous optimisation [19], here we modified BO to incorporate a human-in-the-loop [20, 21]. By relying on the user instead of absolute limits to determine QS, we have created a more generalised and flexible approach to assess and optimise pipeline performance. Without predefined limits on quality (as is most apparent with the *ManualEvaluation* module), our approach can optimise pipelines for segmentation of objects with complex geometric properties (e.g. the mitochondria). We can even extend the pipeline optimisation process for tasks with undefined quality metrics (e.g. illumination and background correction [22] or for curation of images for quality control [23]).

The flexibility of our approach for pipeline optimisation is also extended to the implemented modules, where users have control over: 1) the task; 2) the target QS; 3) modules and settings; 4) weighting between automatic and manual evaluation into a composite evaluation score; and 5) BO hyperparameters. The modularity of the CP also permits multiple BO runs throughout a single pipeline to optimise various tasks. Complex tasks such as focal adhesion segmentation undoubtedly benefit from this scenario, where there are interdependencies between segmented objects.

A few established tools for cell profiling utilise a similar flavor of machine learning (user supervised, and interactive machine learning) to aid object segmentation by requiring the user to provide input. For instance, the ilastik and weka toolkits rely on user-annotated training data at the pixel (classification of pixels) or object level (binary segmentation mask) [8, 9]. To ensure high quality output using these tools, there is a requirement for an adequately sized training dataset. However, segmentation quality reduces over time from fatigue and attentional bias. To some extent, the expense from low-level annotation can be overcome by using an active learning approach that carefully selects the most informative datasets to require user-based annotation [24]. Nonetheless, in all these toolkits, the main aim is to identify and classify objects. As a consequence of this primary goal, these toolkits also require the input of a domain expert to generate high quality training data.

In contrast, our approach only requires the user to provide input on a higher level of abstraction – quality. With the real time visualisation of an output that is related to an image processing task, our approach dismisses the requirement for high quality annotation of datasets for training. By focusing on optimising the configuration and hyperparameters of a pipeline (rather than labelling of training data for individual elements in the pipeline), the user based quality rating is relevant to the task at hand and unconstrained to segmentation or classification. Indeed, our approach certainly reduces the bias against users who may not have the same domain knowledge to pass accurate judgement on training data, and the data expense required to train a machine learning algorithm. Furthermore, the existing infrastructure for classification can be complemented by a dedicated approach for high quality image processing and segmentation offered by our interactive machine learning approach.

The rapidity by which we collect data calls for fully or semi-automatic methods of cell profiling that is adaptable to different experimental designs, biological systems and imaging modalities. Many are developing machine and deep learning methods to eliminate human intervention in the data analysis process. However, it is difficult and often counter productive to eliminate the user, who has expertise to validate, configure, fine-tune parameters and label data under novel conditions. Here, we show that allowing the user to interactively provide feedback to a machine learning algorithm improves both automation and quality of analysis. Our pipeline using an interactive machine learning approach presents a new paradigm wherein human decision-making and oversight is required for robust scientific discovery.

## Supporting information

All supplementary files

## Acknowledgments

We thank the team at the Broad Institute for creating the open-source, image-based cell profiling platform CellProfiler.

NG and MFAC acknowledge support from the European Research Council (ERC) through the FAKIR 648892 Consolidator Award. NG and BSJ acknowledge support from a Cancer Research UK (CRUK) Pioneer Award. MFAC is additionally supported by the University of Glasgow MG Dunlop Bequest and the College of Science and Engineering Scholarship. BSJ acknowledge support from the Engineering and Physical Sciences Research Council (EPSRC) grant EP/R018634/1. The funders had no role in study design, data collection and analysis, decision to publish, or preparation of the manuscript.

## Supporting information

**Supporting Information 1 File.** Pipelines and raw images for the various tasks carried out in user experiments.

**Supporting Information 2 File.** Participant information sheet, consent form and task sheet used for user-based experiments.

**Supporting Information 3 File.** Image masks used for scoring quality of segmentation using the intersection over union score.

**Supporting Information 4 File.** Survey based results on quality score of image segmentation and difficulty of pipeline optimisation.

## Supporting methods

### Image acquisition

#### Cell culture

MC3T3 cells (passages 10-12, ATCC) were cultured using standard cell culture practice. Cells were grown in growth media comprised of α-MEM with nucleosides and L-glutamine without ascorbic acid and supplemented with 10% FBS and 1% penicillin-streptomycin [25]. Cells were seeded at a density of 4000 cells/cm^2^ on injection moulded and surface-texturised polycarbonate substrates [3, 26].

#### Immunofluorescence staining

MC3T3 cells were cultured for 2 days before fixation using 4% paraformaldehyde. Cells were then stained with DAPI, AlexaFluor conjugated-phalloidin (ThermoFisher, 1:200) to detect the nucleus and the actin cytoskeleton, respectively. On the same cells, focal adhesions were visualised using an anti-talin1 (Abcam 71333, 1:200) and an appropriate AlexaFluor conjugated secondary antibody (ThermoFisher, 1:500). Cells were then mounted on 0.17 μm thick glass coverslips before imaging.

#### Fluorescence microscopy

Images of fluorescently stained cells were obtained using an EVOS FL2 Auto system (ThermoFisher) with 40X magnification (numerical aperture = 1.3). Image sets of the nucleus, the cell (visualised using the actin cytoskeleton) and focal adhesions were used to test the newly developed CellProfiler modules.

### Participant recruitment

Participants were informed that the purpose of the study was to evaluate the performance of our CP modules for object segmentation. Participants provided consent to participate by signing the informed consent forms. The participants were required to have a basic understanding of computer aided image analysis. Participants were offered minor monetary compensation (10 GBP), and the possibility to win a larger amount (50 GBP) in a raffle. A total of 16 participants were recruited, all of whom showed varying levels of experience in image analysis. The participant information sheet, consent form and task sheets are provided in Supporting Information 2.

### Supporting figures

**Fig S1.**
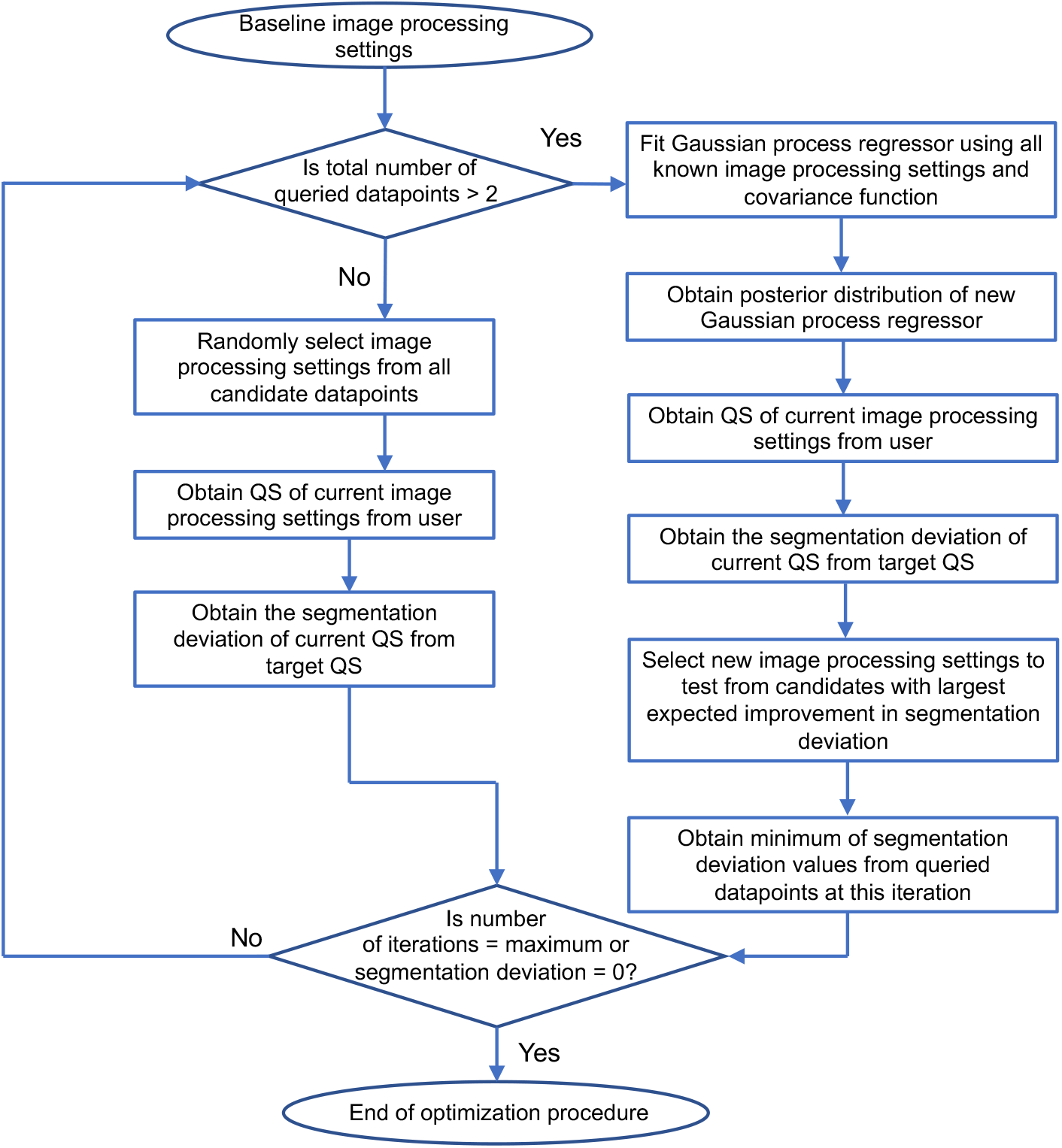
Flowchart of the specific incarnation of the BO algorithm used in the experiments. The BO algorithm is first initialised with two randomly generated settings for pipeline configurations. A Gaussian process (GP) is estimated from all evaluated pipeline configurations and its corresponding QS (acquired from the evaluation modules at each iteration as the current QS). The GP generates a predictive distribution for all pipeline configurations, each with an expected QS and uncertainty. To choose the next pipeline configuration to evaluate, the BO algorithm uses an Expected Improvement function to trade off maximisation of QS with the need to fully learn the GP. From the chosen pipeline configuration, a current QS is obtained from the user. This two-step process of (i) estimating the GP using all evaluated pipeline configurations and corresponding QS, and (ii) selecting the pipeline configuration to evaluate is repeated until the deviation of the current from the target QS is minimised or the user-defined maximum number of iterations have been reached.

**Fig S2.**
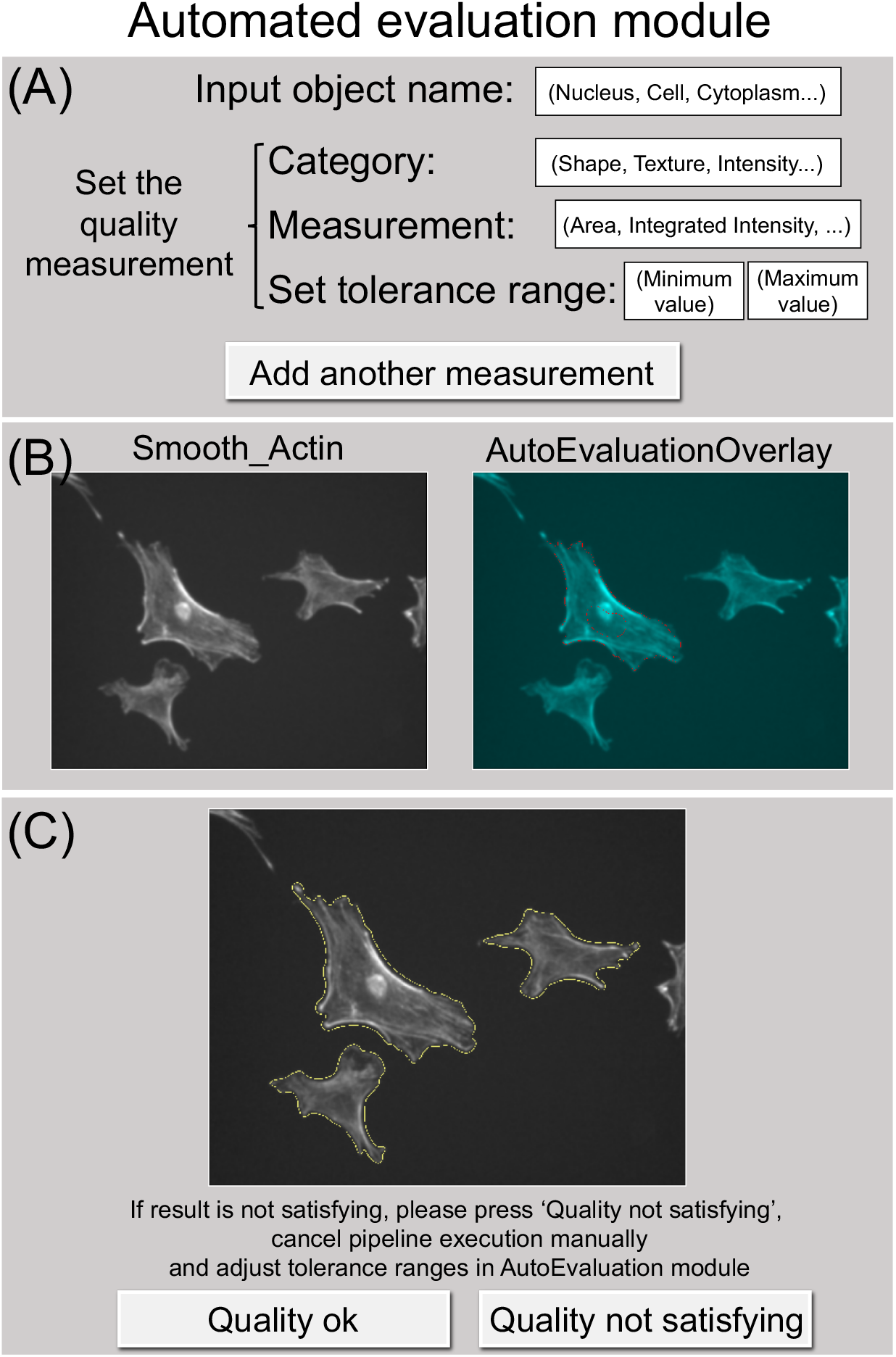
The *AutomatedEvaluation* module. A representation of the *AutomatedEvaluation* module shows the settings that need to be defined by the user. Values placed inside parenthesis show examples of possible input in each module setting. (A) The module allows the user to define an image for visualisation of the segmented object. The segmented object to be optimised is specified in the first level of objects to display. Optionally, other objects that require visualisation can be added to the same image. The module requires tolerance ranges or limits for at least one object measurement (e.g. Area, Perimeter) that define optimal segmentation. The user can define a maximum of 4 different object measurements, which are aggregated to calculate the target QS. (B) At every iteration, the *AutomatedEvaluation* module displays the segmented object resulting from the current pipeline configuration. (C) At the end of the BO procedure (i.e. when current QS meets or exceeds the target QS), the segmented object obtained from the optimum pipeline settings is displayed.

**Fig S3.**
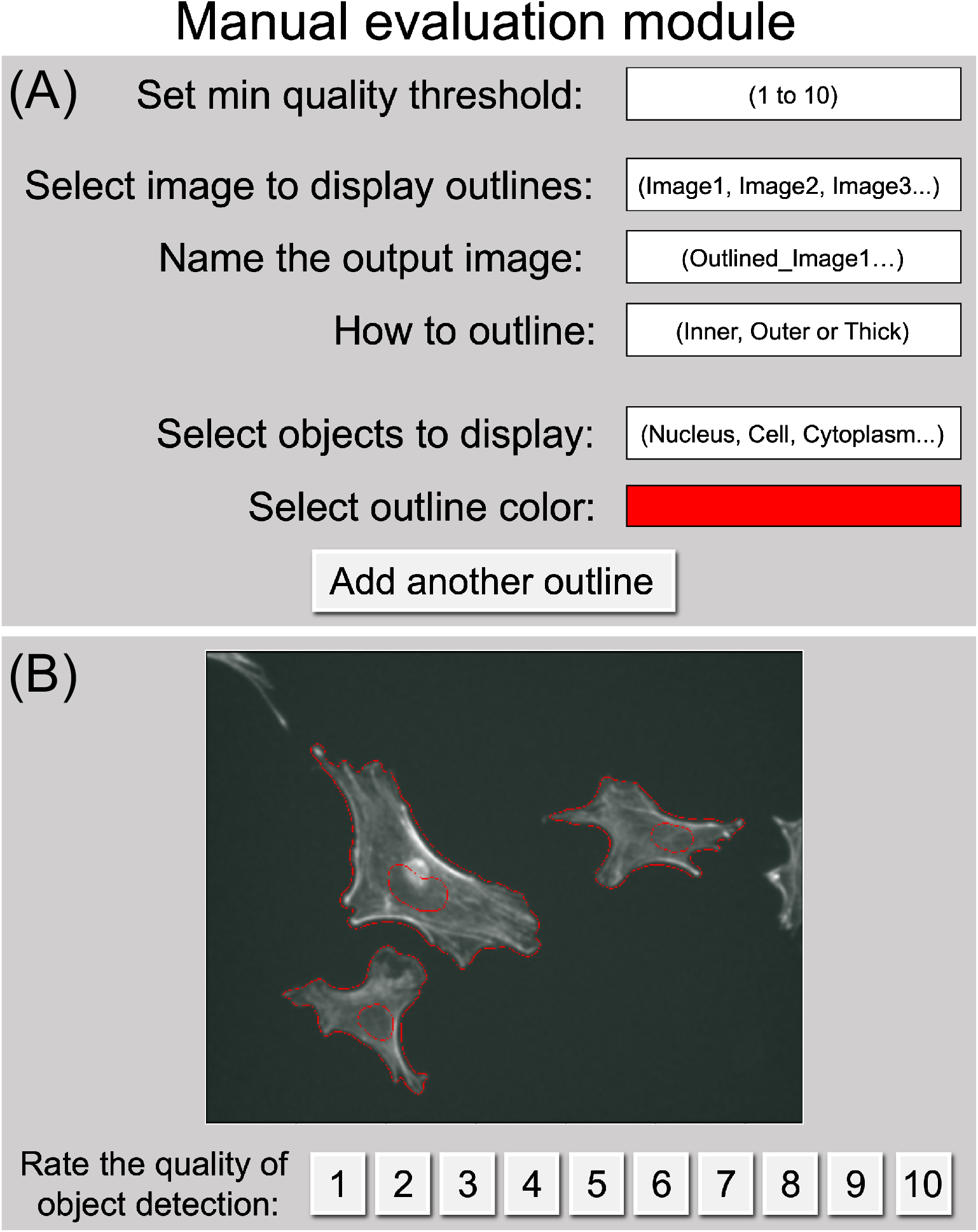
The *ManualEvaluation* module. A representation of the *ManualEvaluation* module shows the settings that need to be defined by the user. Values placed inside parenthesis show examples of possible input in each module setting. (A) The target QS is defined by the user using a scale of 1 (poor quality) to 10 (excellent quality). For visualisation, the image on which to overlay segmentation outlines, the name of the output image and the type of object outline needs to be defined by the user. The segmented object to be optimised is specified in the first level of objects to display. Optionally, other objects that require visualisation can be added to the same image. (B) At each iteration of the BO process, a pop-up window displays the segmented object from the current recent pipeline configuration. (F) The user is required to rate the segmented object to provide the current QS.

**Fig S4.**
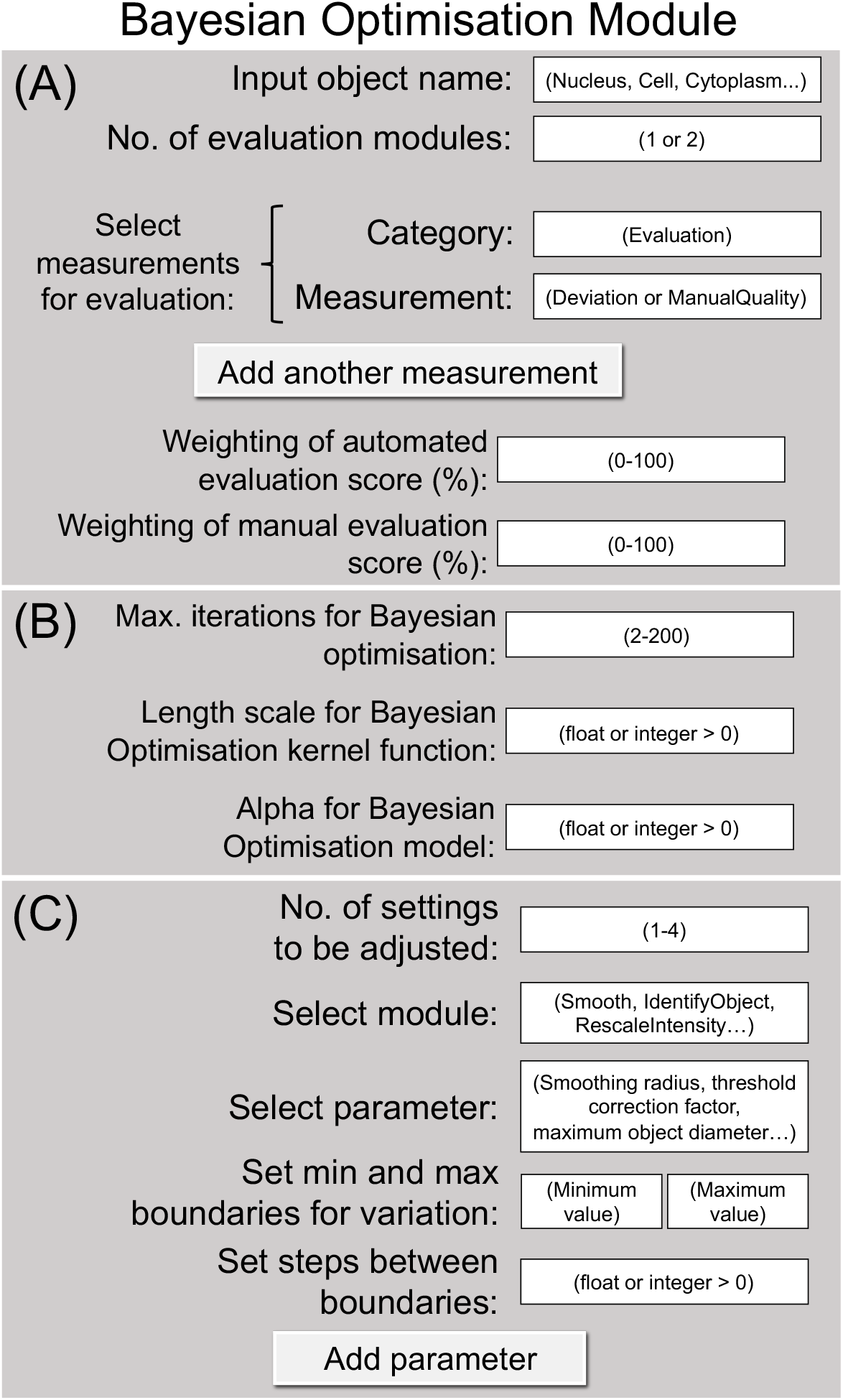
The *BayesianOptimisation* module. A representation of the *BayesianOptimisation* module shows the settings that need to be defined by the user. Values placed inside parenthesis show examples of possible input in each module setting. (A) The target object and the number of evaluation modules to be used are first specified. The evaluation modes to be used will automatically propagate values for evaluation, depending on the available evaluation modules placed upstream of *BayesianOptimisation*. When using both *ManualEvaluation* and *AutomatedEvaluation* modules, the weighted contribution of the current QS from each evaluation module can be explicitly defined. (B) Parameters of the BO algorithm, such as the maximum number of iterations and covariance function hyperparameters, can be tuned by the user. (C) The pipeline parameters to be optimised by the BO process is easily customised by the user. A minimum of one pipeline parameter needs to be optimised for the BO algorithm to proceed. The specific parameters requiring optimisation is defined individually and explicitly (including minimum, maximum, and interval). All evaluated pipeline parameters and its corresponding QS are saved in *.txt* files.

**Fig S5.**
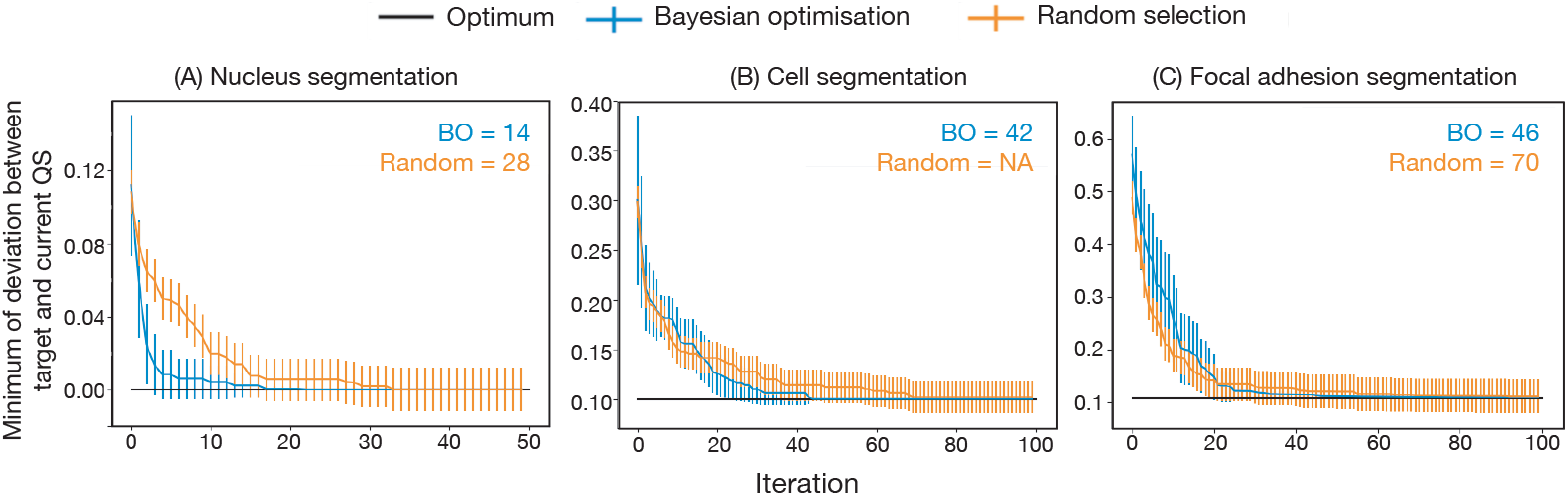
Efficiently optimising a pipeline for object segmentation using an interactive machine learning approach. Segmentation of (A) nuclei, (B) the cell body, and (C) adhesions were tested. Our BO-based approach was used to rapidly minimise the segmentation deviation between the target (black) and the current QS (blue). Random selection (orange) of pipeline parameters was used as a comparison. Data are presented as mean *±* standard deviation/2 from (A) n=50 and (B)(C) n=100 repetitions.

**Fig S6.**
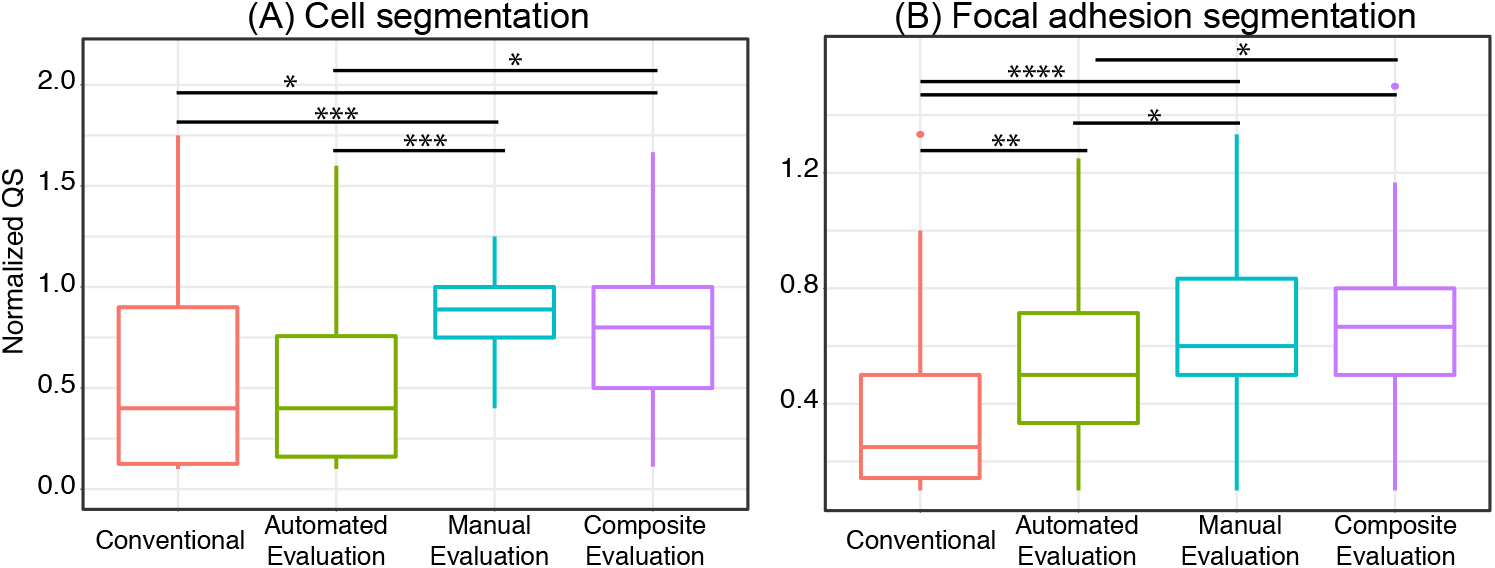
Improved QS using an interactive machine learning approach. To optimise cell profiling pipeline configuration, *BayesianOptimisation* was tested together with *AutomatedEvaluation*, *ManualEvaluation* or both evaluation types (Composite Evaluation). QS of an image obtained from the indicated optimisation mode was normalized against QS of the same image processed by a CP expert (’Normalised QS’). For segmentation of (A) cells and (B) focal adhesions, normalized QS increased by using our interactive machine learning approach. Statistical analysis was performed on raw QS of each image across different optimisation modes using Friedman test for rank based analysis of paired samples with Dunn’s post-hoc test for pairwise comparison. We denote statistical significance of * at p <0.05, ** at p <0.01, *** at p <0.005, **** at p <0.0001. (A) n = 72 image sets from 8 participants and (B) n = 80 image sets from 8 participants.

### Supporting tables

**Table S1.** Pipeline parameters automatically optimised in cell segmentation using the interactive machine learning approach.

| Object | Module | Setting | Minimum value | Maximum Value | Interval |
| --- | --- | --- | --- | --- | --- |
| Nucleus | Identify Primary Objects | Threshold correction factor | 0.9 | 1.5 | 0.1 |
| Cell body | Identify Secondary Objects | Size of adaptive window | 50 | 350 | 25 |
|  |  | Threshold correction factor | 0.9 | 1.5 | 0.05 |
|  | Smooth | Typical artifact diameter | 2 | 10 | 1 |

**Table S2.** Pipeline parameters automatically optimised for focal adhesion segmentation using the interactive machine learning approach.

| Object | Module | Setting | Minimum value | Maximum Value | Interval |
| --- | --- | --- | --- | --- | --- |
| Cell | Identify Secondary Object | Size of adaptive window | 50 | 350 | 50 |
| Focal adhesions | Identify Primary Objects | Threshold correction factor | 0.8 | 1.5 | 0.05 |
|  |  | Size of smoothing | 0 | 20 | 1 |
|  |  | Suppress local maxima that are closer than this minimum allowed distance | 0 | 20 | 1 |

