## Supplementary material for "Interactive machine learning for fast and robust cell profiling": All supplementary files: ParticipantInformationSheet.pdf

### Participant Information Sheet

**Project title**: Interactive machine learning for accelerated analysis of microscopy images.

***Investigators***: Lisa Laux and Marie FA Cutiongco under the supervision of Bjørn Sand Jensen

#### Invitation

You are invited to take part in a research study. Before you decide, it is important for you to understand why the research is being done and what it will involve. Please take time to read the following information carefully. Ask us if there is anything that is not clear or if you would like more information. Take time to decide whether or not you wish to take part. Thank you for reading this.

#### Purpose of the study and procedure

The aim of this experiment is to evaluate the performance of a new software plugin for the image-based cell profiling tool CellProfiler, which was built in order to reduce the time spent on manually adjusting settings manually for enhanced quality of object segmentation, while still retaining the customization capability of CellProfiler.

For this purpose, we designed an experimental study to record the performance of the new software.

At the start of the experiment, you will be provided a test computer running CellProfiler and a sample pipeline for image analysis. You will then have to modify the standard CellProfiler pipeline to evaluate a randomly assigned dataset by completing specific tasks (handed to you on a separate sheet) that uses different functionalities of the plugin. The times for completing the tasks will be recorded, along with the quality of the solution as indicated by you.

Using a questionnaire after completing the tasks, you will then evaluate the different modules.

The session will last up to 150 minutes but will include several breaks and periods where you will simply observe the system operation. Your participation is rewarded by £10 and the possibility to gain an additional £50 in a random draw among all participants.

#### Why would your participation be useful?

Rapid and high throughput techniques in microscopy are still burdened by the need for human-based intervention, thereby hindering important processes such as cell profiling for drug response. There is a need to improve the efficiency of biological image processing and analysis by using computational tools (e.g. machine learning, computer vision). However, such techniques are relatively new and must be informed by experimentation. The data you provide in the experiment will thus be used to

University of Glasgow | College of Science & Engineering

### Participant Consent Form

**Project title:** Interactive machine learning for accelerated analysis of microscopy images.

**Investigators**: Lisa Laux and Marie FA Cutiongco under the supervision of Dr Bjørn Sand Jensen

[ ] I confirm that I have read and understood the *Participant Information Sheet* for the above study and have had the opportunity to ask questions.

[ ] I understand that my participation is voluntary and that I am free to withdraw at any time, without giving any reason, that I may decline to answer any questions, and that this does not affect the compensation.

[ ] I acknowledge that participants will be referred to by pseudonym. I acknowledge that there will be no effects on my situation arising from my participation or non-participation in this research.

All names and other material likely to identify individuals will be anonymised. The material will always be treated as confidential and kept in secure storage. The material may be used in future publications, both print and online.

[ ] I agree to waive my copyright to any data collected as part of this project

[ ] I acknowledge that I have received £10 and the opportunity to receive an additional £50 for my participation in this study.

[ ] I wish to be informed about the outcome of the study via email.

**[ ] I agree to take part in the study.**

**Participant:**

Full name: ________________________________

Email (optional): ________________________________

Signature: ____________________________ Date: ______________

**Investigator:**

Full name: ________________________________

E-mail: ________________________________

Office address: ________________________________

Signature: ____________________________ Date: ______________

**Supervisor:**
Bjørn Sand Jensen, PhD, , +44 0141 330 1639, School of Computing Science, 18 Lilybank Gardens , University of Glasgow

accelerate microscopy analysis via machine learning and verify that such an approach is beneficial for speeding up analysis and understanding cell behaviour.

#### Do you have to take part?

Taking part in this research is entirely voluntary: it is up to you to decide whether or not to take part. If you do decide to take part, you will be given this information sheet to keep and be asked to sign a consent form. Note that even if you decide to take part you are still free to withdraw at any time and without giving a reason.

#### Possible Risks

The use and interaction with the software can be tiring after a while. Several breaks will be offered and can be provided anytime you feel the need. Note that you also have the right to withdraw from the study at any point without having to provide a reason.

#### Inclusion and Exclusion criteria

You are eligible to take part if you have experience with microscopy analysis using computational tools, are 18 years of age or older, able to provide consent and have no mental or physical disabilities.

#### Publication of Results

Results of the study may be submitted for publication, but your identity will be strictly confidential. You can request now to be notified of the publication.

#### Who has reviewed this study?

This study has been reviewed and approved by the Ethics Committee of the College of Science and Engineering at the University of Glasgow.

#### Contacts:

For further information about this study, please contact:

**Lisa Laux**


**Marie F.A. Cutiongco**


Supervisor:

**Bjørn Sand Jensen**

+44 0141 330 1639
School of Computing Science
18 Lilybank Gardens
University of Glasgow

### Debrief
### CellProfiler plugin user evaluation

The aim of this experiment was to evaluate the performance of new plugins for the image-based cell profiling tool CellProfiler. These plugins were built in order to increase the efficiency of obtaining the parameters for optimal object segmentation and enhance the maximum quality of object segmentation.

We were particularly interested on your opinion about the usability of the plugins and your impression regarding its impact on the time required for adjusting pipelines. The surveys provided us with information regarding the ease of use and perceived usefulness of the plugins for different tasks. We were also interested on the impact of the plugins on the quality of object segmentation. During the experiment, we presented you with the output of the processing of image sets using the pipelines from all 5 tasks to determine segmentation quality.

Do you have any comments or questions about the experiment? Please take a note of our email address:


Marie Cutiongco,

Lisa Laux,


You can also contact the supervisor for this project:


Bjørn Jensen,


Please let us know if you have any further questions about this experiment. Thank you for your help.
