## Supplementary material for "Interactive machine learning for fast and robust cell profiling": All supplementary files: TaskSheet.pdf

University of Glasgow | College of Science & Engineering

### Participant Instruction Sheet

**Project title**: Interactive machine learning for accelerated analysis of microscopy images.

**Investigators**: Lisa Laux and Marie FA Cutiongco under the supervision of Bjørn Sand Jensen.

#### Contacts:

For further information about this study, please contact:

**Lisa Laux**


**Marie Cutiongco**


Supervisor:

**Bjørn Sand Jensen**

+44 0141 330 1639
School of Computing Science
18 Lilybank Gardens
University of Glasgow

### Introduction: CellProfiler

CellProfiler is an open source software created for rapid and high-throughput analysis of images for biological purposes. CellProfiler has a modular configuration that allows bespoke pipelines to be made to perform the standard tasks of image pre-processing, object detection and segmentation, feature extraction from objects and object classification. The modular quality of CellProfiler is an advantageous tool that allows many different types of datasets to be analysed by one platform. However, the modular approach means that there is a large number of parameters that need to be manually tuned to obtain high quality object segmentation. In some cases and with high complexity of objects, high quality of segmentation cannot be achieved in a reasonable timescale.

### Tasks completion sheet: Evaluation of CellProfiler plugins

Please follow the instructions given on the following pages to complete the tasks of the experiment.

Your tasks will involve optimizing and running the results of a CellProfiler pipeline on one image set using a new plugin. After each task, you will be asked to evaluate the quality of object segmentation resulting from the use of the resulting pipeline on a new image set. Furthermore, you will be asked questions regarding the ease of the task performed. After completing all 5 tasks, you will be presented with a questionnaire (and some output images) to evaluate the performance of the plugins.

<u>The objective of today's experiment and a short overview of CellProfiler will be explained to you now.</u>

#### General note: please intermittently save your pipelines

##### Task 1: Manually adjust pipeline settings to enhance object segmentation

*Your task is to manually adjust the parameters of a given CellProfiler pipeline to optimise the object segmentation of nuclei, cytoskeleton and adhesions.*

*You have a maximum of* ***20 minutes*** *for this task.*

1. Drag and drop “UserOptimized_Task1.cpproj” into the left hand side of the CellProfiler window.
2. Go into test mode by clicking *Start Test Mode* at the bottom left hand.
3. Enhance the pipeline output (which will be displayed during and after running the pipeline on an image set) by adjusting the setting parameters of the pipeline.

    a. Only modules that have the open eye icon 👁 can be adjusted.
    b. Only parameter settings that contain integer or floating-point values (no range of values) within the module can be changed.
    c. If in doubt about what the specific parameter in a module indicates, please click on the ? button at the right-hand side of the screen.
    d. You are allowed to run the given pipeline on any of the 5 image sets available. Please choose an image set by clicking on *Test* → *Choose image set* found at the top part

**BREAK: 5 mins**

**Task 1: Evaluation of images resulting from a manually configured pipeline**

The pipeline you have just optimized will be used to segment objects within new images. These segmented objects will be shown to you. Rate the quality of the object segmentation for each image set by choosing an appropriate rating between 1 and 10 in the table below. A rating of 1 indicates poor object segmentation and 10 indicates excellent object segmentation.

| **Image set** | **Object segmentation rating**<br>Poor | | | | | | | | | Excellent |
|---|---|---|---|---|---|---|---|---|---|---|
| A | 1 | 2 | 3 | 4 | 5 | 6 | 7 | 8 | 9 | 10 |
| B | 1 | 2 | 3 | 4 | 5 | 6 | 7 | 8 | 9 | 10 |
| C | 1 | 2 | 3 | 4 | 5 | 6 | 7 | 8 | 9 | 10 |
| D | 1 | 2 | 3 | 4 | 5 | 6 | 7 | 8 | 9 | 10 |
| E | 1 | 2 | 3 | 4 | 5 | 6 | 7 | 8 | 9 | 10 |
| F | 1 | 2 | 3 | 4 | 5 | 6 | 7 | 8 | 9 | 10 |
| G | 1 | 2 | 3 | 4 | 5 | 6 | 7 | 8 | 9 | 10 |
| H | 1 | 2 | 3 | 4 | 5 | 6 | 7 | 8 | 9 | 10 |
| I | 1 | 2 | 3 | 4 | 5 | 6 | 7 | 8 | 9 | 10 |
| J | 1 | 2 | 3 | 4 | 5 | 6 | 7 | 8 | 9 | 10 |

**SURVEY: Circle the appropriate response to questions related to Task 1**

| Question | Answers | | | | |
|---|---|---|---|---|---|
| Changing the parameters of the pipeline to achieve optimal object segmentation is easy. | Strongly disagree | Disagree | Neutral | Agree | Strongly agree |
| The amount of time provided was sufficient to configure the pipeline for optimal object segmentation. | Strongly disagree | Disagree | Neutral | Agree | Strongly agree |
| Object segmentation is easy and straightforward. | Strongly disagree | Disagree | Neutral | Agree | Strongly agree |
| I am satisfied with the performance of my pipeline | Strongly disagree | Disagree | Neutral | Agree | Strongly agree |
| I have explored all possibilities in pipeline configuration within the given timeframe | Strongly disagree | Disagree | Neutral | Agree | Strongly agree |

**Task 2: Implement a plugin for automated parameter configuration and segmentation evaluation**

*Your task is to use the new plugins* AutomatedEvaluation *and* BayesianOptimisation*. Take a look at them and read through the module description. Please note that there is a known bug in the way the pipeline executes. It will eventually stop running at some point and not execute all available modules in the pipeline. If this happens during any time of your task execution, please simply click on the* Run *button again. The data already gathered will be persisted automatically.*

*You have a maximum of **20 minutes** for this task.*

1. Drag and drop the file named "Automatic_Task2.cpproj" into the left hand side of the CellProfiler window.
2. Select the module *AutomatedEvaluation* and indicate the measurement range of the object that indicates successful segmentation. Example measurements are:

    a. Minimum and maximum expected area of the object
    b. Minimum and maximum expected eccentricity (ellipticity) of the object
    c. Minimum and maximum expected number of objects in one image

3. Click *Start Test Mode* → *Run* found at the bottom left hand side
4. A pop-up window will be displayed when the pipeline reaches the *AutomatedEvaluation* module. The *AutomatedEvaluation* modules will alter other module parameters to reach the desired values of the object as defined in Step 2.

    You are encouraged to let the *AutomatedEvaluation* module run for at least 5 iterations/rounds before adjusting any settings, as described below:

    If you are not satisfied with the optimized segmentation:

    - change the image set by clicking *Next Image Set* found at the bottom left hand side;
    - or change the parameters defined in Step 2 (as suggested by the *AutomatedEvaluation* module at the end of the current optimisation). Before resuming run, remove all previous data by going to *BayesianOptimisation* module → *Delete previous Data* at the bottom of the page.

    However, if you are satisfied with the current optimisation or, want to use the pipeline on the next image set you can simply move on to the next image set. Interrupt the pipeline run in this case and choose a new image set by clicking *Stop* → *Next Image Set*.

**BREAK: 5 mins**

**Task 2: Evaluation of images resulting from an automatically configured pipeline**

The pipeline will be used to segment objects within new images, which will be shown to you. Rate the quality of the object segmentation for each image set by choosing an appropriate rating between 1 and 10 in the table below. A rating of 1 indicates poor object segmentation and 10 indicates excellent object segmentation.

| Image set | Object segmentation rating<br>Poor | | | | | | | | | Excellent |
|---|---|---|---|---|---|---|---|---|---|---|
| A | 1 | 2 | 3 | 4 | 5 | 6 | 7 | 8 | 9 | 10 |
| B | 1 | 2 | 3 | 4 | 5 | 6 | 7 | 8 | 9 | 10 |
| C | 1 | 2 | 3 | 4 | 5 | 6 | 7 | 8 | 9 | 10 |
| D | 1 | 2 | 3 | 4 | 5 | 6 | 7 | 8 | 9 | 10 |
| E | 1 | 2 | 3 | 4 | 5 | 6 | 7 | 8 | 9 | 10 |
| F | 1 | 2 | 3 | 4 | 5 | 6 | 7 | 8 | 9 | 10 |
| G | 1 | 2 | 3 | 4 | 5 | 6 | 7 | 8 | 9 | 10 |
| H | 1 | 2 | 3 | 4 | 5 | 6 | 7 | 8 | 9 | 10 |
| I | 1 | 2 | 3 | 4 | 5 | 6 | 7 | 8 | 9 | 10 |
| J | 1 | 2 | 3 | 4 | 5 | 6 | 7 | 8 | 9 | 10 |

**SURVEY: Circle the appropriate response to questions related to Task 2**

| Question | Answers | | | | |
|---|---|---|---|---|---|
| The automated configuration of the pipeline parameters is easy to implement. | Strongly disagree | Disagree | Neutral | Agree | Strongly agree |
| Setting measurement ranges that denote optimum object segmentation is easy and straightforward. | Strongly disagree | Disagree | Neutral | Agree | Strongly agree |
| The evaluation of object segmentation is easy and straightforward. | Strongly disagree | Disagree | Neutral | Agree | Strongly agree |
| I observed improvement in object segmentation with the implementation of the new plugins compared to my pipeline | Strongly disagree | Disagree | Neutral | Agree | Strongly agree |
| The amount of time provided was sufficient to configure the pipeline for optimal object segmentation. | Strongly disagree | Disagree | Neutral | Agree | Strongly agree |
| I am satisfied with the performance of the automatically configured pipeline | Strongly disagree | Disagree | Neutral | Agree | Strongly agree |

**Task 3: Implement a plugin for automated parameter configuration with user-based feedback on segmentation quality**

*Your task is to use the new plugins* ManualEvaluation *and* BayesianOptimisation. *These modules will be placed after the module which identifies object of interest. Take a look at them and read through the module description. Please note that there is a known bug in the way the pipeline executes. It will eventually stop running at some point and not execute all available modules in the pipeline. If this happens during any time of your task execution, please simply click on the* Run *button again. The data already gathered will be persisted automatically.*

*You have a maximum of **20 minutes** for this task.*

1. Drag and drop "Manual_Task3.cpproj" into the left hand side of the CellProfiler window.
2. Go to the module *ManualEvaluation* and indicate your threshold for segmentation quality. A threshold of 1 indicates low threshold for object segmentation and 10 indicates high threshold for object segmentation.
3. Click *Start Test Mode* → *Run* found at the bottom left hand side
4. A pop-up window will be displayed when the pipeline reaches the *ManualEvaluation* module. Rate the quality of the object segmentation for each image set by choosing an appropriate rating between 1 and 10. A rating of 1 indicates poor object segmentation and 10 indicates excellent object segmentation.

   The *BayesianOptimisation* plugin will continue running until segmentation quality threshold is reached.

   If you feel that your optimisation never reaches the required quality, or want to use the pipeline on the next image set you can simply move on to the next image set. Interrupt the pipeline run in this case and choose a new image set by clicking *Stop* → *Next Image Set*

5. Repeat step 3 for the 5 image sets available. It is advisable to train the pipeline on more than one image set.

#### BREAK: 5 mins

#### Task 3: Evaluation of images resulting from an automatically configured pipeline with user-based feedback

The pipeline will be used to segment objects within new images, which will be shown to you. Rate the quality of the object segmentation for each image set by choosing an appropriate rating between 1 and 10 in the table below. A rating of 1 indicates poor object segmentation and 10 indicates excellent object segmentation.

| **Image set** | **Object segmentation rating**<br>Poor | | | | | | | | | Excellent |
|---|---|---|---|---|---|---|---|---|---|---|
| A | 1 | 2 | 3 | 4 | 5 | 6 | 7 | 8 | 9 | 10 |
| B | 1 | 2 | 3 | 4 | 5 | 6 | 7 | 8 | 9 | 10 |
| C | 1 | 2 | 3 | 4 | 5 | 6 | 7 | 8 | 9 | 10 |
| D | 1 | 2 | 3 | 4 | 5 | 6 | 7 | 8 | 9 | 10 |
| E | 1 | 2 | 3 | 4 | 5 | 6 | 7 | 8 | 9 | 10 |
| F | 1 | 2 | 3 | 4 | 5 | 6 | 7 | 8 | 9 | 10 |
| G | 1 | 2 | 3 | 4 | 5 | 6 | 7 | 8 | 9 | 10 |
| H | 1 | 2 | 3 | 4 | 5 | 6 | 7 | 8 | 9 | 10 |
| I | 1 | 2 | 3 | 4 | 5 | 6 | 7 | 8 | 9 | 10 |
| J | 1 | 2 | 3 | 4 | 5 | 6 | 7 | 8 | 9 | 10 |

**SURVEY: Circle the appropriate response to questions related to Task 3**

| Question | Answers | | | | |
|---|---|---|---|---|---|
| Giving feedback on object segmentation is easy and straightforward | Strongly disagree | Disagree | Neutral | Agree | Strongly agree |
| I observed improvement in object segmentation with the implementation of the new plugins compared to my pipeline | Strongly disagree | Disagree | Neutral | Agree | Strongly agree |
| I am satisfied with the performance of a pipeline relying on user feedback | Strongly disagree | Disagree | Neutral | Agree | Strongly agree |
| The amount of time provided was sufficient to configure the pipeline for optimal object segmentation. | Strongly disagree | Disagree | Neutral | Agree | Strongly agree |

**Task 4: Implement a plugin for automated parameter configuration with both automated and user-based evaluation of segmentation quality**

*Your task is to use the new plugins* AutomatedEvaluation*,* ManualEvaluation *and* BayesianOptimisation *simultaneously.*

*You have a maximum of **20 minutes** for this task.*

1. Drag and drop the file named "Semiauto_Task5.cpproj" into the left hand side of the CellProfiler window.
2. Go to the module *ManualEvaluation* and indicate your threshold for segmentation quality. A threshold of 1 indicates low threshold for object segmentation and 10 indicates high threshold for object segmentation.
3. Go to the module *AutomatedEvaluation* and indicate the measurements of the object and the range that indicates successful segmentation.
    - Minimum and maximum expected area of the object
    - Minimum and maximum expected eccentricity (ellipticity) of the object
    - Minimum and maximum expected number of objects in one image
4. Click *Start Test Mode* → *Run* found at the bottom left hand side
5. Pop-up windows will be displayed when the pipeline reaches the *ManualEvaluation* and *AutomatedEvaluation* modules.
   For the *ManualEvaluation* pop-up window, rate the quality of the object segmentation for each image set by choosing an appropriate rating between 1 and 10. A rating of 1 indicates poor object segmentation and 10 indicates excellent object segmentation.

   You are encouraged to let the AutomatedEvaluation module run for at least 5 iterations/rounds before adjusting any settings. If you are not satisfied with the segmentation shown in the *AutomatedEvaluation* module.

   - change the image set by clicking *Next Image Set* found at the bottom left hand side;
   - or change the parameters defined in Step 2 (as suggested by the AutomatedEvaluation module at the end of the current optimisation). Before resuming run, remove all previous data by going to *BayesianOptimisation* module → *Delete previous Data* at the bottom of the page.

   The *BayesianOptimisation* plugin will continue running to automatically optimise the parameters of the pipeline until segmentation quality threshold is reached.
6. Repeat step 3 for the 5 image sets available. It is advisable to train the pipeline on more than one image set.

**BREAK: 5 mins**

**Task 4: Evaluation of images resulting from an automatically configured pipeline**

The pipeline will be used to segment objects within new images, which will be shown to you. Rate the quality of the object segmentation for each image set by choosing an appropriate rating between 1 and 10 in the table below. A rating of 1 indicates poor object segmentation and 10 indicates excellent object segmentation.

| Image set | Object segmentation rating<br>Poor | | | | | | | | | Excellent |
|---|---|---|---|---|---|---|---|---|---|---|
| A | 1 | 2 | 3 | 4 | 5 | 6 | 7 | 8 | 9 | 10 |
| B | 1 | 2 | 3 | 4 | 5 | 6 | 7 | 8 | 9 | 10 |
| C | 1 | 2 | 3 | 4 | 5 | 6 | 7 | 8 | 9 | 10 |
| D | 1 | 2 | 3 | 4 | 5 | 6 | 7 | 8 | 9 | 10 |
| E | 1 | 2 | 3 | 4 | 5 | 6 | 7 | 8 | 9 | 10 |
| F | 1 | 2 | 3 | 4 | 5 | 6 | 7 | 8 | 9 | 10 |
| G | 1 | 2 | 3 | 4 | 5 | 6 | 7 | 8 | 9 | 10 |
| H | 1 | 2 | 3 | 4 | 5 | 6 | 7 | 8 | 9 | 10 |
| I | 1 | 2 | 3 | 4 | 5 | 6 | 7 | 8 | 9 | 10 |
| J | 1 | 2 | 3 | 4 | 5 | 6 | 7 | 8 | 9 | 10 |

**SURVEY: Circle the appropriate response to questions related to Task 4**

| Question | Answers | | | | |
|---|---|---|---|---|---|
| The automated configuration of the pipeline parameters is easy to implement. | Strongly disagree | Disagree | Neutral | Agree | Strongly agree |
| Setting measurement ranges that denote optimum object segmentation is easy and straightforward. | Strongly disagree | Disagree | Neutral | Agree | Strongly agree |
| Giving feedback on object segmentation is easy and straightforward. | Strongly disagree | Disagree | Neutral | Agree | Strongly agree |
| I observed improvement in object segmentation with the implementation of the new plugins compared to my pipeline | Strongly disagree | Disagree | Neutral | Agree | Strongly agree |
| The combination of the Automated and Manual Evaluation modules with the BayesianOptimisation module helped to enhance the segmentation quality | Strongly disagree | Disagree | Neutral | Agree | Strongly agree |
| The amount of time provided was sufficient to configure the pipeline for optimal object segmentation. | Strongly disagree | Disagree | Neutral | Agree | Strongly agree |

**Task 5: Evaluation of images resulting from a fixed pipeline**

Open the folder “Results_FixedPipeline”, where the image set resulting from a pipeline configured by a CellProfiler expert is provided.

Rate the quality of the object segmentation for each image set by choosing an appropriate rating between 1 and 10 in the table below. A rating of 1 indicates poor object segmentation and 10 indicates excellent object segmentation.

| Image set | Object segmentation rating | | | | | | | | | |
|---|---|---|---|---|---|---|---|---|---|---|
| | Poor | | | | | | | | | Excellent |
| A | 1 | 2 | 3 | 4 | 5 | 6 | 7 | 8 | 9 | 10 |
| B | 1 | 2 | 3 | 4 | 5 | 6 | 7 | 8 | 9 | 10 |
| C | 1 | 2 | 3 | 4 | 5 | 6 | 7 | 8 | 9 | 10 |
| D | 1 | 2 | 3 | 4 | 5 | 6 | 7 | 8 | 9 | 10 |
| E | 1 | 2 | 3 | 4 | 5 | 6 | 7 | 8 | 9 | 10 |
| F | 1 | 2 | 3 | 4 | 5 | 6 | 7 | 8 | 9 | 10 |
| G | 1 | 2 | 3 | 4 | 5 | 6 | 7 | 8 | 9 | 10 |
| H | 1 | 2 | 3 | 4 | 5 | 6 | 7 | 8 | 9 | 10 |
| I | 1 | 2 | 3 | 4 | 5 | 6 | 7 | 8 | 9 | 10 |
| J | 1 | 2 | 3 | 4 | 5 | 6 | 7 | 8 | 9 | 10 |

**SURVEY: Circle the appropriate response to questions related to Task 1**

| Question | Answers | | | | |
|---|---|---|---|---|---|
| The evaluation of object segmentation is easy and straightforward | Strongly disagree | Disagree | Neutral | Agree | Strongly agree |
| I am satisfied with the object segmentation presented to me | Strongly disagree | Disagree | Neutral | Agree | Strongly agree |

**Thank you! You are now done with the tasks**

**and can continue filling out the questionnaire in the next page.**

### Questionnaire: Evaluation of CellProfiler plugins

Please answer the following questions by circling one of the answers per question.

| Question | Answers | | | | |
|---|---|---|---|---|---|
| Compared to manually adjusting parameters, the use of the new plugins for automatic parameter testing and successful object segmentation is ... | much more time-consuming | more time consuming | takes equal time | less time-consuming | much less time-consuming |
| Understanding the modules with the information provided in the module (and help windows) was ... | Very difficult | Difficult | Neutral | Easy | Very easy |
| Overall, using the new modules to adjust pipeline settings was ... | Very complex and/or very irritating | Complex and/or irritating | Neutral | Easy | Very easy |
| I would regularly use the new plugins to automatically optimise my image segmentation pipeline | Strongly disagree | Disagree | Neutral | Agree | Strongly agree |
| Which module do you prefer to implement to automatically optimise your image segmentation pipeline? | *AutomaticEvaluation* | *ManualEvaluation* | Both | None | |

| Do you have any comments or remarks on the plugins? Do you think there is functionality missing? Do you think they are too complex? What would you change? |
|---|

### Questionnaire: About yourself

| Question | Answers | | | | | |
|---|---|---|---|---|---|---|
| How often do you perform biological image analysis (on any software/platform)? | Never | Rarely | Sometimes | Frequently | Often/ Regularly | Prefer not to say |
| How often do you use CellProfiler? | Never | Rarely | Sometimes | Frequently | Often/ Regularly | Prefer not to say |
| How old are you? | 18-25 | 26-33 | 34-41 | 42-48 | >48 | Prefer not to say |
| What is your occupation at the University of Glasgow? | Undergraduate student | Masters student | PhD Student | Postdoctoral researcher | Scientist/ Researcher | Prefer not to say |

**Thank you for answering these questions. You are now finished with the experiment.**
