## Supplementary figures and images for "Interactive machine learning for fast and robust cell profiling"

### A_groundtruth.jpg

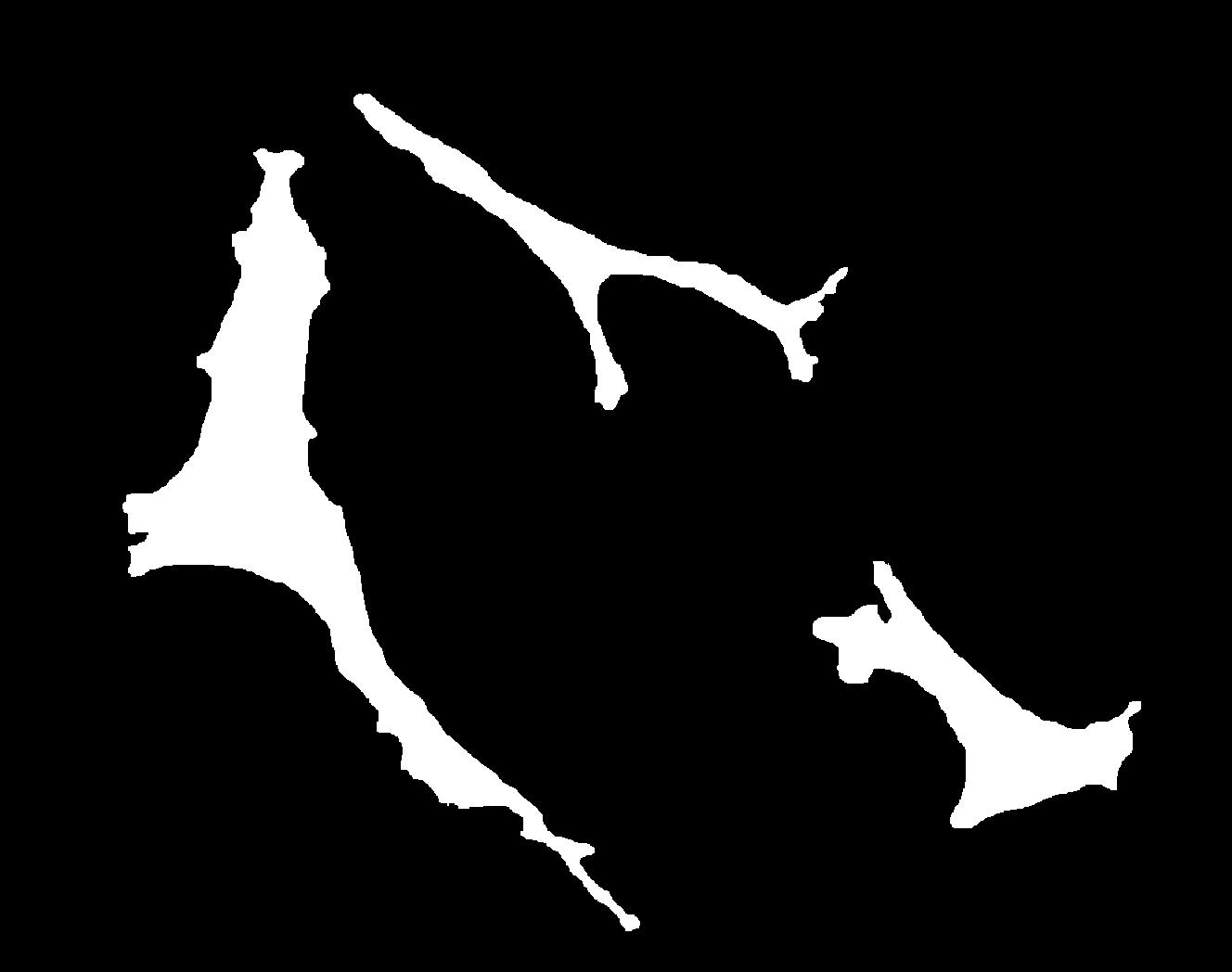

### Automated_Participant1_mask_cell_A.jpg

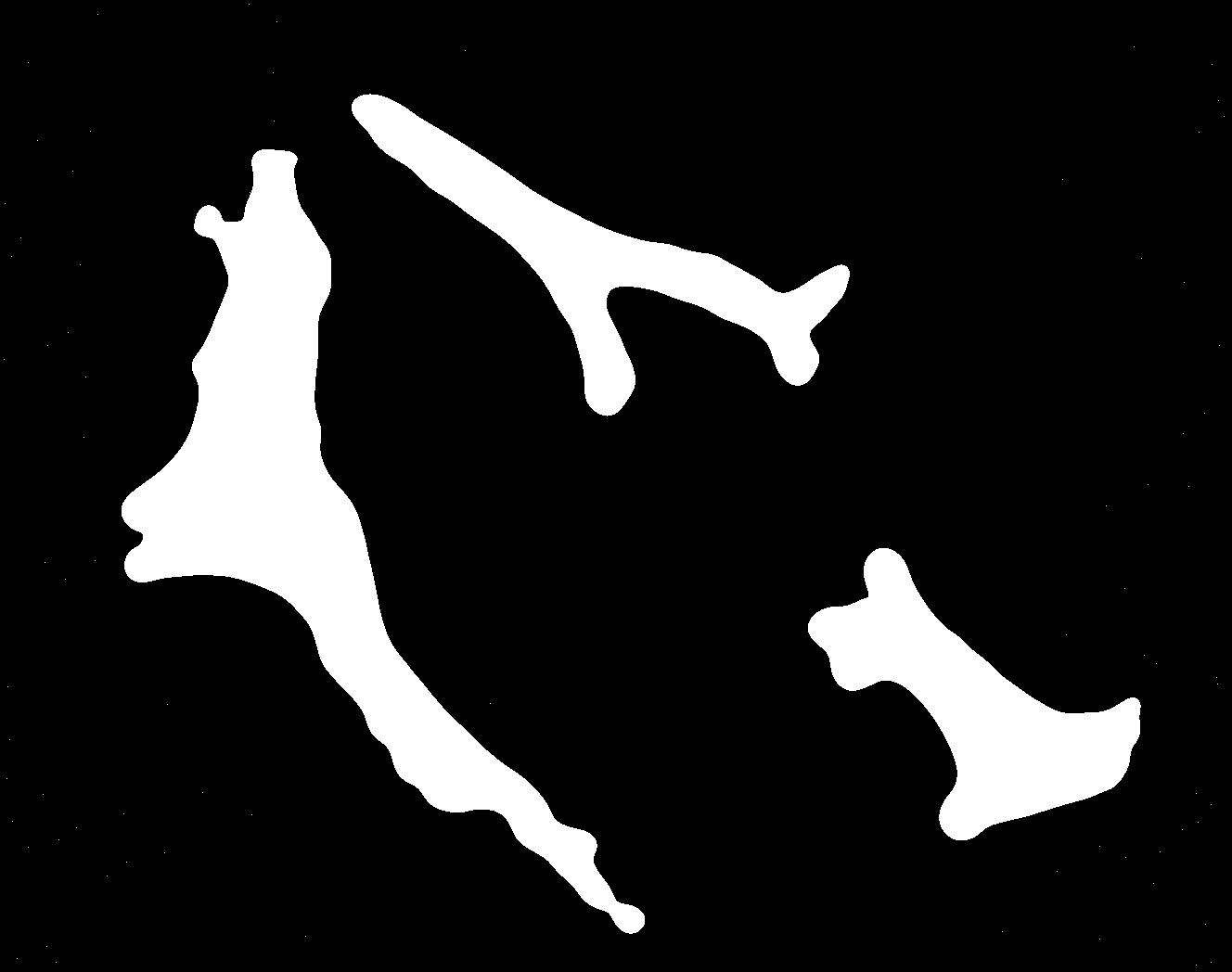

### Automated_Participant1_mask_cell_B.jpg

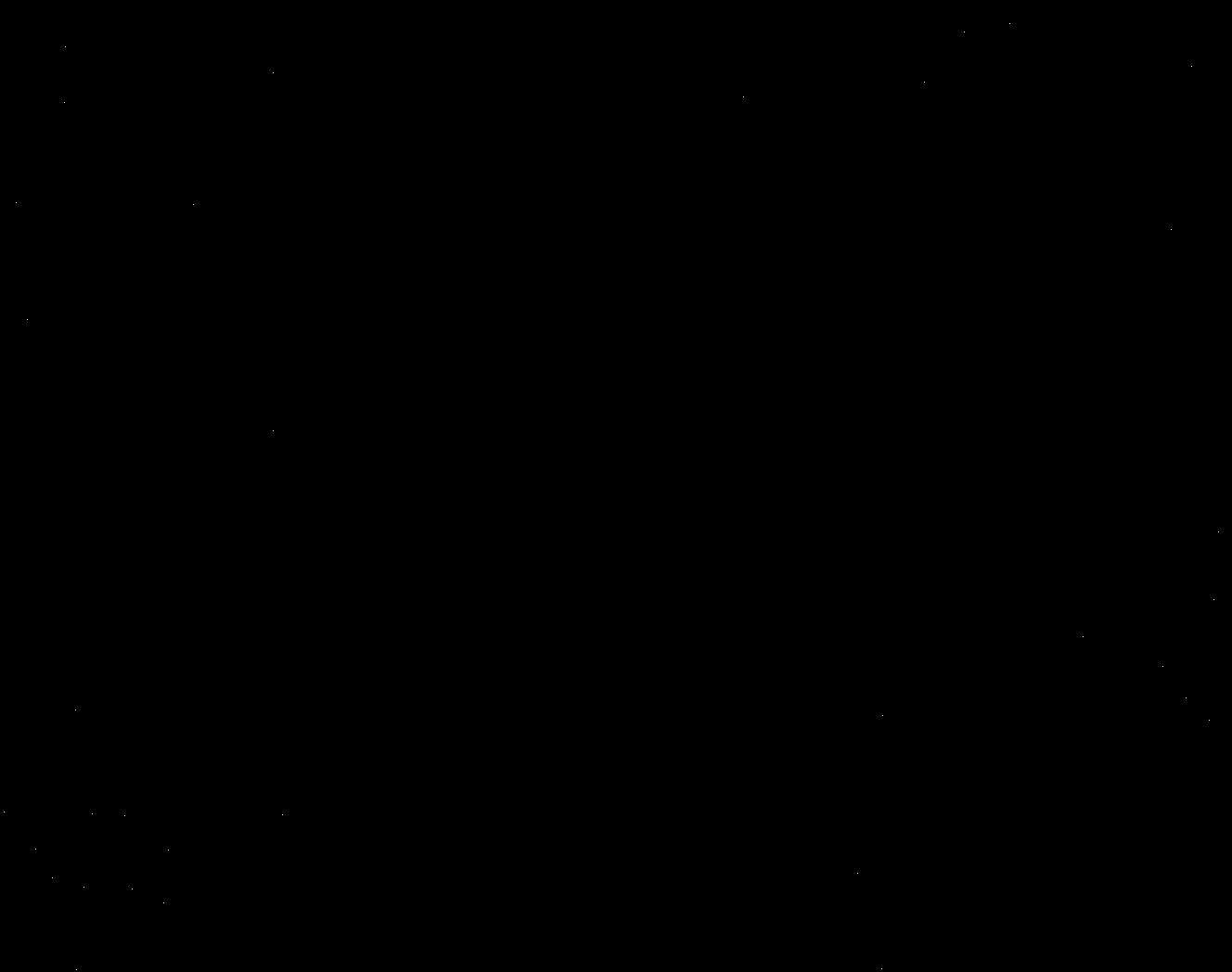

### Automated_Participant1_mask_cell_C.jpg

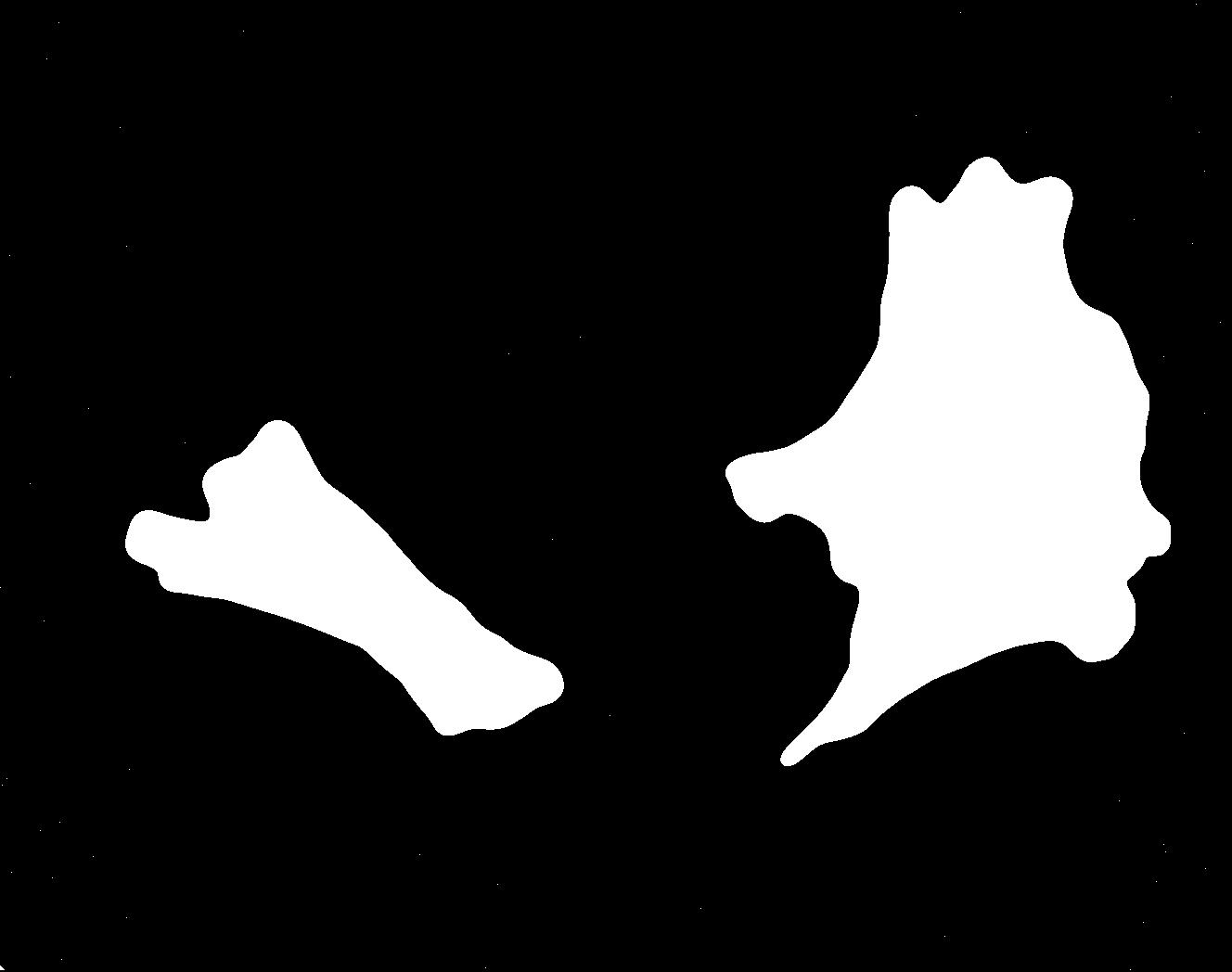

### Automated_Participant1_mask_cell_D.jpg

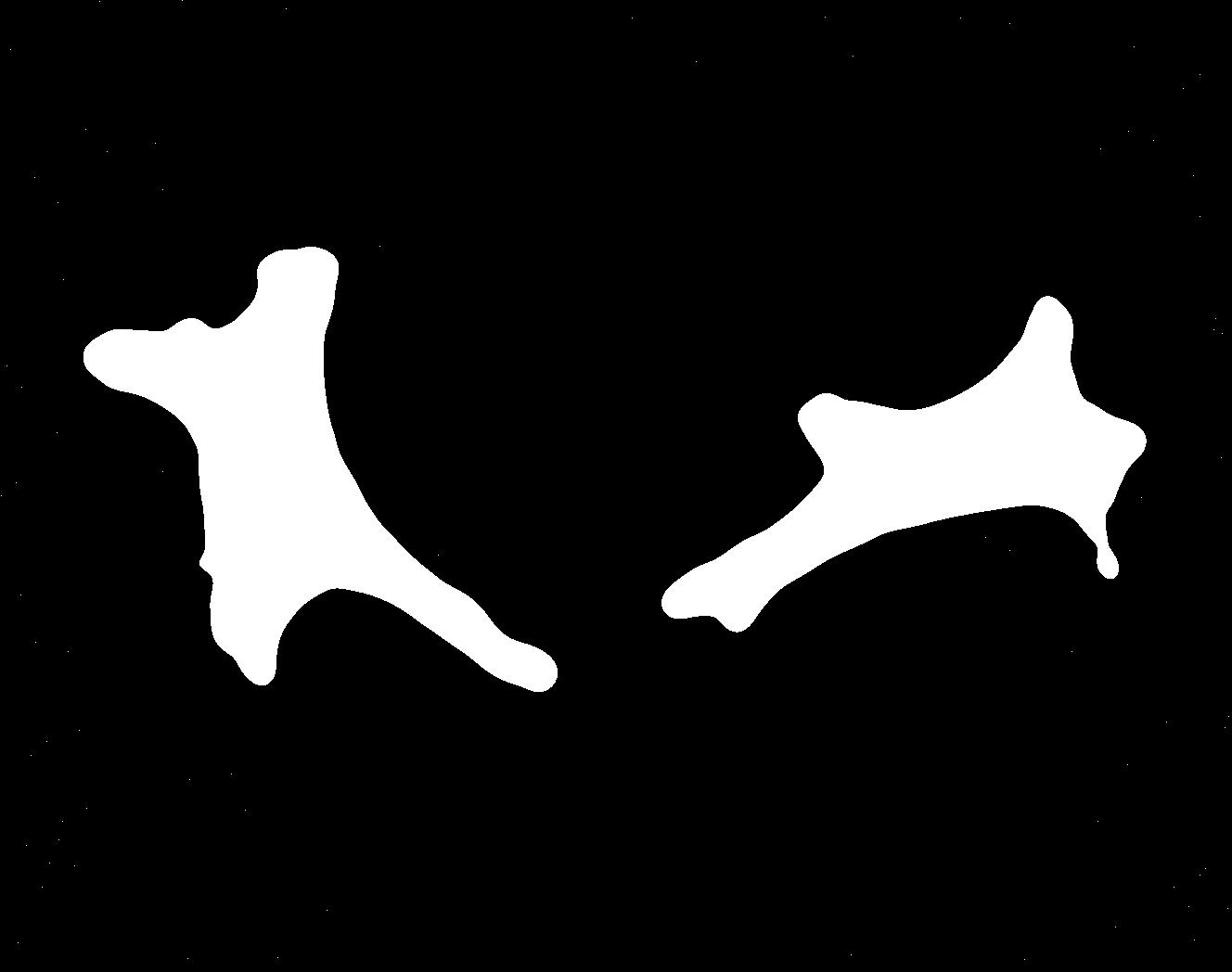

### B_groundtruth.jpg

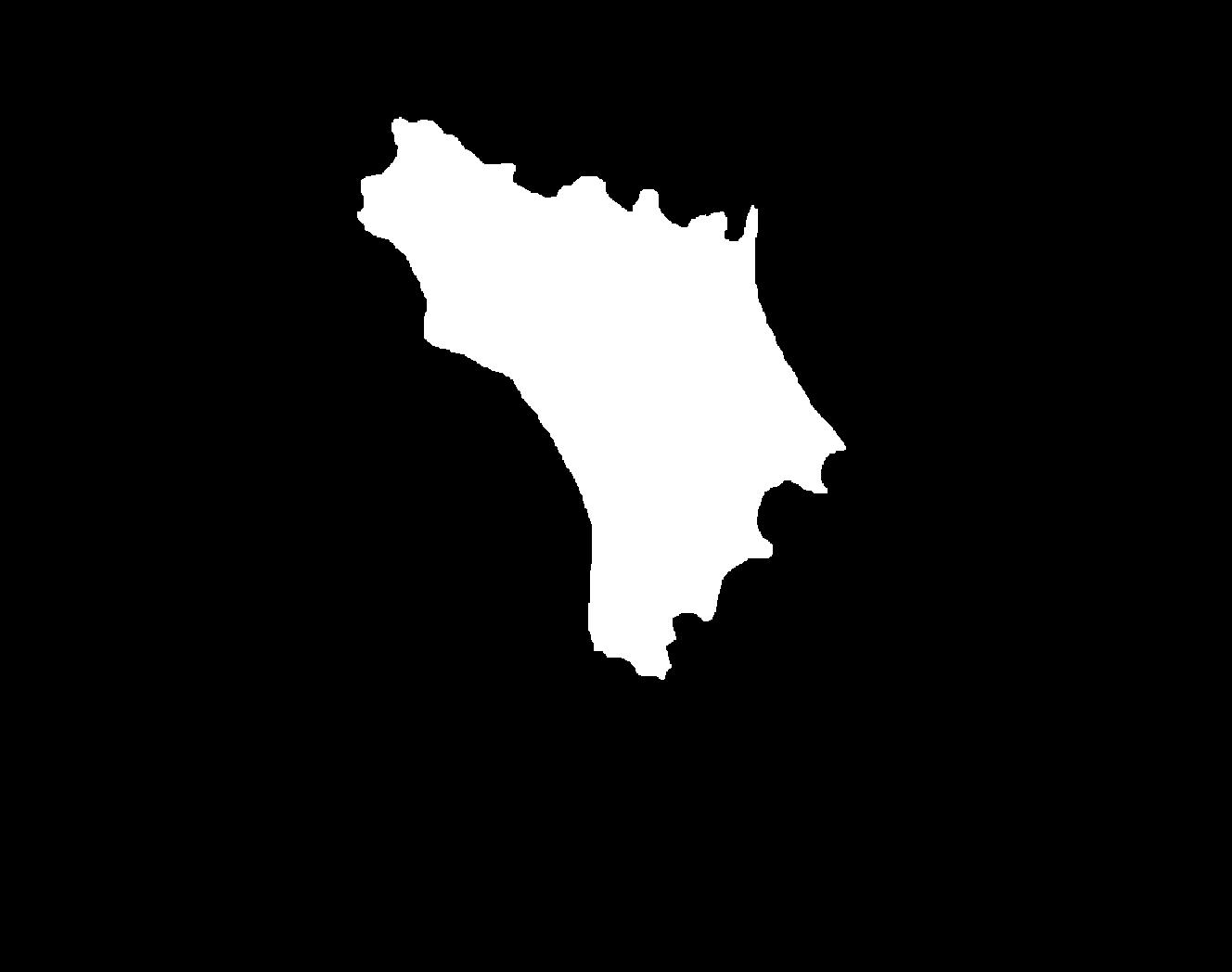

### C_groundtruth.jpg

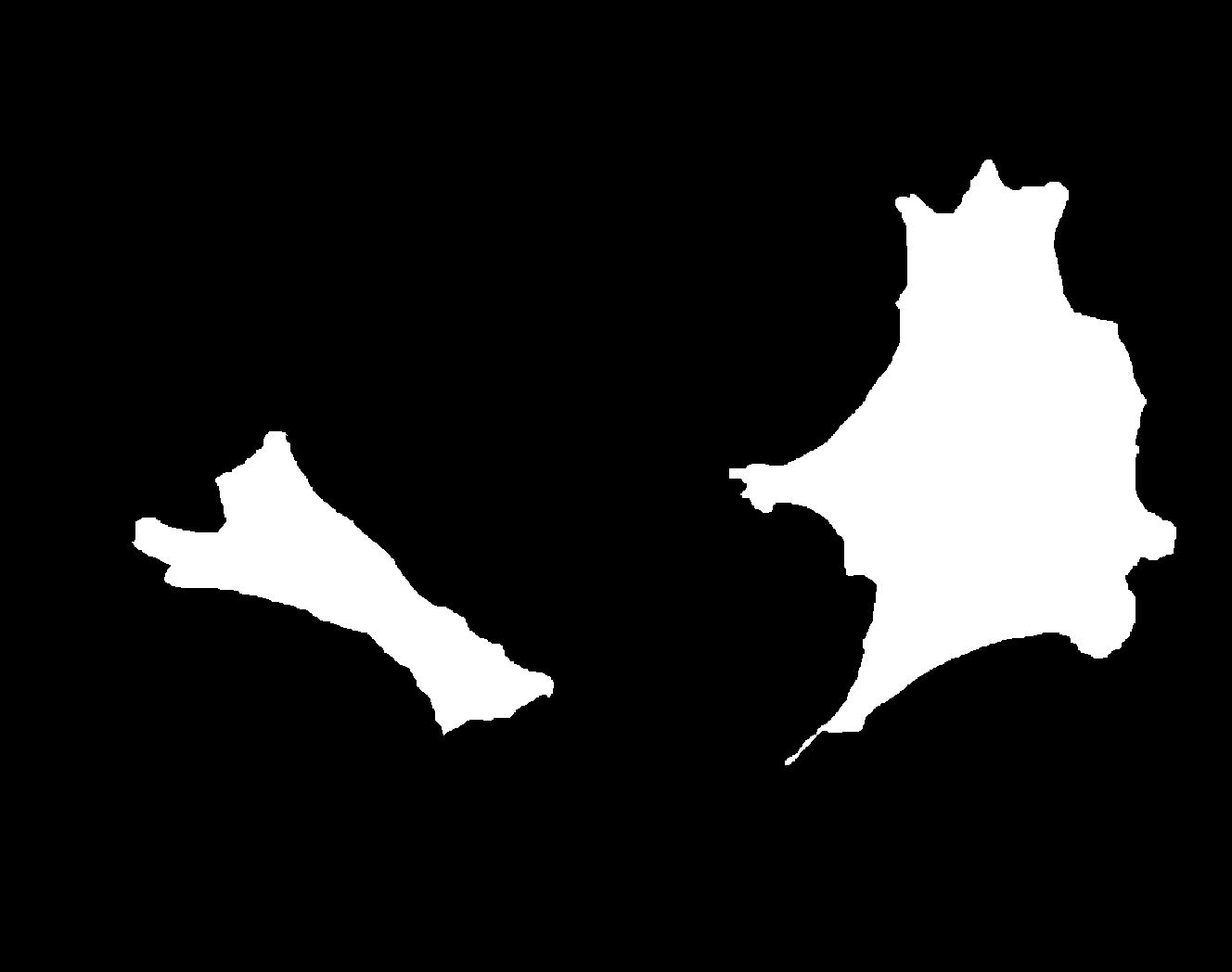

### C_Plate_R_p00_0_A01f17d0.TIF

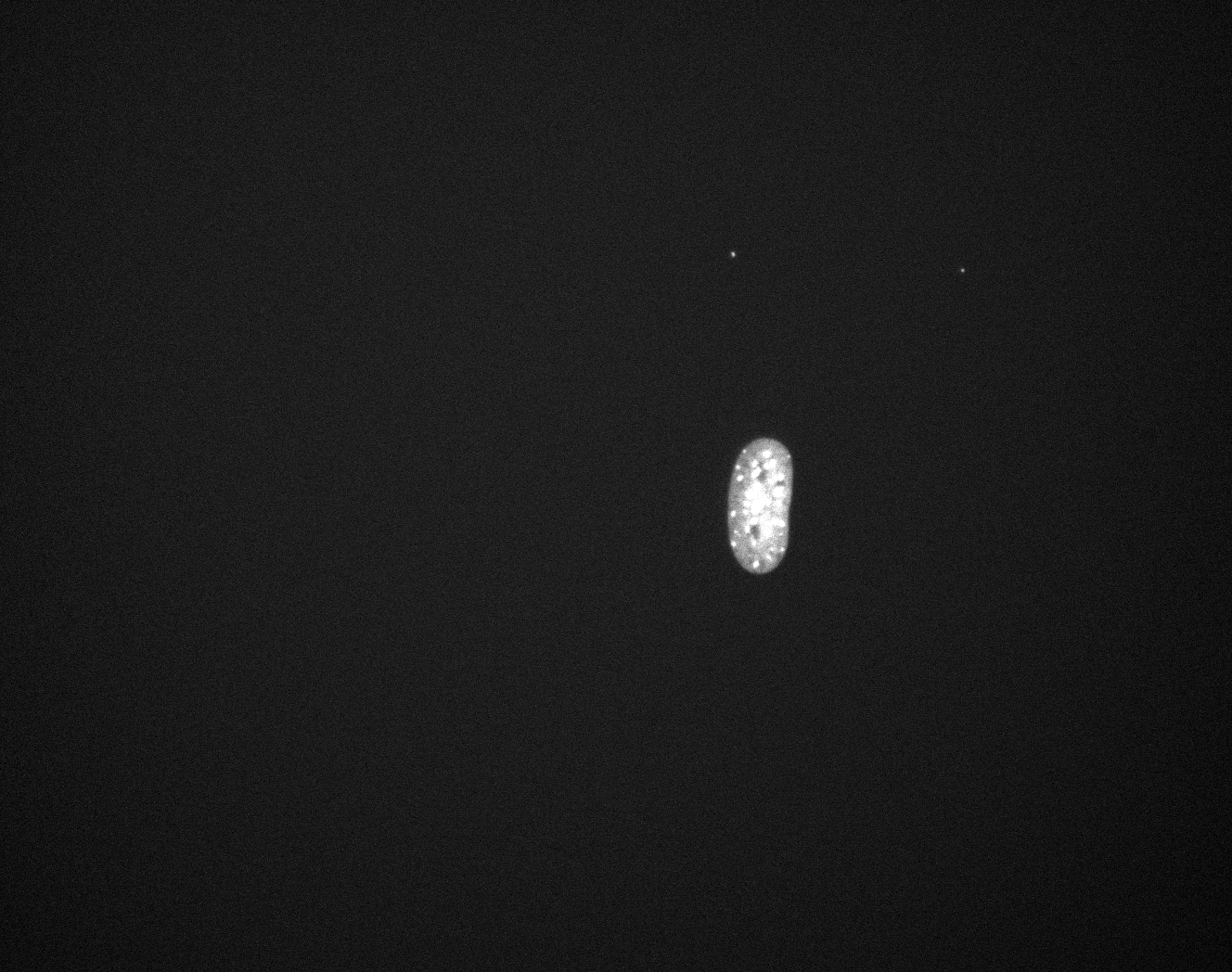

### C_Plate_R_p00_0_A01f17d1.TIF

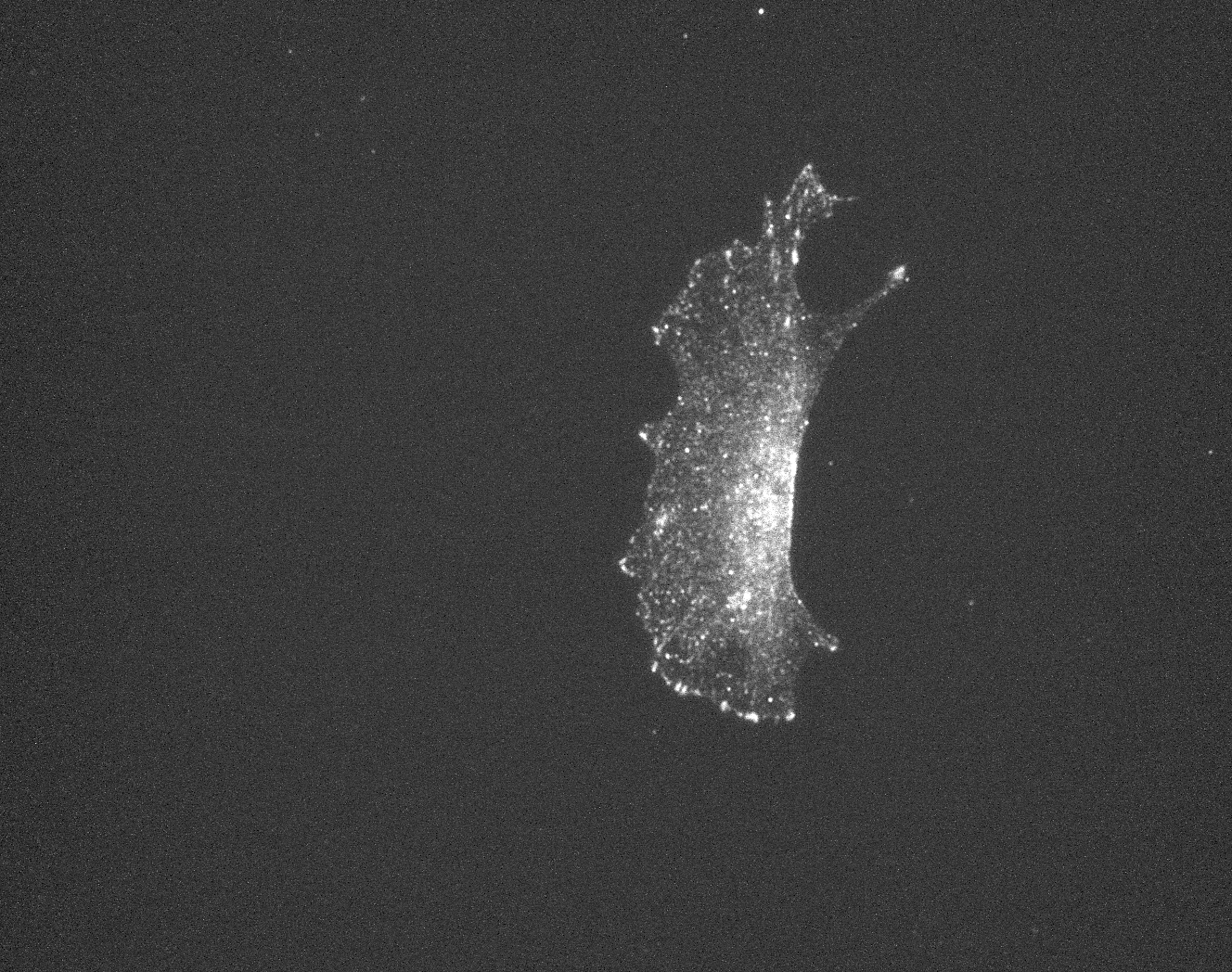

### C_Plate_R_p00_0_A01f17d2.TIF

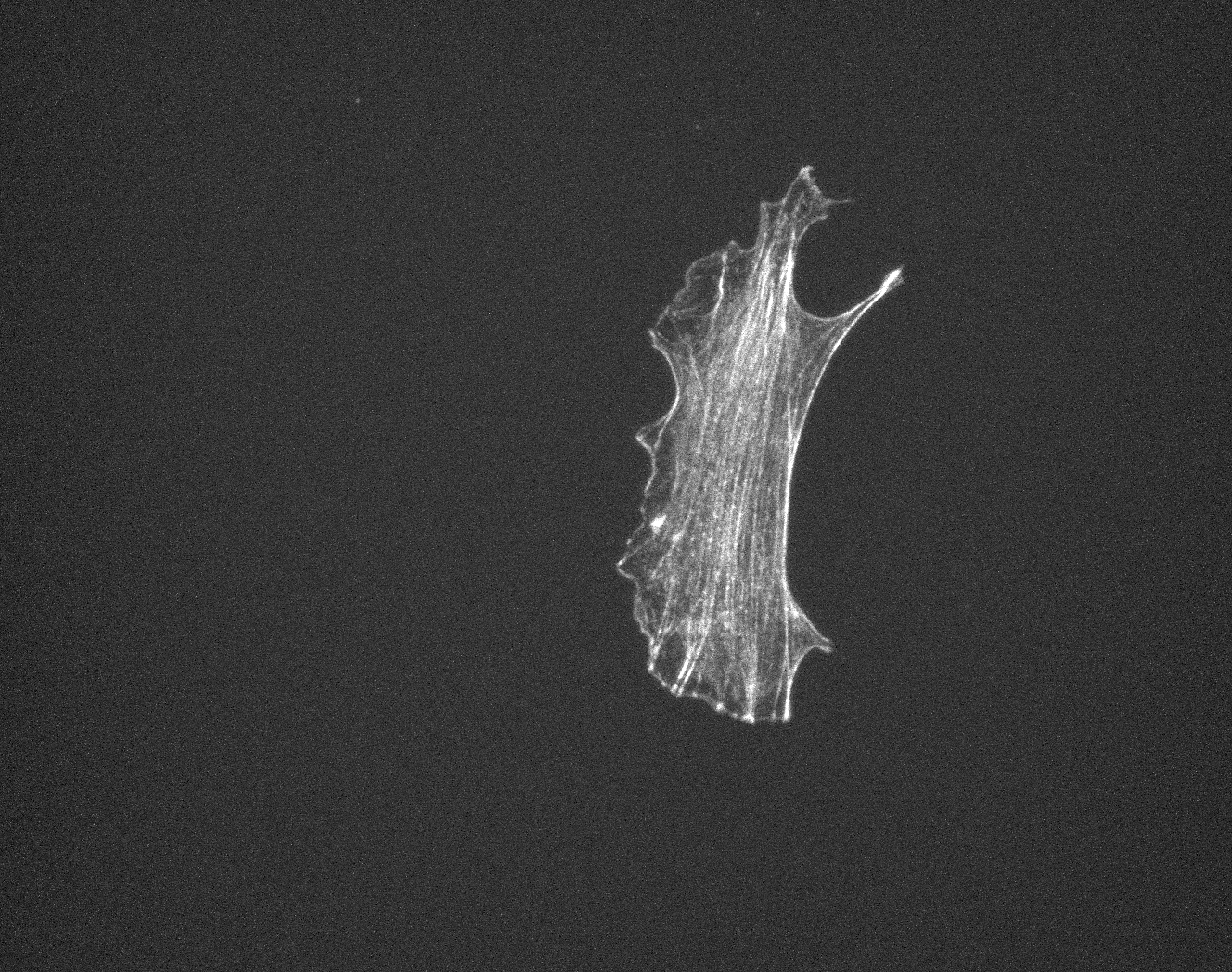

### C_Plate_R_p00_0_A01f17d3.TIF

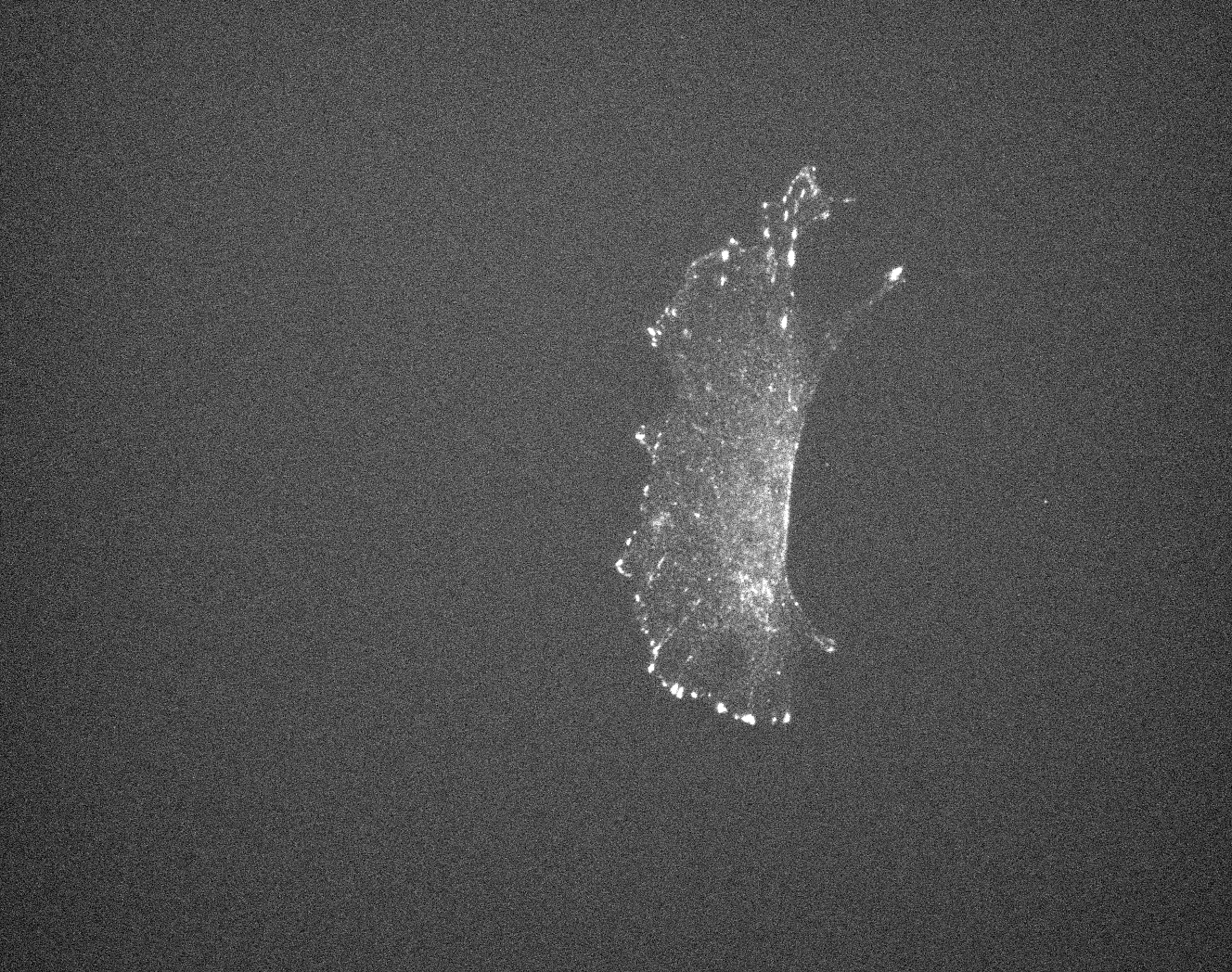

### D_groundtruth.jpg

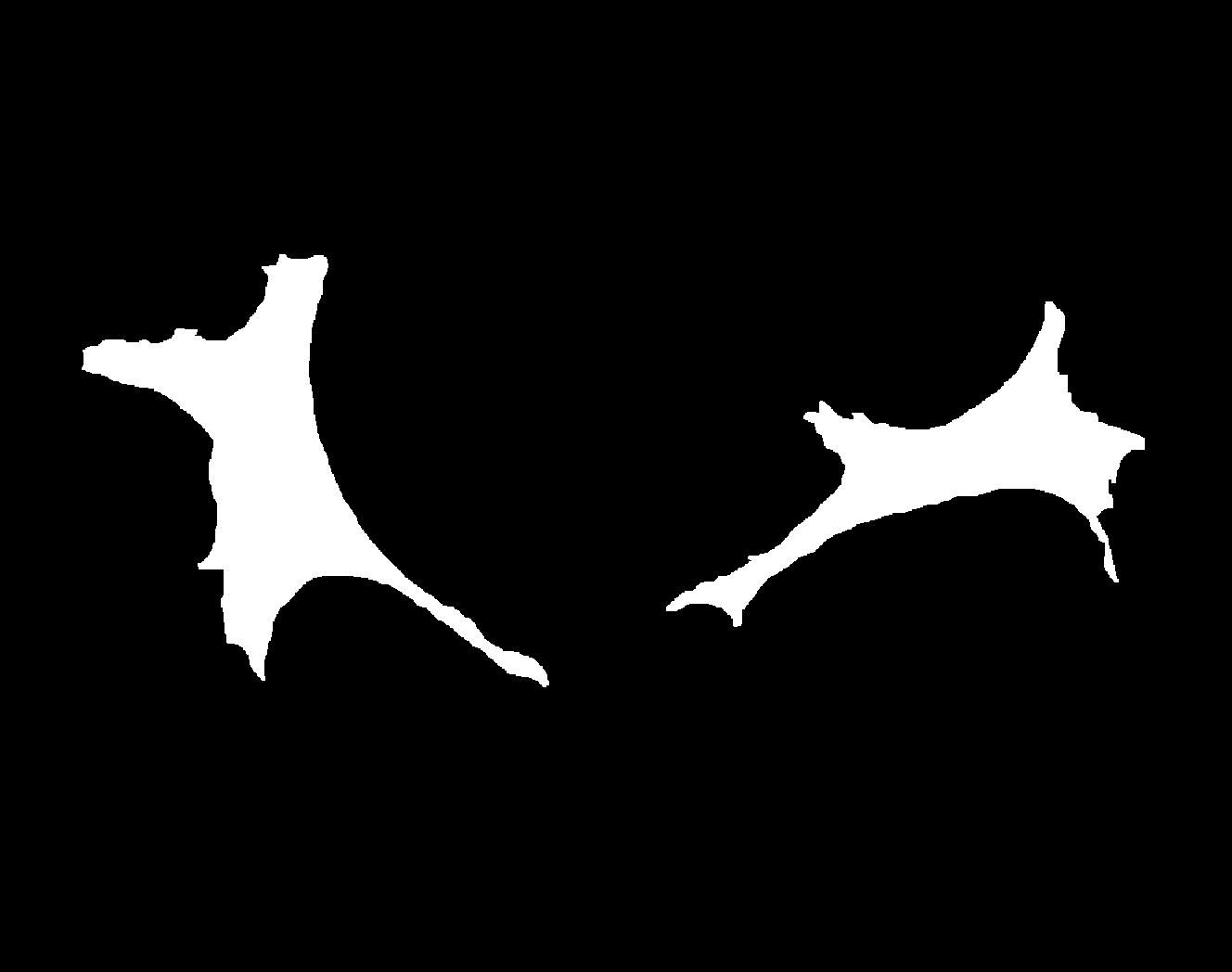

### D_Plate_R_p00_0_A01f15d0.TIF

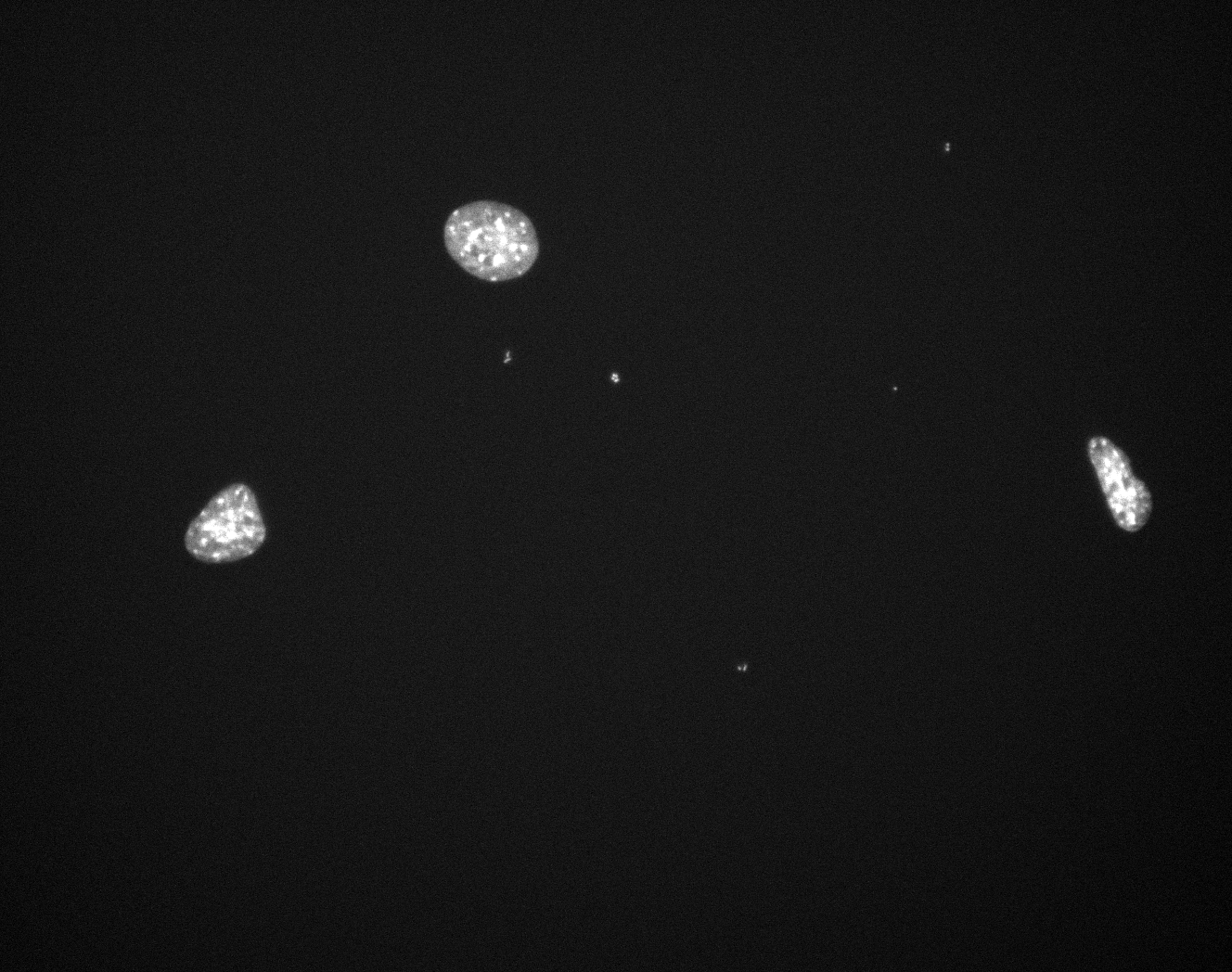

### D_Plate_R_p00_0_A01f15d1.TIF

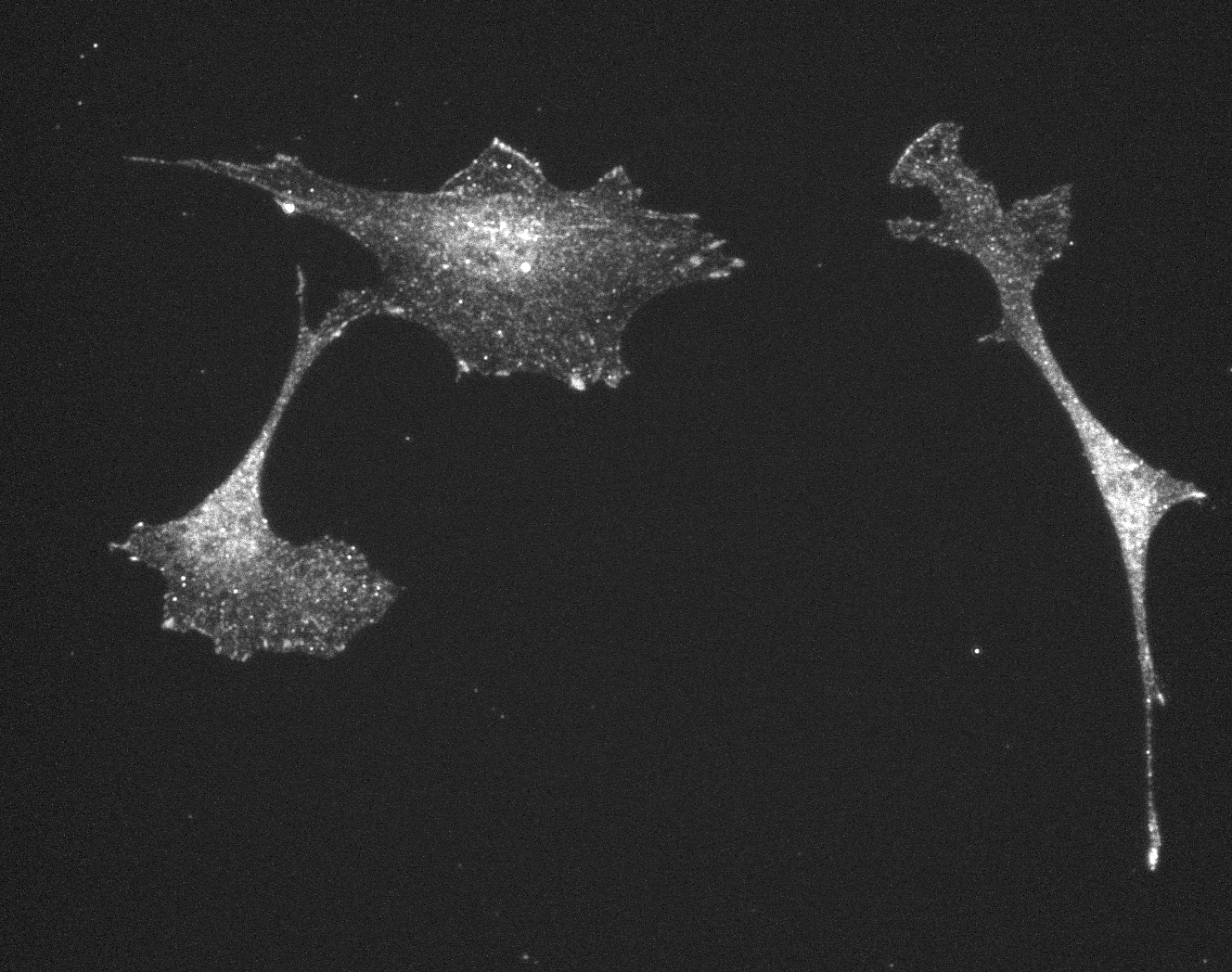

### D_Plate_R_p00_0_A01f15d2.TIF

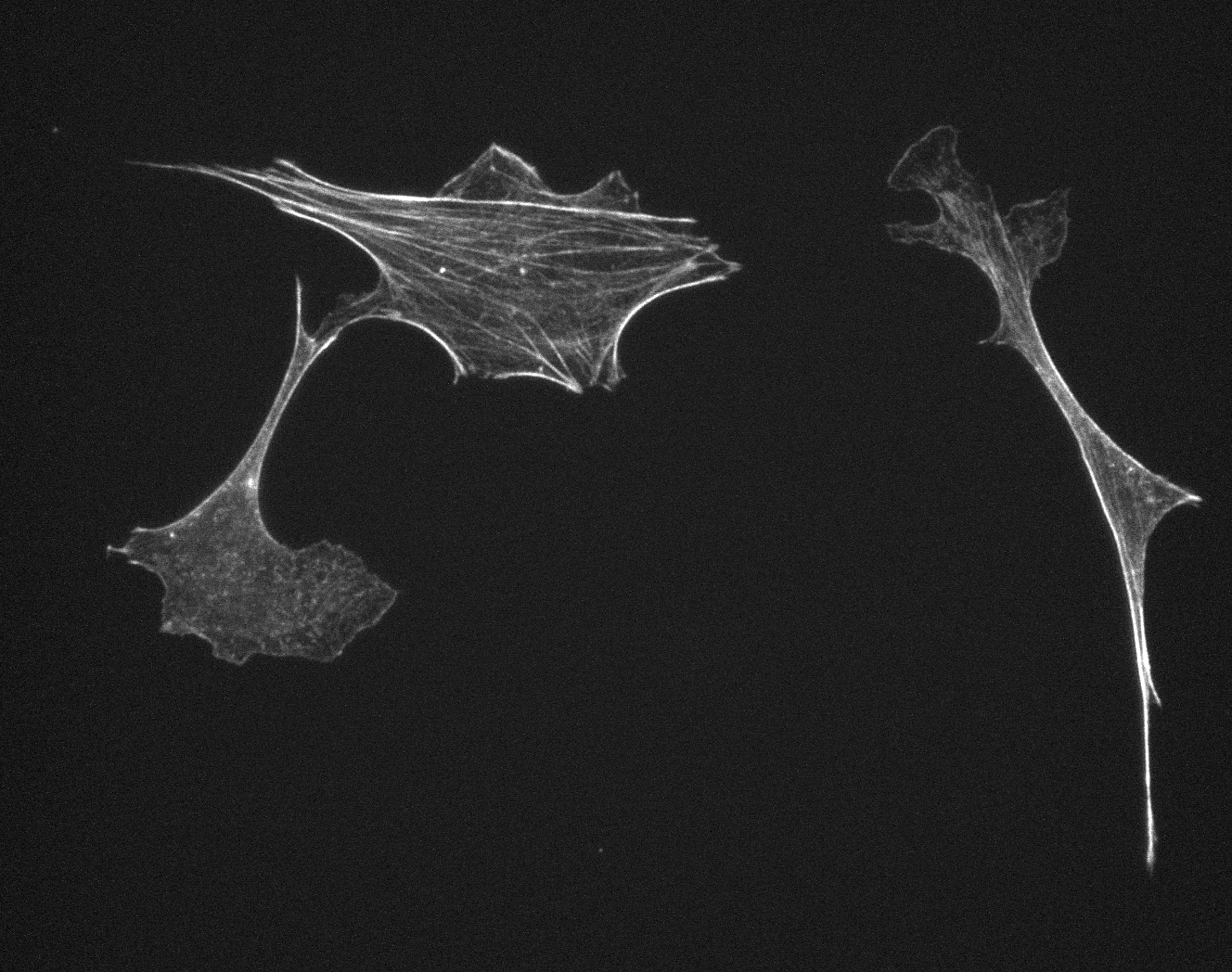

### D_Plate_R_p00_0_A01f15d3.TIF

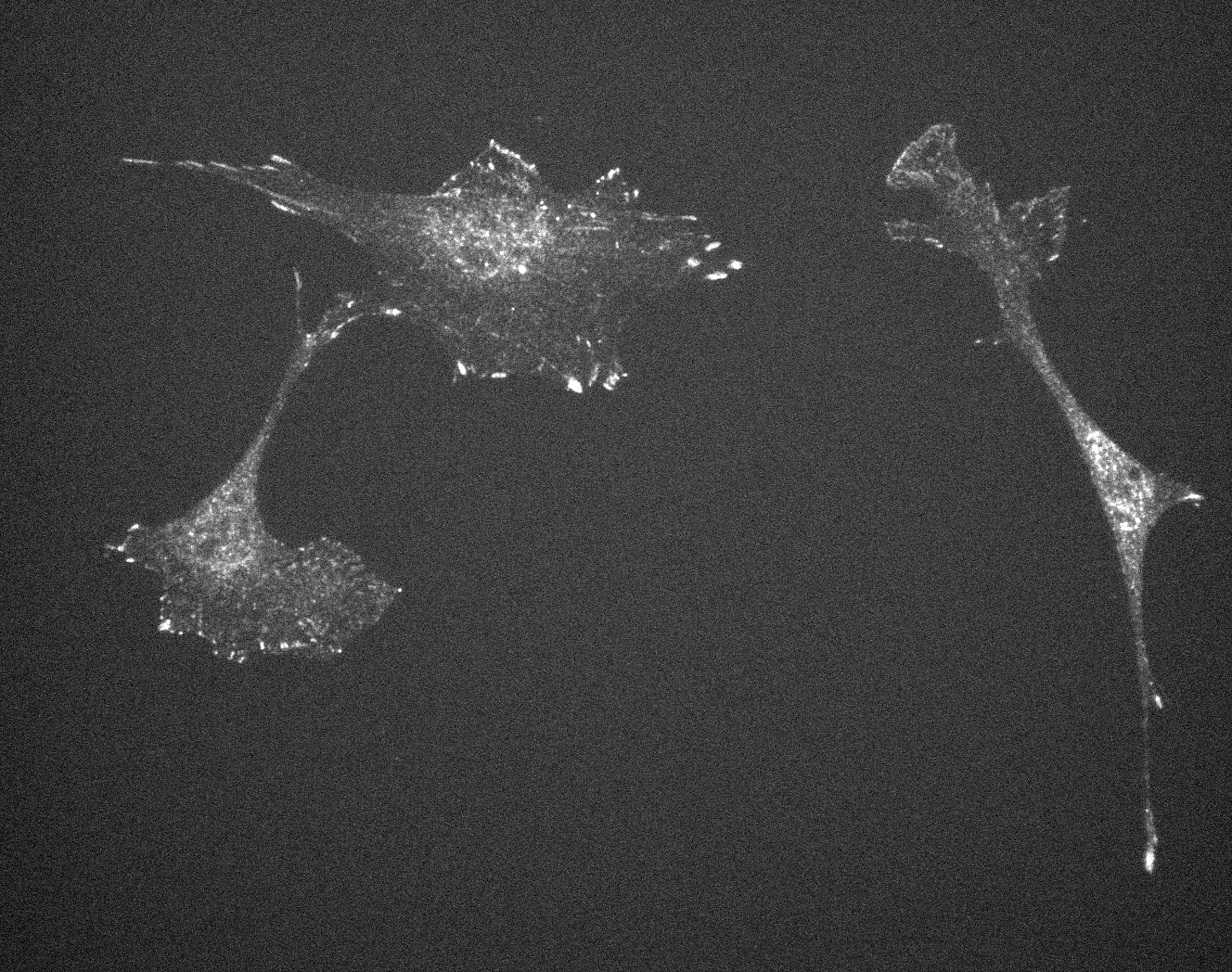

### D_Plate_R_p00_0_A01f59d0.TIF

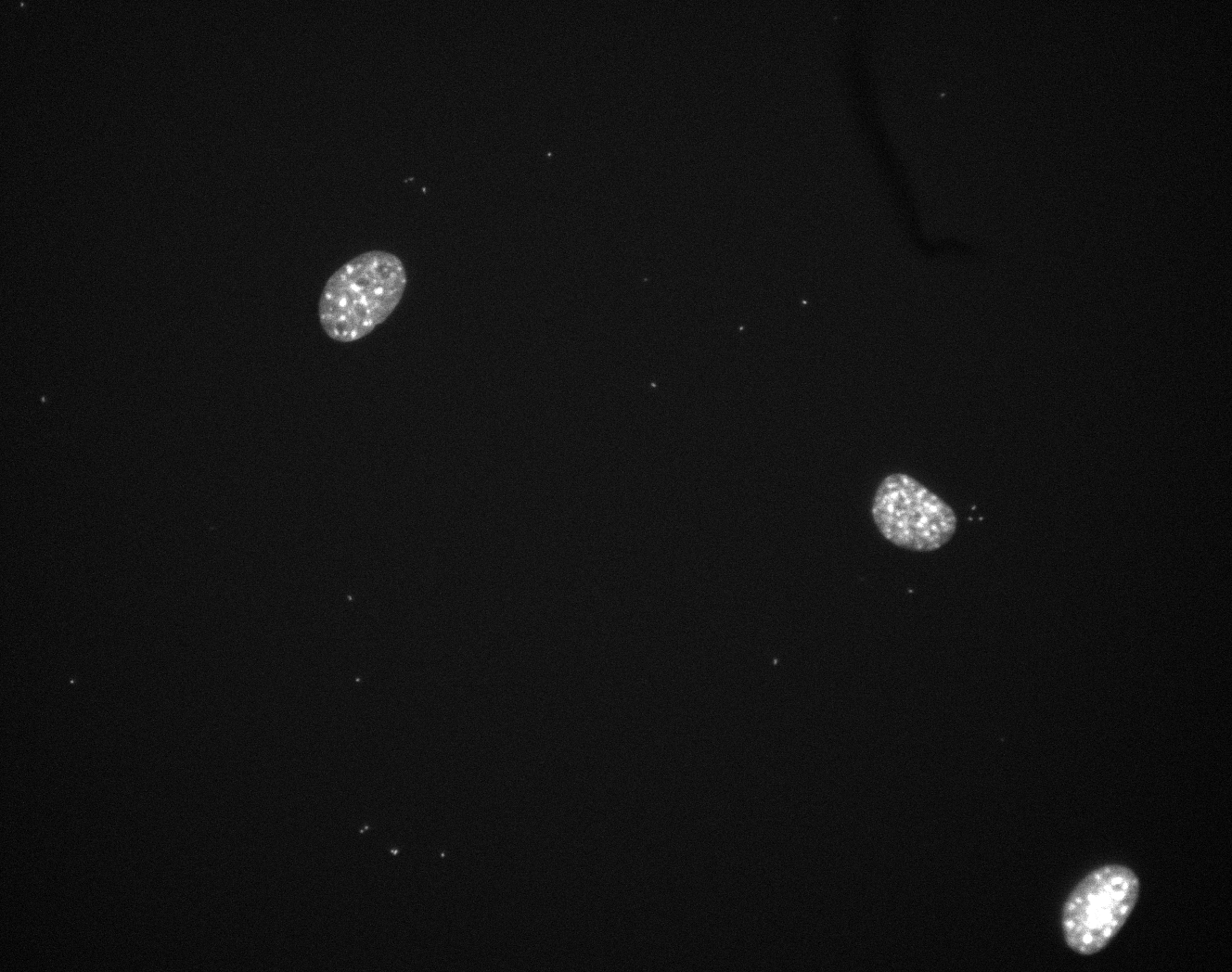

### D_Plate_R_p00_0_A01f59d1.TIF

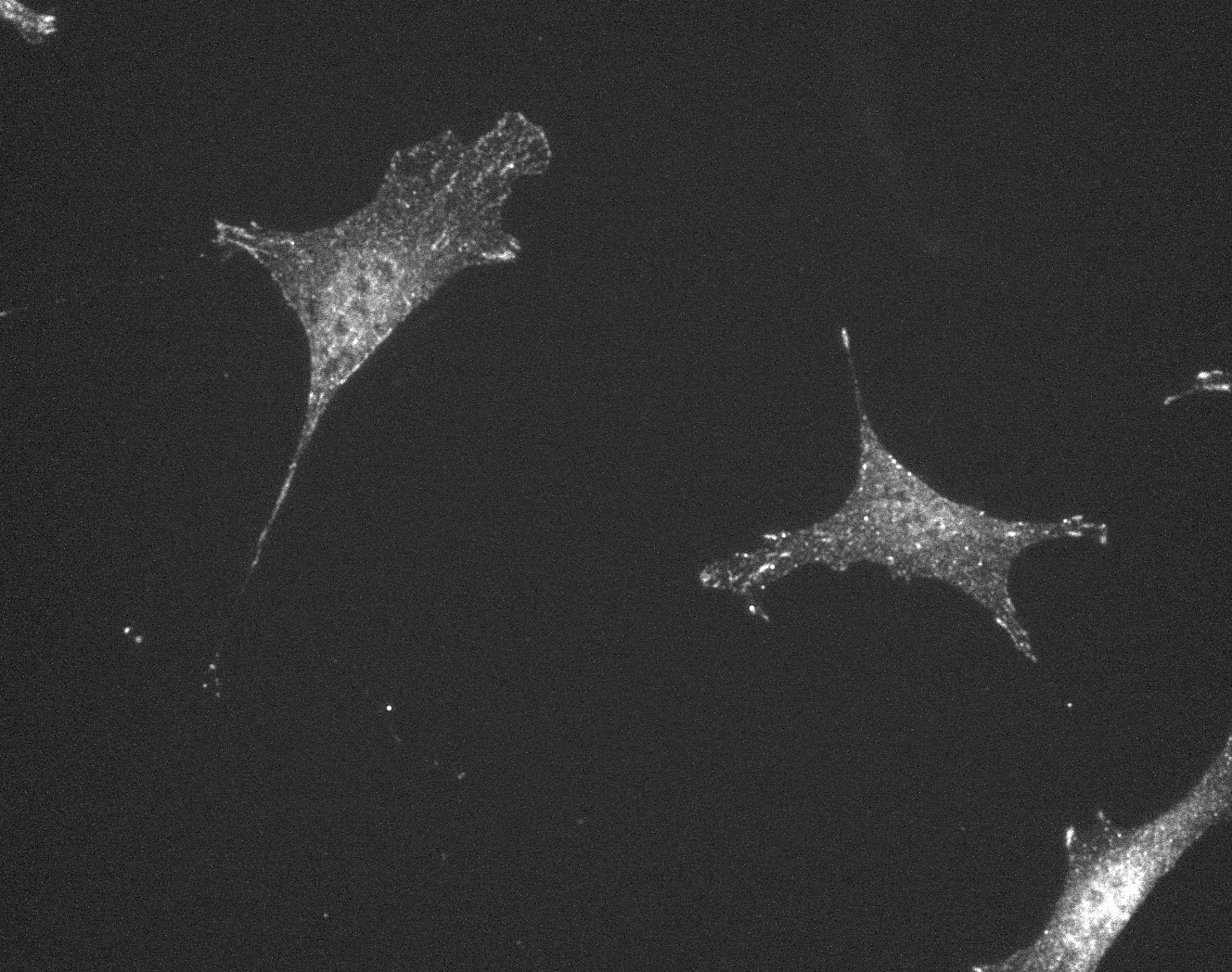

### D_Plate_R_p00_0_A01f59d2.TIF

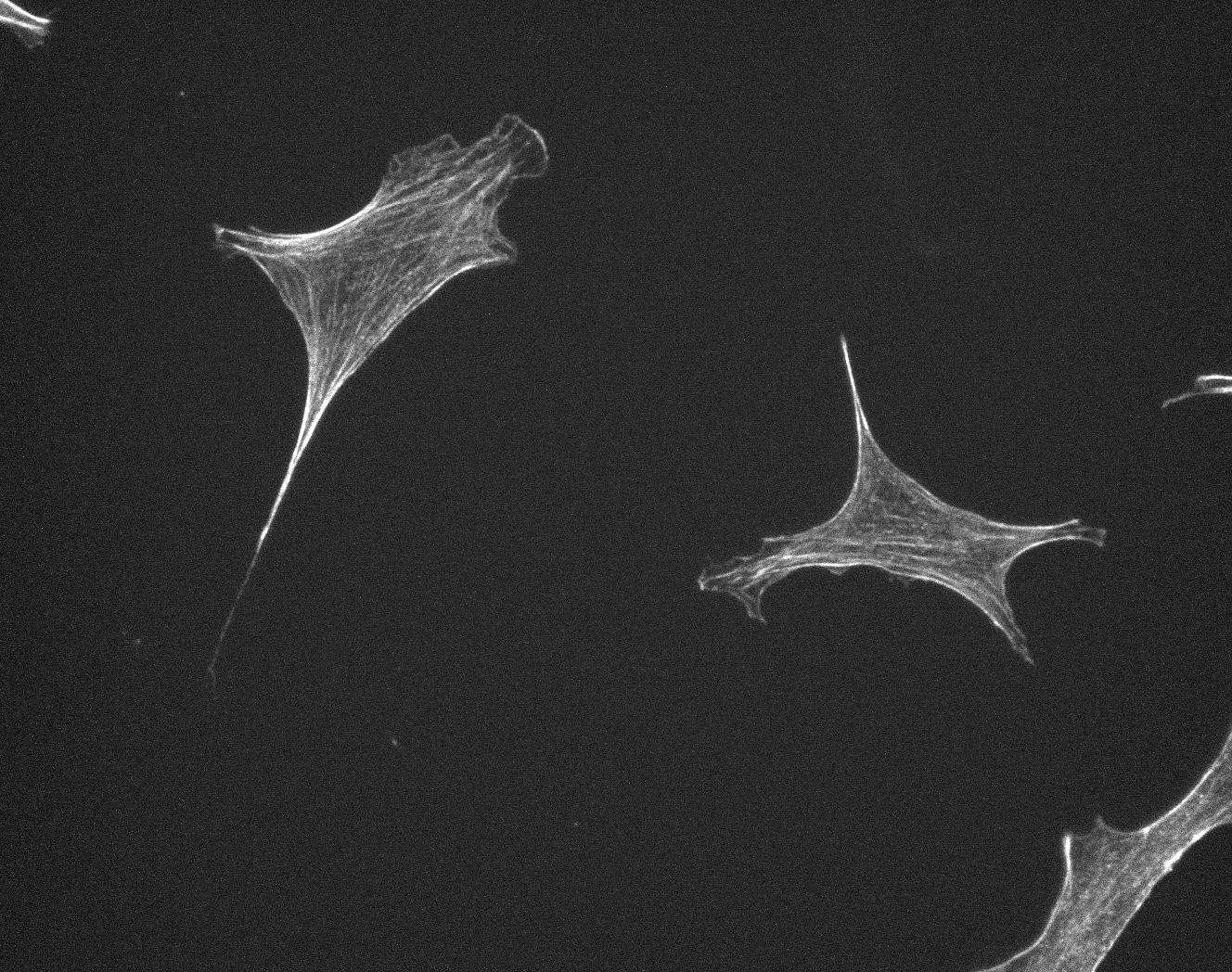

### D_Plate_R_p00_0_A01f59d3.TIF

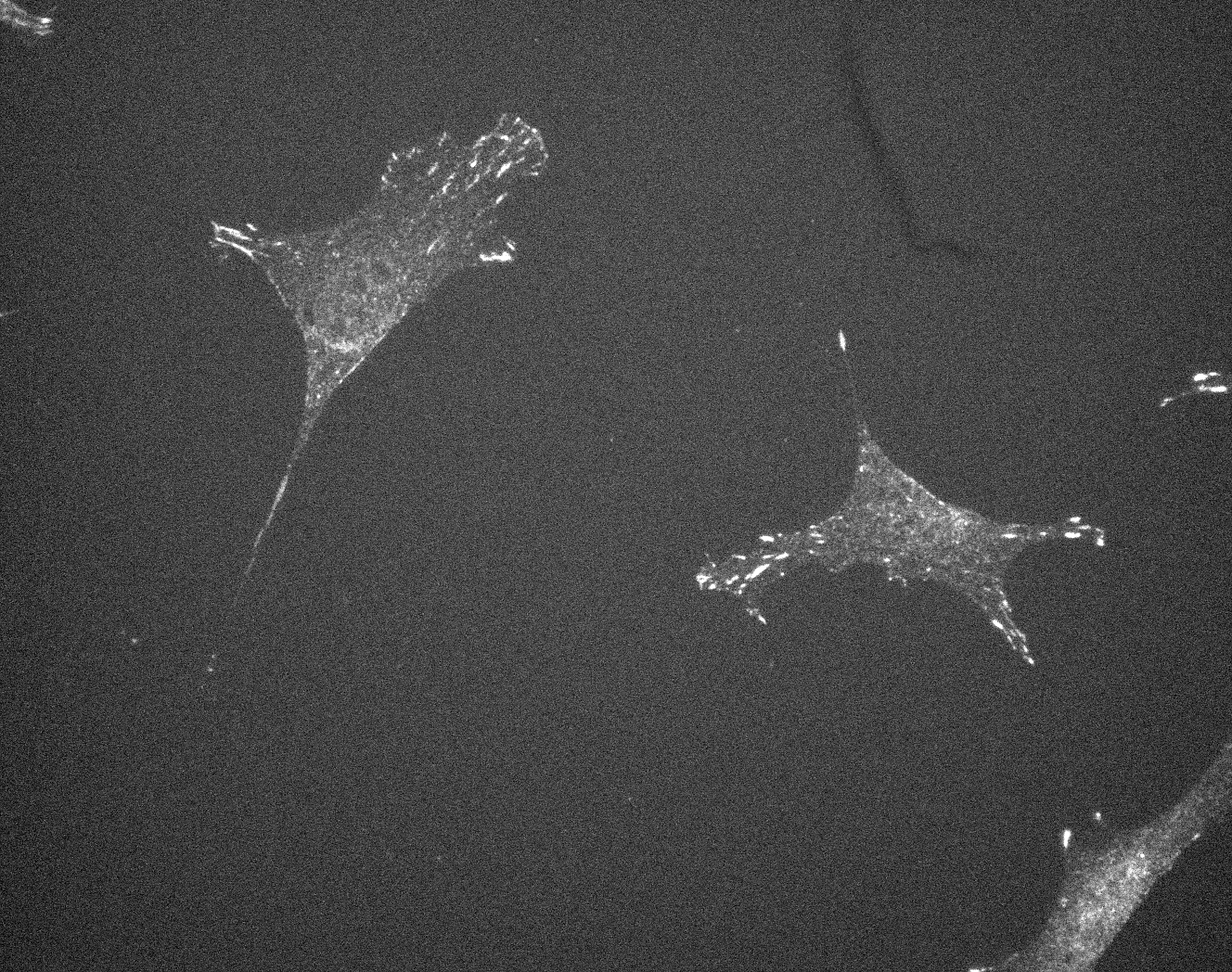

### E_groundtruth.jpg

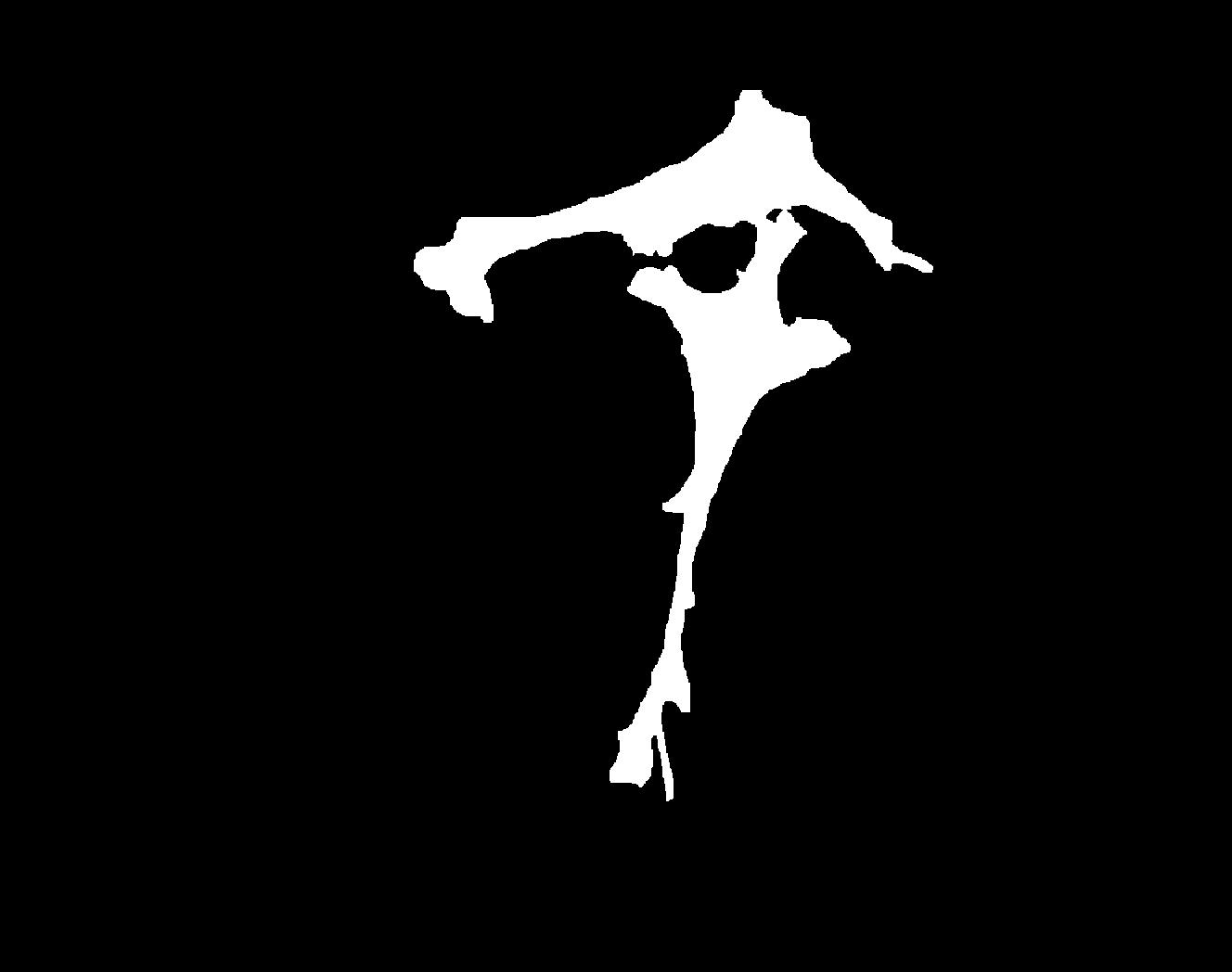

### E_Plate_R_p00_0_A01f16d0.TIF

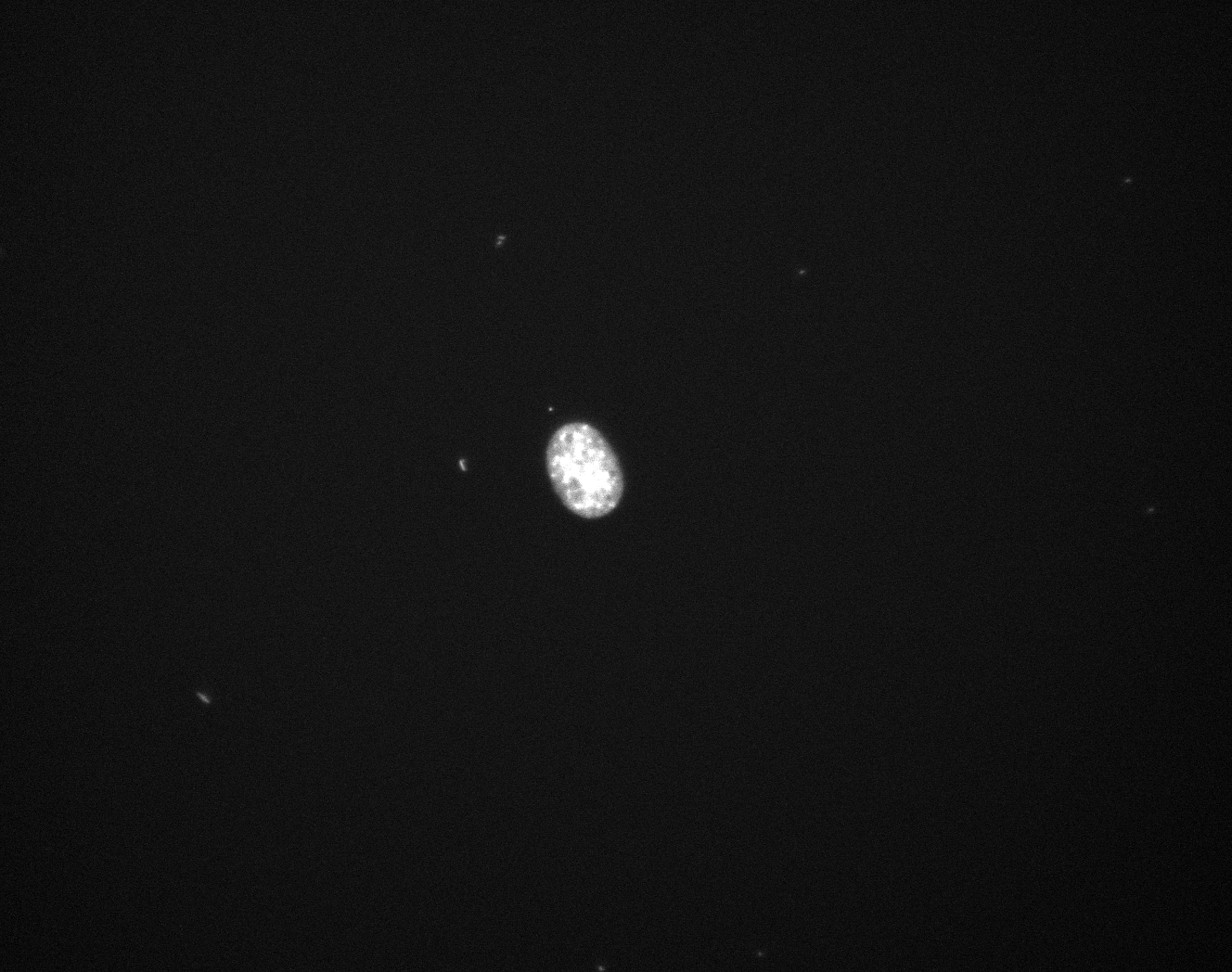

### E_Plate_R_p00_0_A01f16d1.TIF

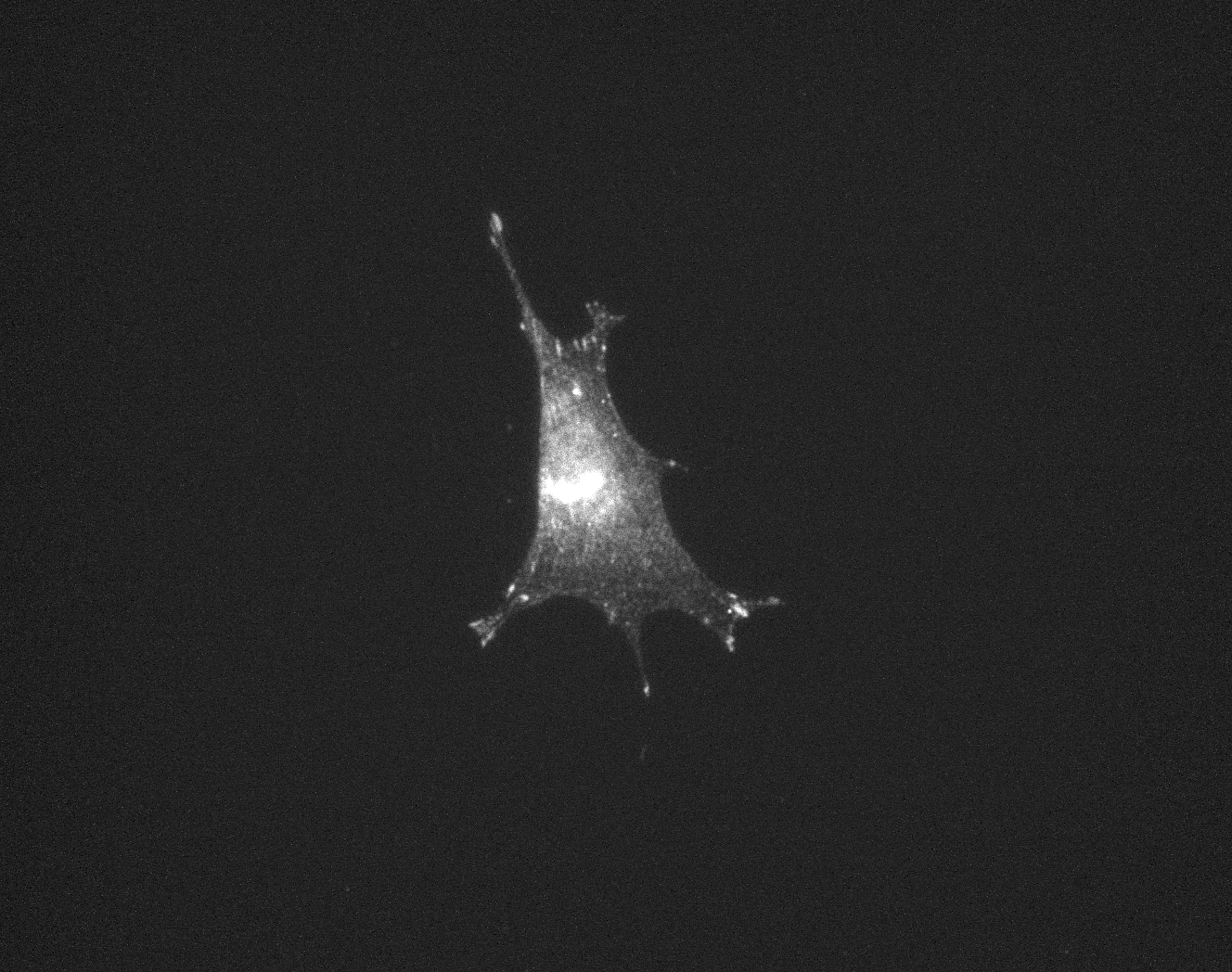

### E_Plate_R_p00_0_A01f16d2.TIF

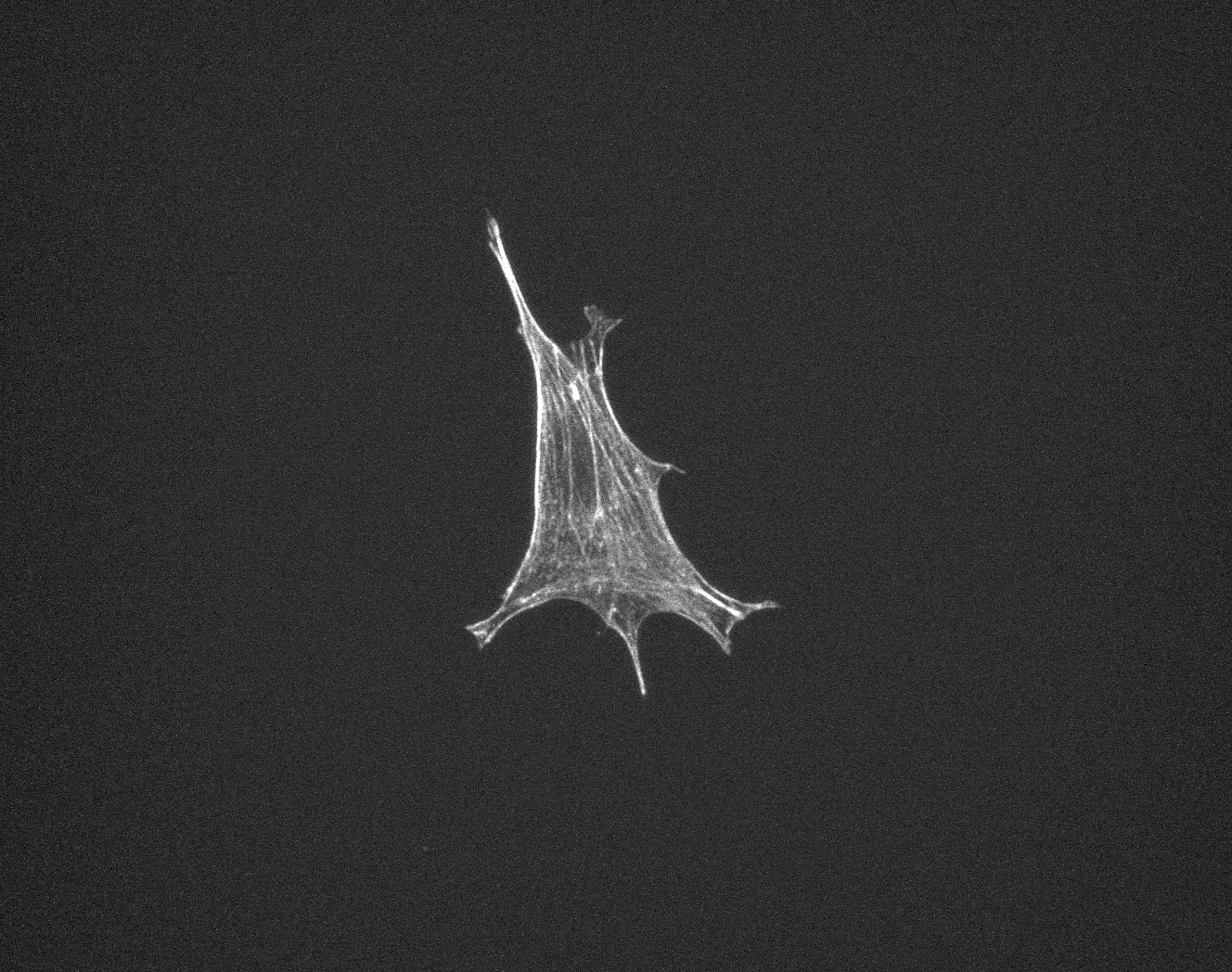

### E_Plate_R_p00_0_A01f16d3.TIF

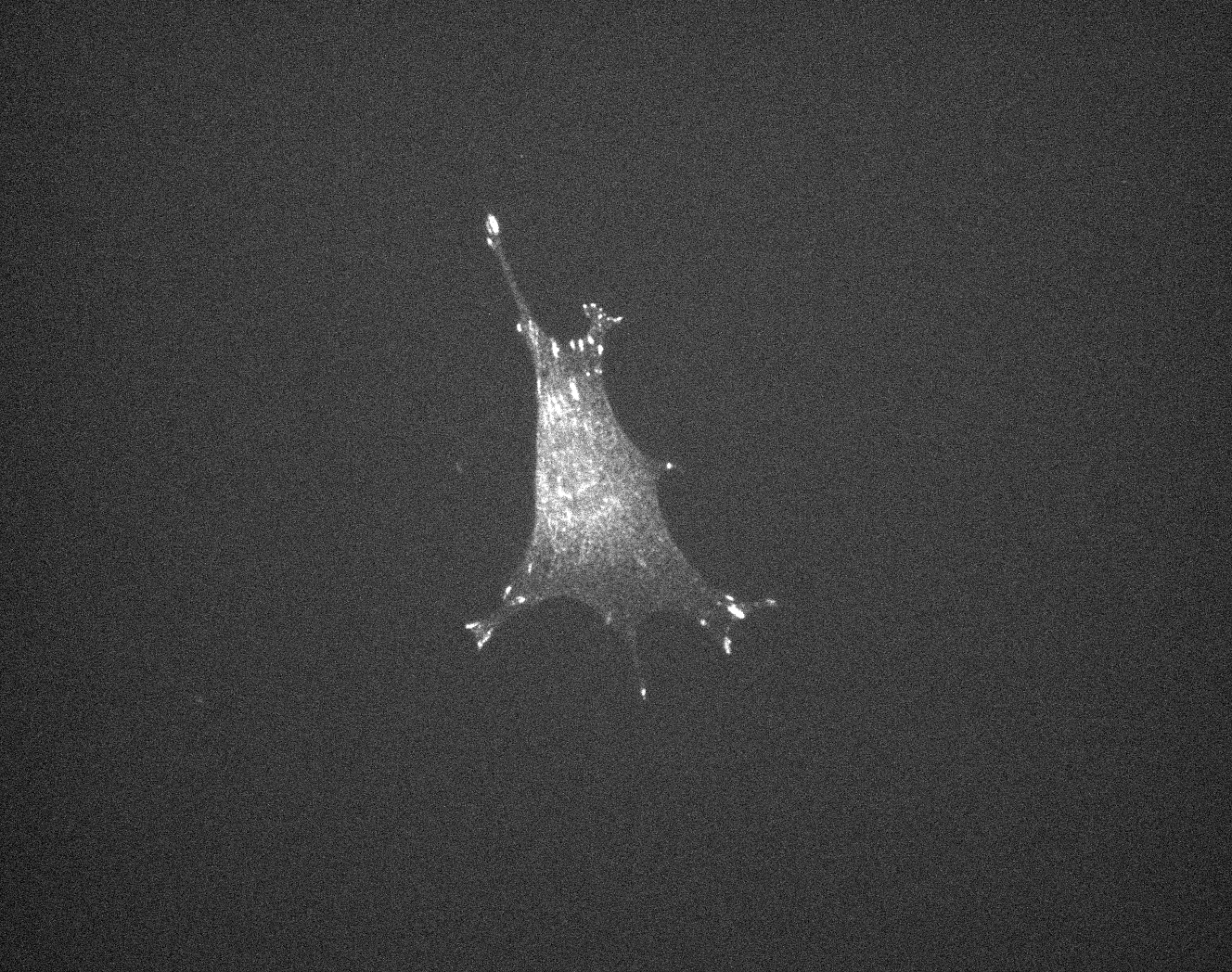

### E_Plate_R_p00_0_A01f34d0.TIF

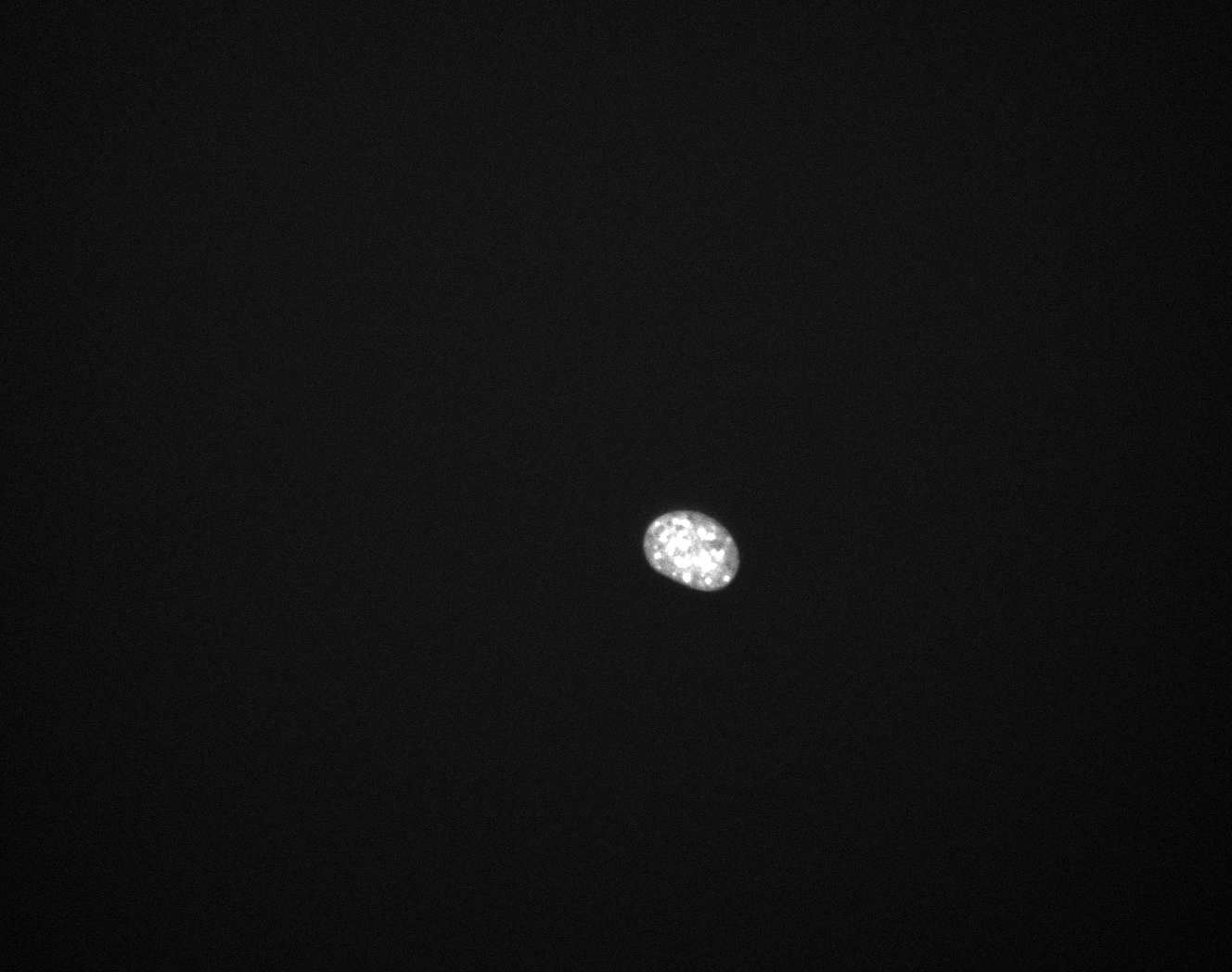

### E_Plate_R_p00_0_A01f34d1.TIF

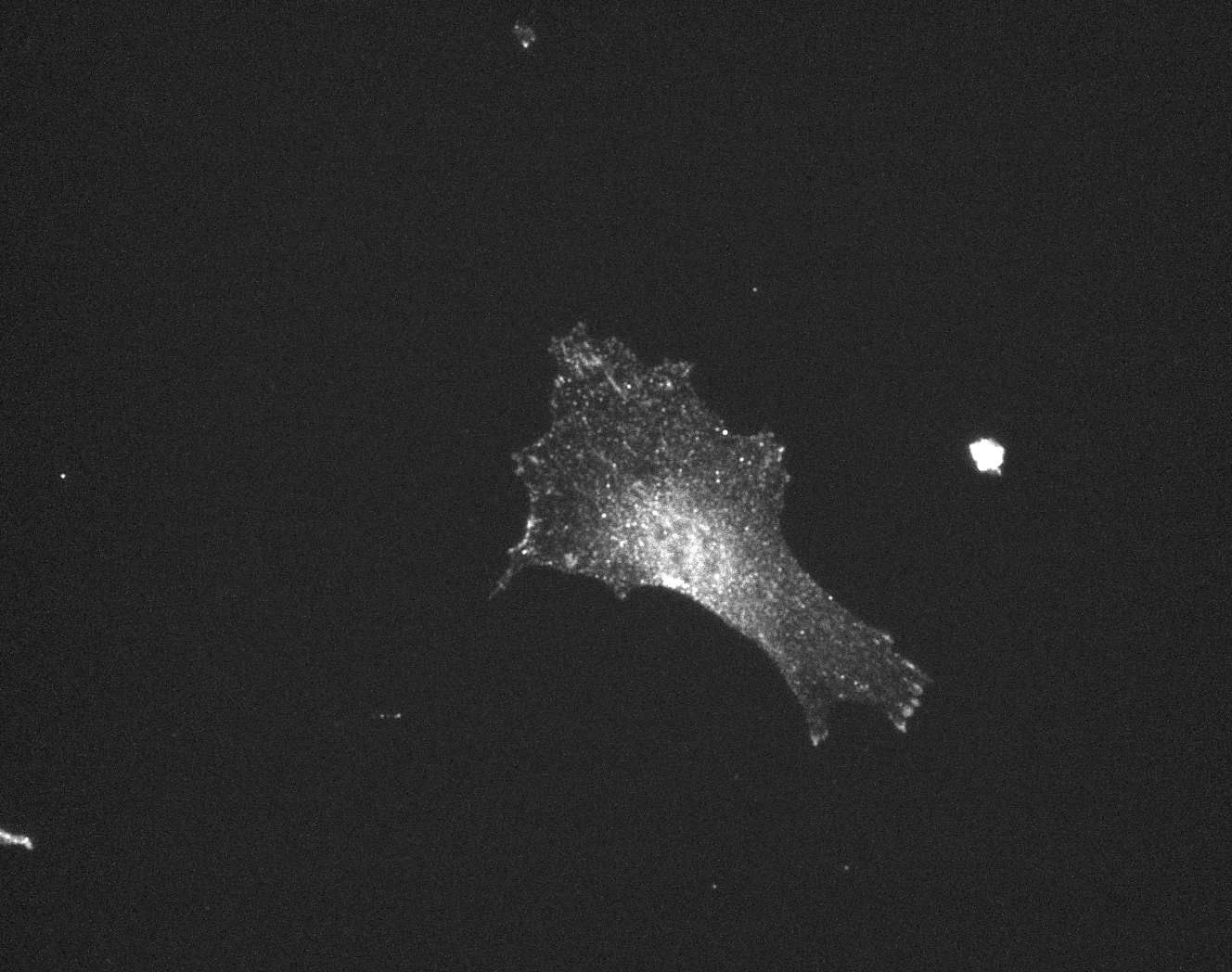

### E_Plate_R_p00_0_A01f34d2.TIF

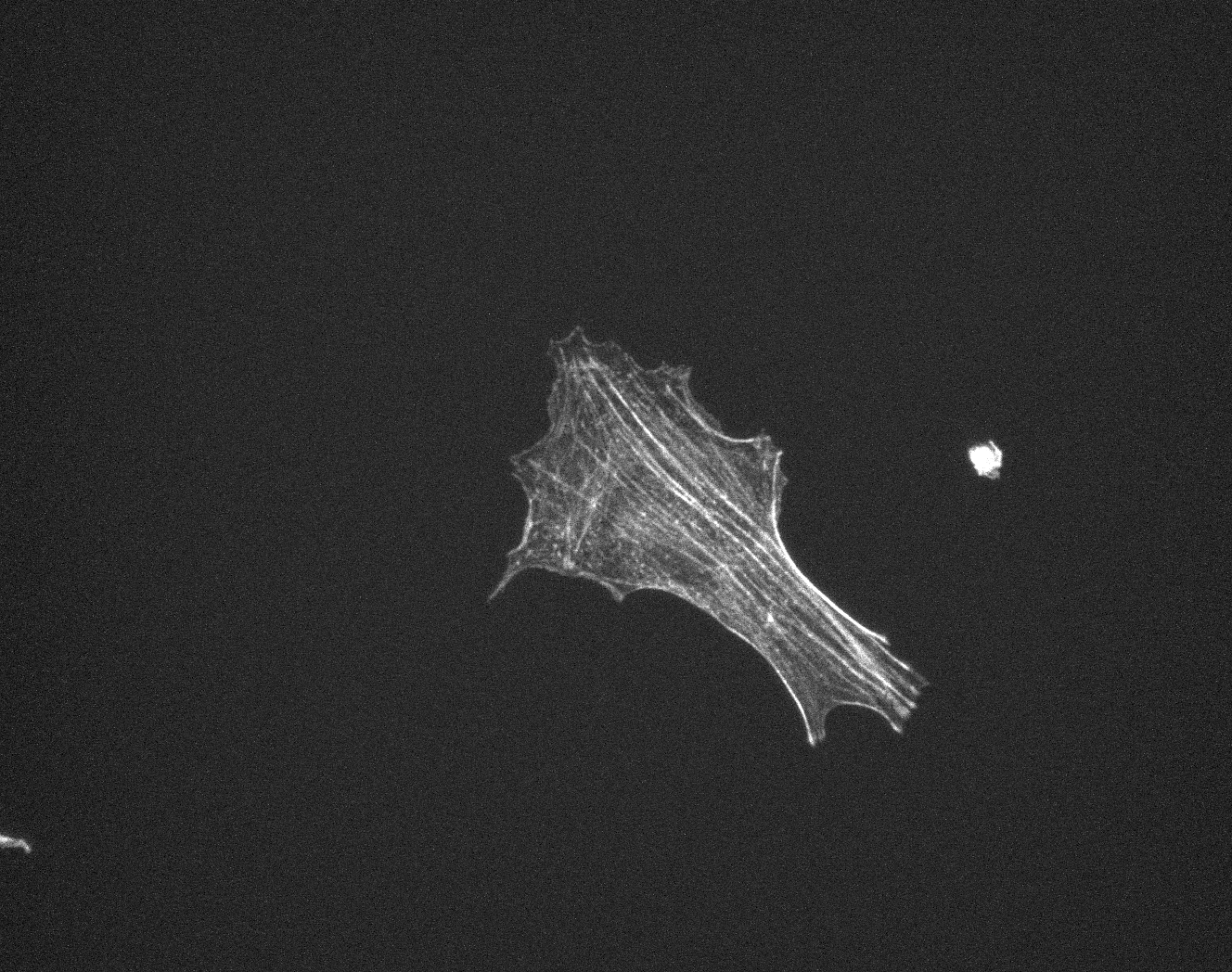

### E_Plate_R_p00_0_A01f34d3.TIF

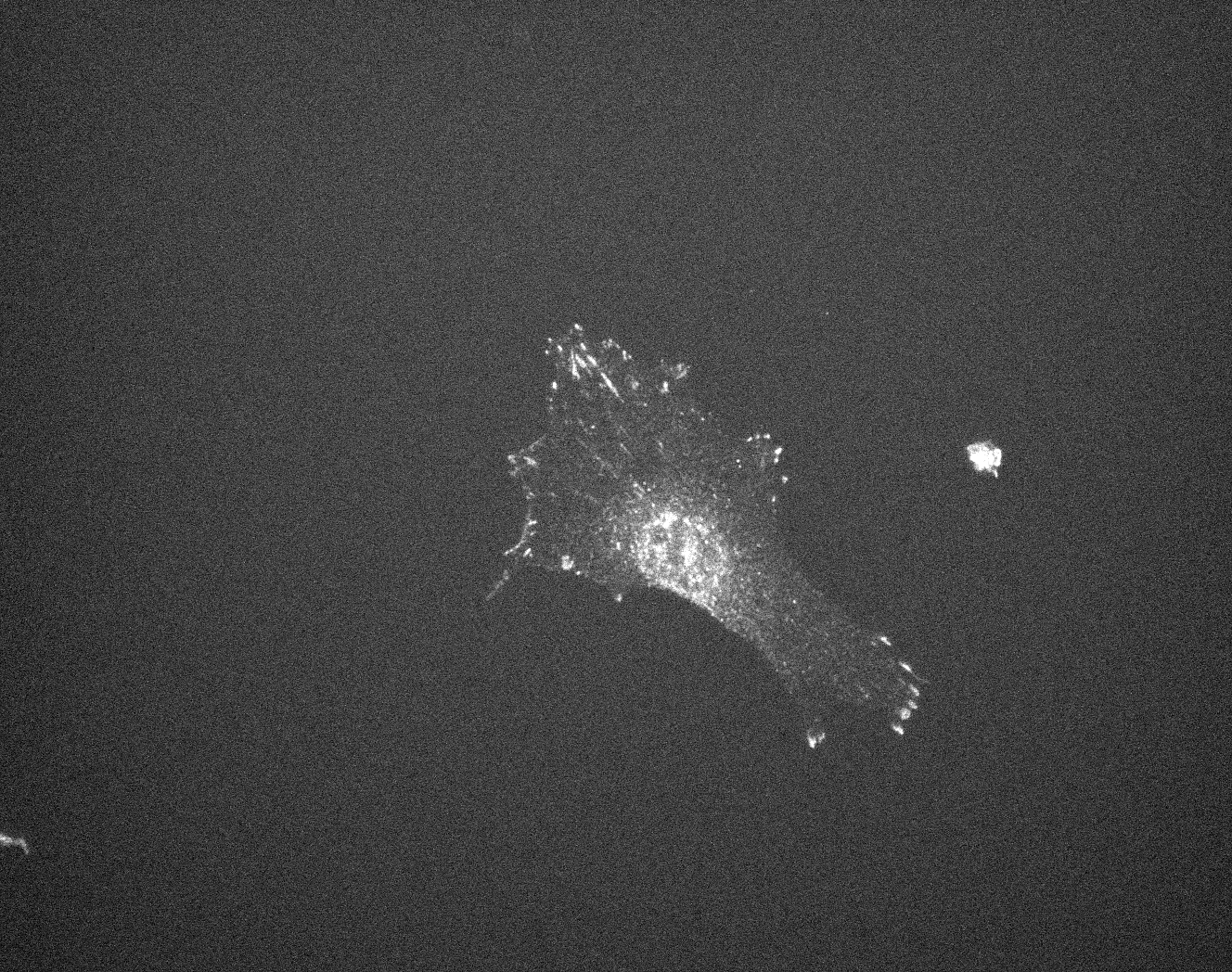

### F_groundtruth.jpg

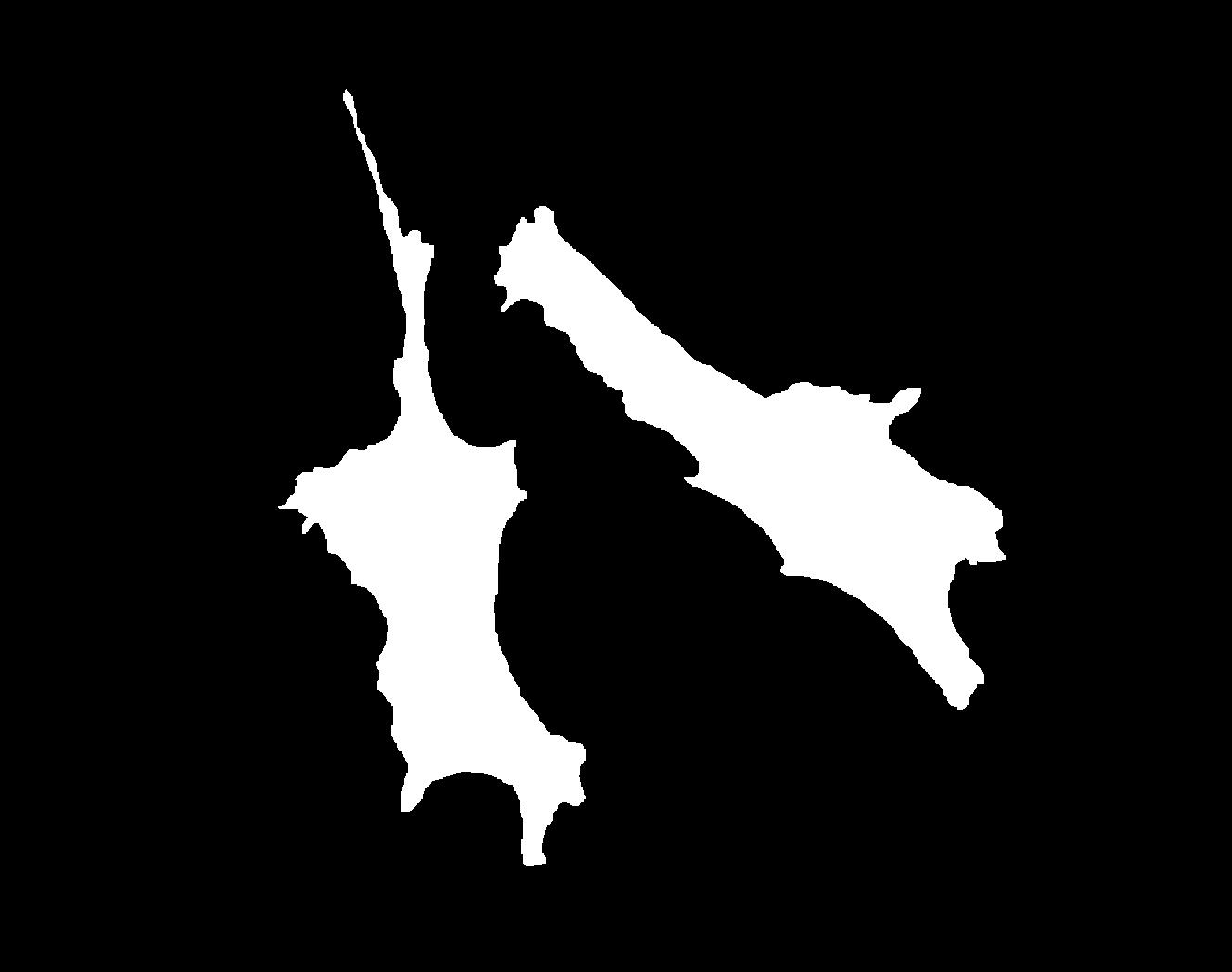

### F_Plate_R_p00_0_A01f15d0.TIF

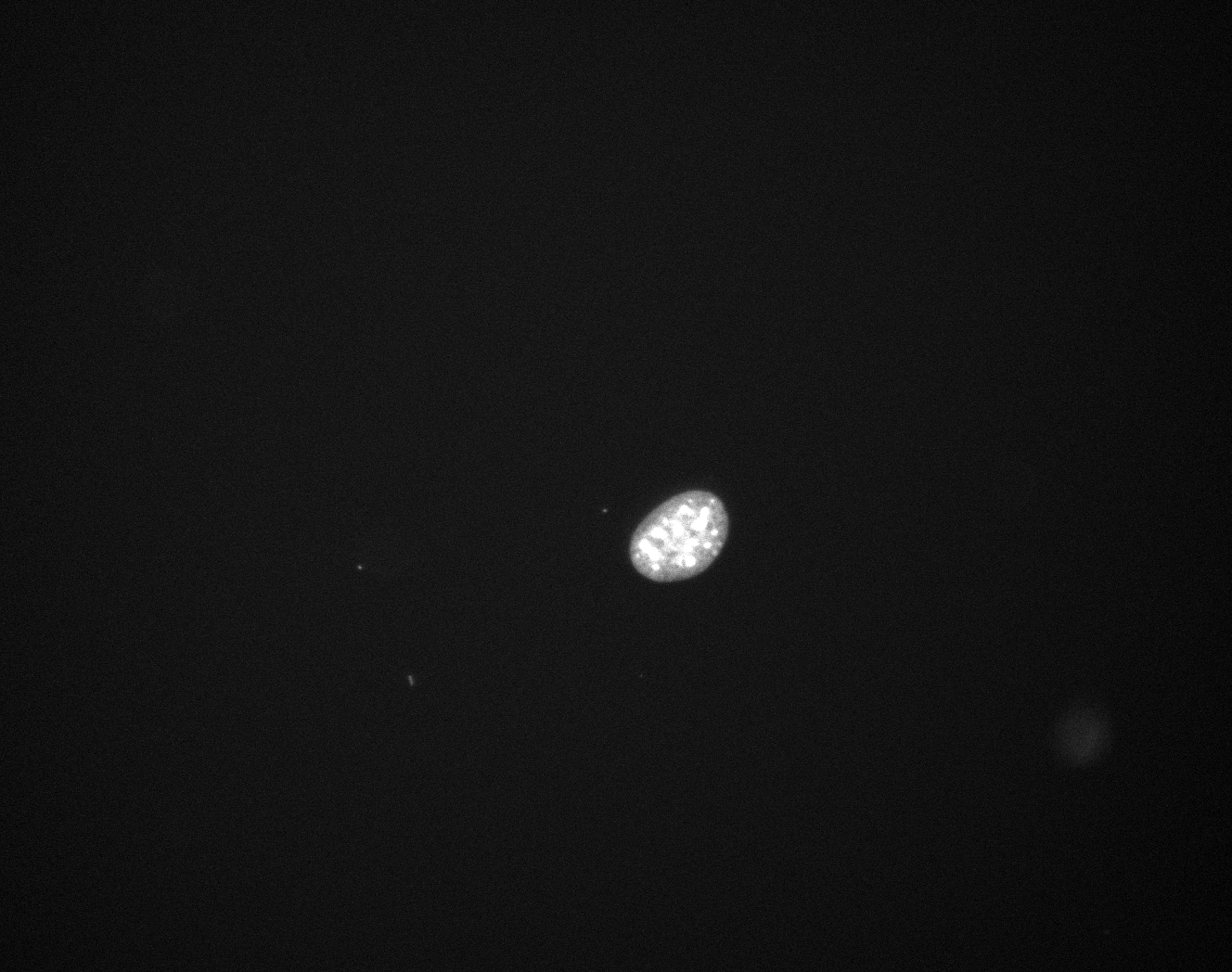

### F_Plate_R_p00_0_A01f15d1.TIF

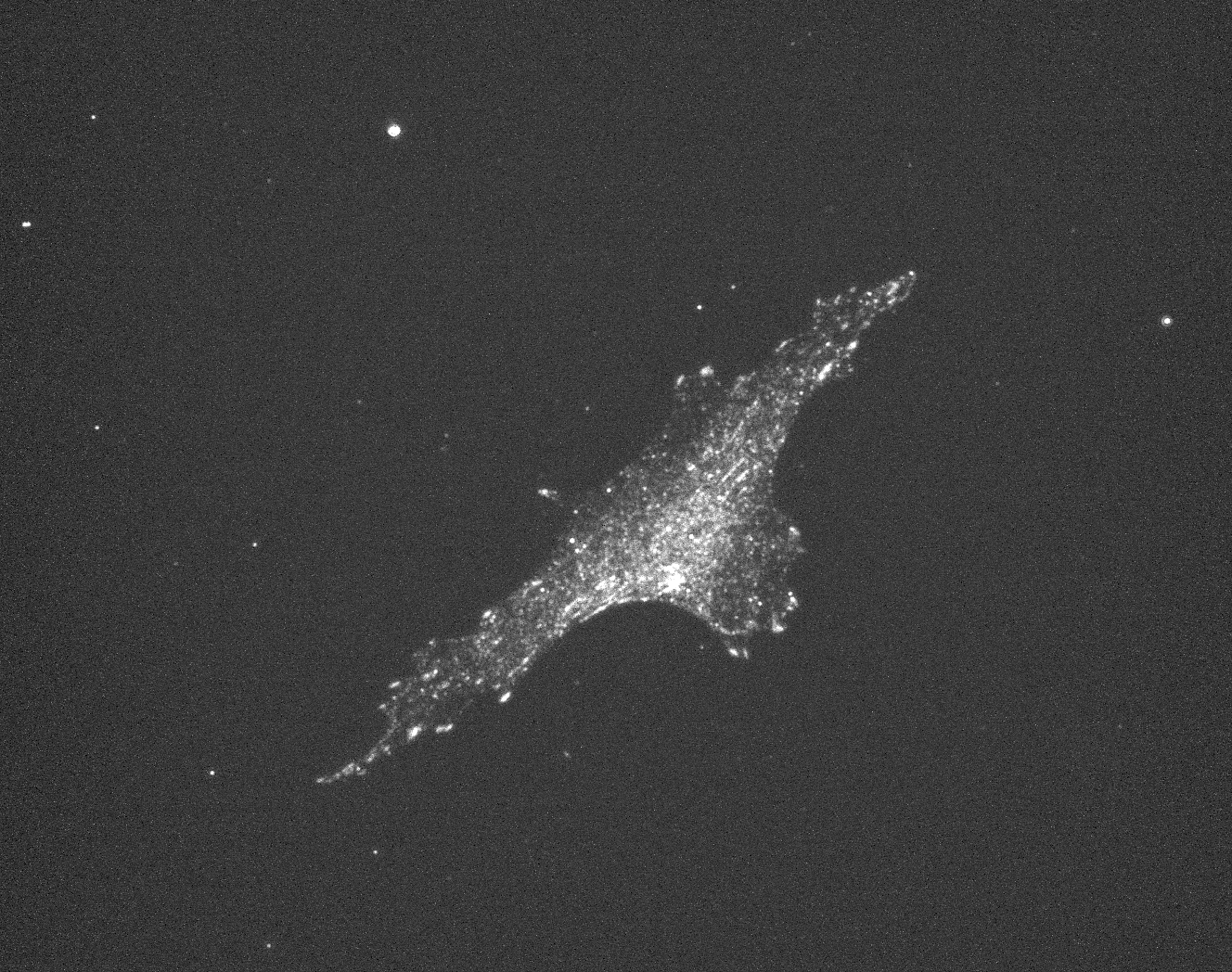

### F_Plate_R_p00_0_A01f15d2.TIF

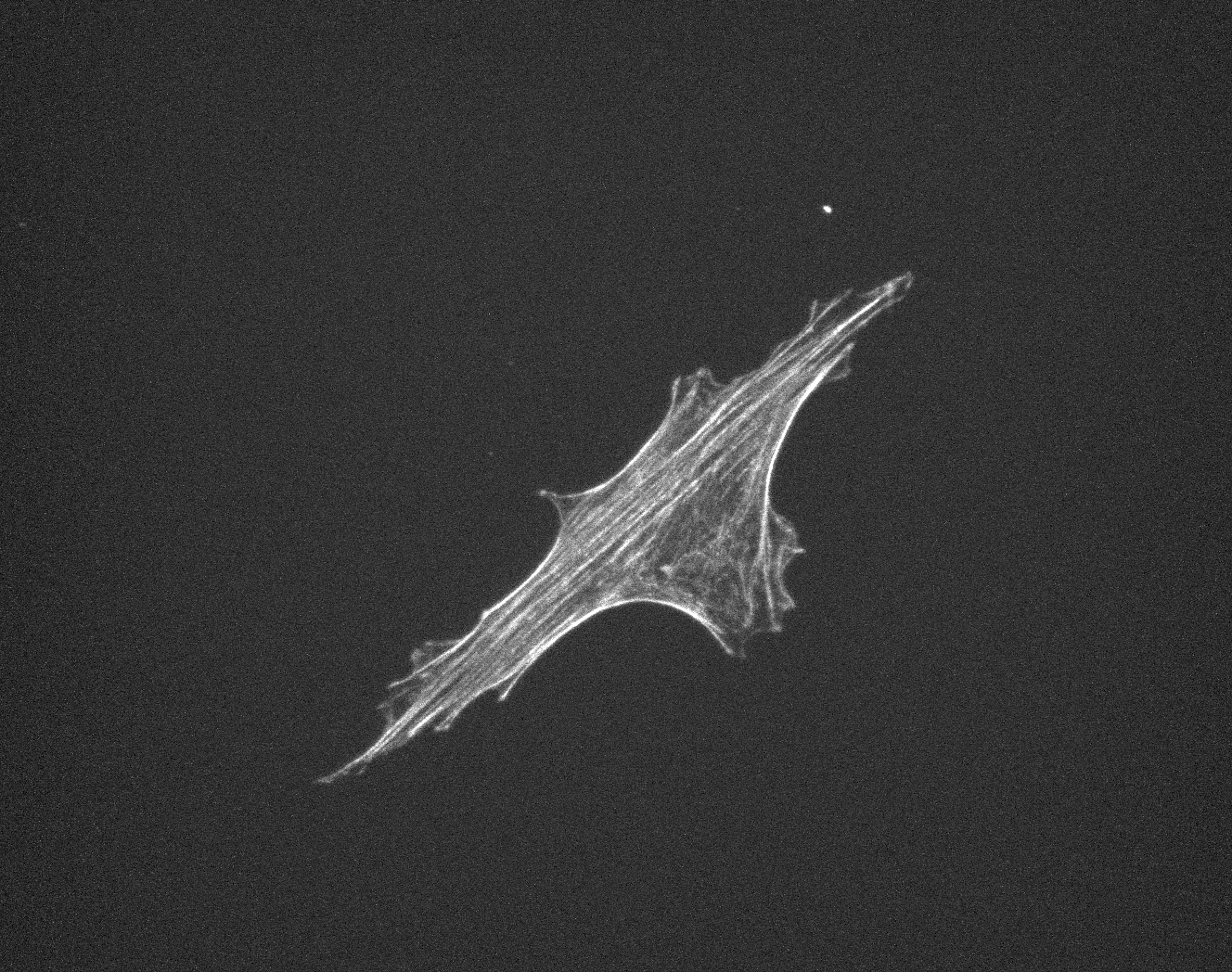

### F_Plate_R_p00_0_A01f15d3.TIF

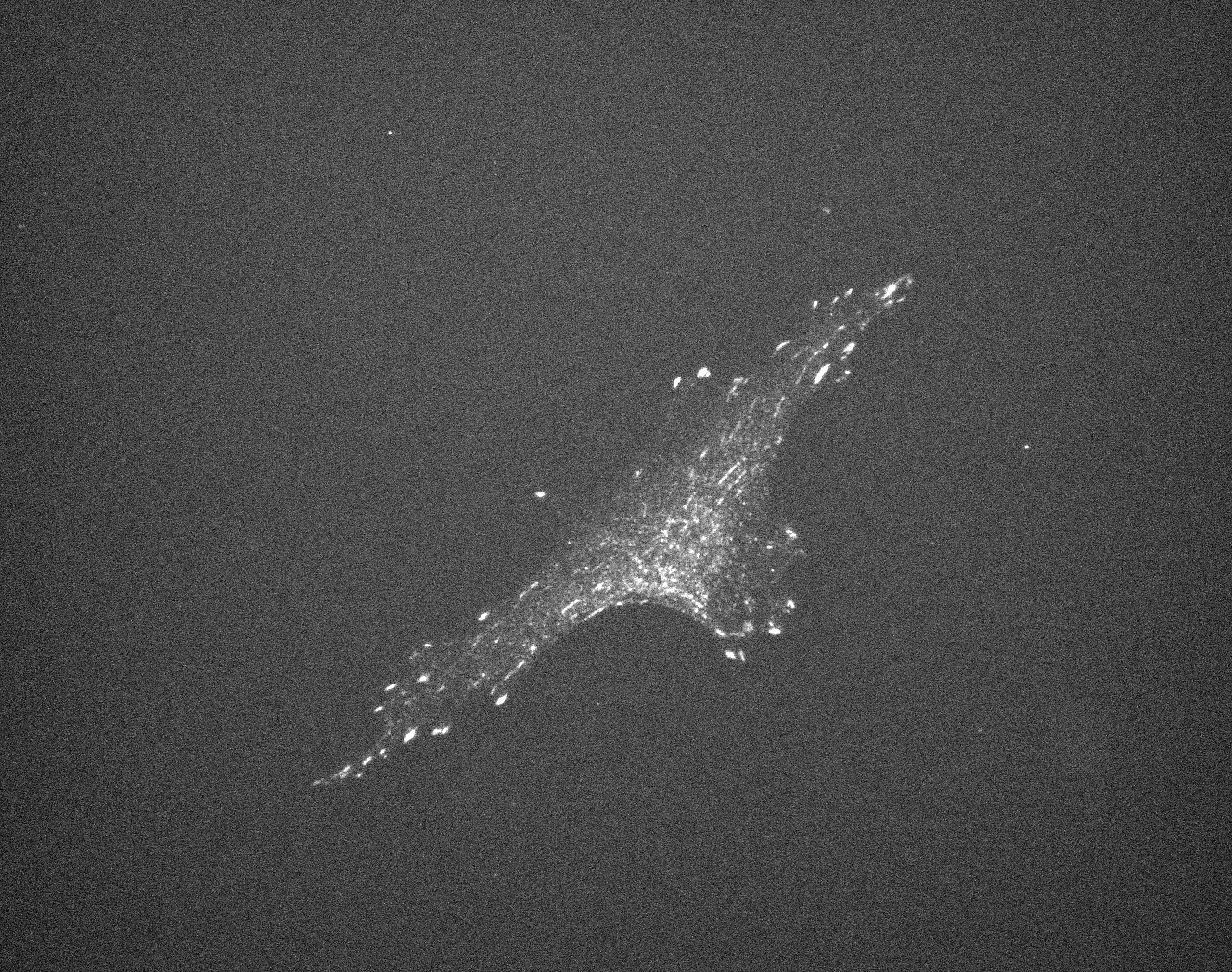

### F_Plate_R_p00_0_A01f37d0.TIF

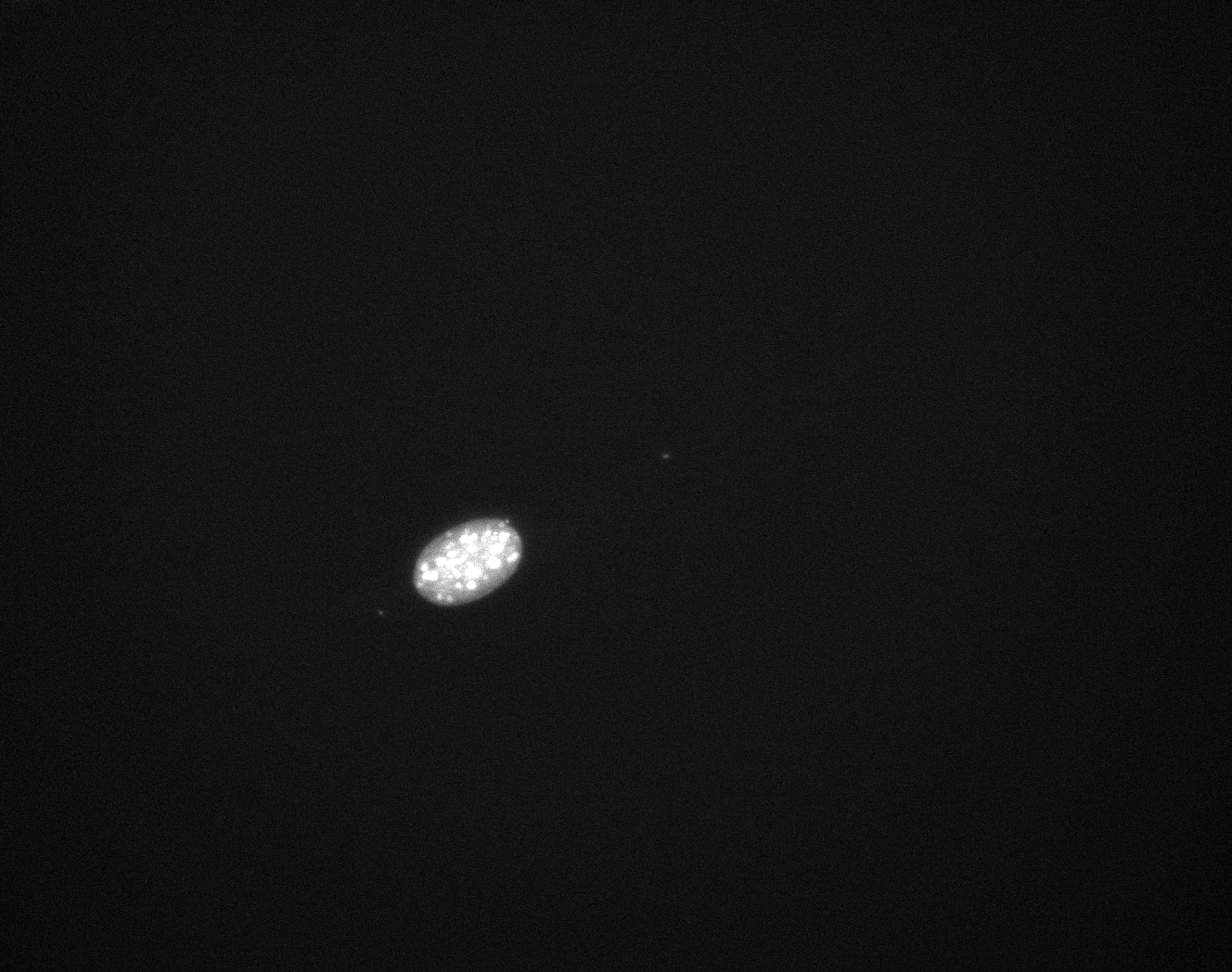

### F_Plate_R_p00_0_A01f37d1.TIF

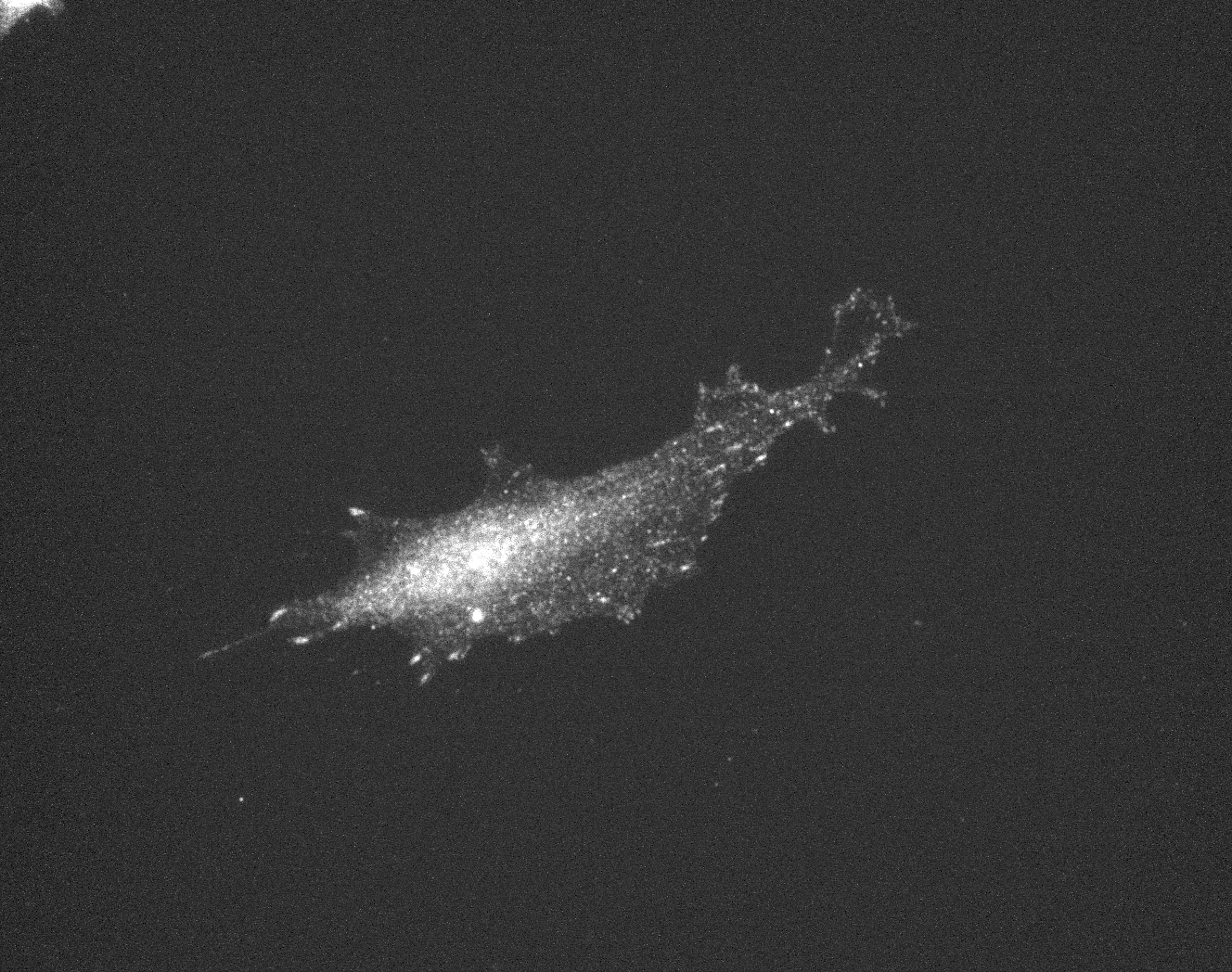

### F_Plate_R_p00_0_A01f37d2.TIF

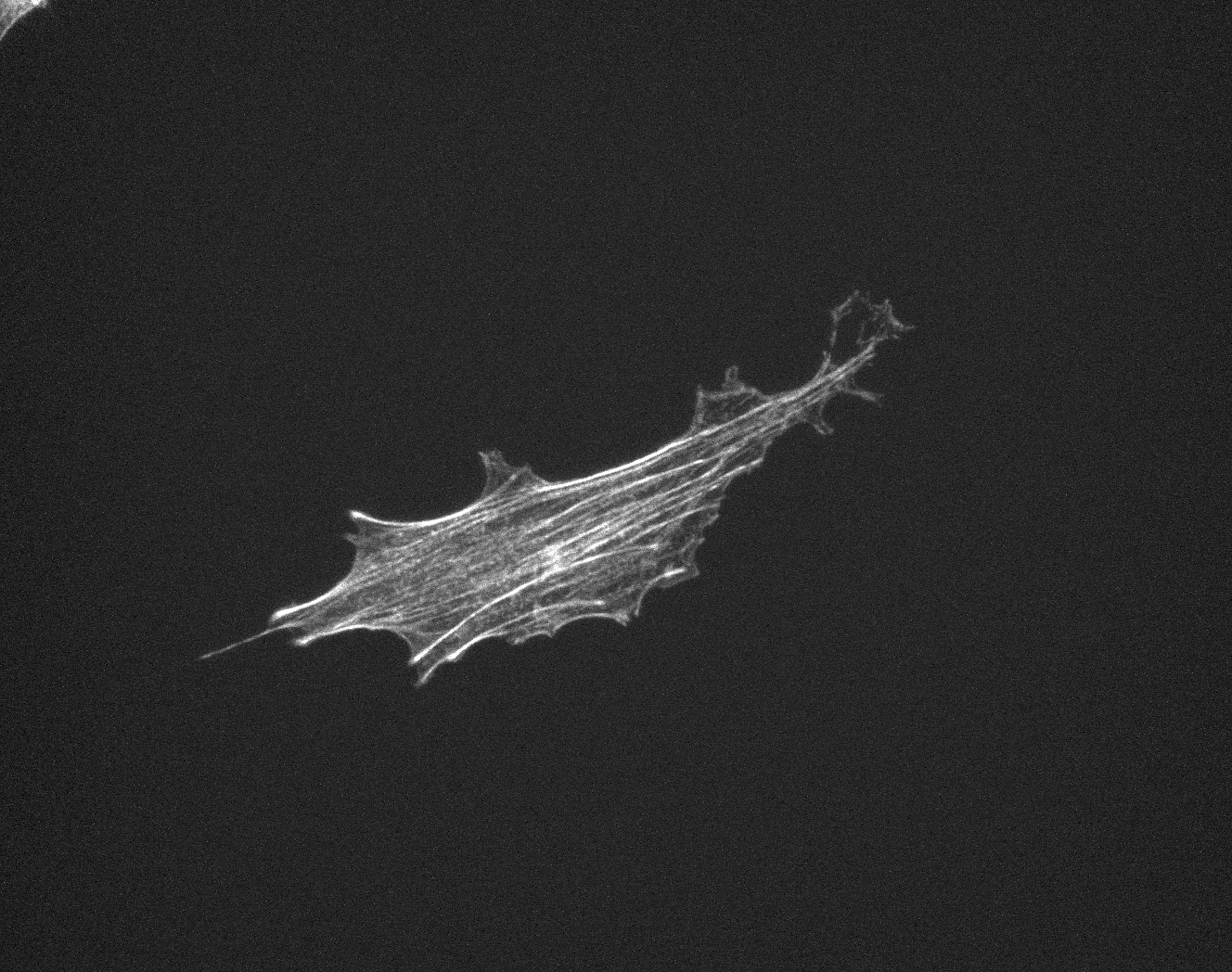

### F_Plate_R_p00_0_A01f37d3.TIF

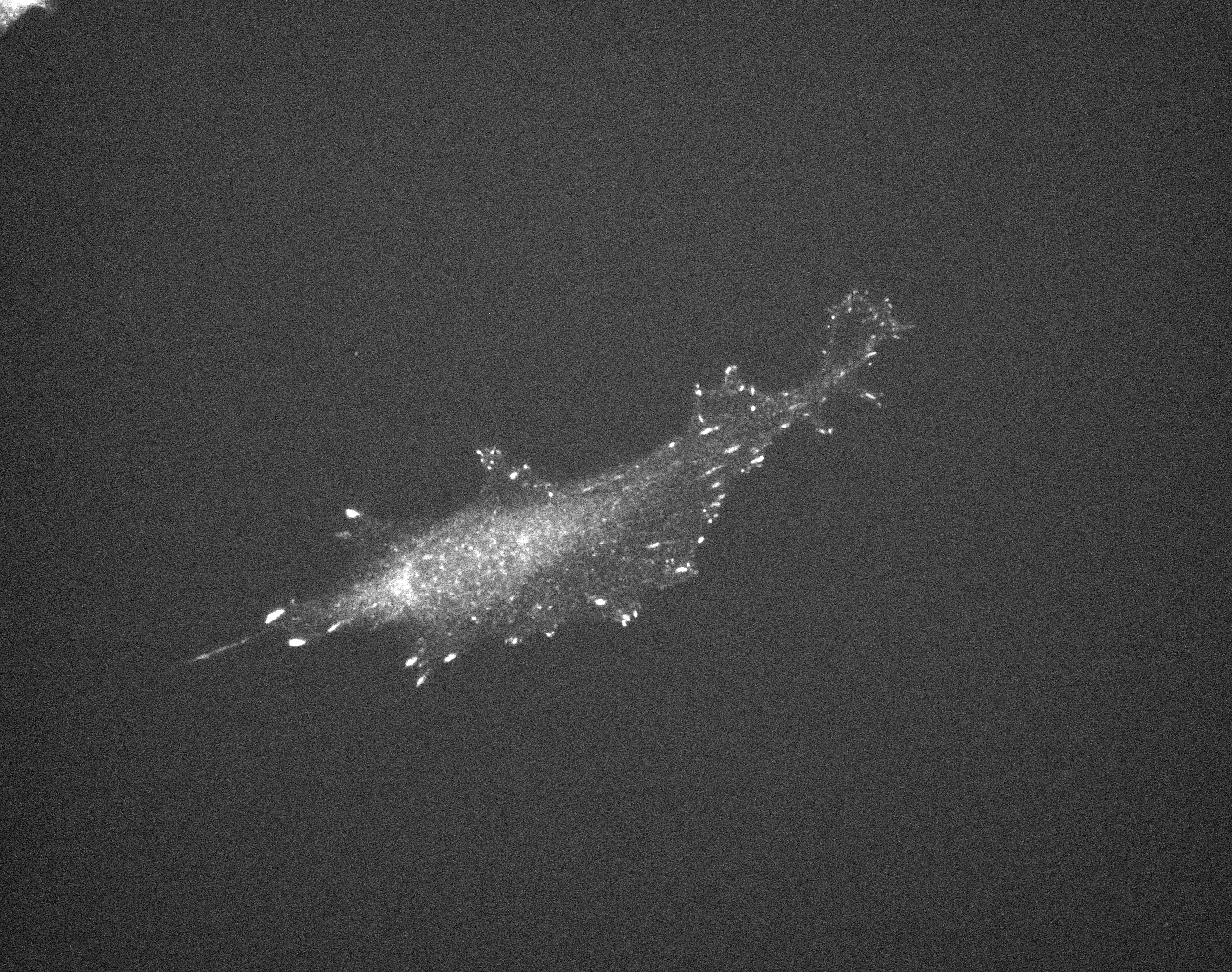

### fixedPipeline_mask_cell_A.jpg

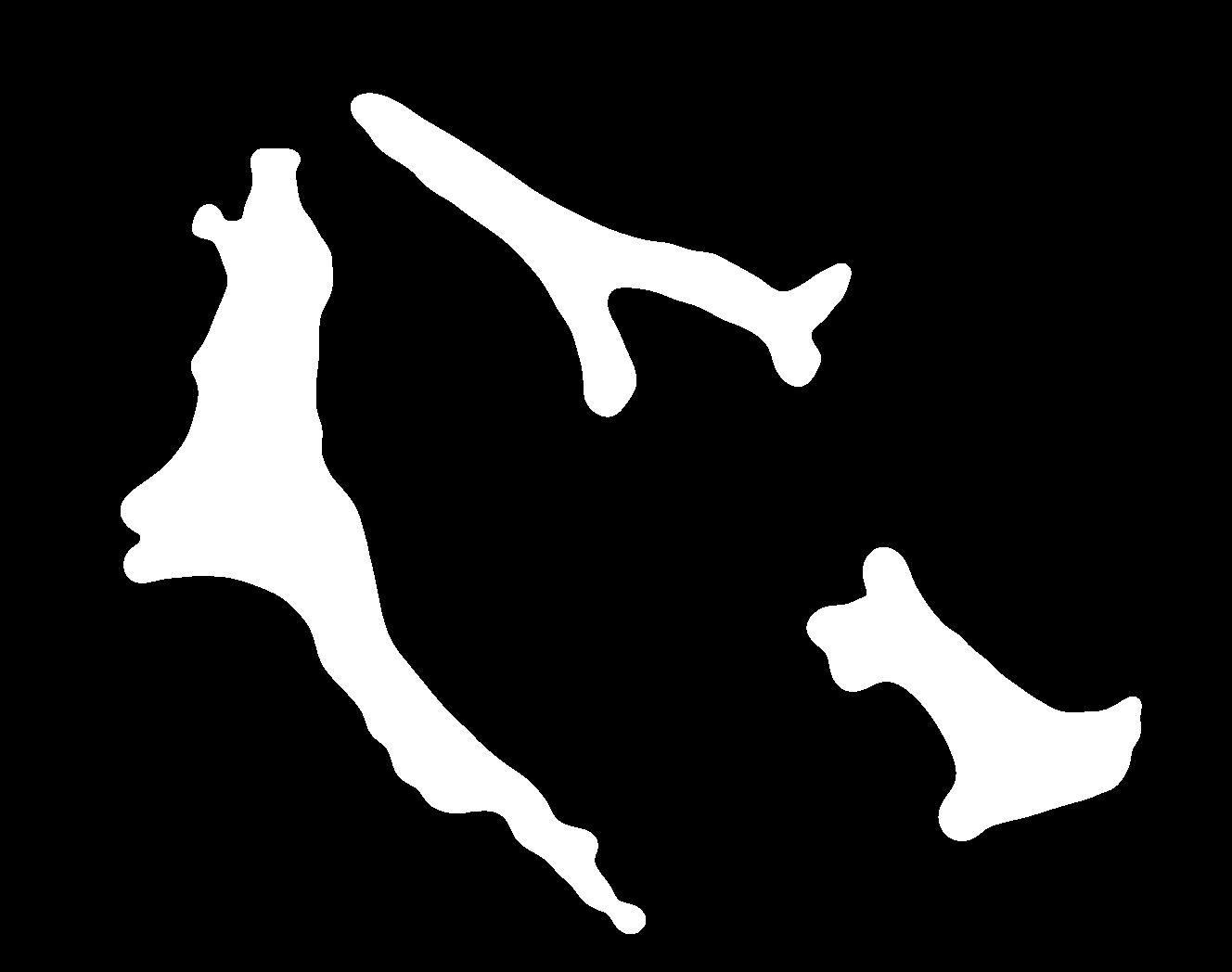

### fixedPipeline_mask_cell_B.jpg

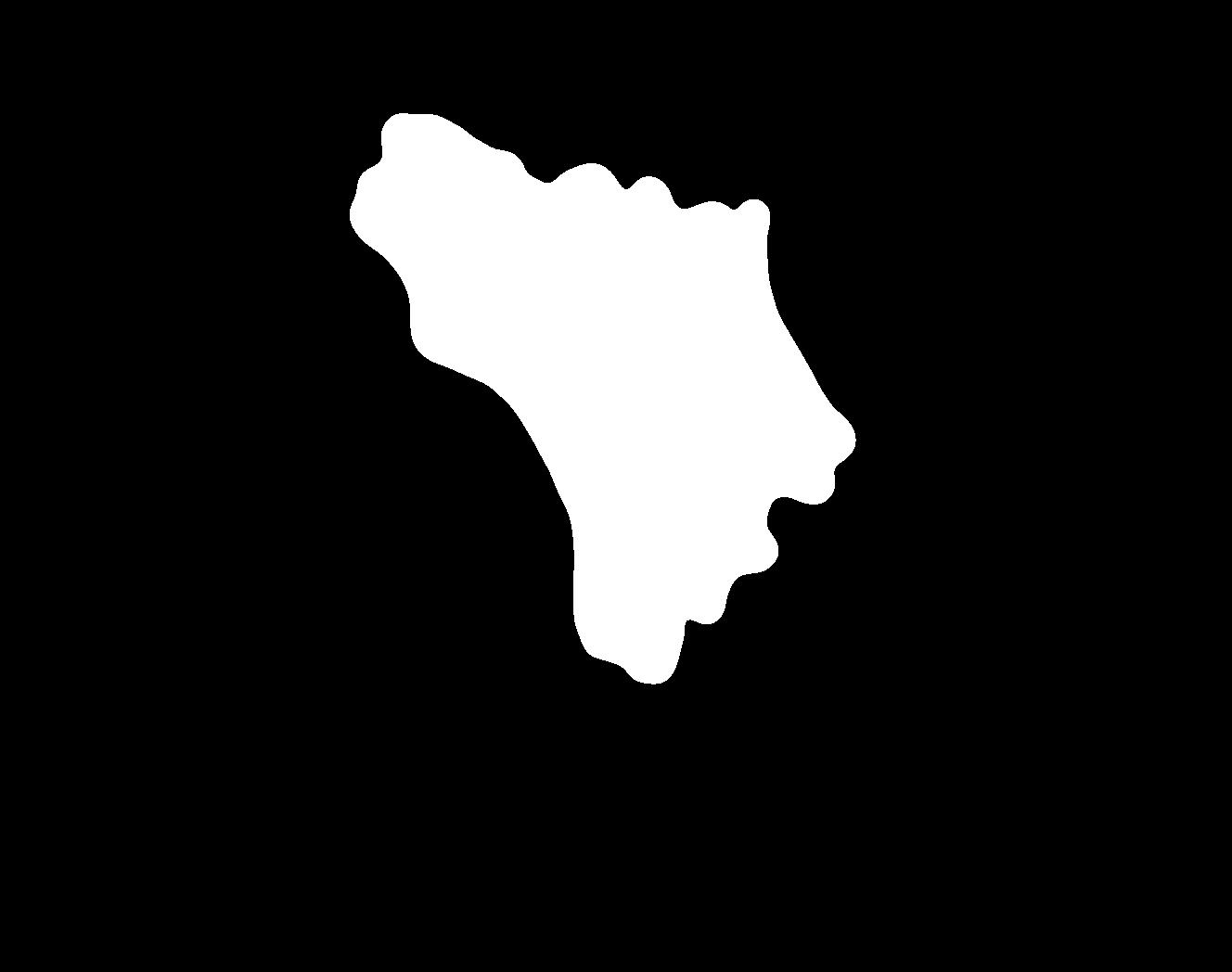

### fixedPipeline_mask_cell_C.jpg

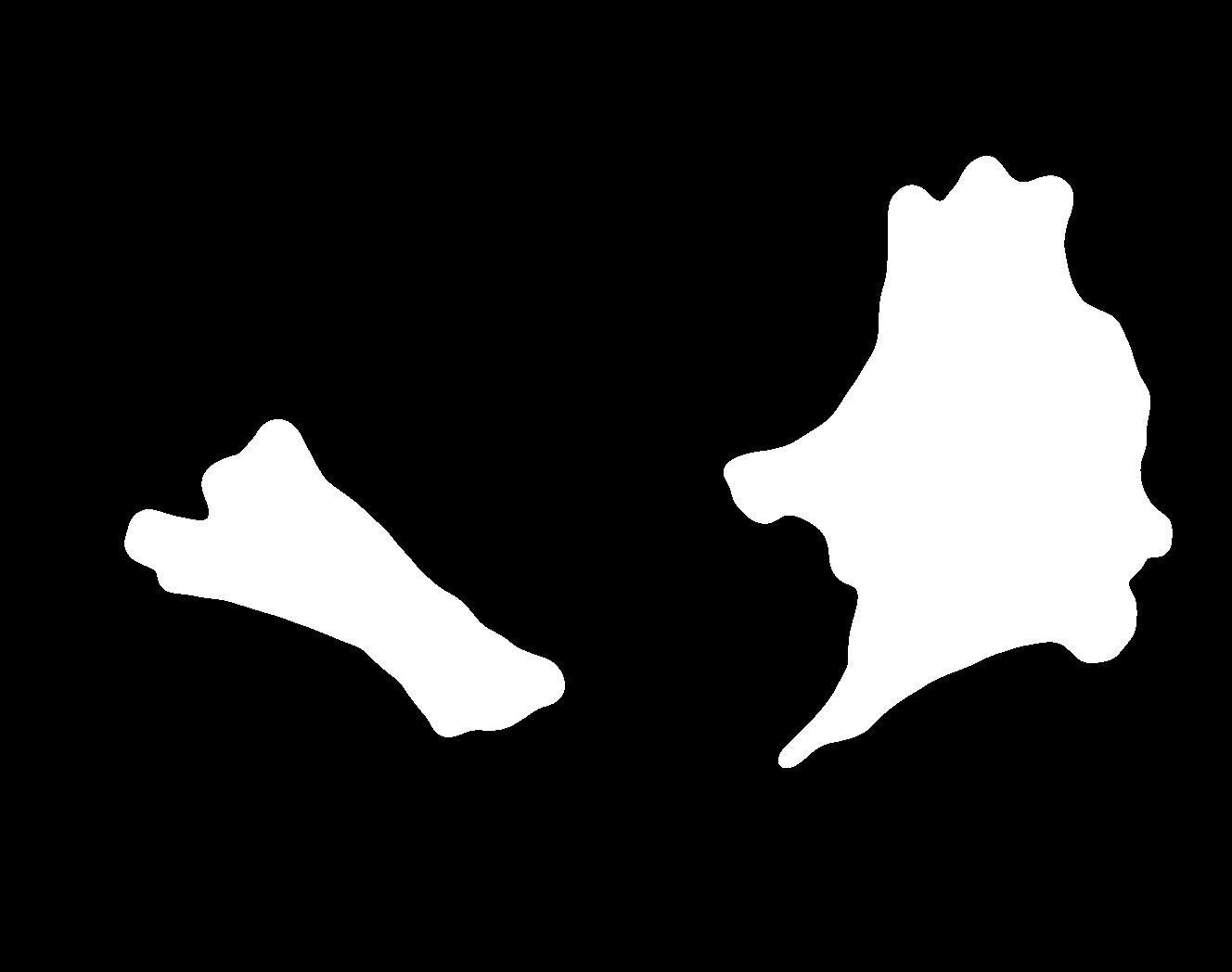

### fixedPipeline_mask_cell_D.jpg

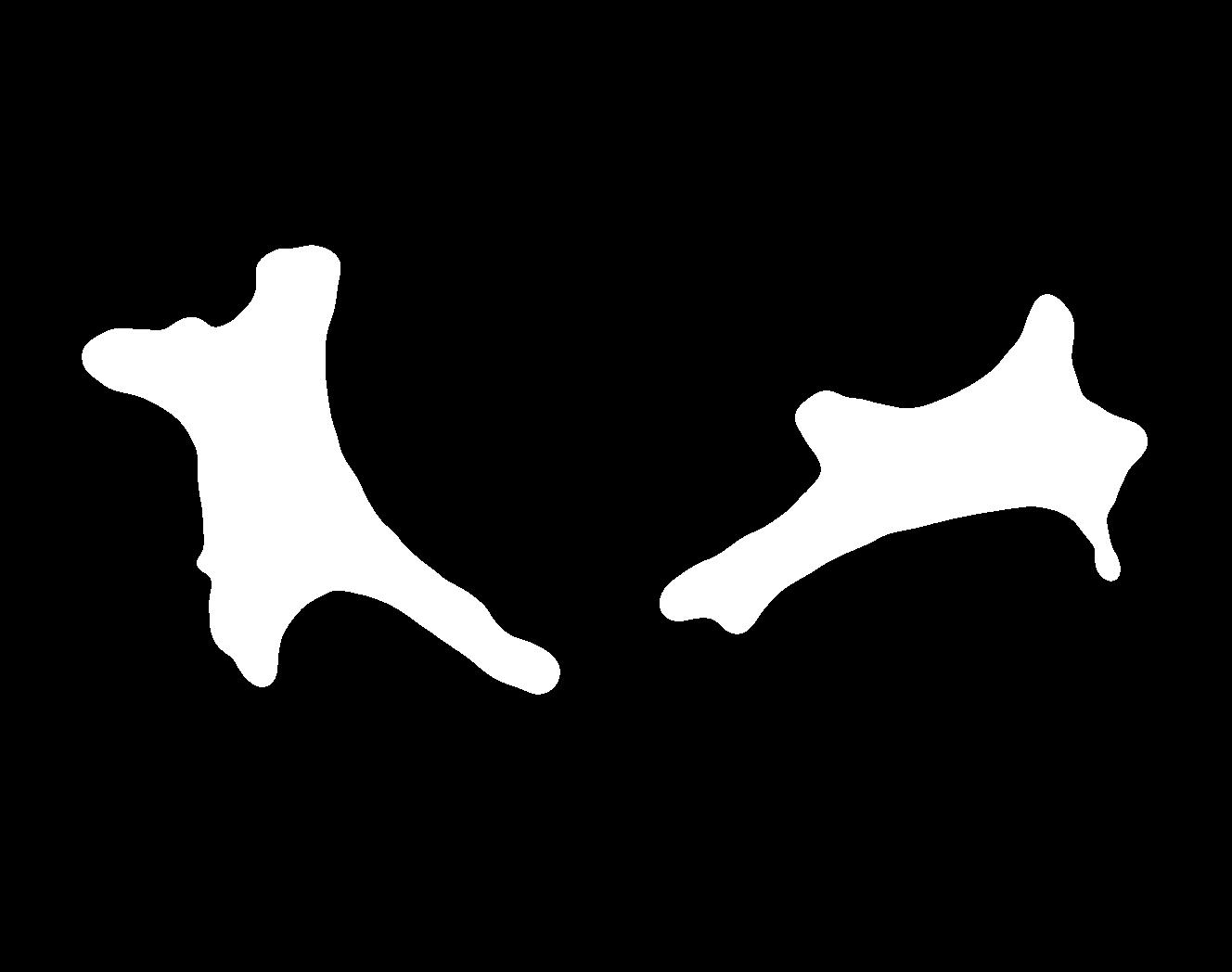

### fixedPipeline_mask_cell_E.jpg

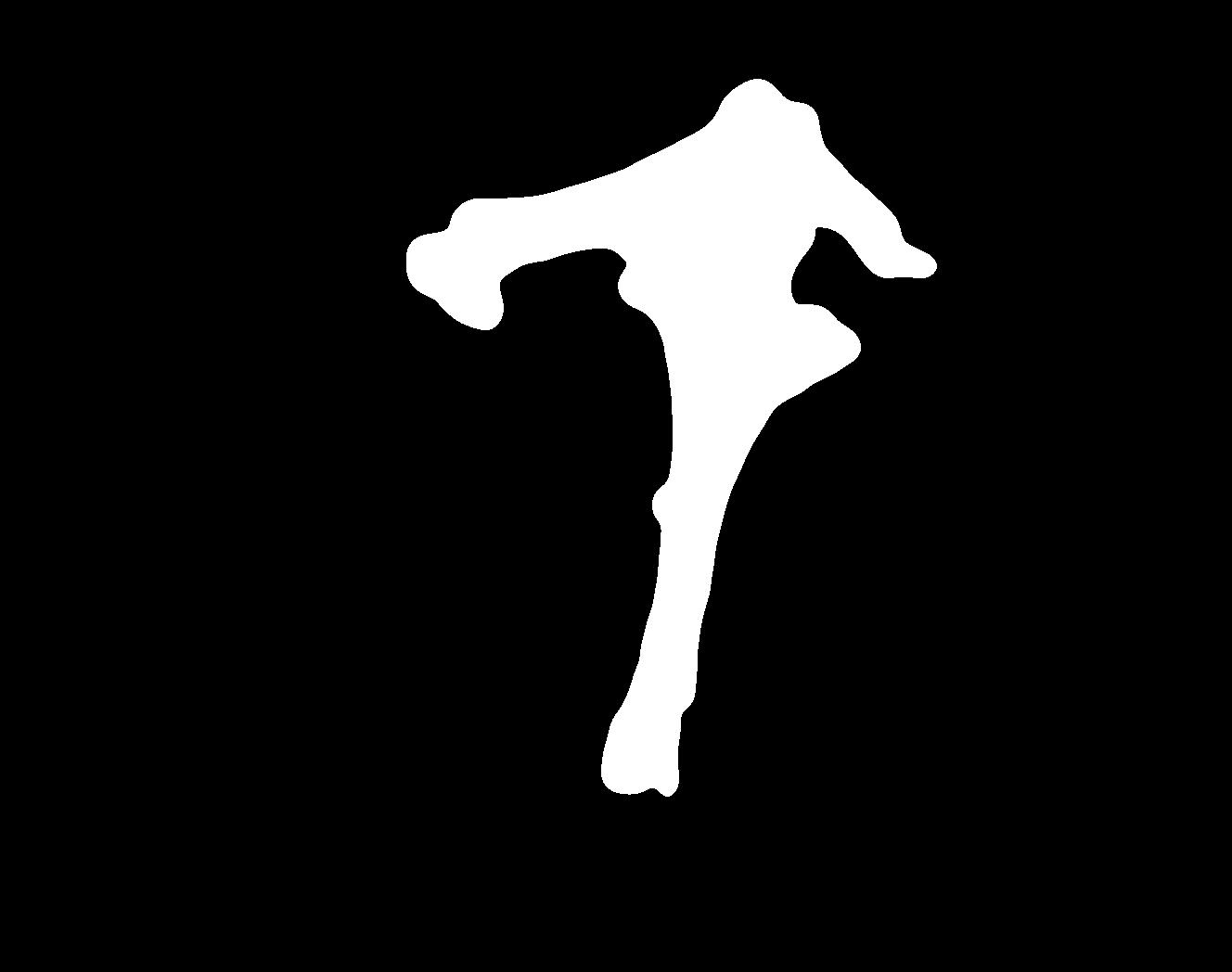

### fixedPipeline_mask_cell_F.jpg

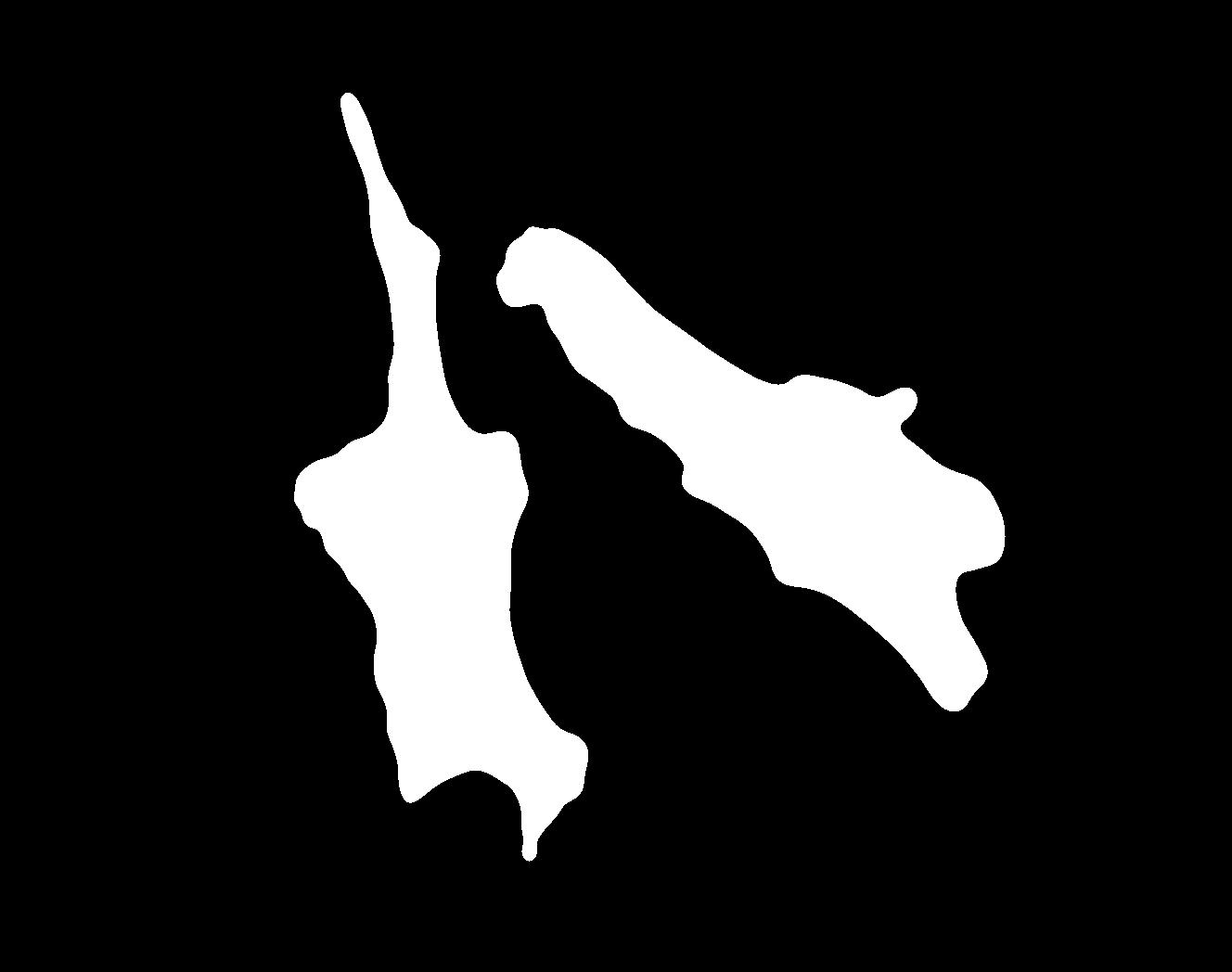

### fixedPipeline_mask_cell_G.jpg

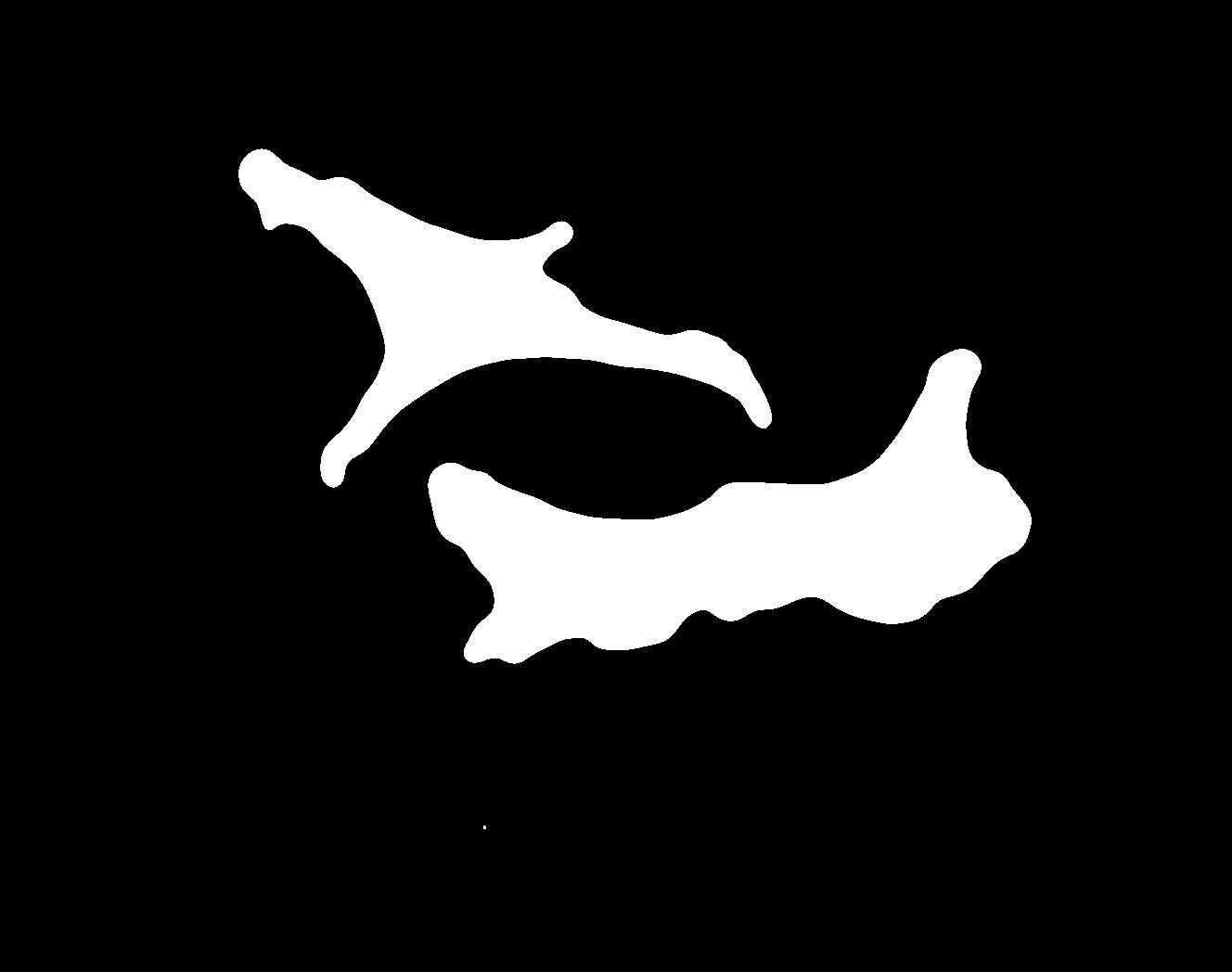

### fixedPipeline_mask_cell_H.jpg

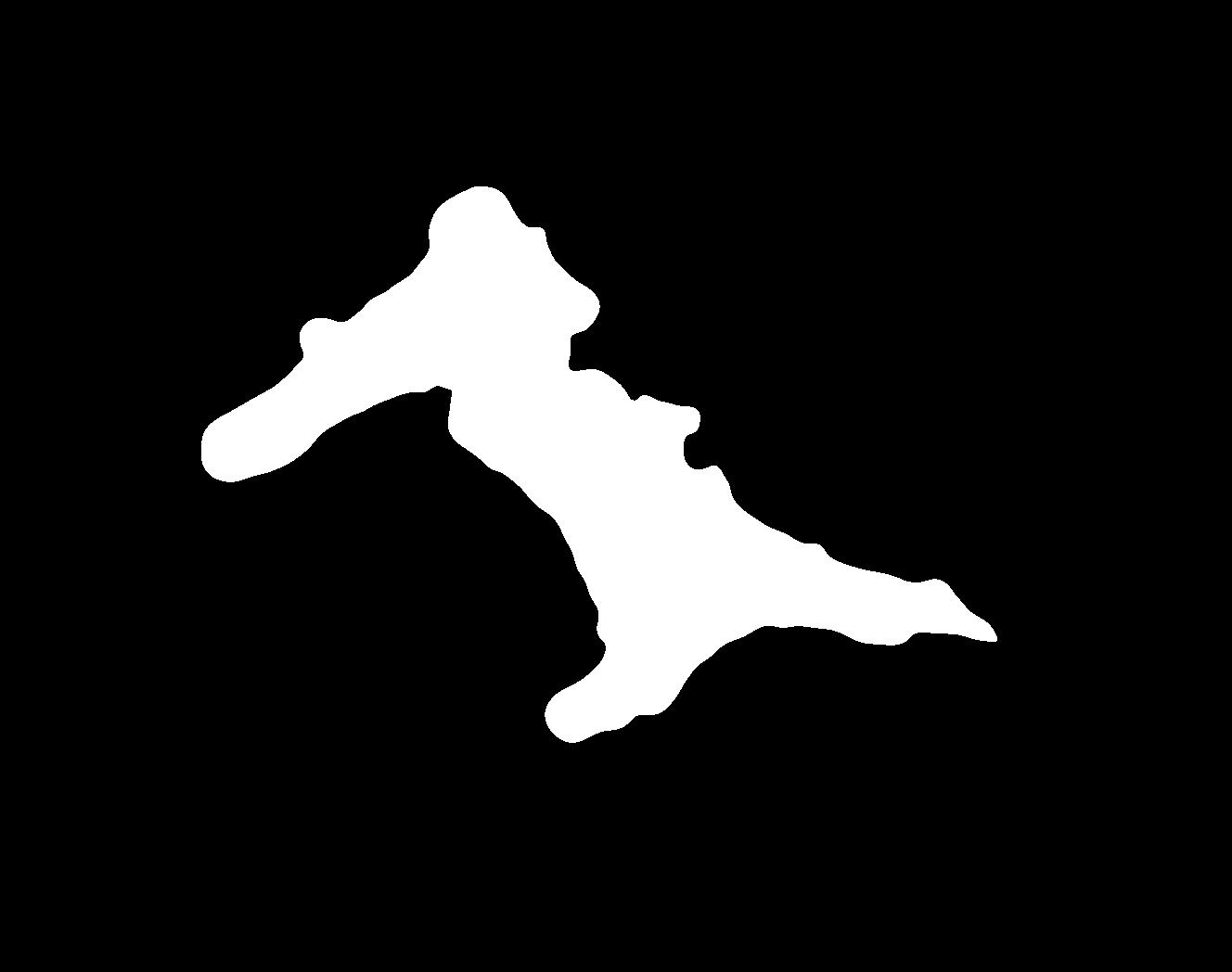

### fixedPipeline_mask_cell_I.jpg

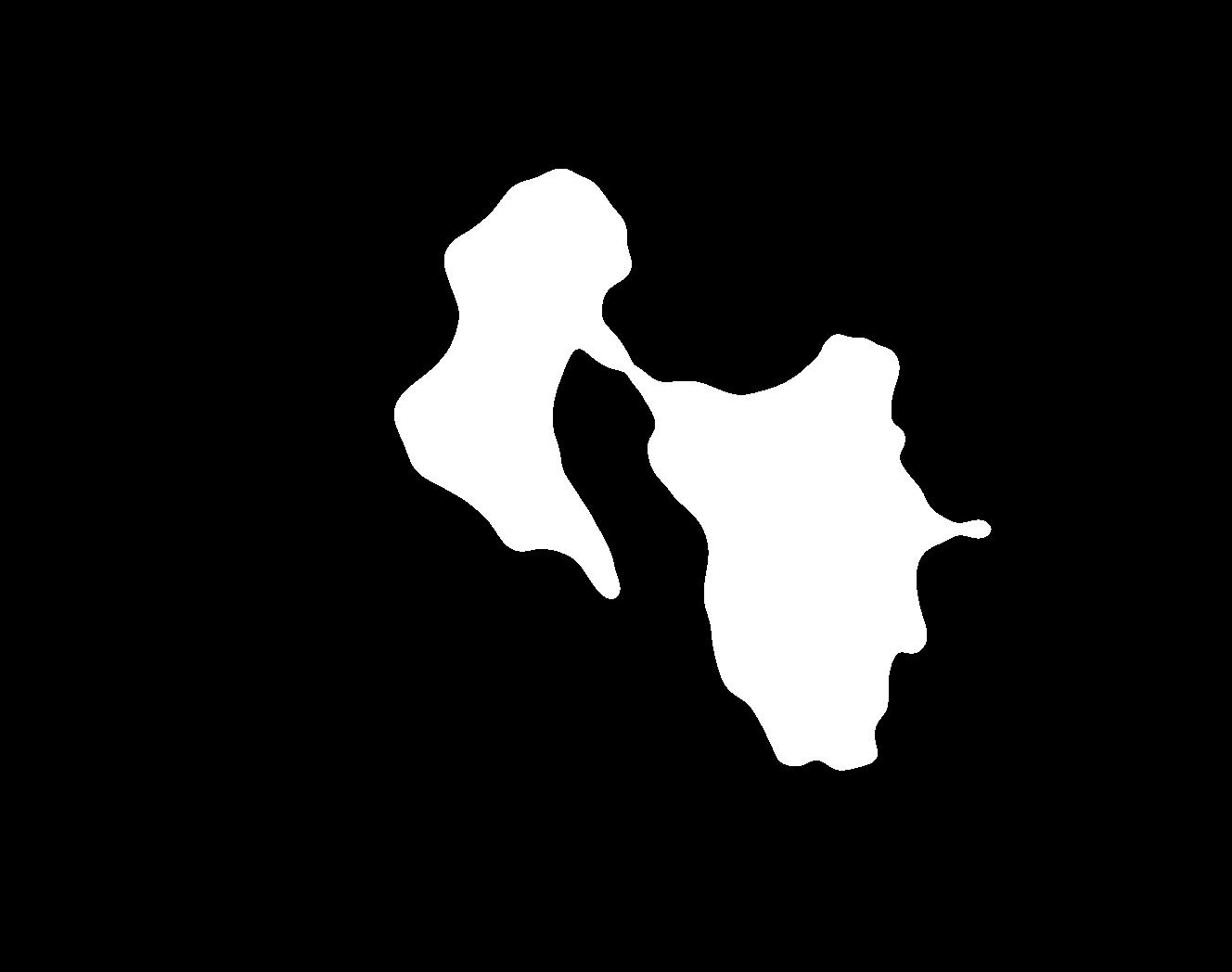

### fixedPipeline_mask_cell_J.jpg

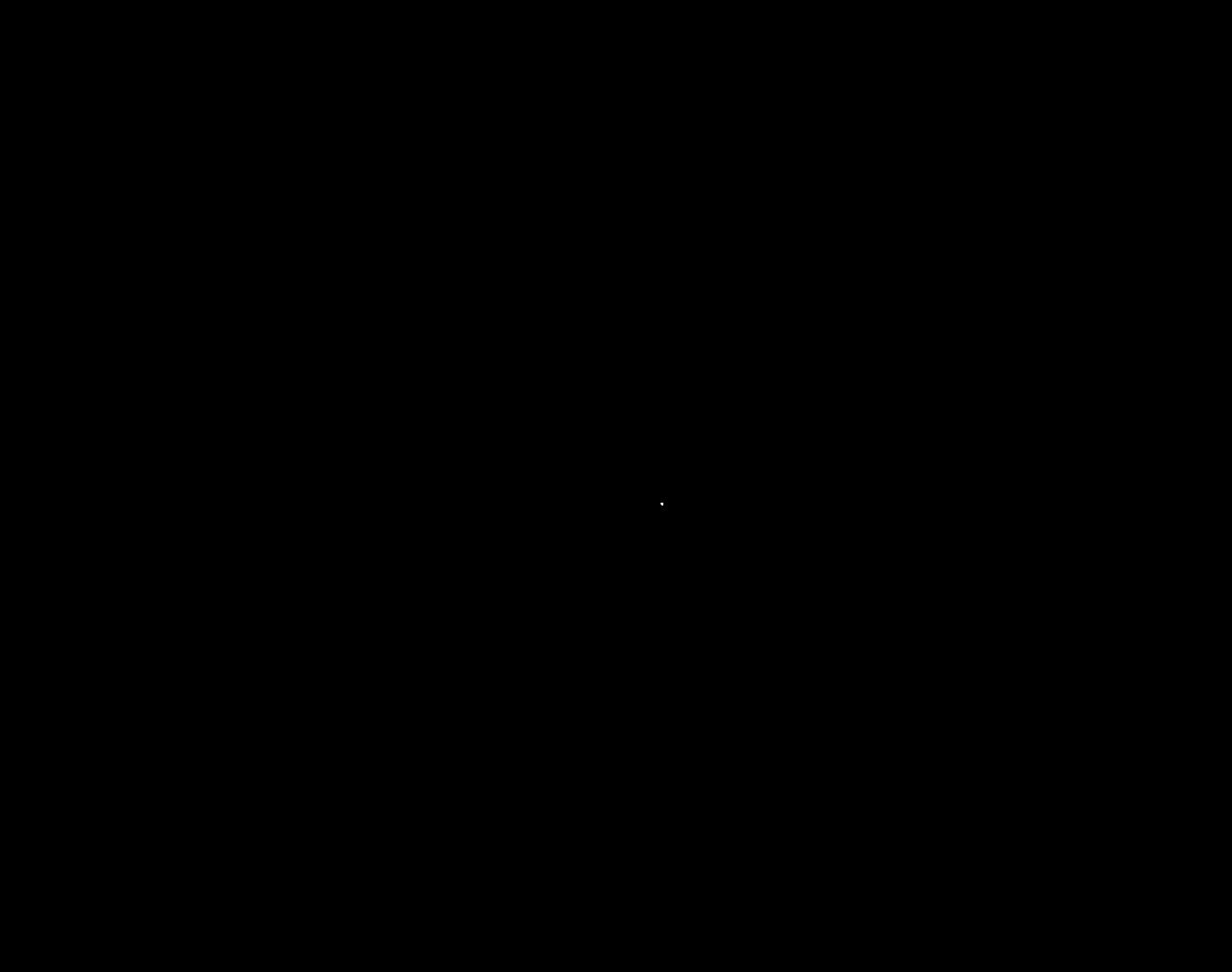

### G_groundtruth.jpg

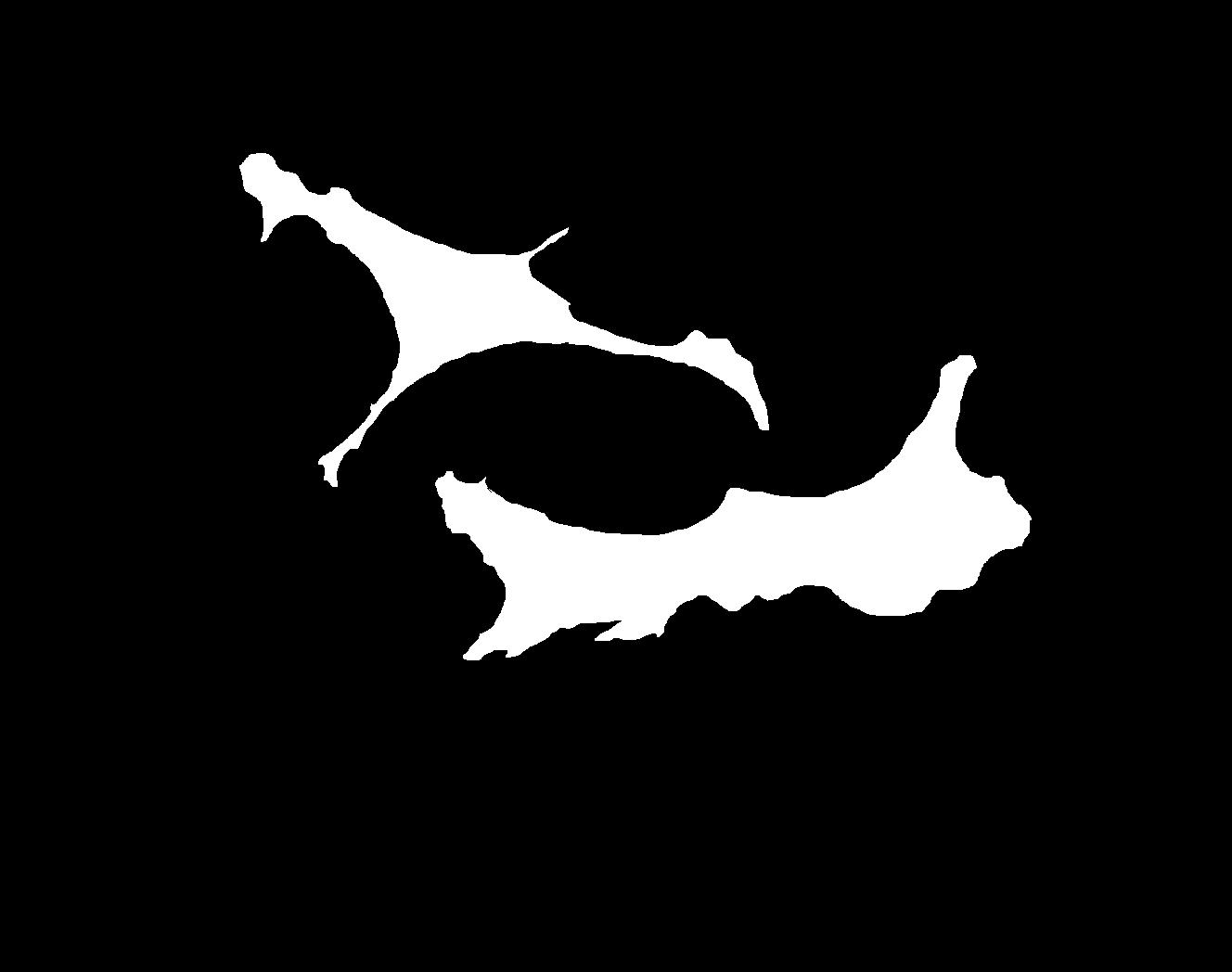

### G_Plate_R_p00_0_A01f25d0.TIF

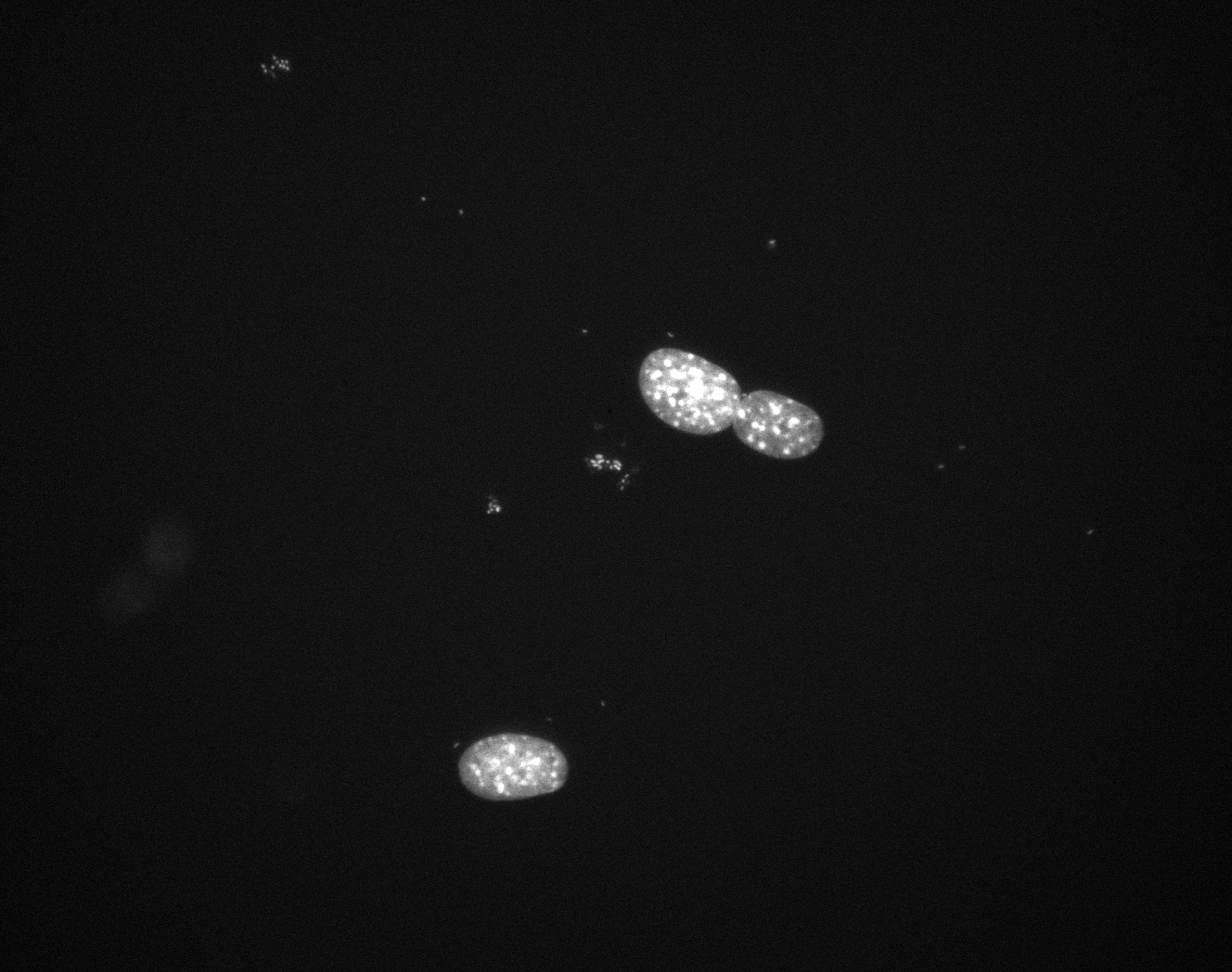

### G_Plate_R_p00_0_A01f25d1.TIF

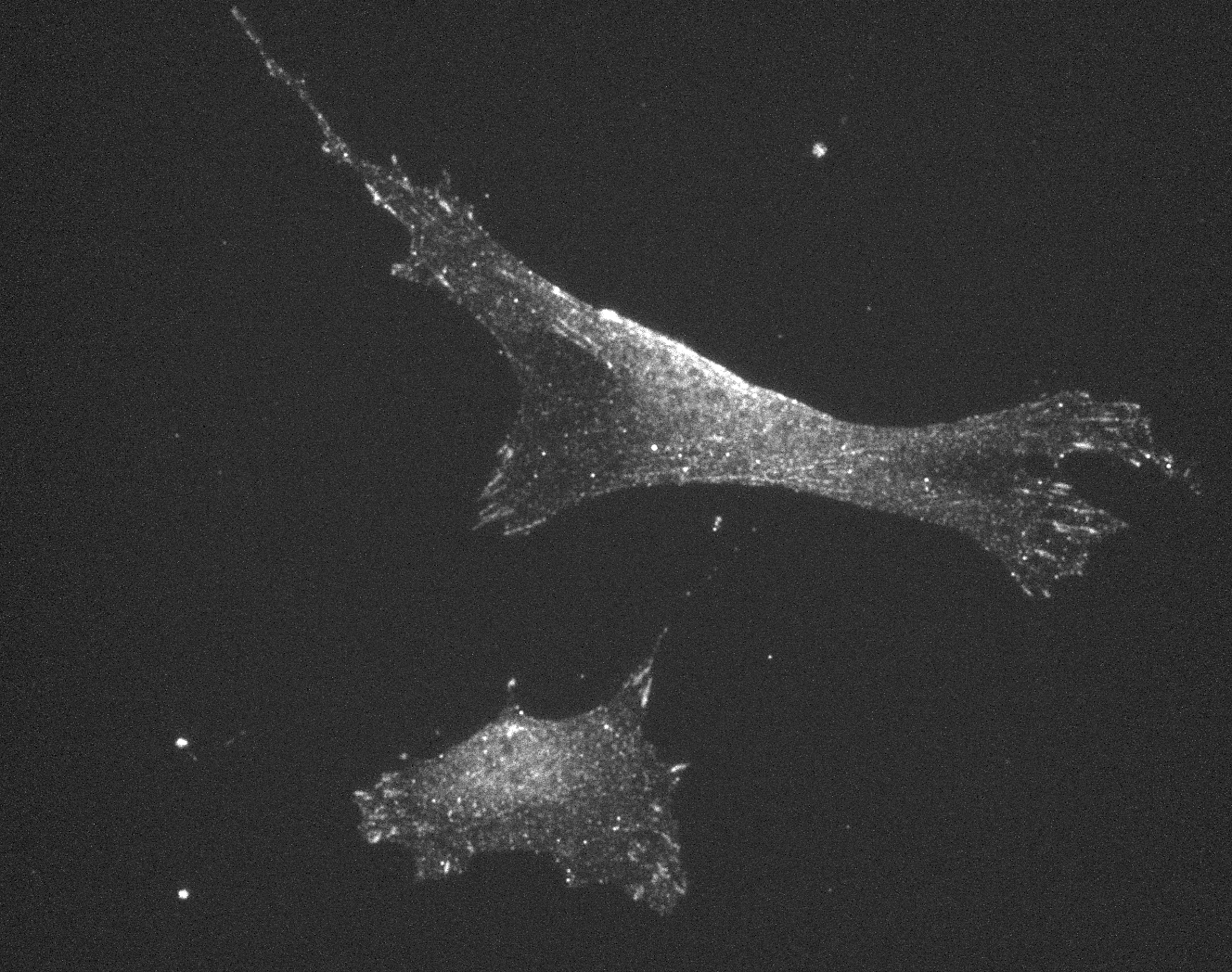

### G_Plate_R_p00_0_A01f25d2.TIF

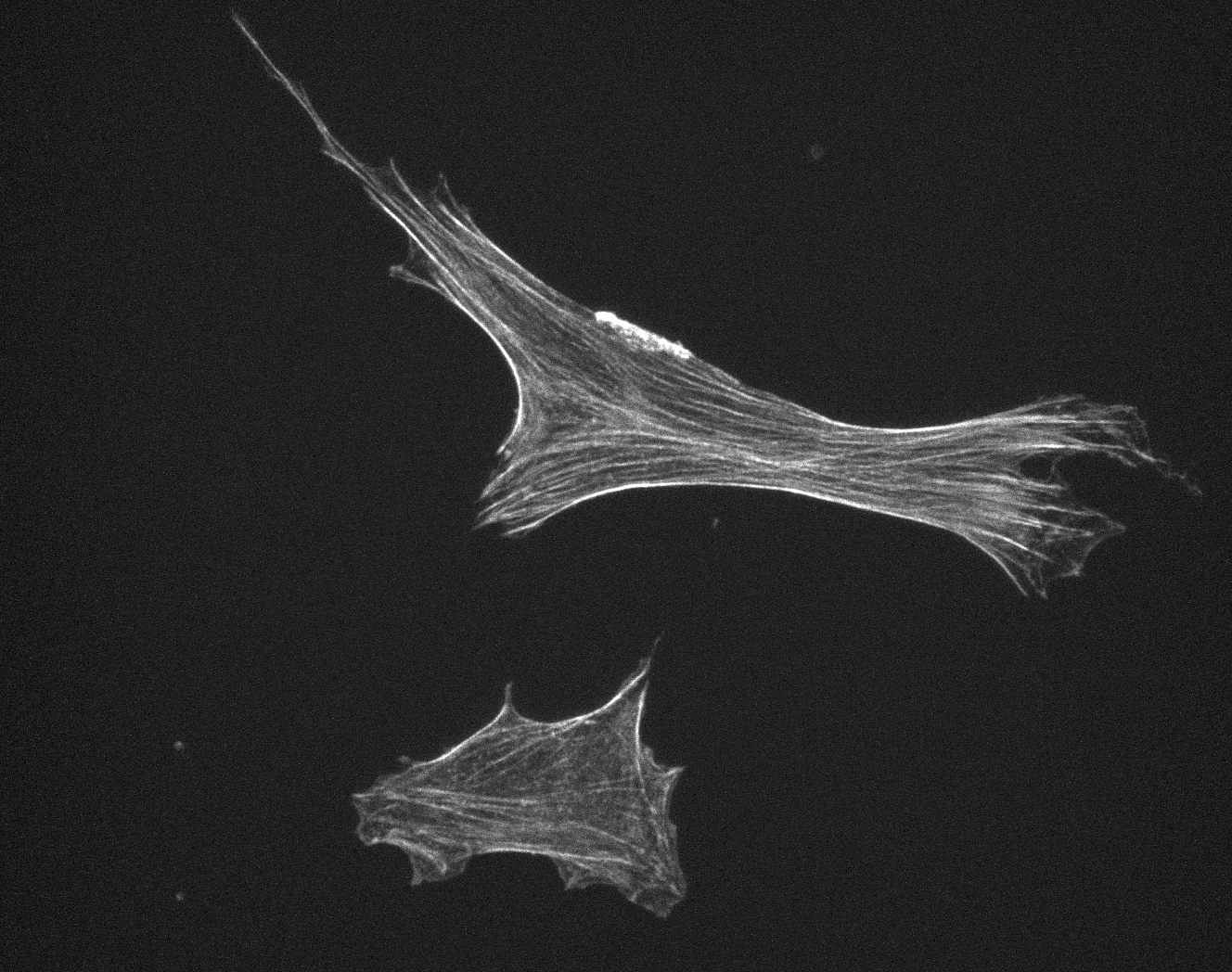

### G_Plate_R_p00_0_A01f25d3.TIF

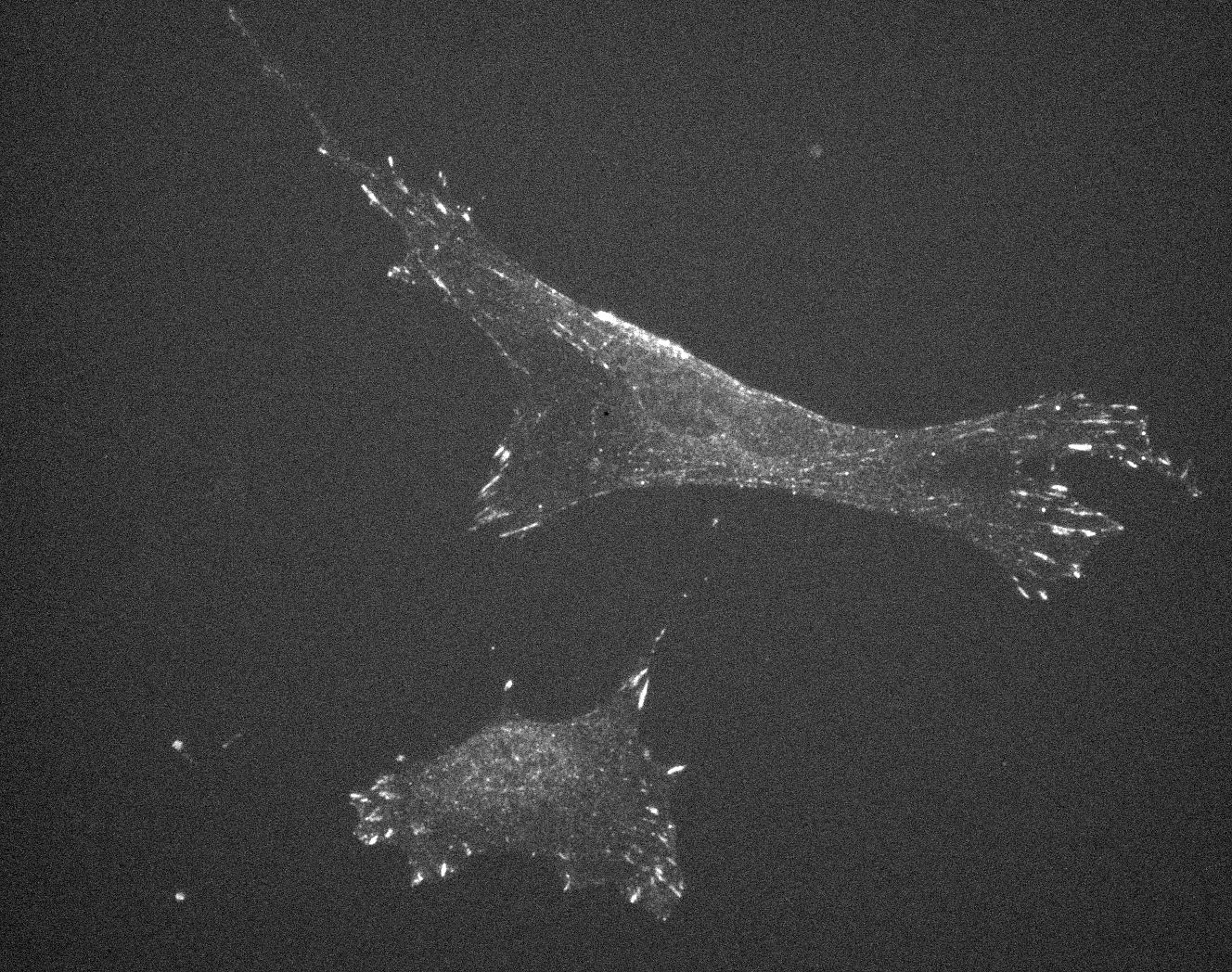

### G_Plate_R_p00_0_A01f26d0.TIF

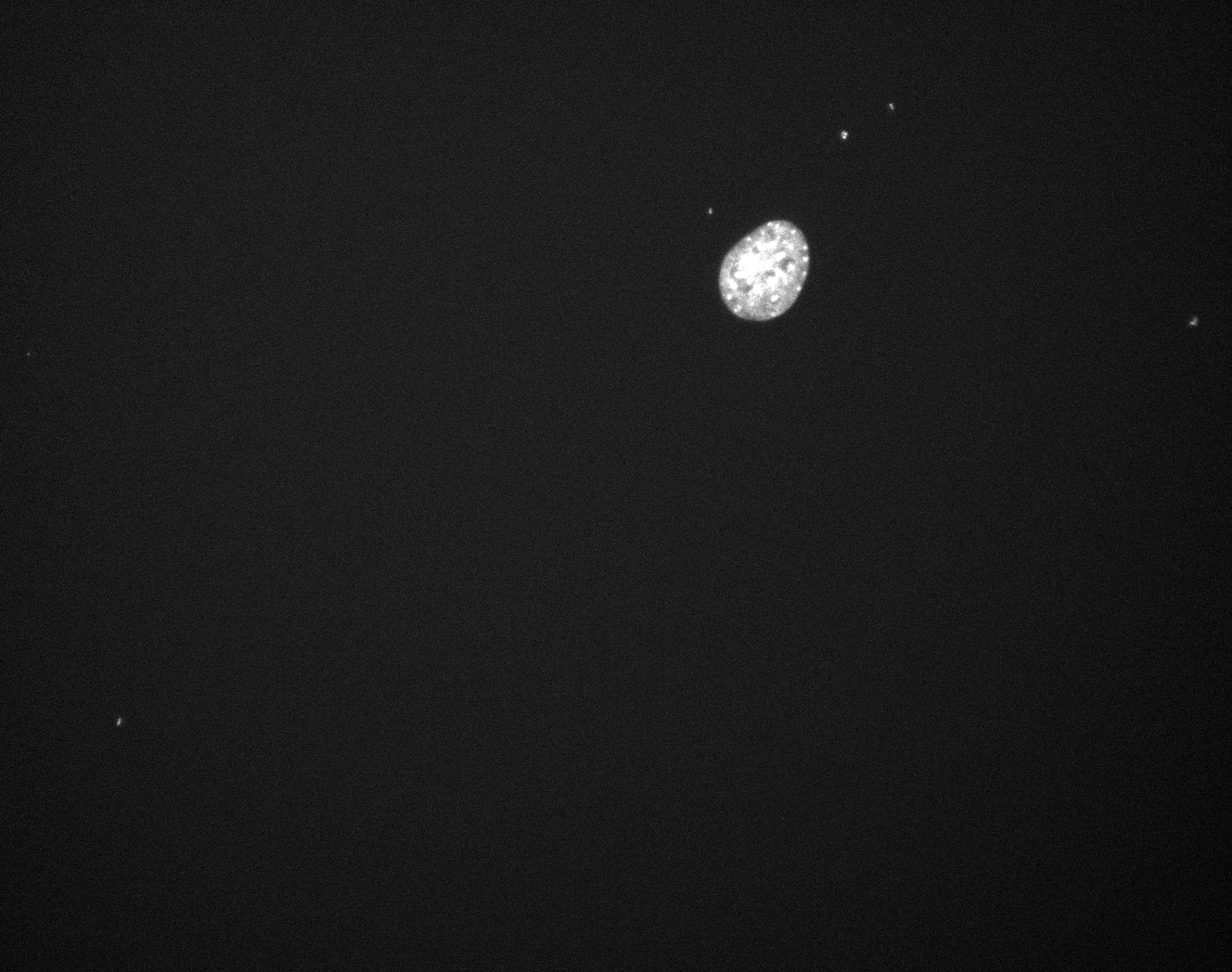

### G_Plate_R_p00_0_A01f26d1.TIF

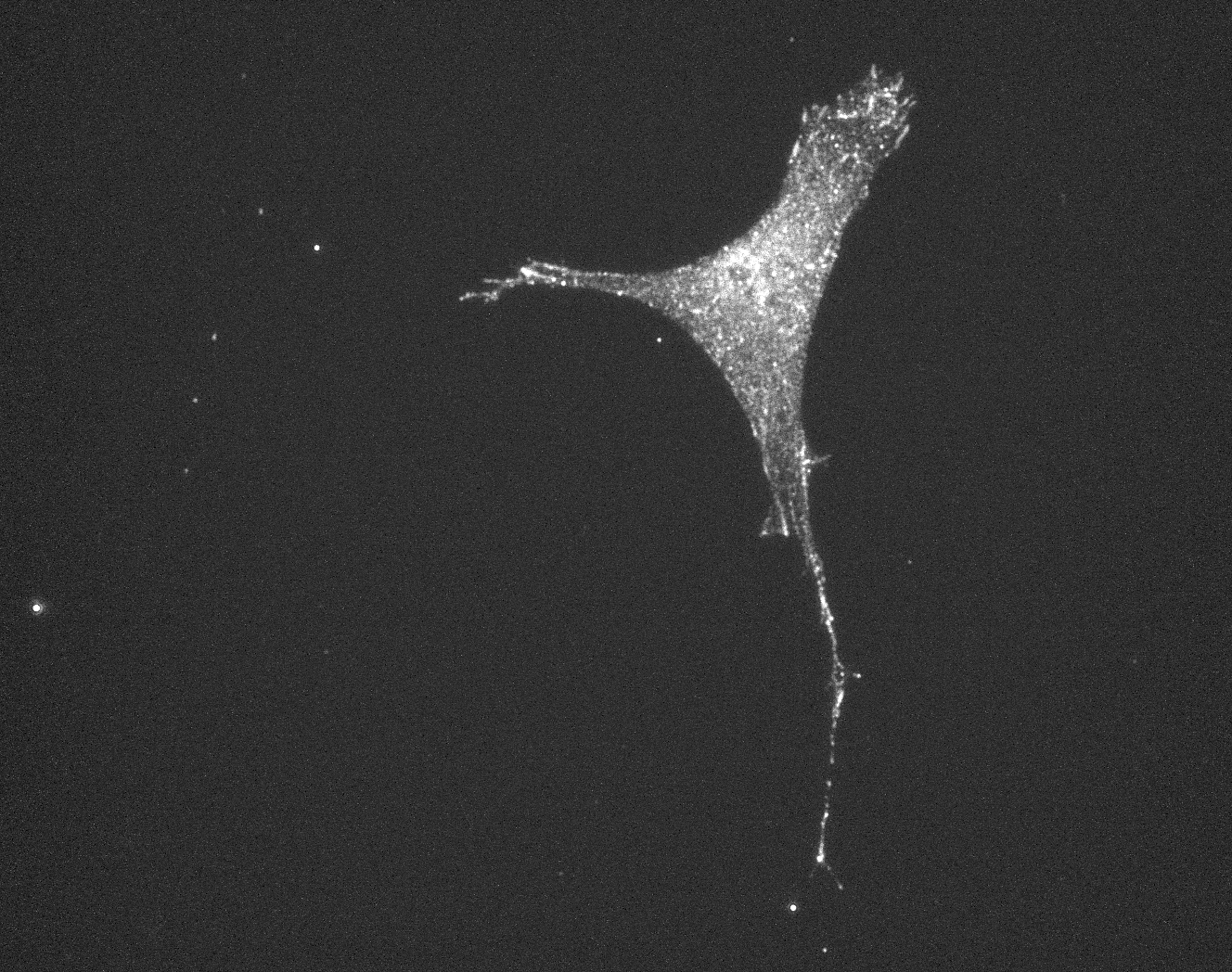

### G_Plate_R_p00_0_A01f26d2.TIF

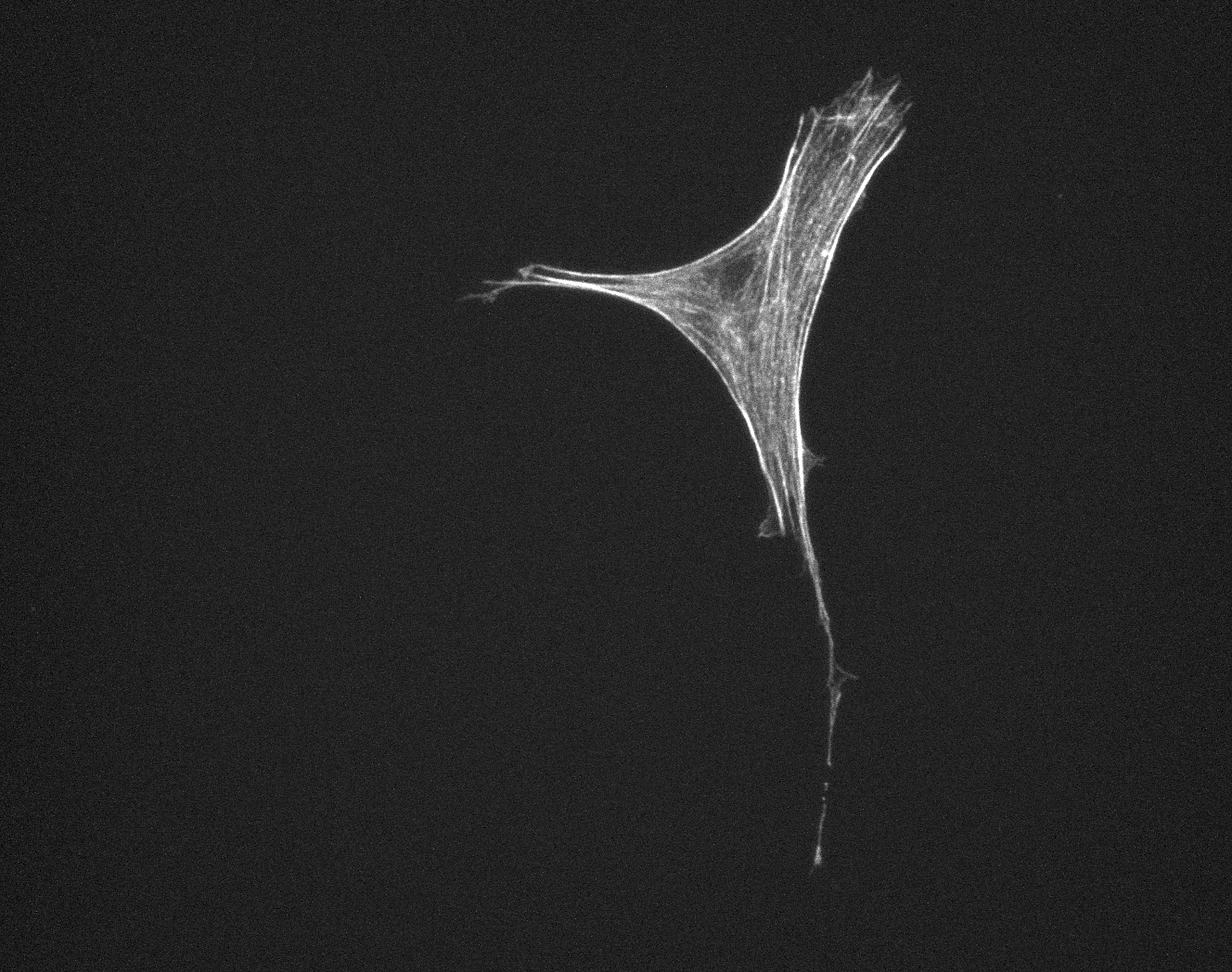

### G_Plate_R_p00_0_A01f26d3.TIF

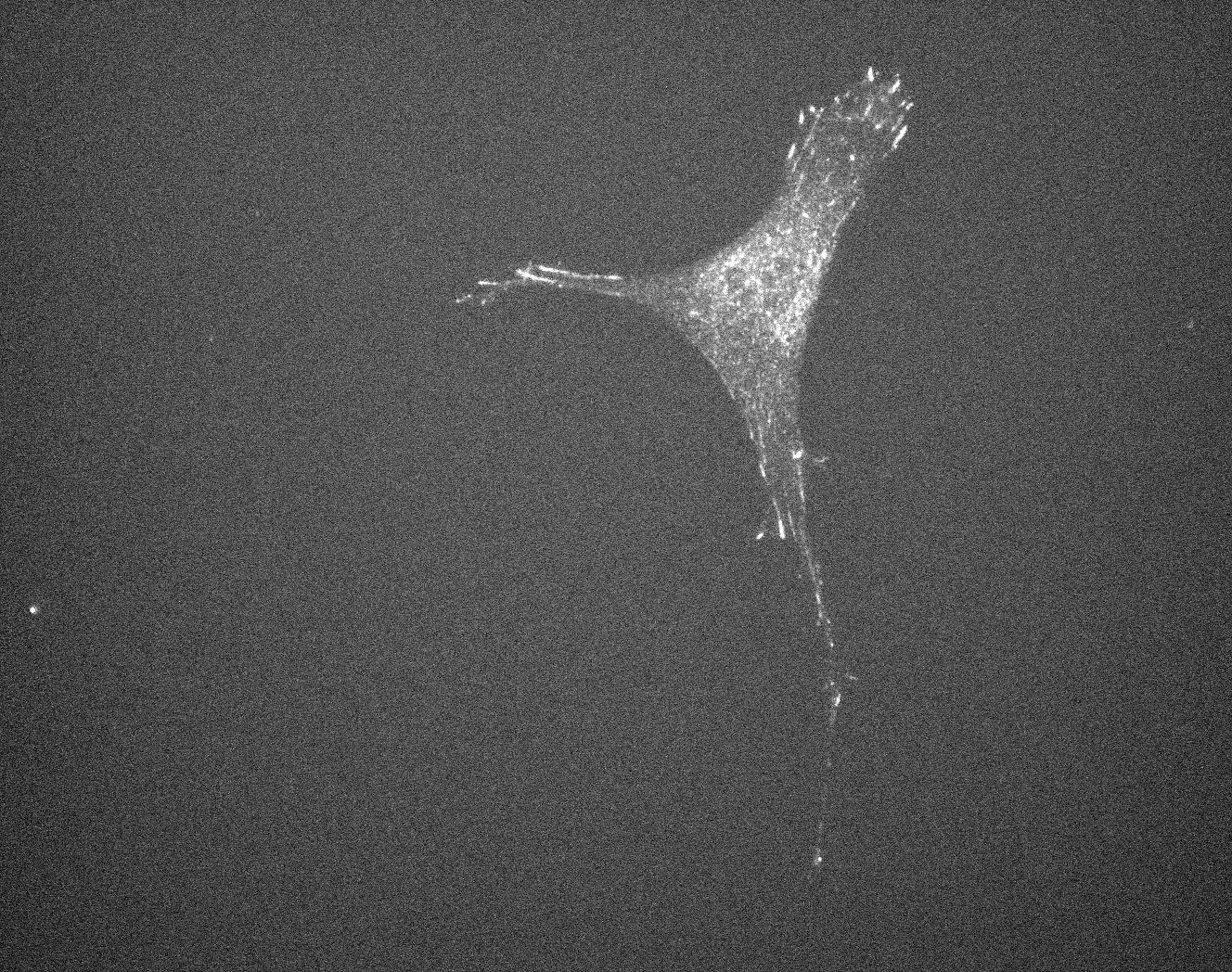

### H_groundtruth.jpg

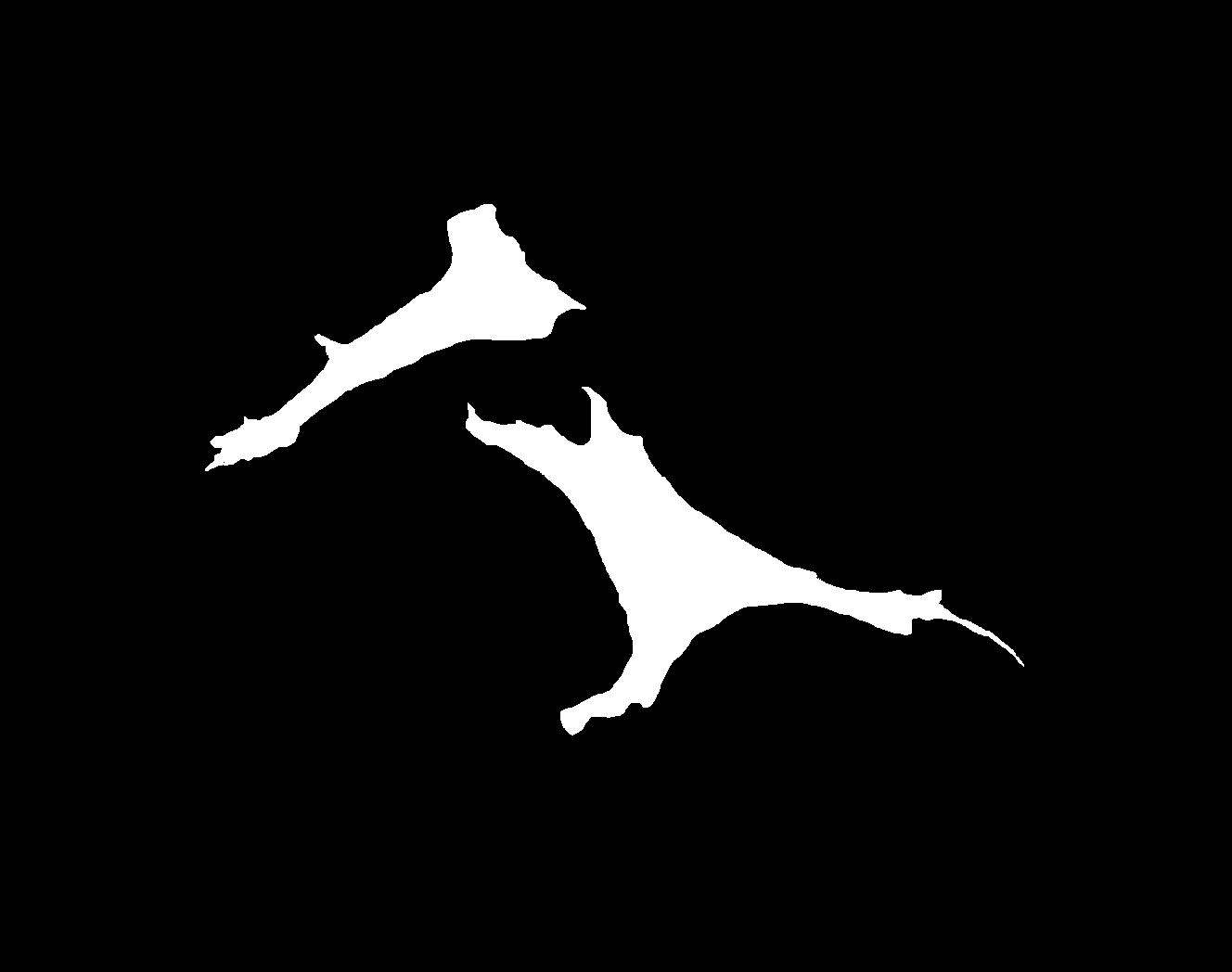

### H_Plate_R_p00_0_A01f41d0.TIF

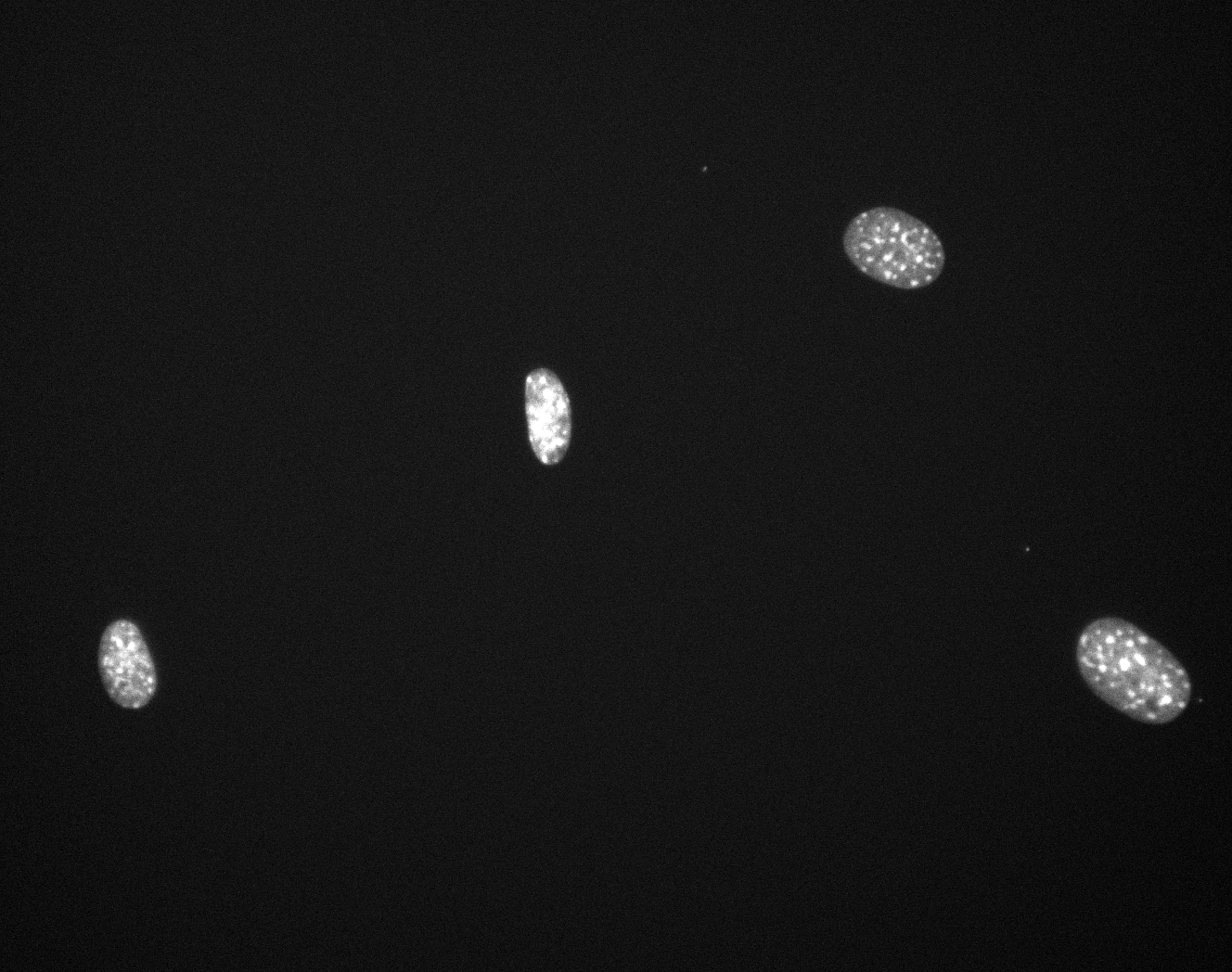

### H_Plate_R_p00_0_A01f41d1.TIF

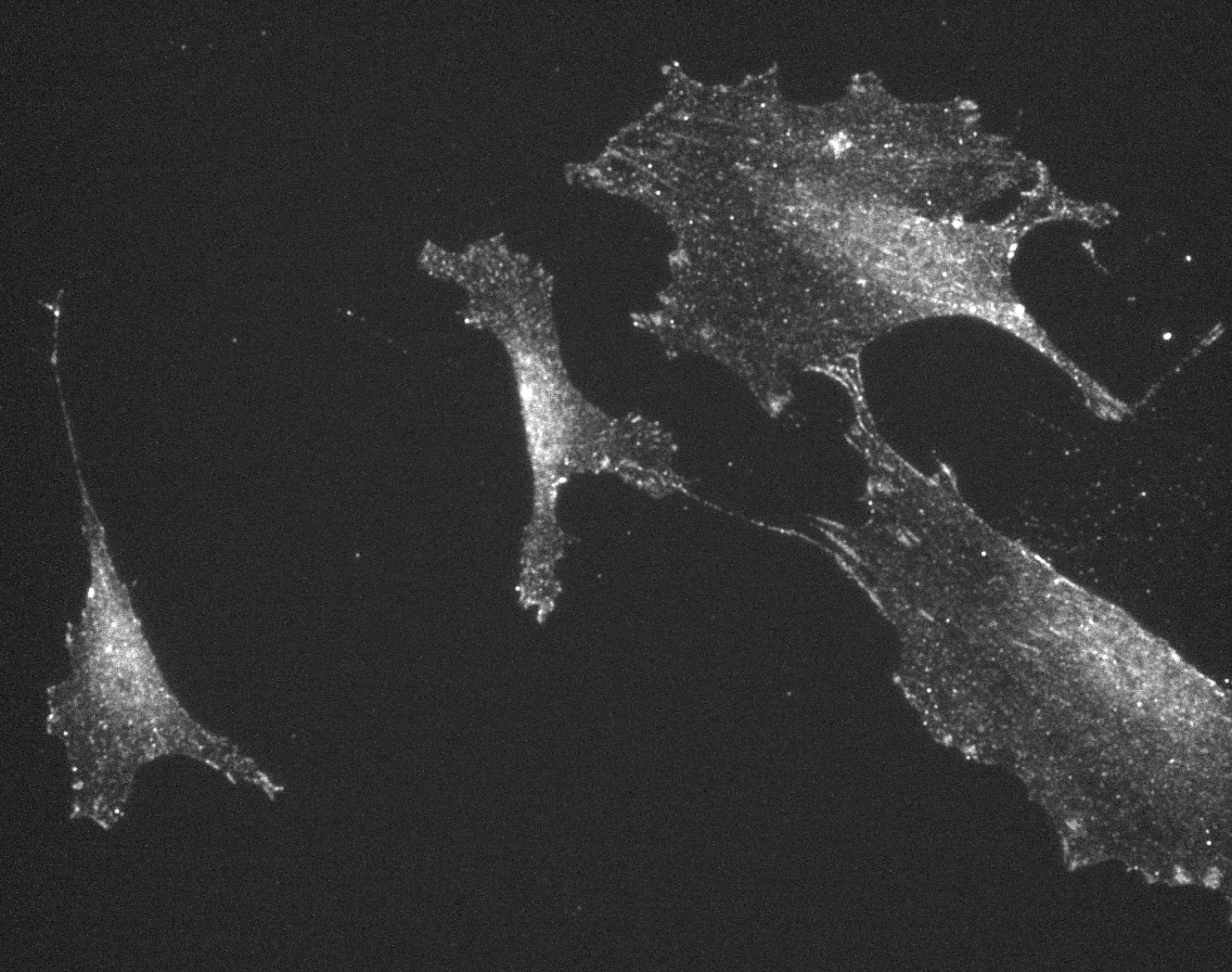

### H_Plate_R_p00_0_A01f41d2.TIF

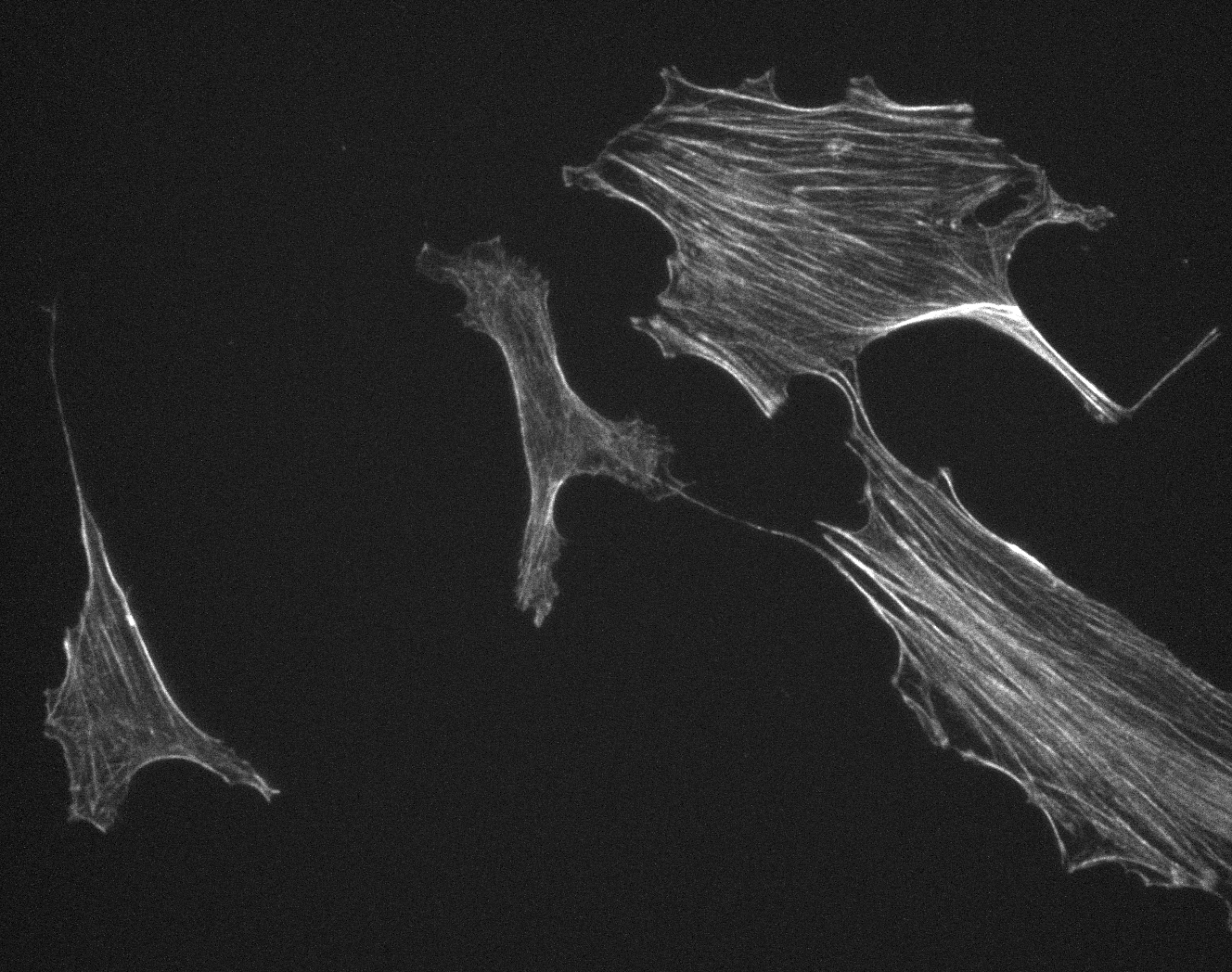

### H_Plate_R_p00_0_A01f41d3.TIF

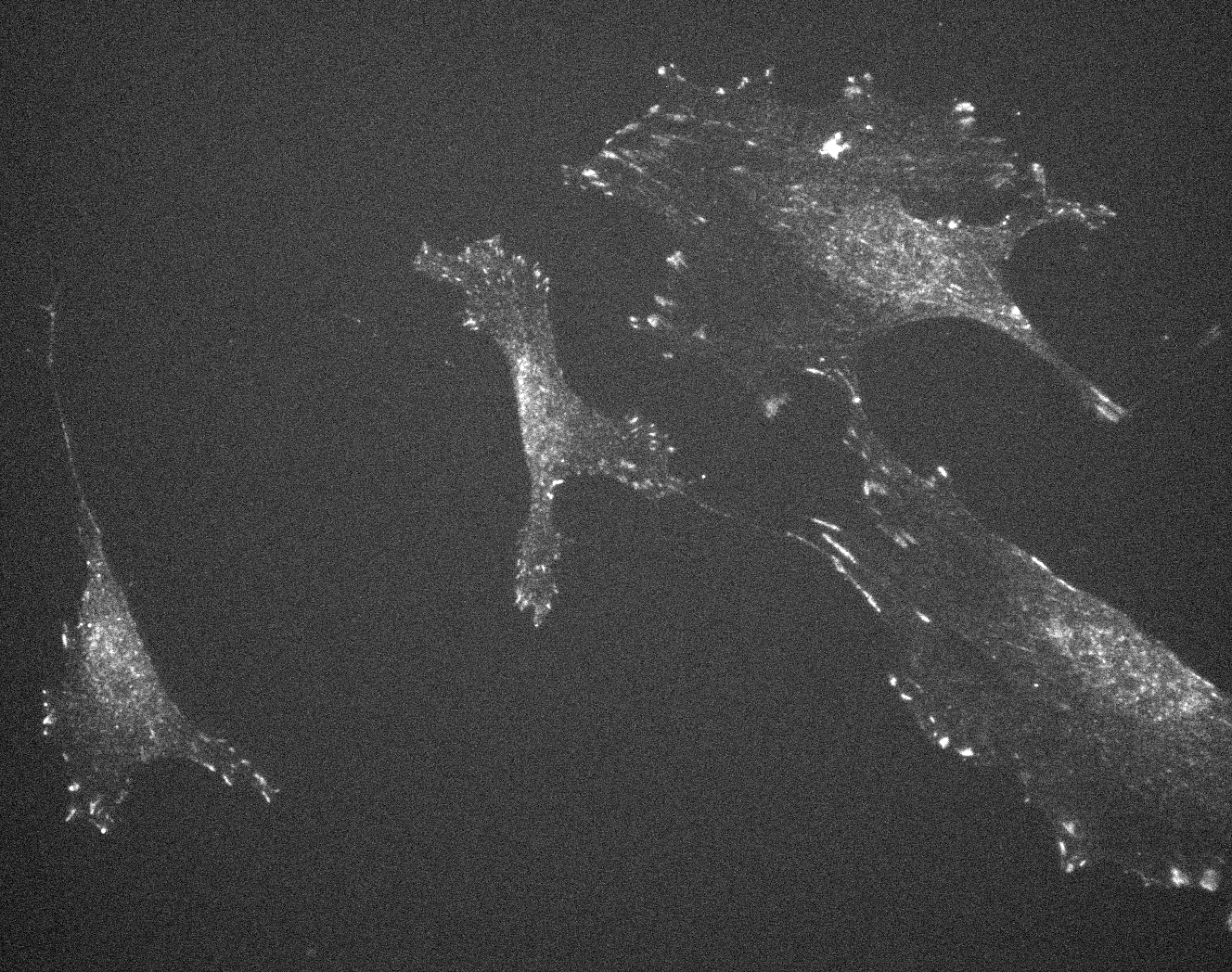

### I_groundtruth.jpg

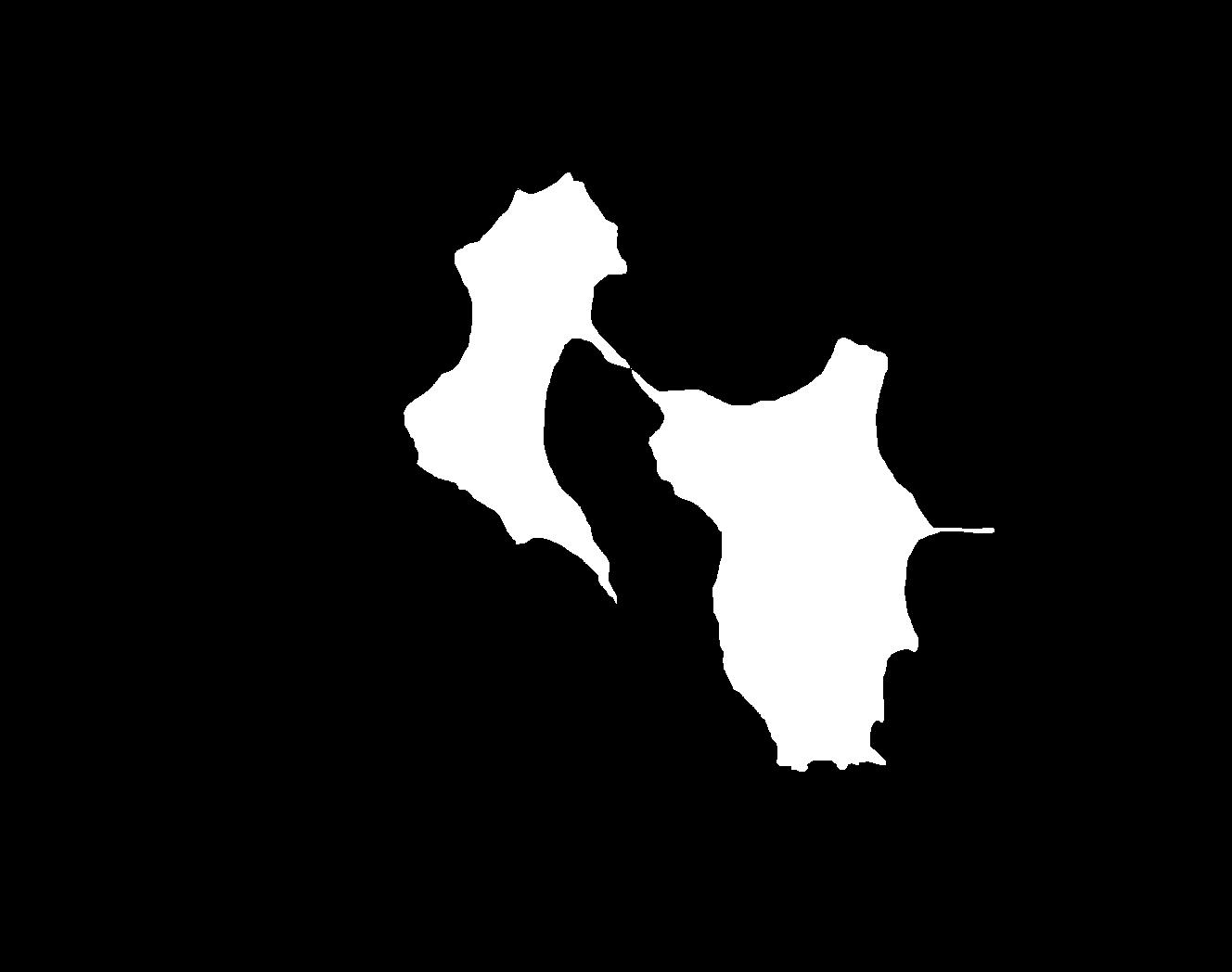

### I_Plate_R_p00_0_A01f39d0.TIF

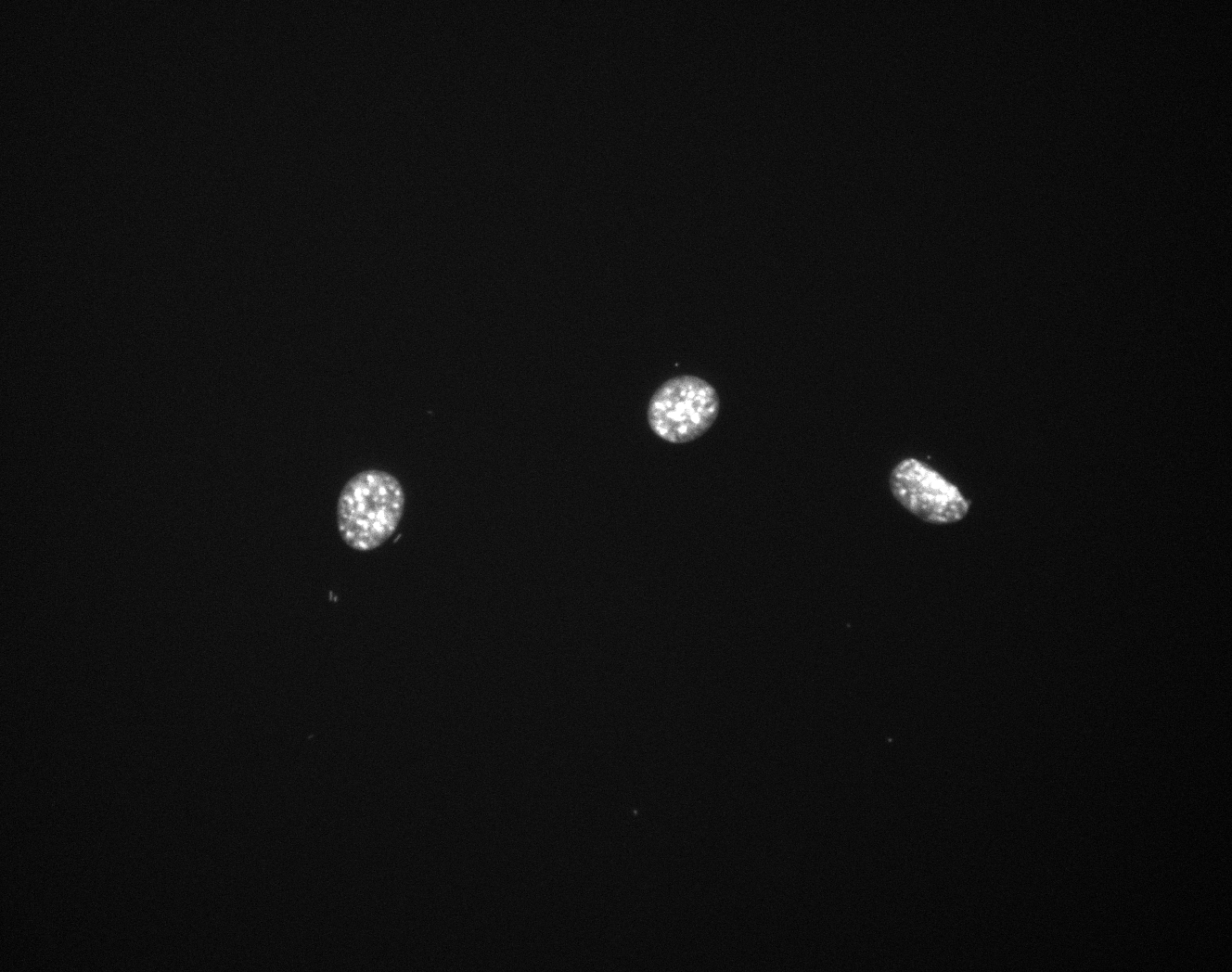

### I_Plate_R_p00_0_A01f39d1.TIF

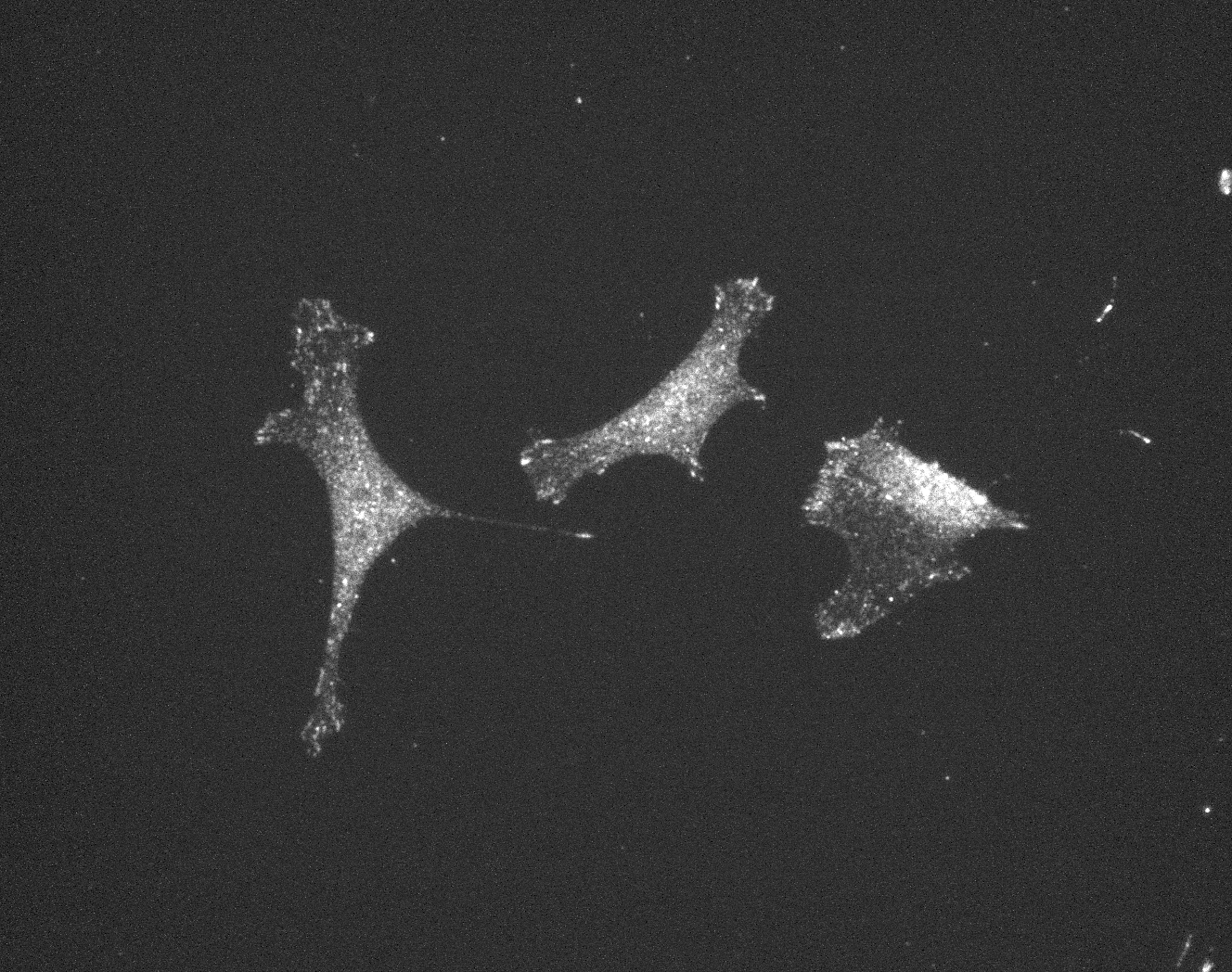

### I_Plate_R_p00_0_A01f39d2.TIF

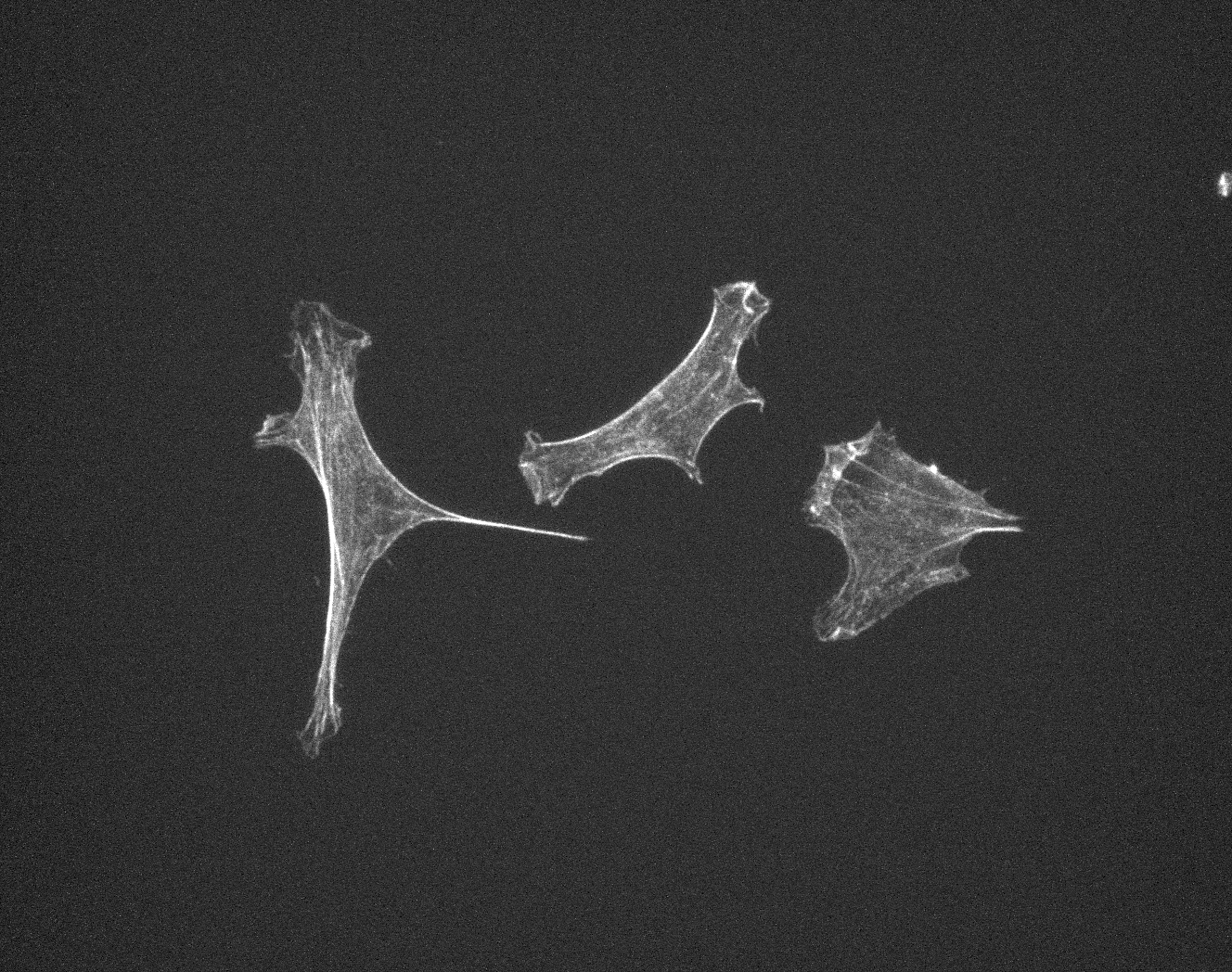

### I_Plate_R_p00_0_A01f39d3.TIF

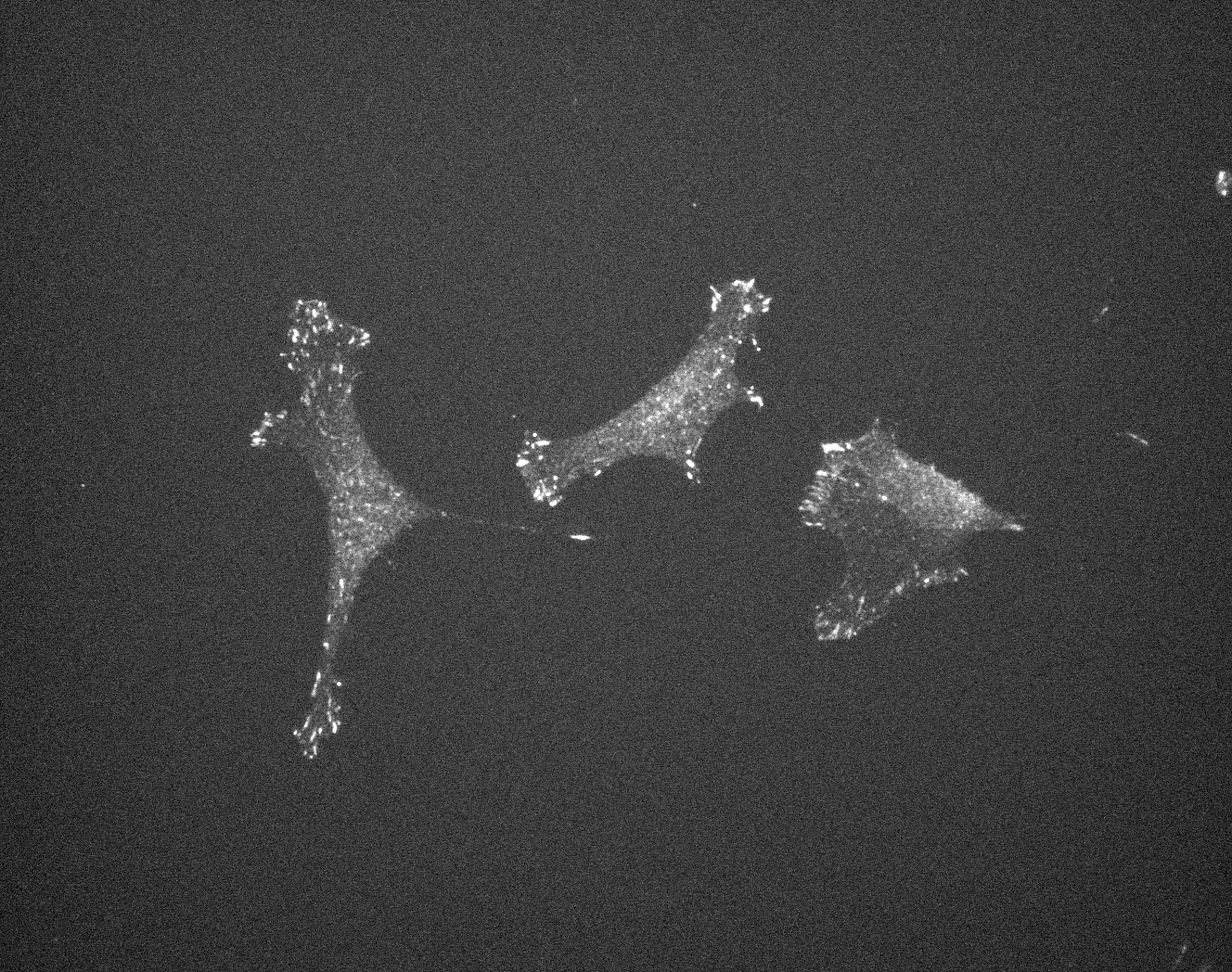

### J_groundtruth.jpg

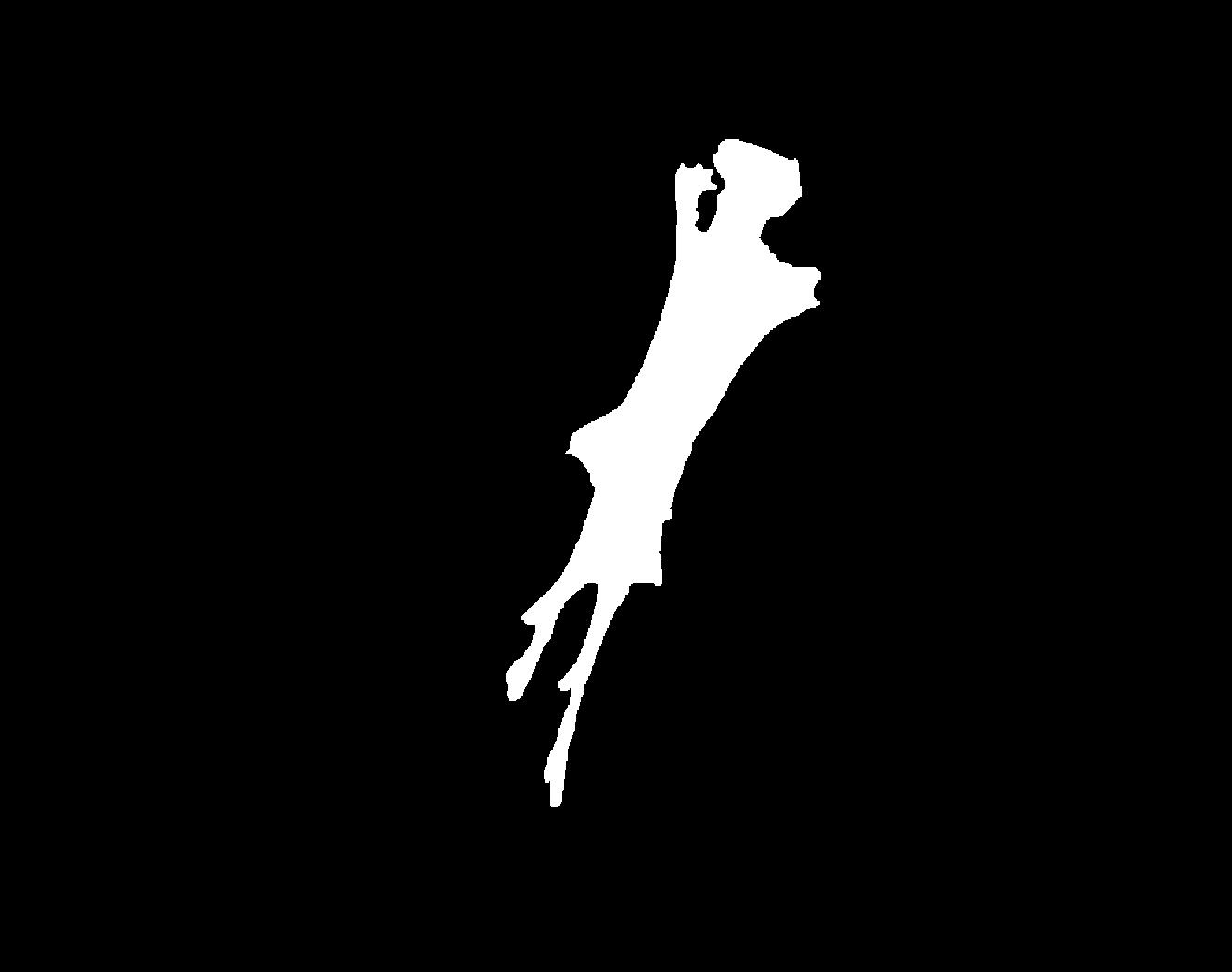

### J_Plate_R_p00_0_A01f49d0.TIF

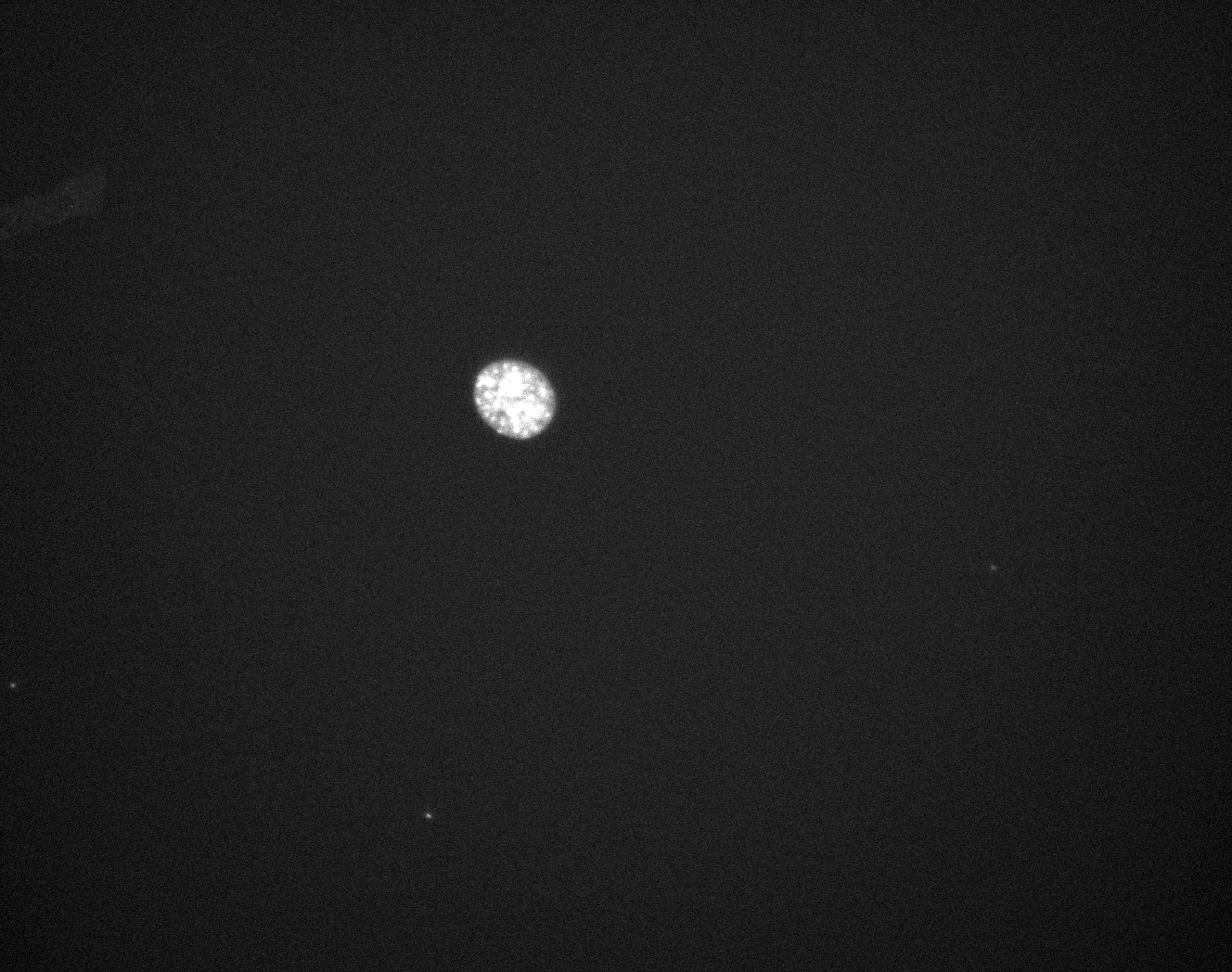

### J_Plate_R_p00_0_A01f49d1.TIF

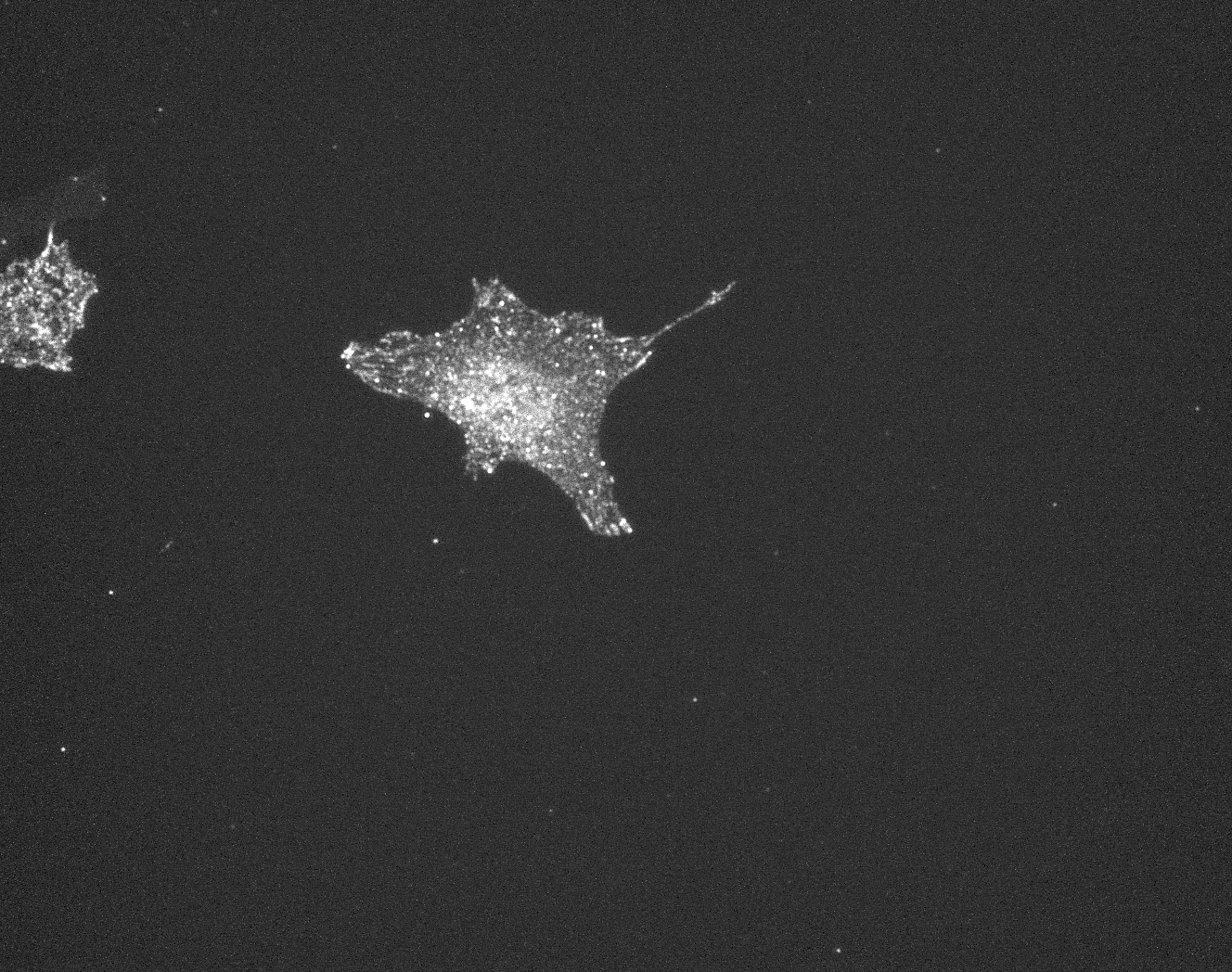

### J_Plate_R_p00_0_A01f49d2.TIF

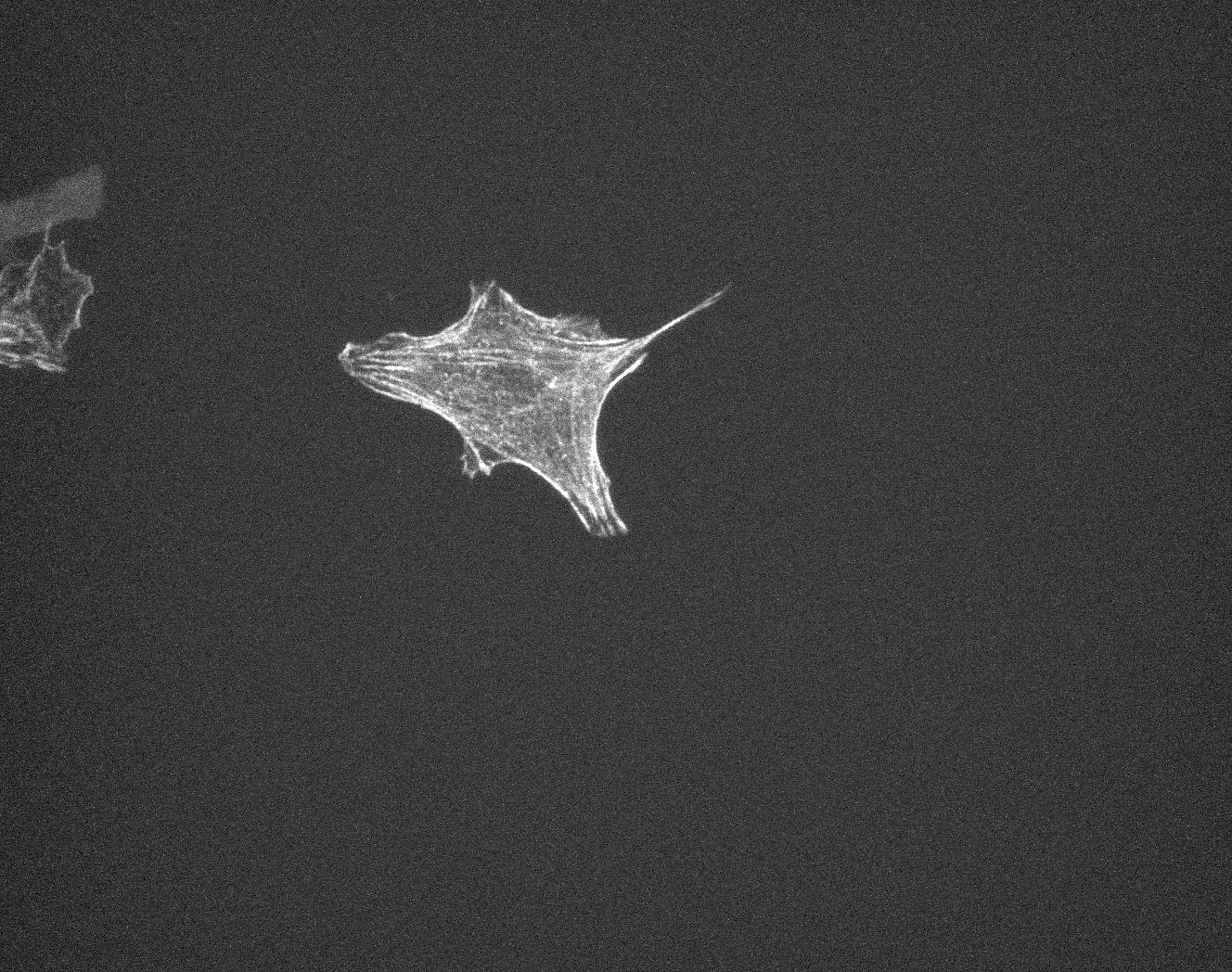

### J_Plate_R_p00_0_A01f49d3.TIF

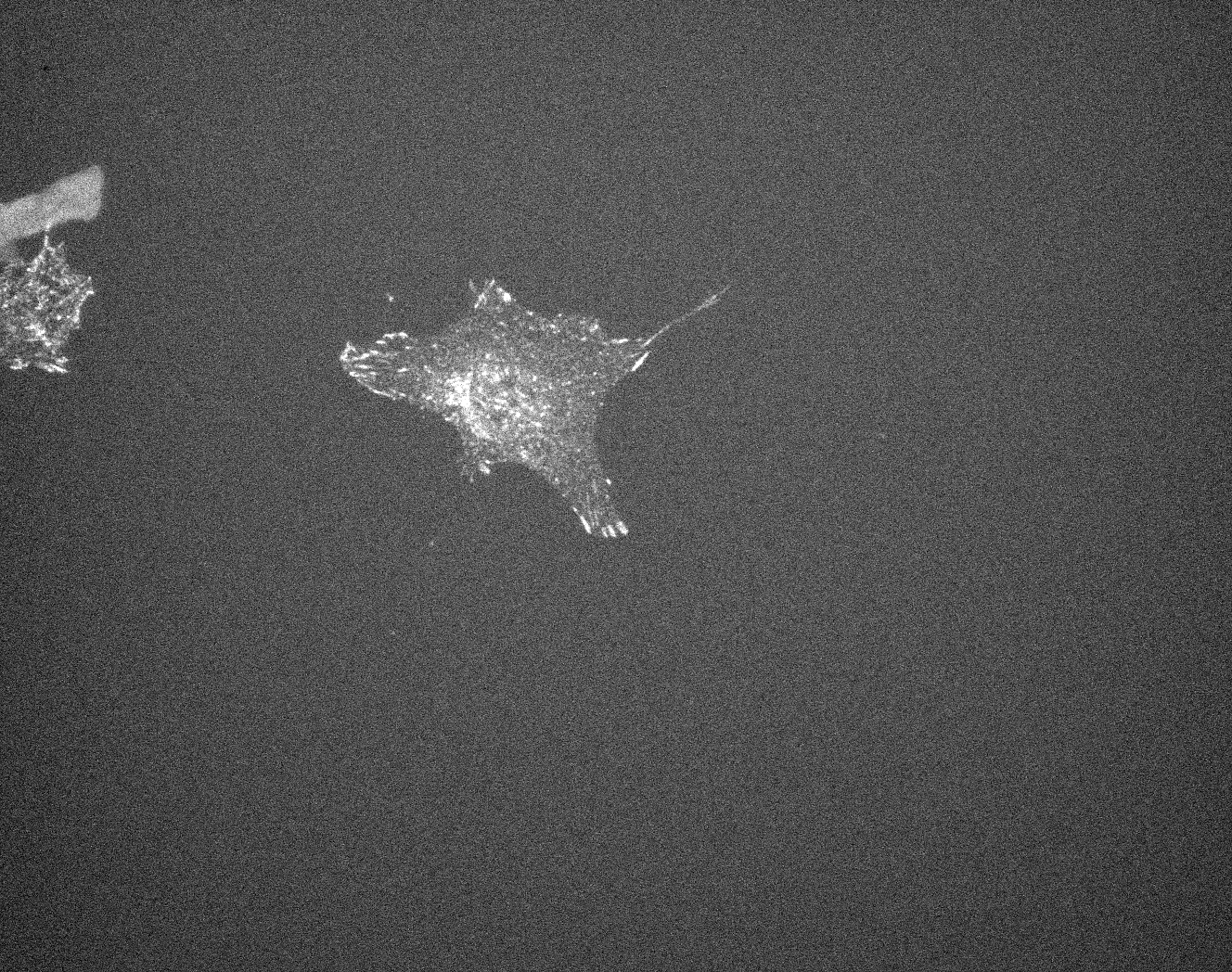
