## Supplementary figures and images for "Interactive machine learning for fast and robust cell profiling"

### A_Plate_R_p00_0_A01f01d0.TIF

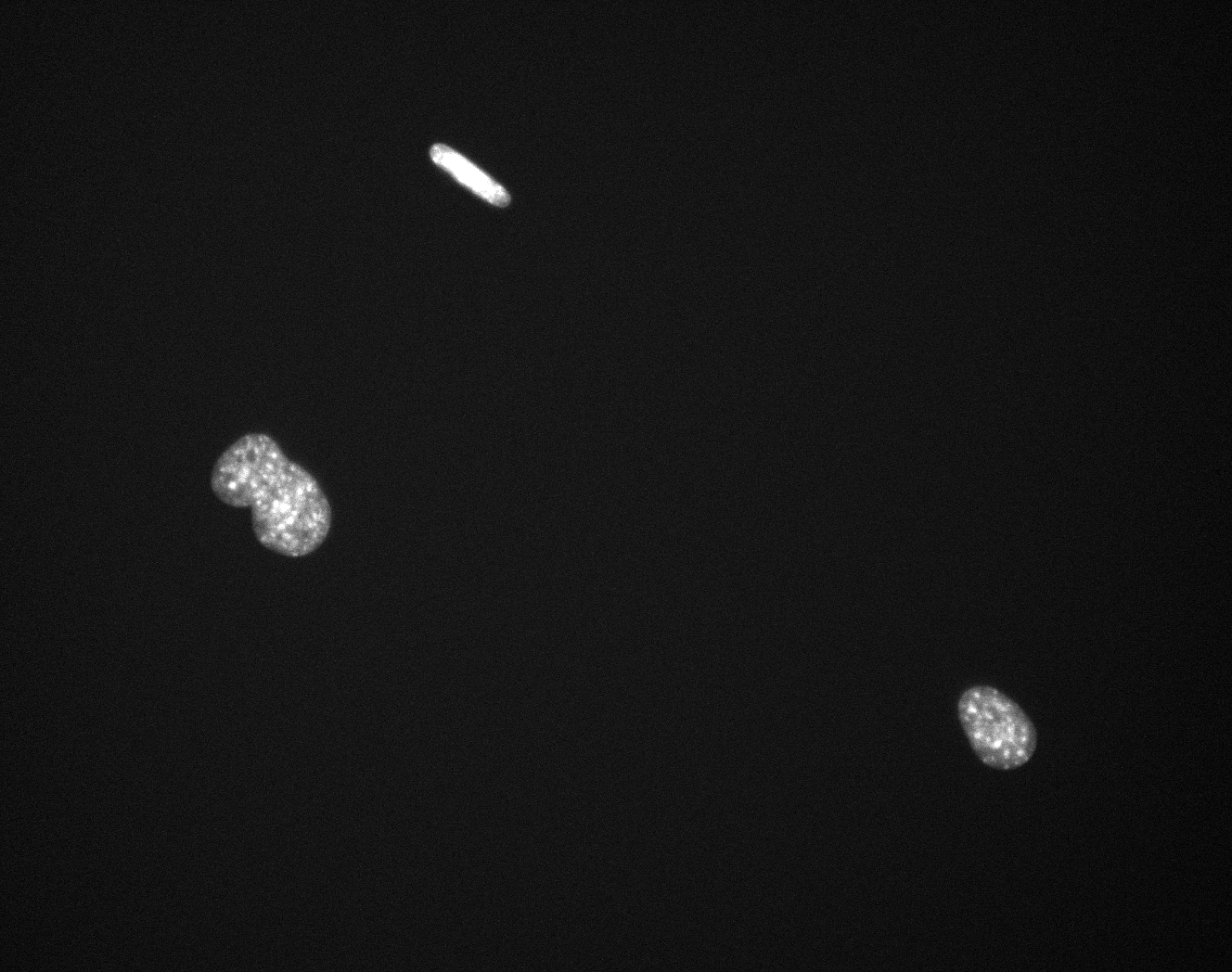

### A_Plate_R_p00_0_A01f01d1.TIF

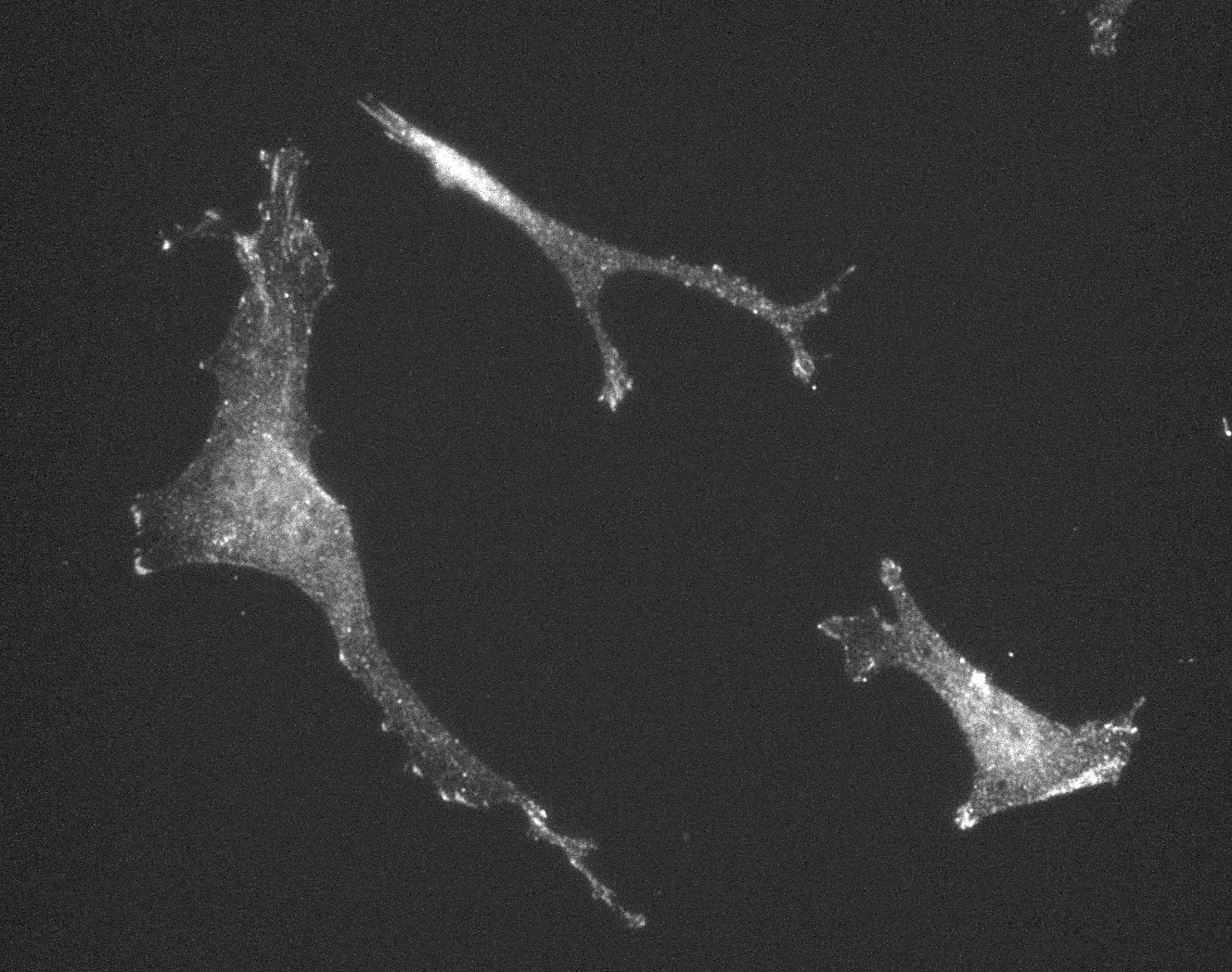

### A_Plate_R_p00_0_A01f01d2.TIF

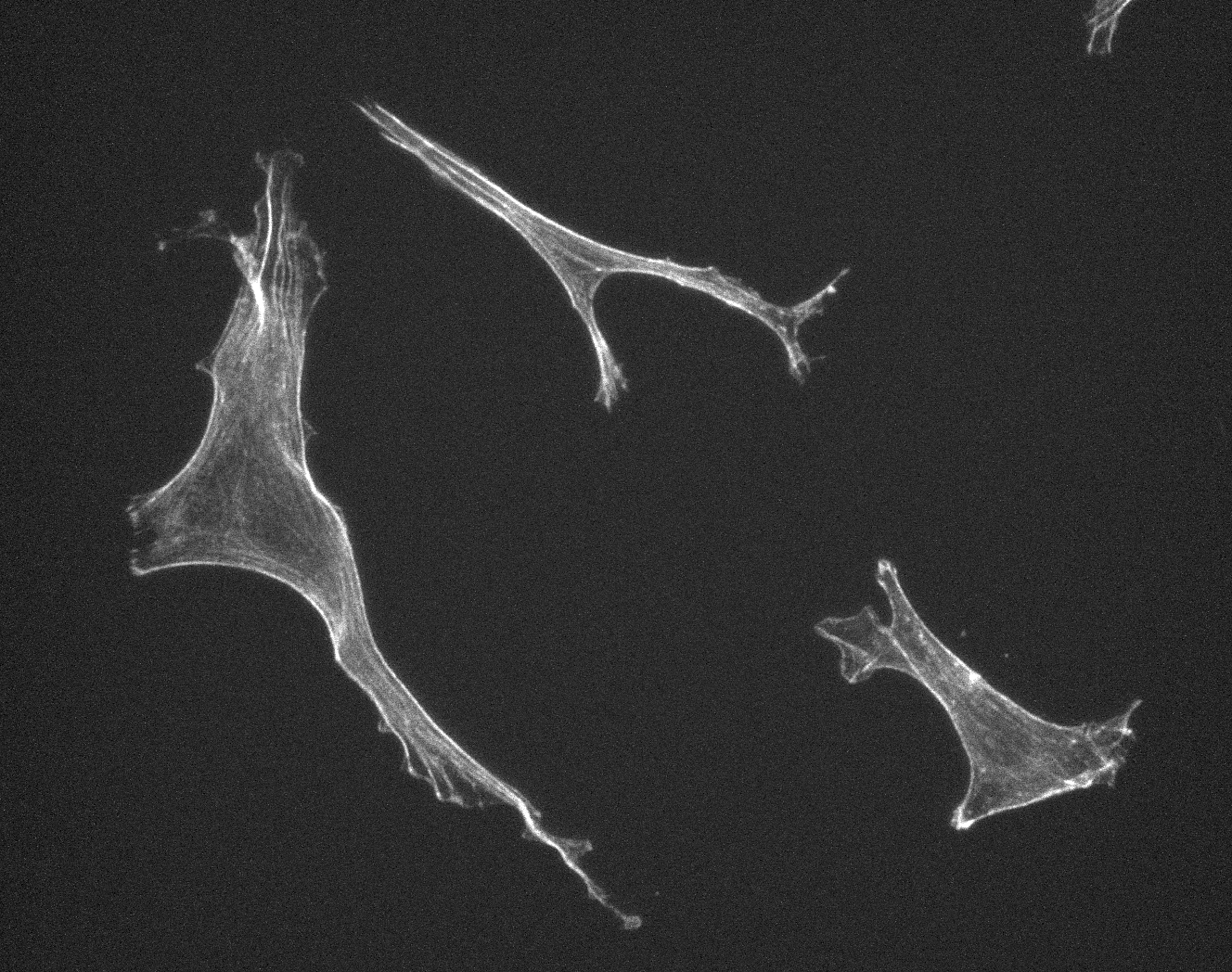

### A_Plate_R_p00_0_A01f01d2_CytoplasmOutline.tiff

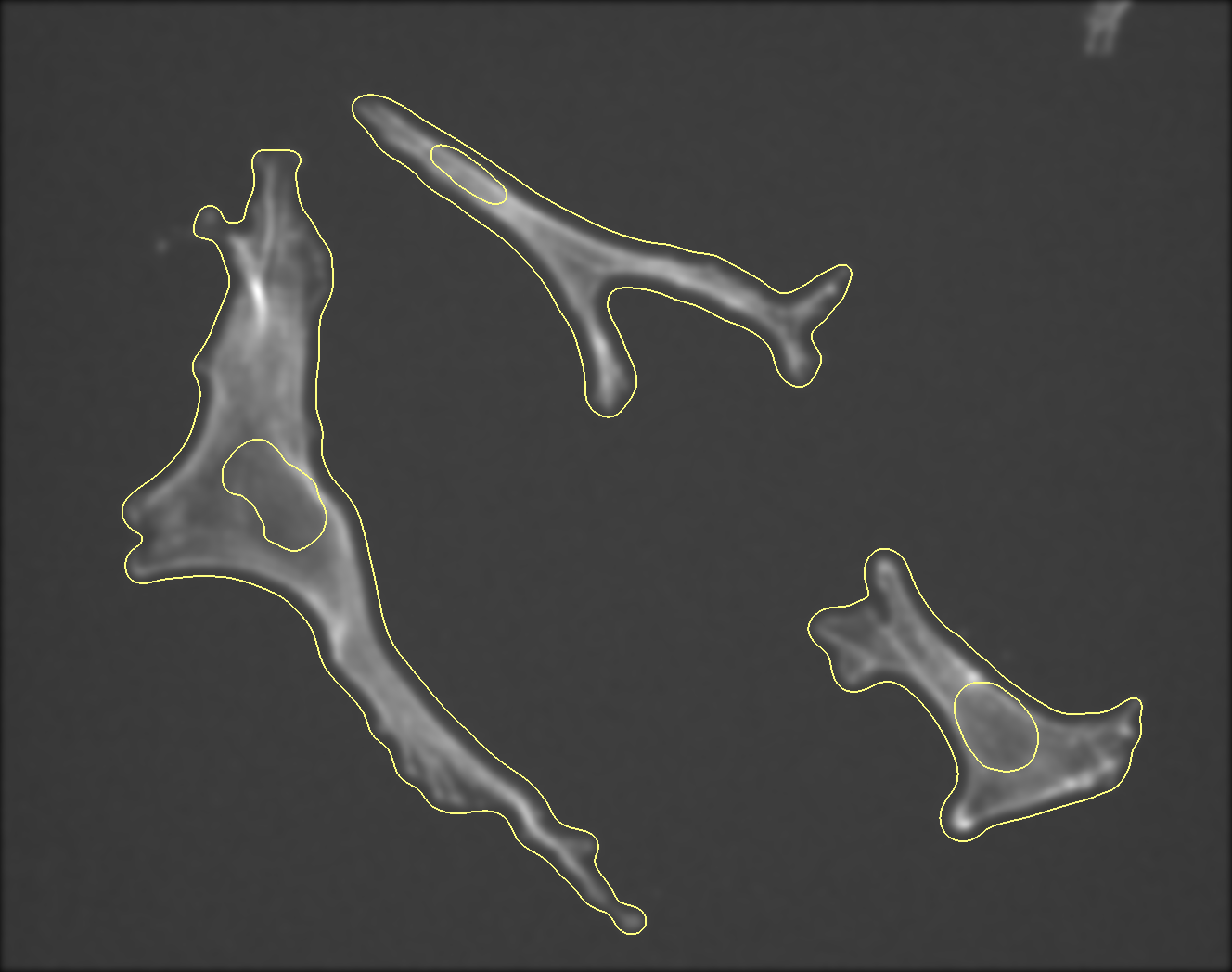

### A_Plate_R_p00_0_A01f01d3.TIF

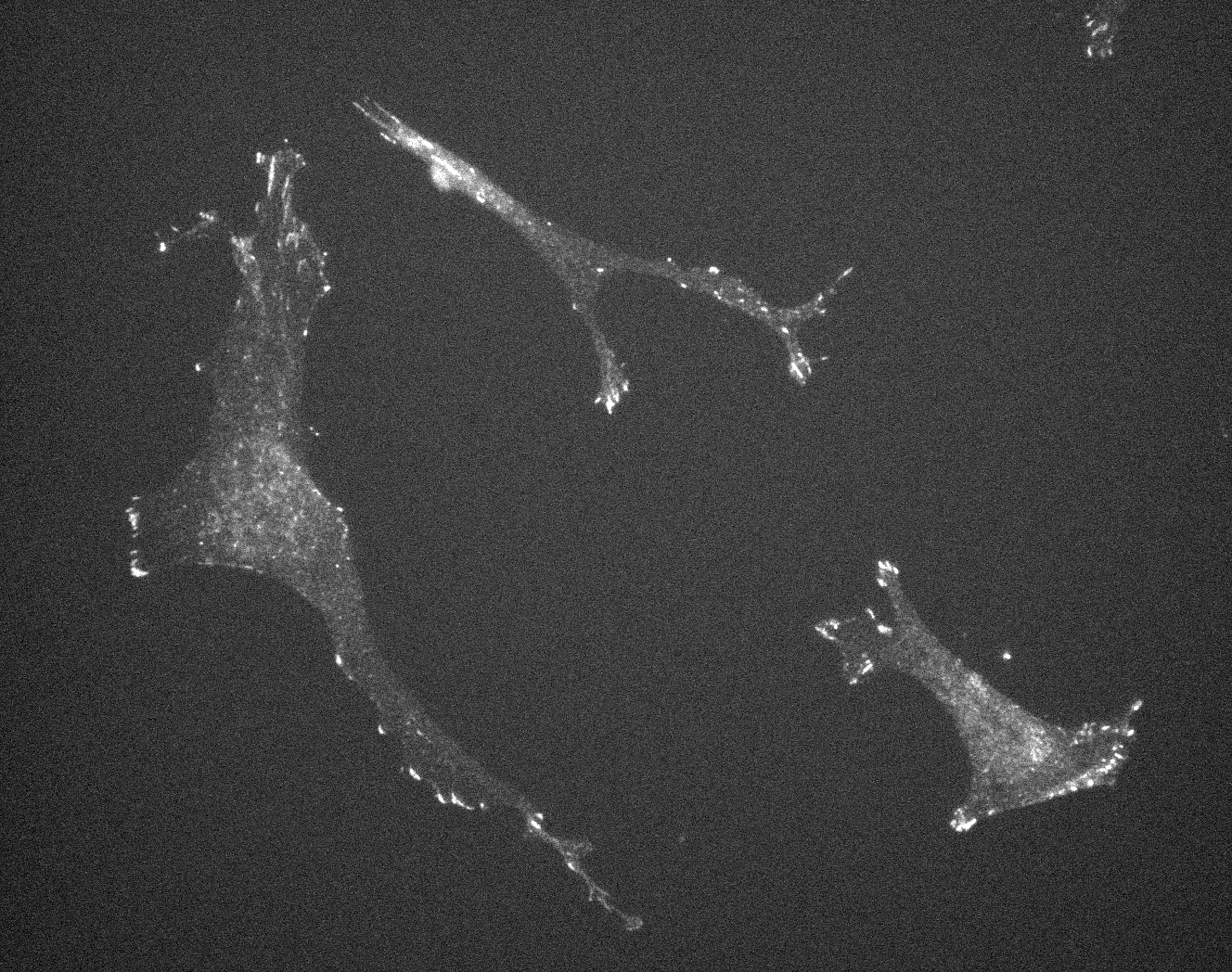

### A_Plate_R_p00_0_A01f11d0.TIF

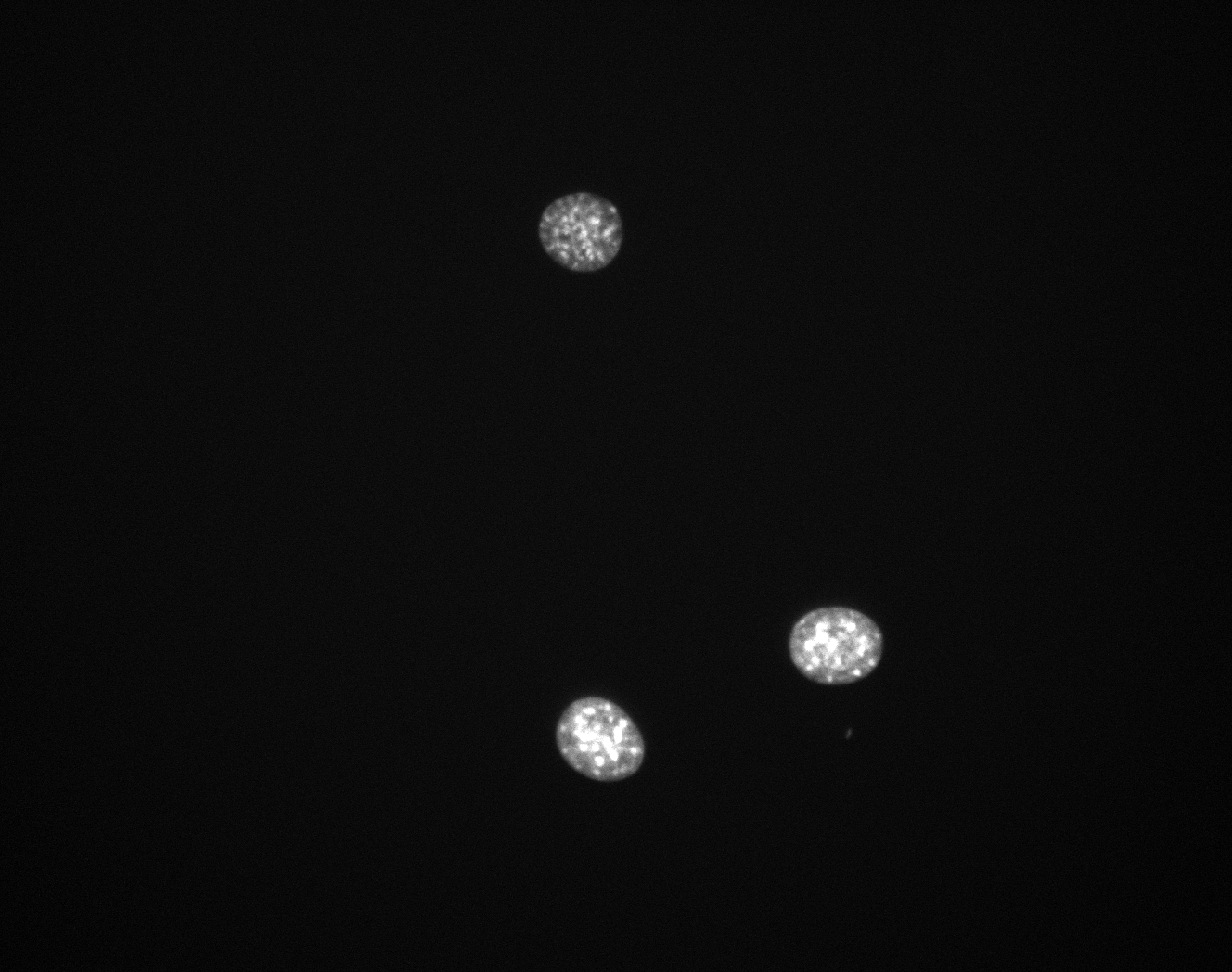

### A_Plate_R_p00_0_A01f11d1.TIF

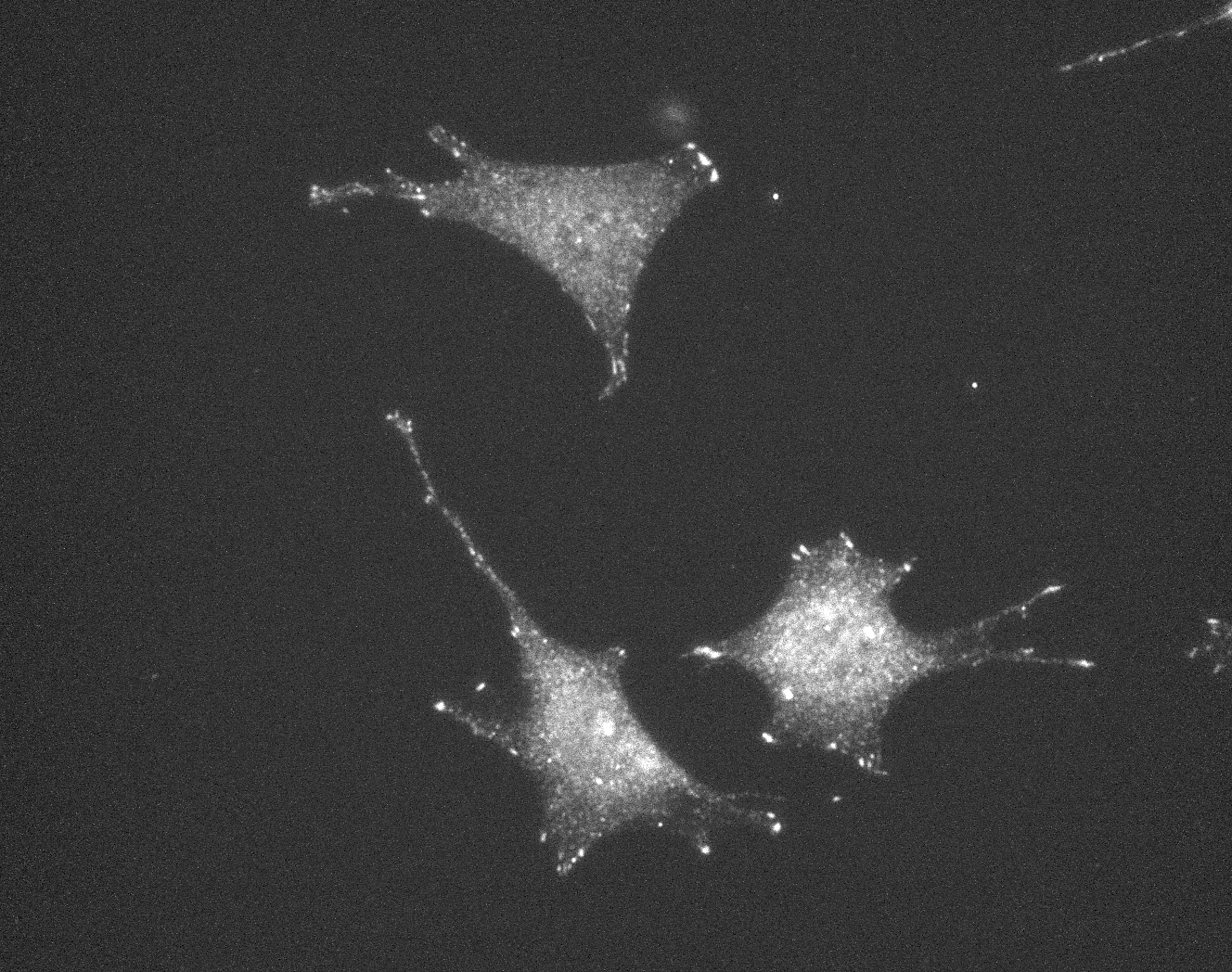

### A_Plate_R_p00_0_A01f11d2.TIF

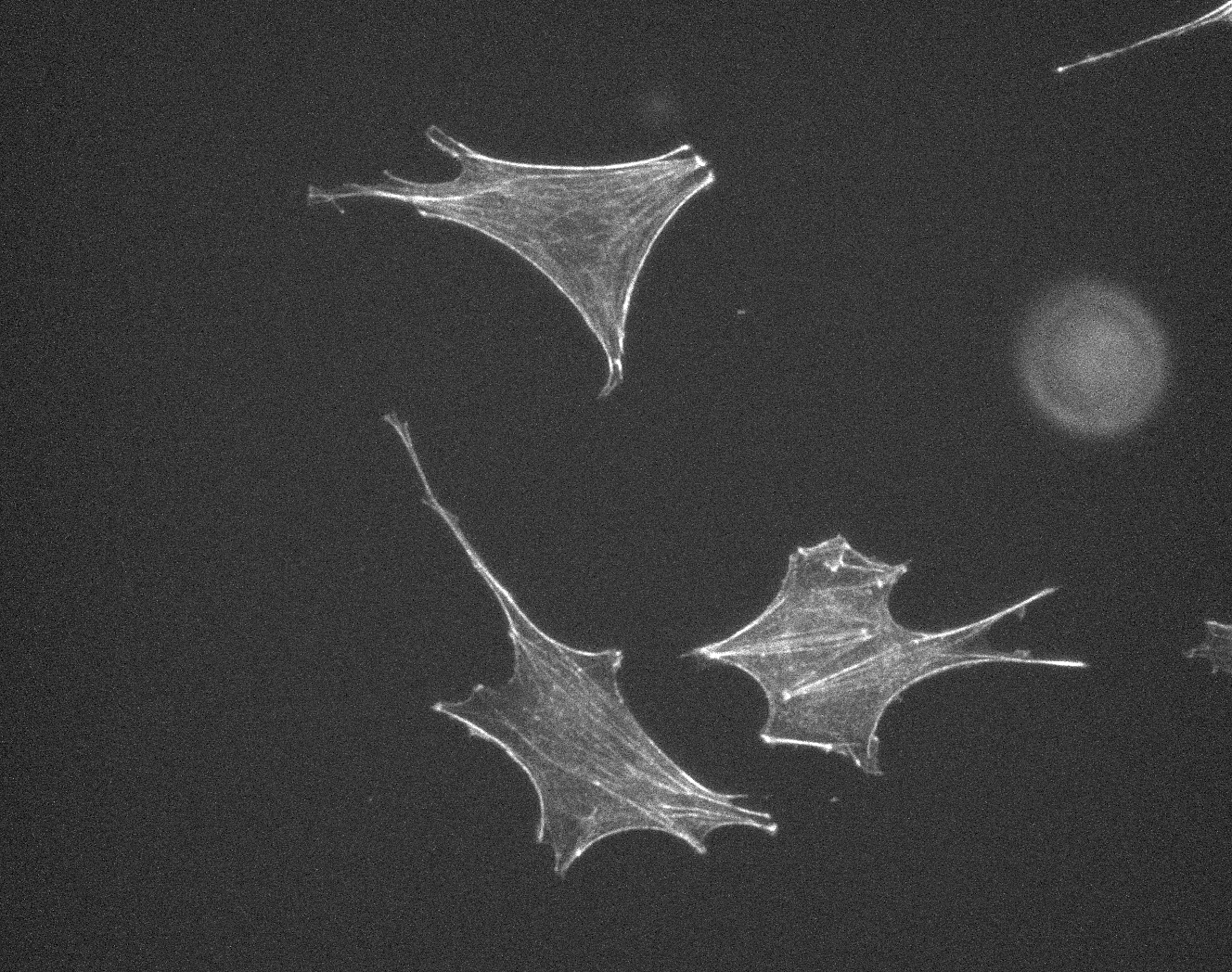

### A_Plate_R_p00_0_A01f11d3.TIF

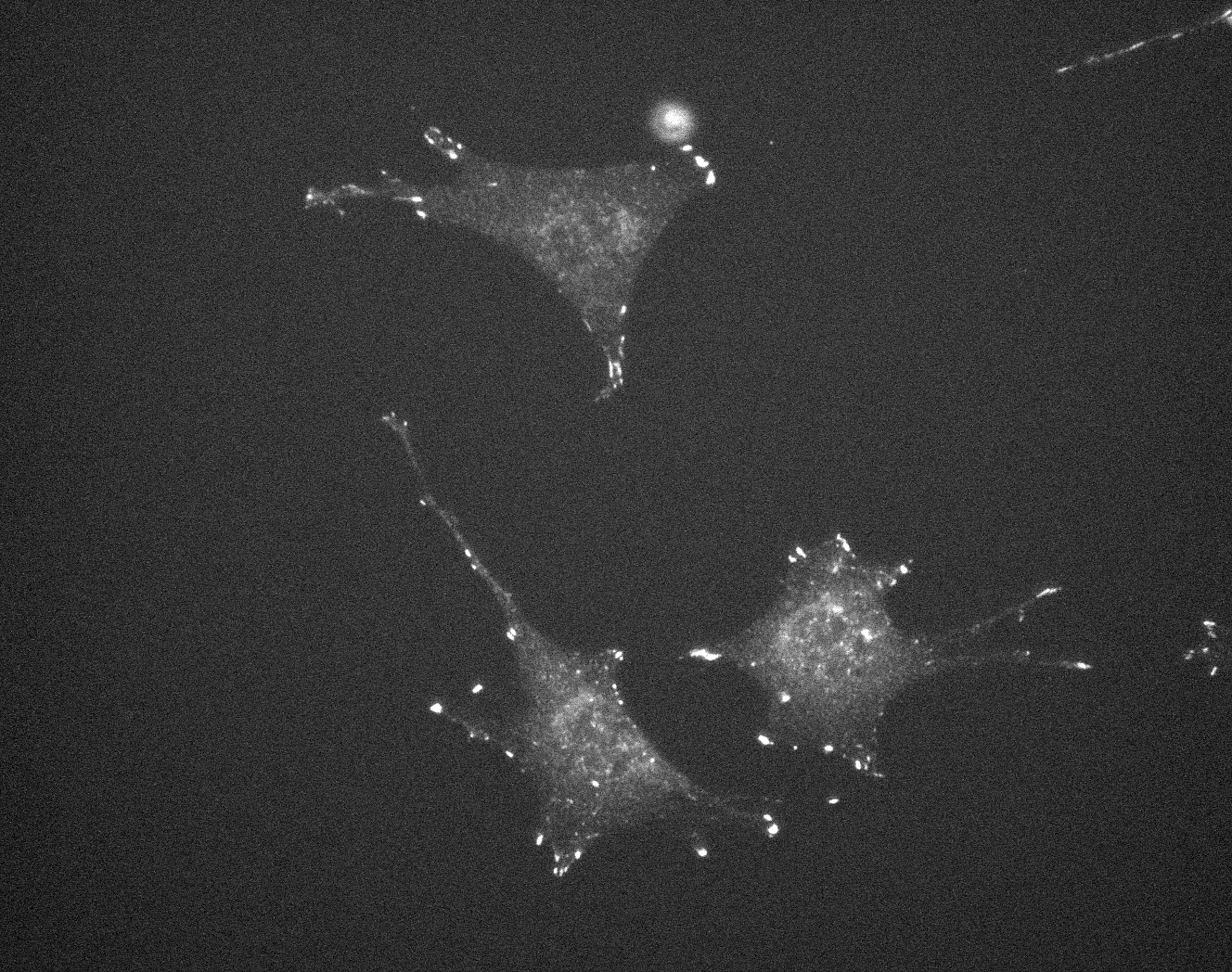

### A_Plate_R_p00_0_A01f11d3_FAOutline.tiff

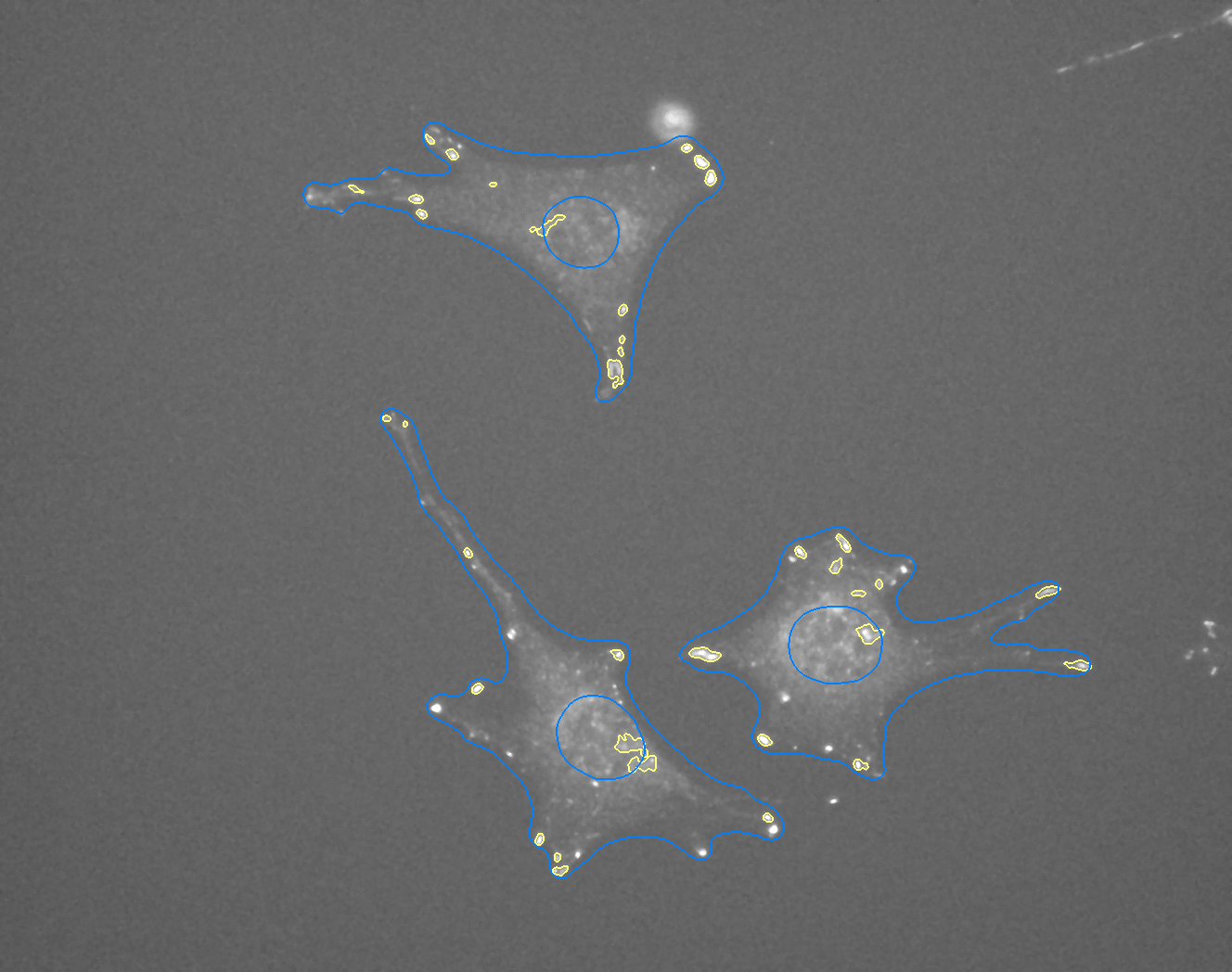

### A_Plate_R_p00_0_A01f27d0.TIF

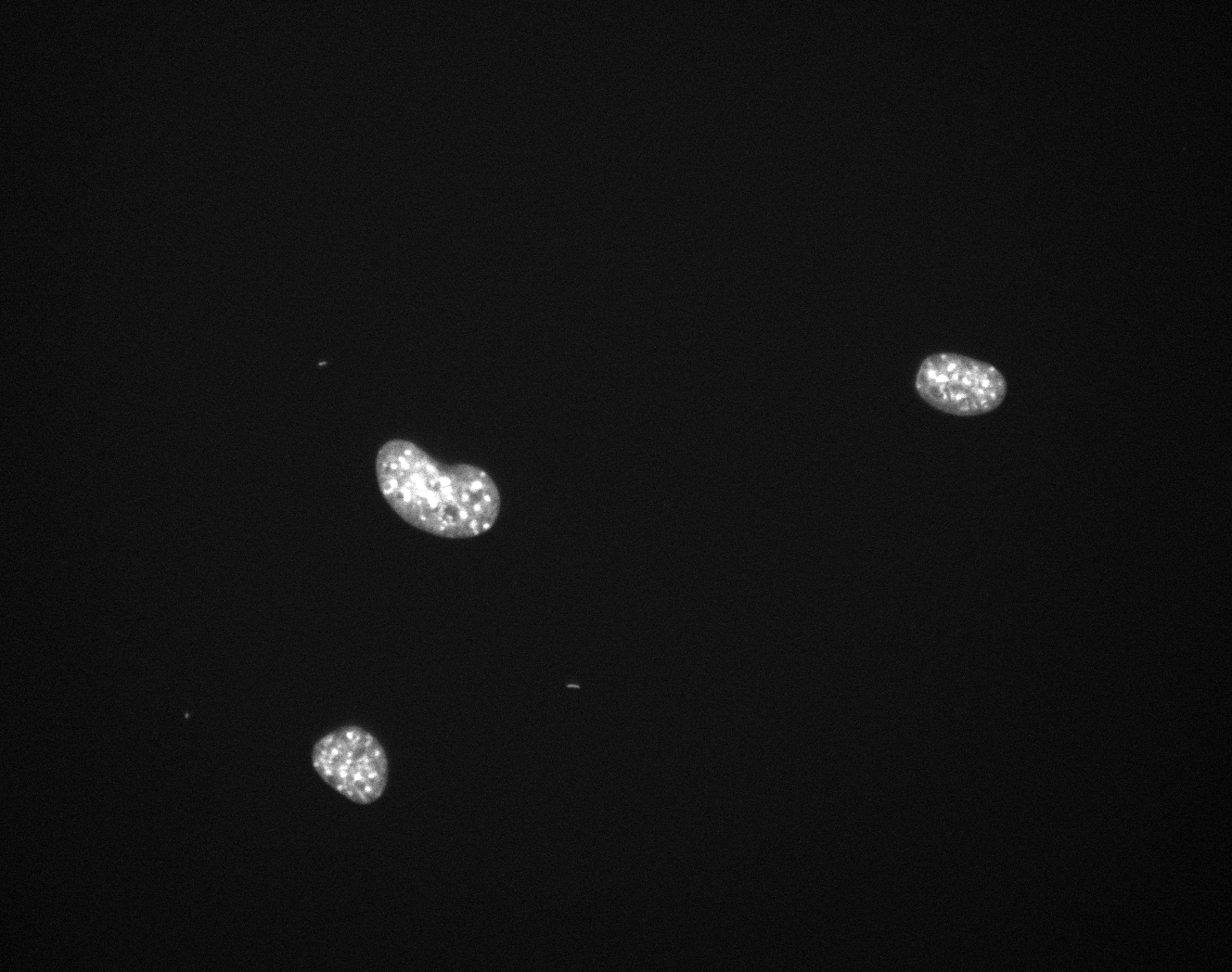

### A_Plate_R_p00_0_A01f27d1.TIF

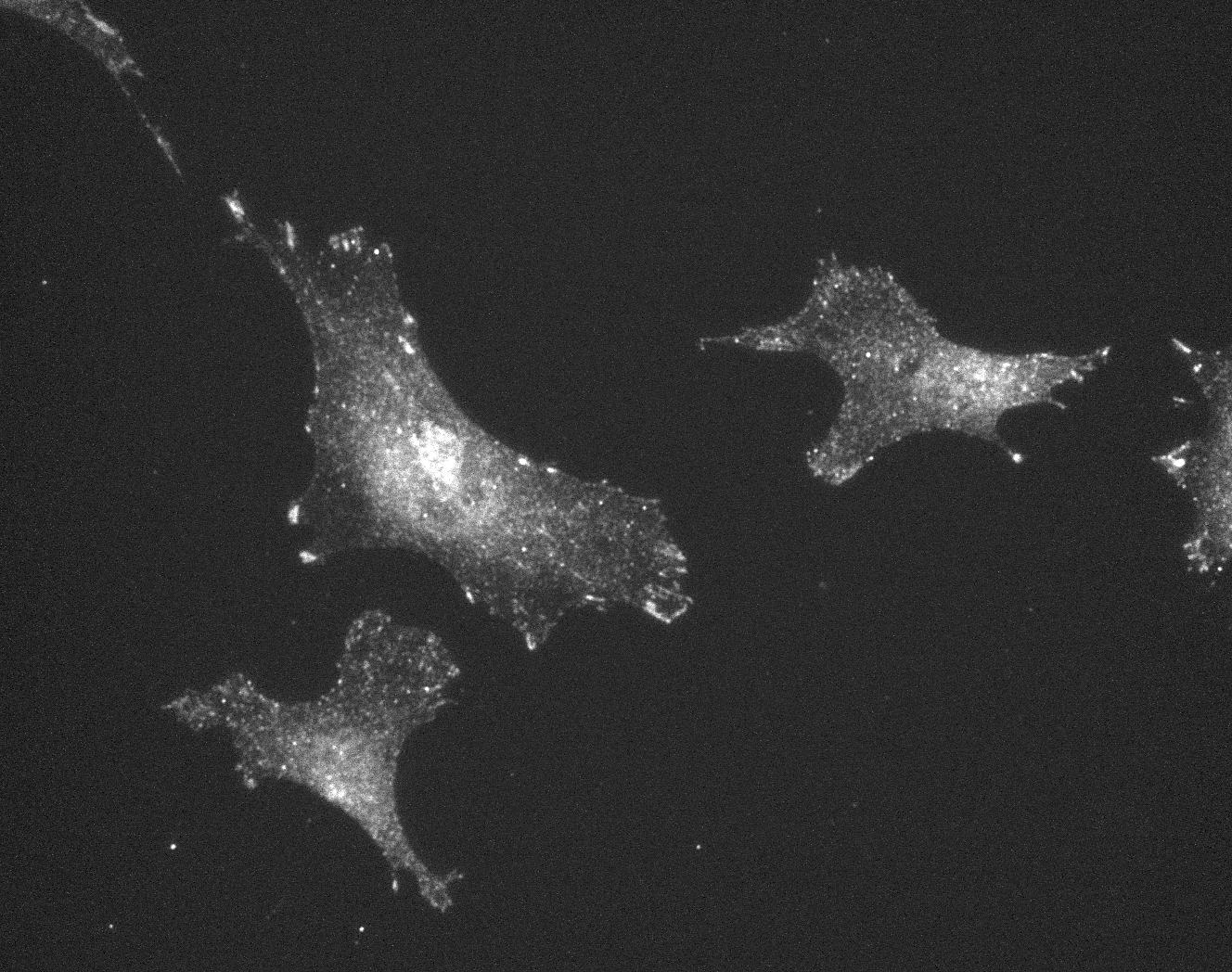

### A_Plate_R_p00_0_A01f27d2.TIF

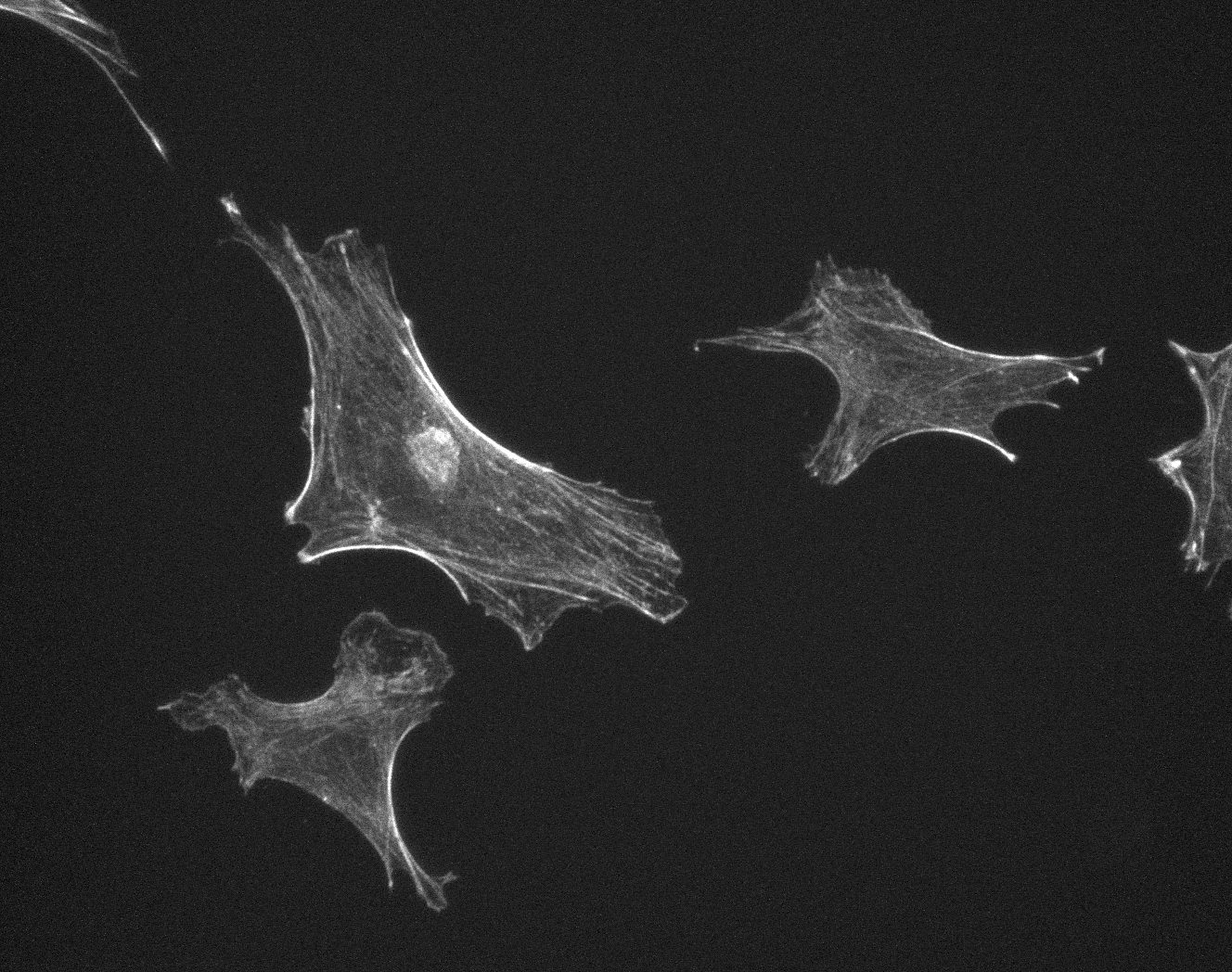

### A_Plate_R_p00_0_A01f27d3.TIF

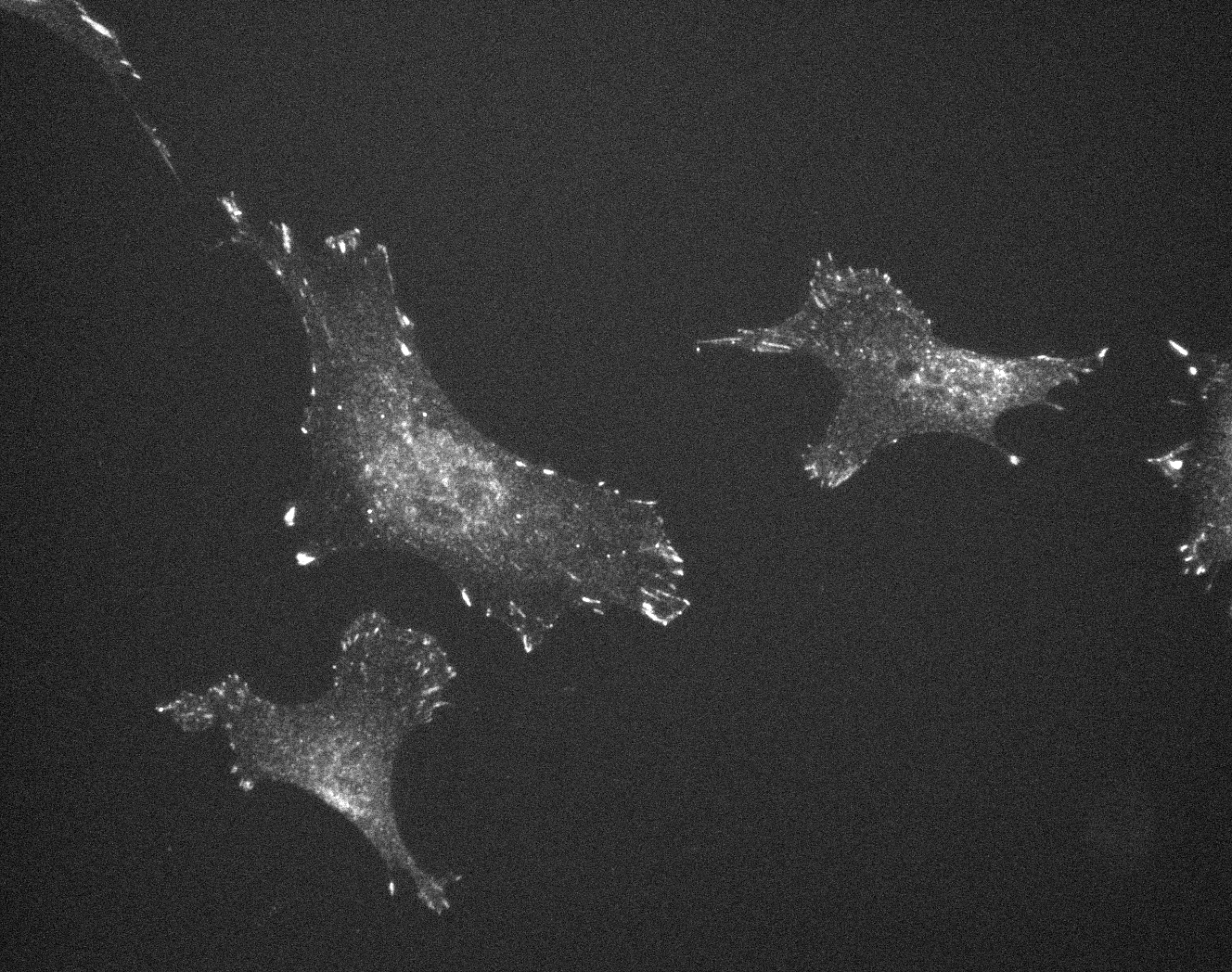

### B_Plate_R_p00_0_A01f04d0.TIF

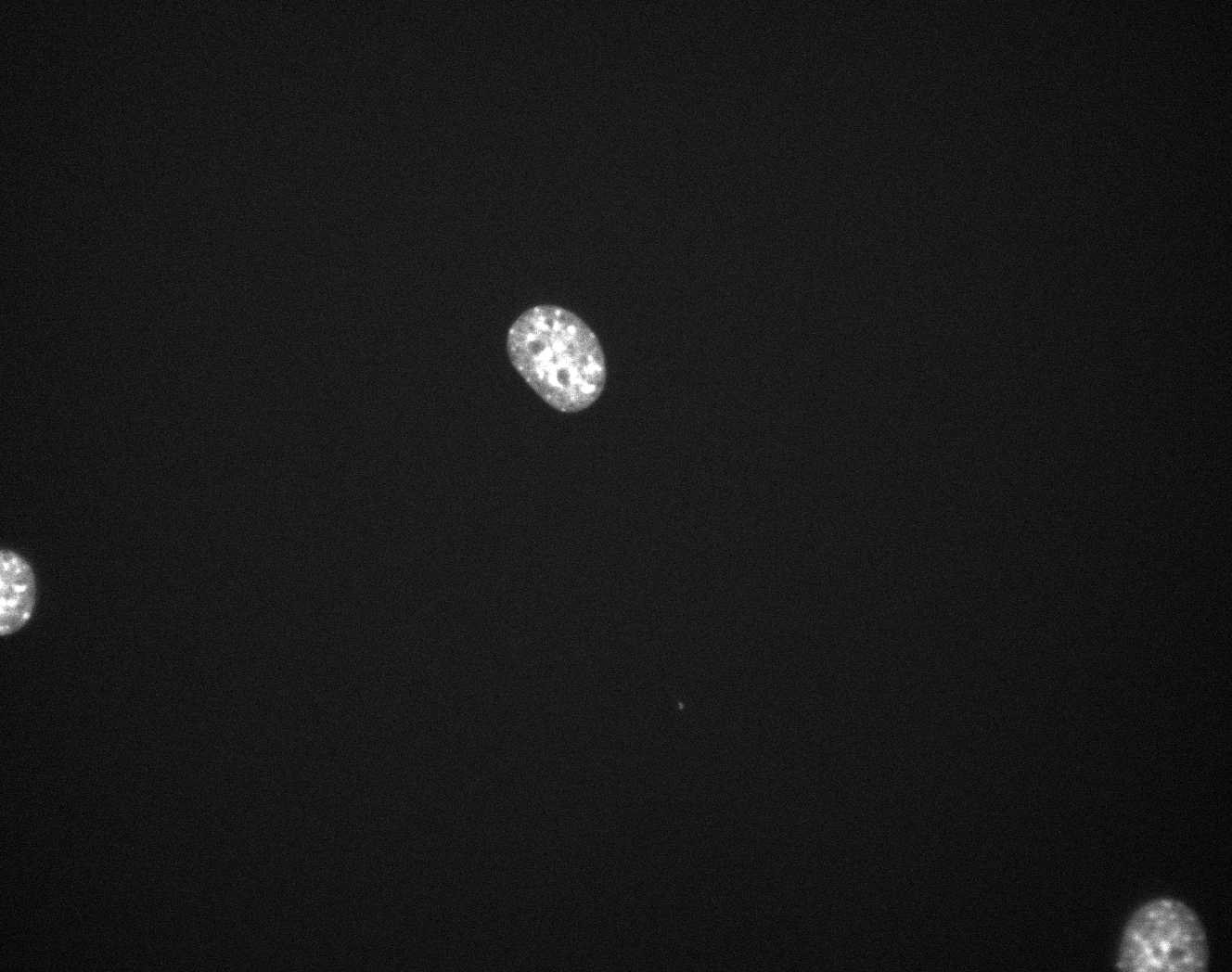

### B_Plate_R_p00_0_A01f04d1.TIF

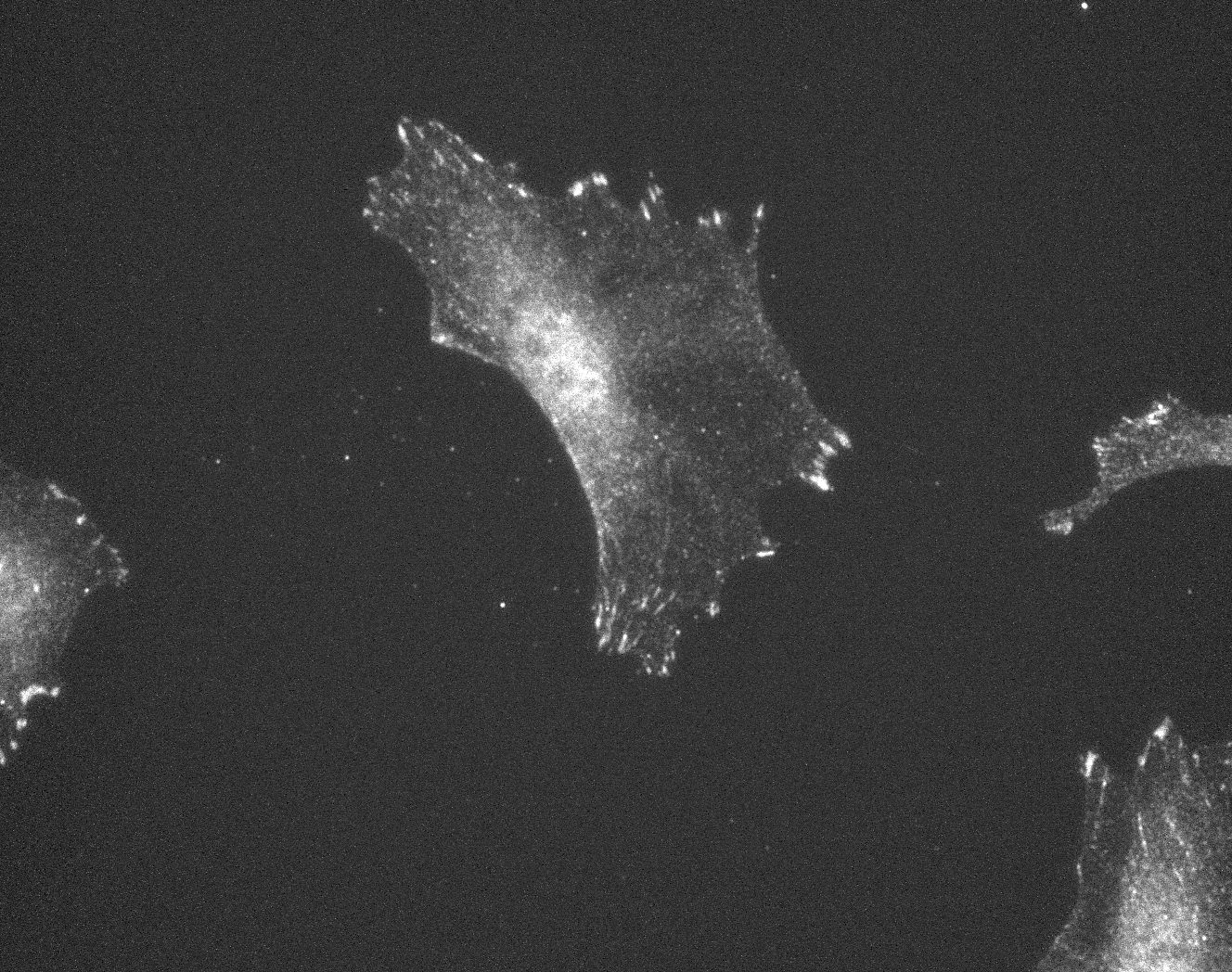

### B_Plate_R_p00_0_A01f04d2.TIF

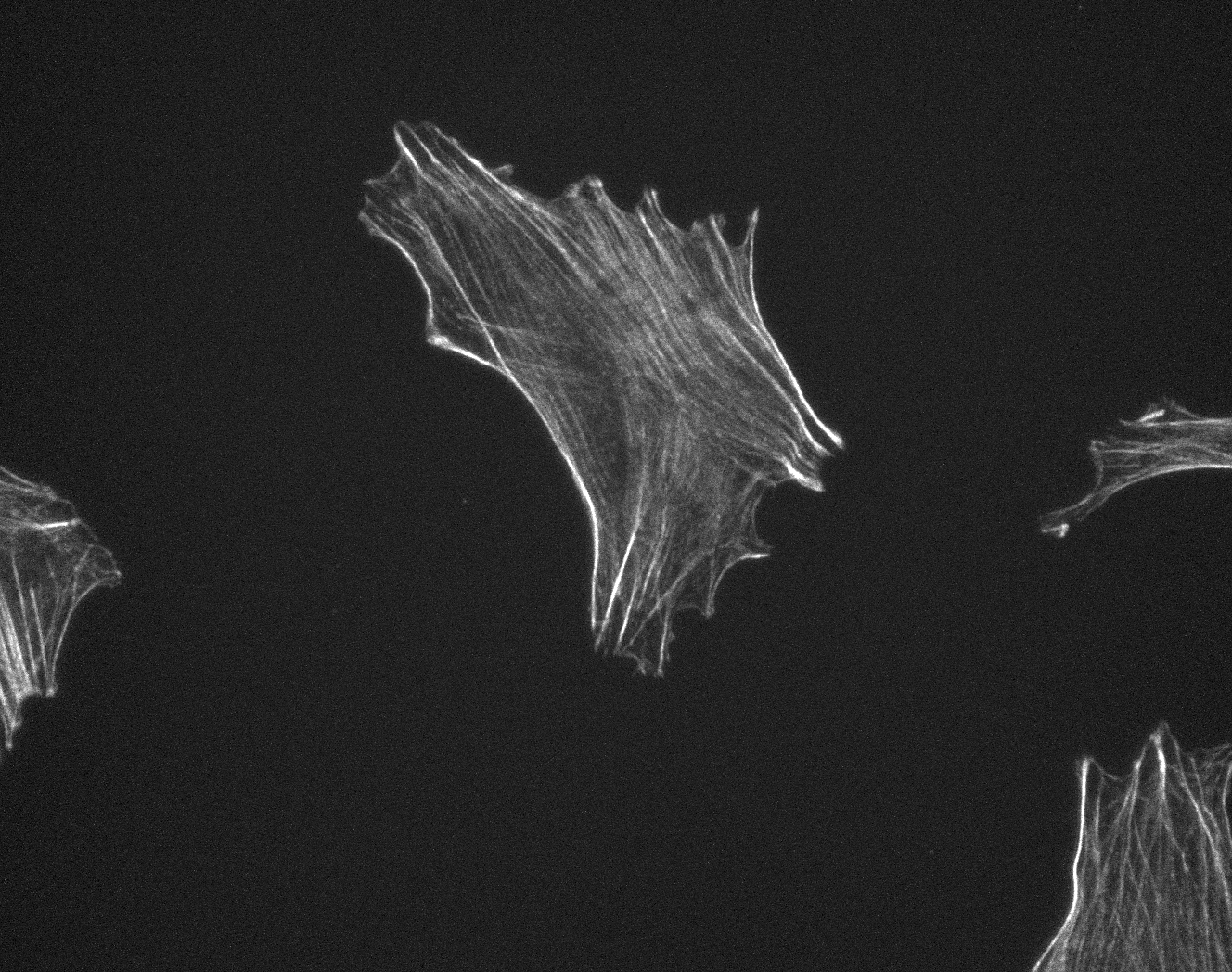

### B_Plate_R_p00_0_A01f04d2_CytoplasmOutline.tiff

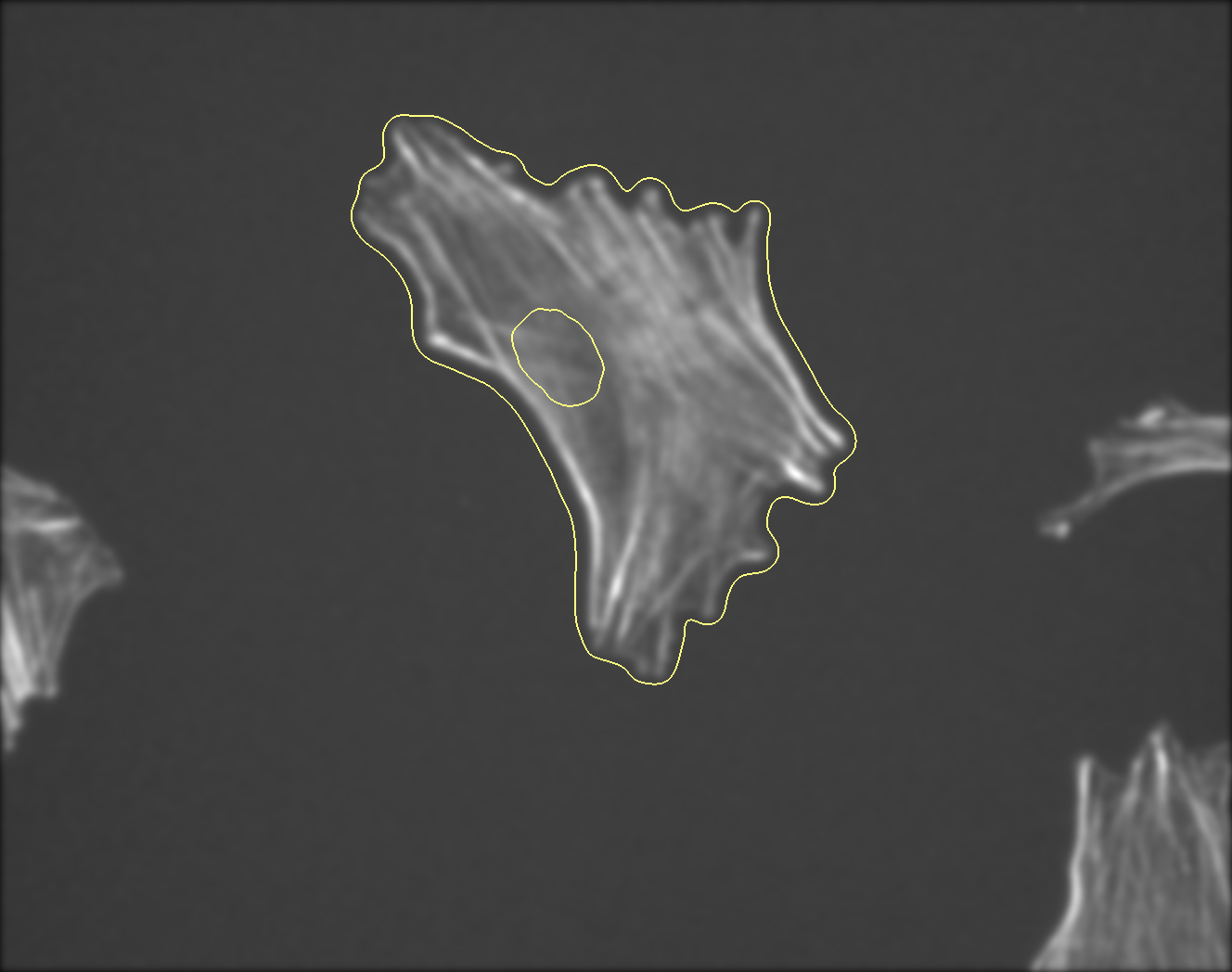

### B_Plate_R_p00_0_A01f04d3.TIF

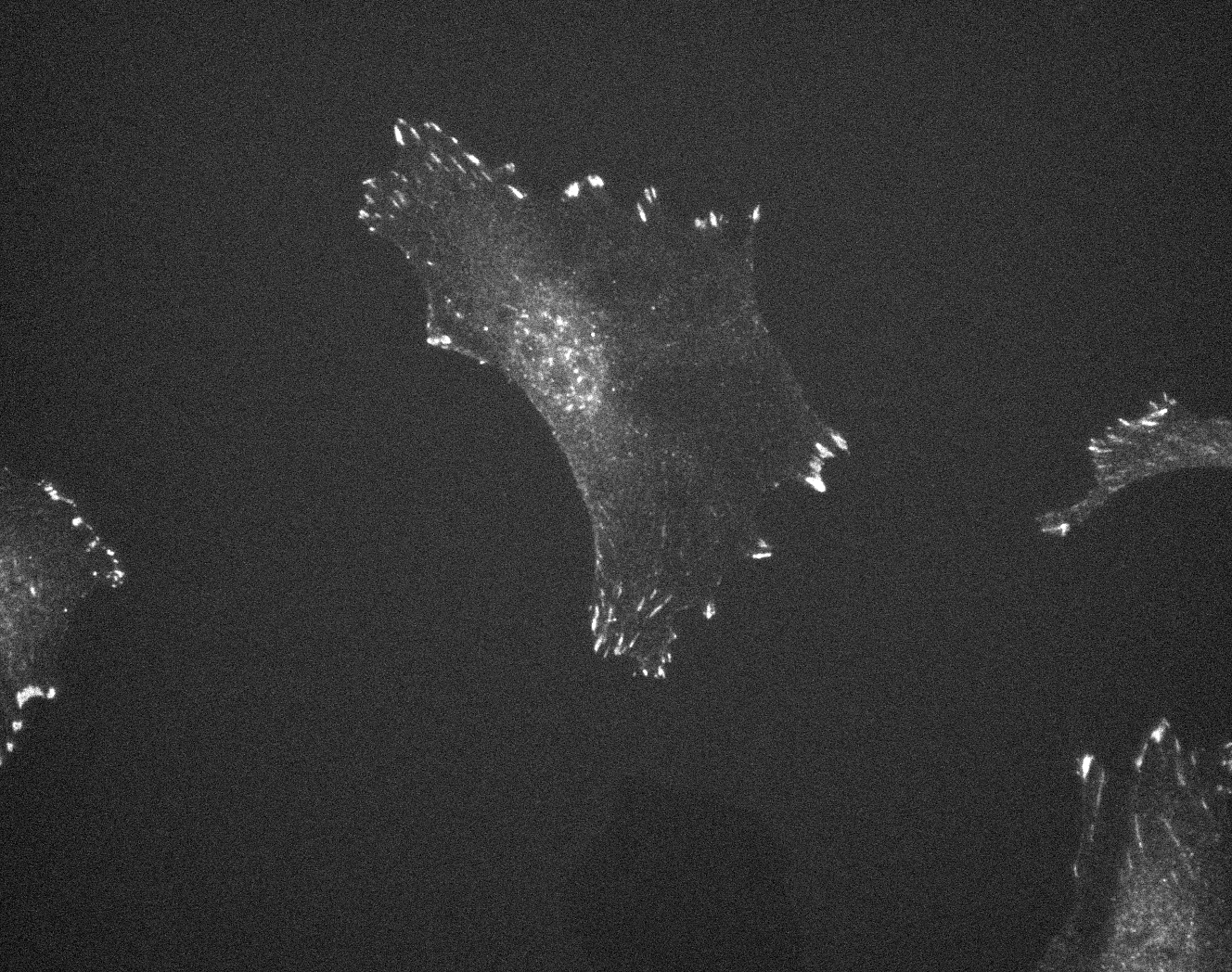

### B_Plate_R_p00_0_A01f14d0.TIF

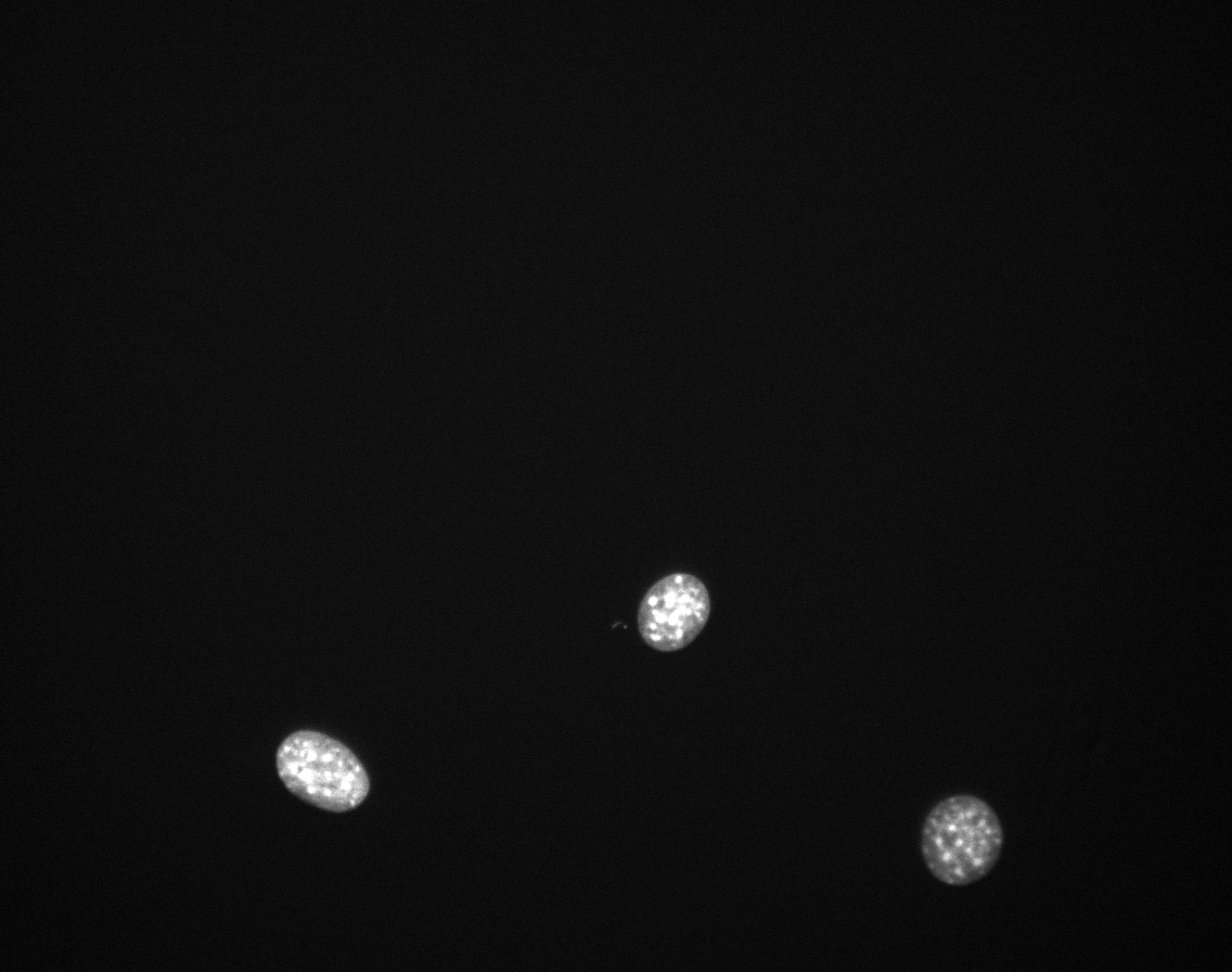

### B_Plate_R_p00_0_A01f14d1.TIF

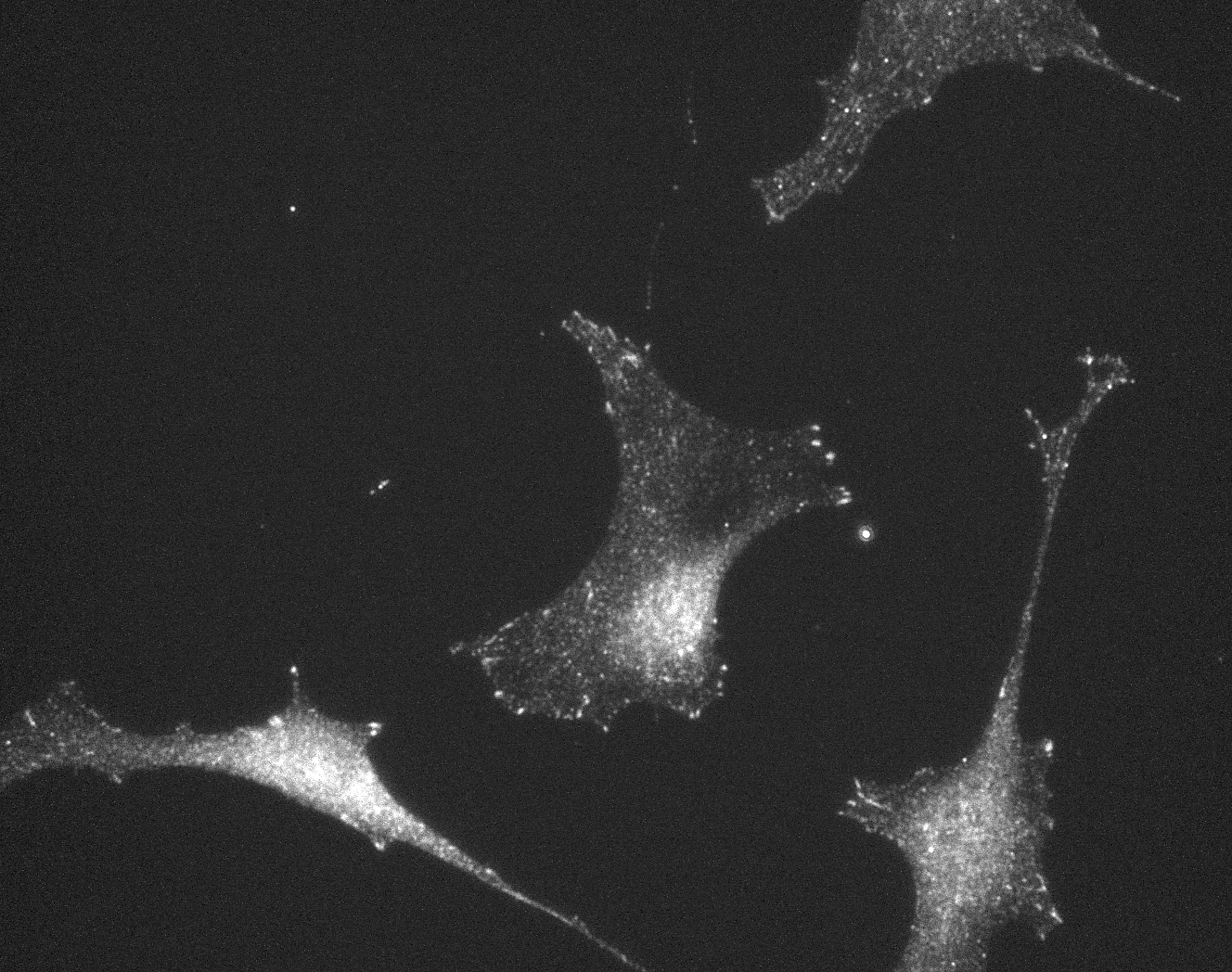

### B_Plate_R_p00_0_A01f14d2.TIF

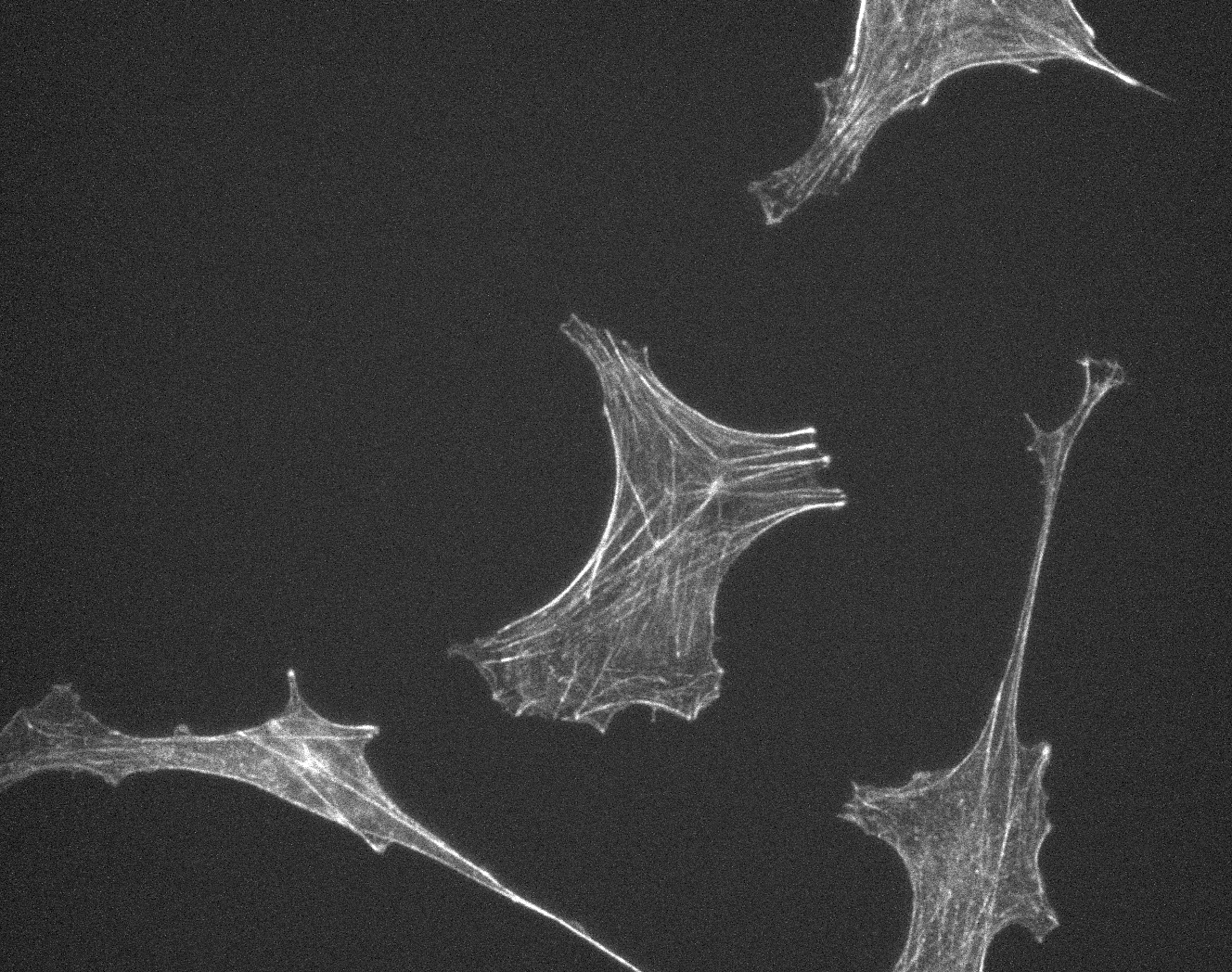

### B_Plate_R_p00_0_A01f14d3.TIF

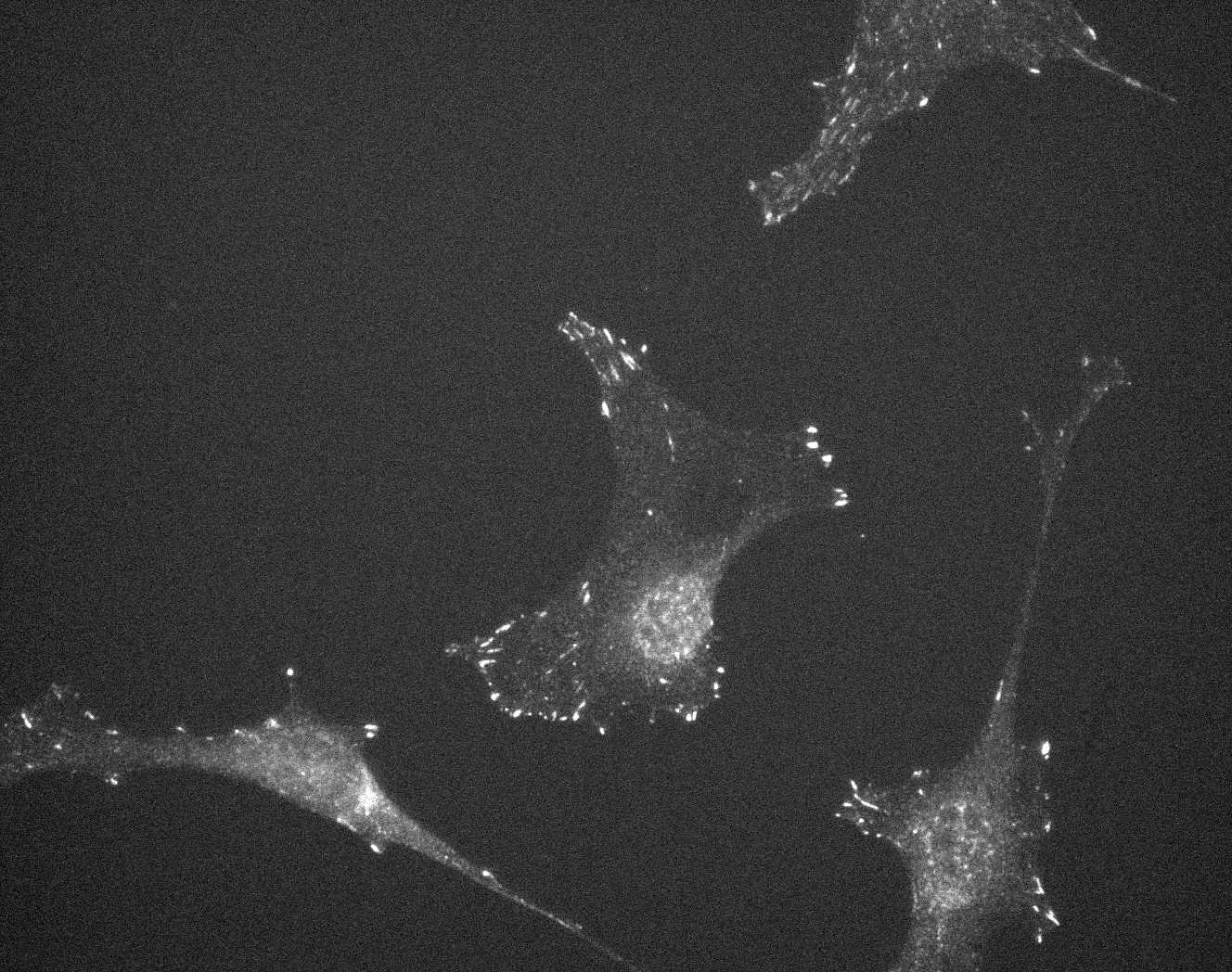

### B_Plate_R_p00_0_A01f14d3_FAOutline.tiff

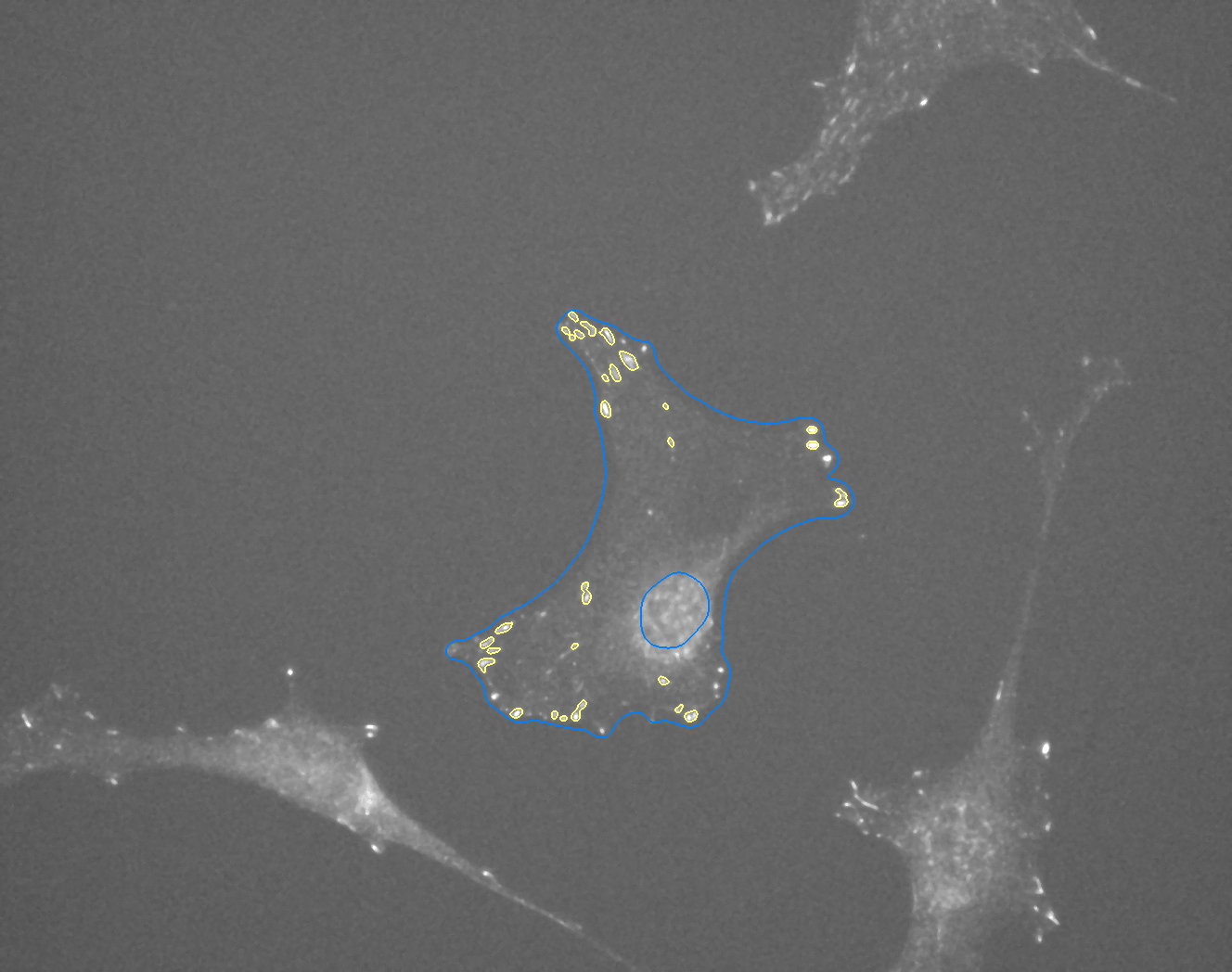

### B_Plate_R_p00_0_A01f37d0.TIF

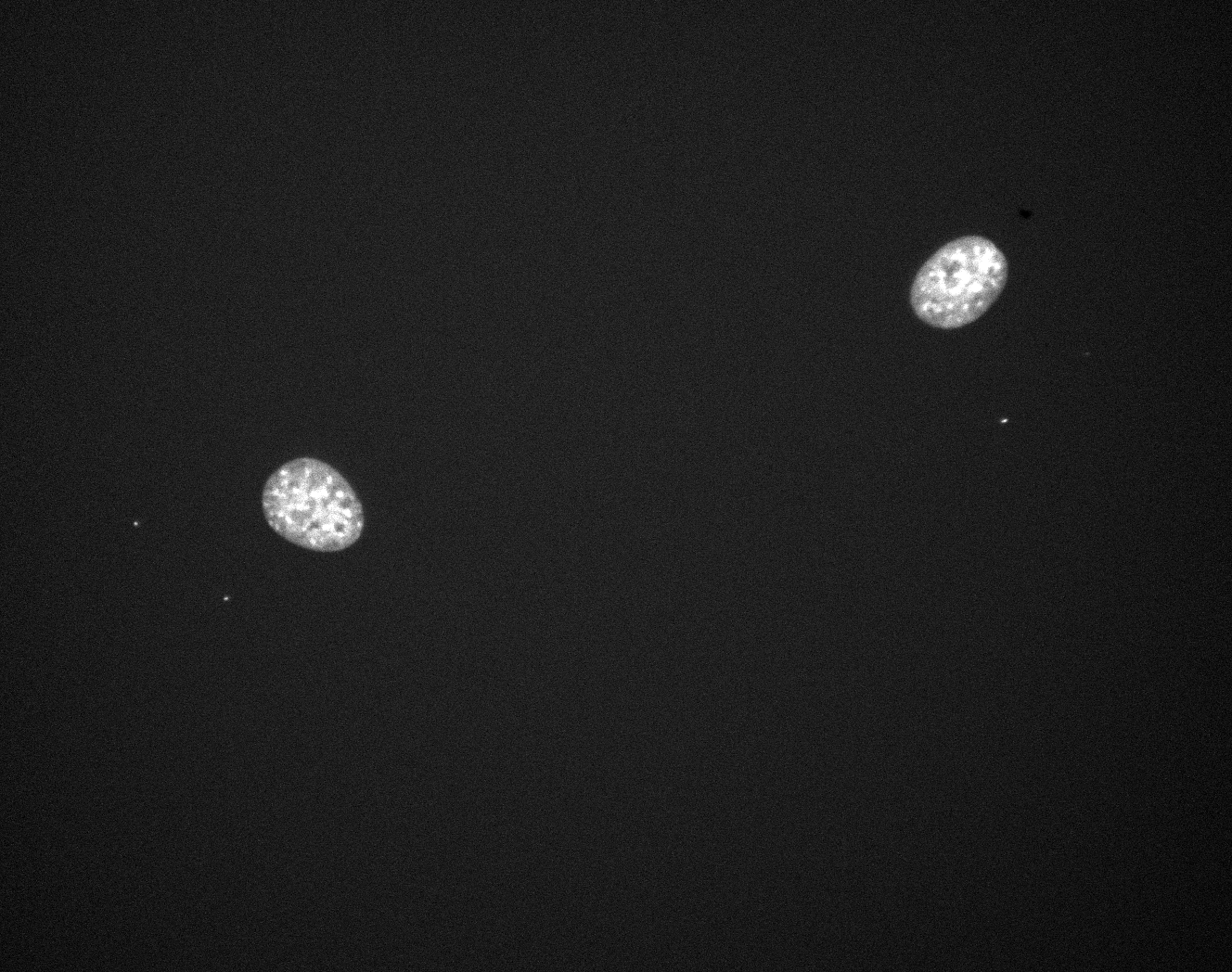

### B_Plate_R_p00_0_A01f37d1.TIF

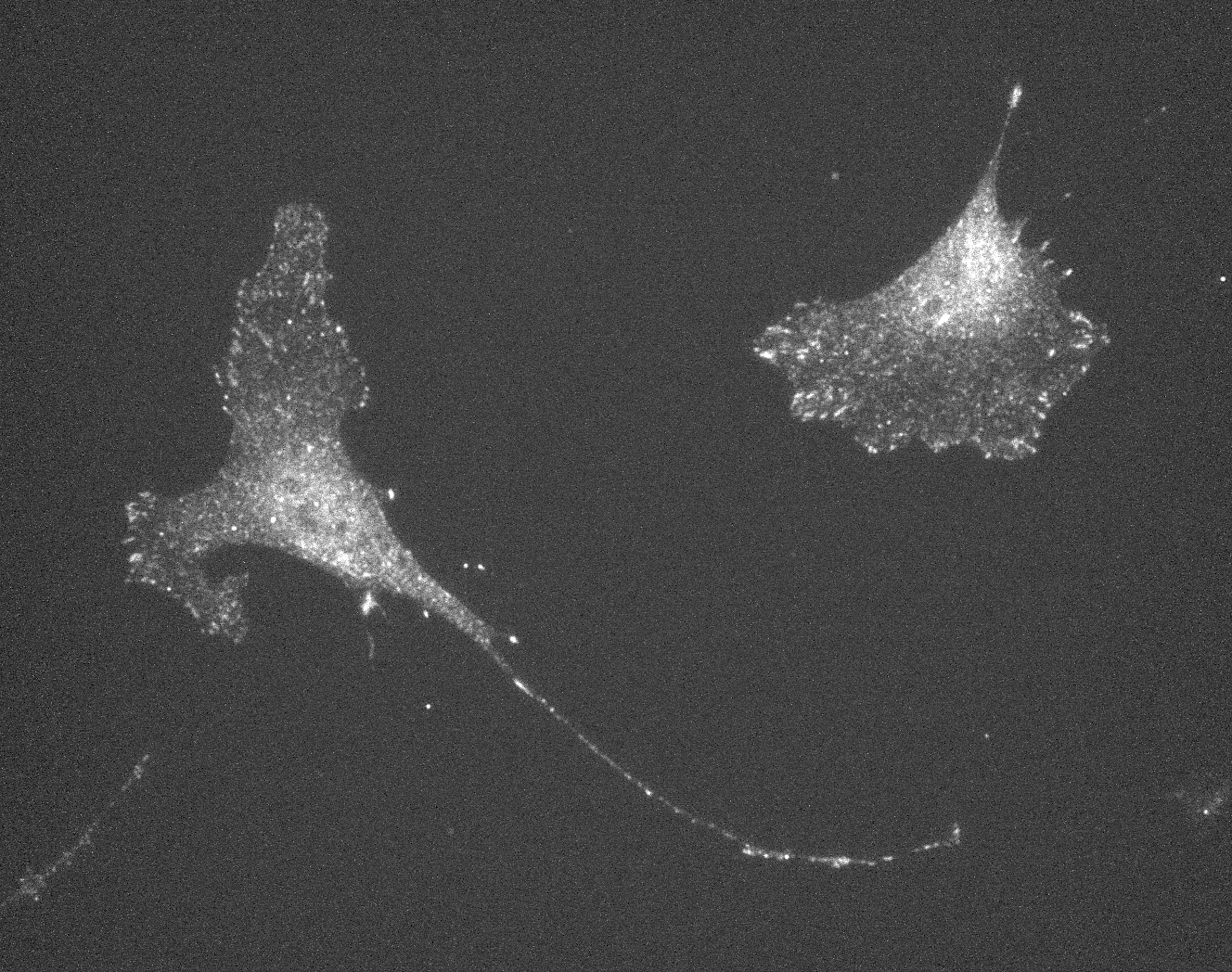

### B_Plate_R_p00_0_A01f37d2.TIF

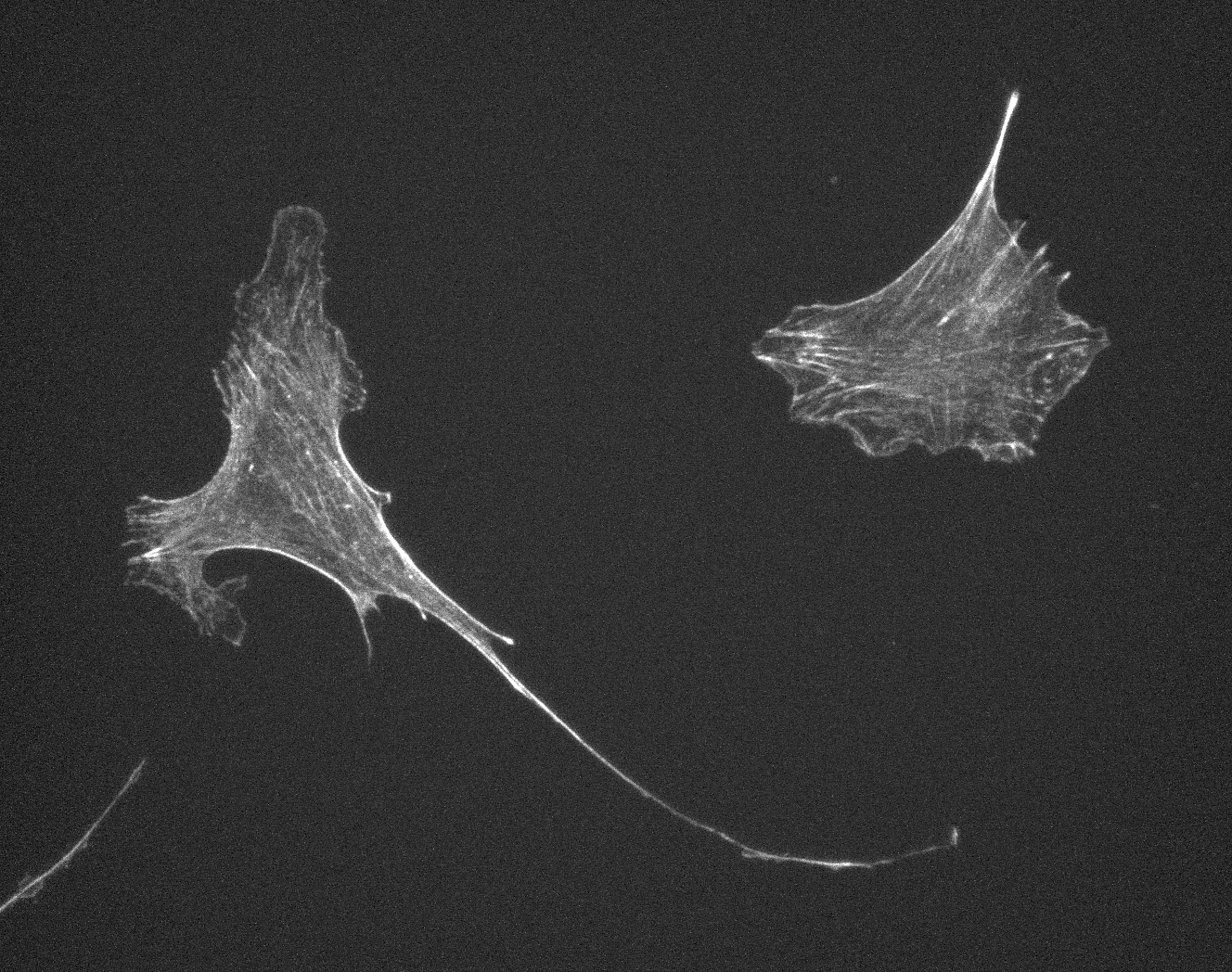

### B_Plate_R_p00_0_A01f37d3.TIF

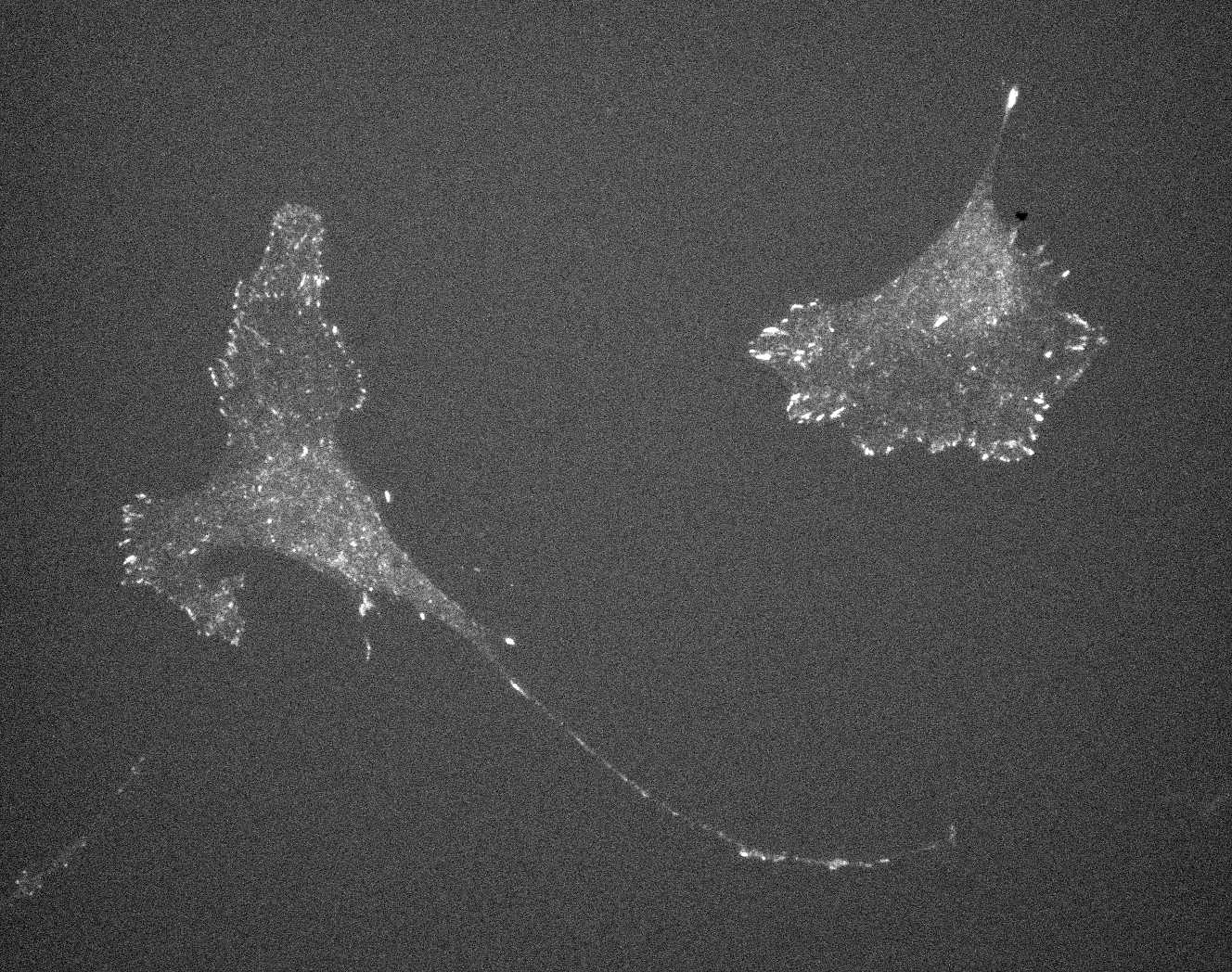

### C_Plate_R_p00_0_A01f05d0.TIF

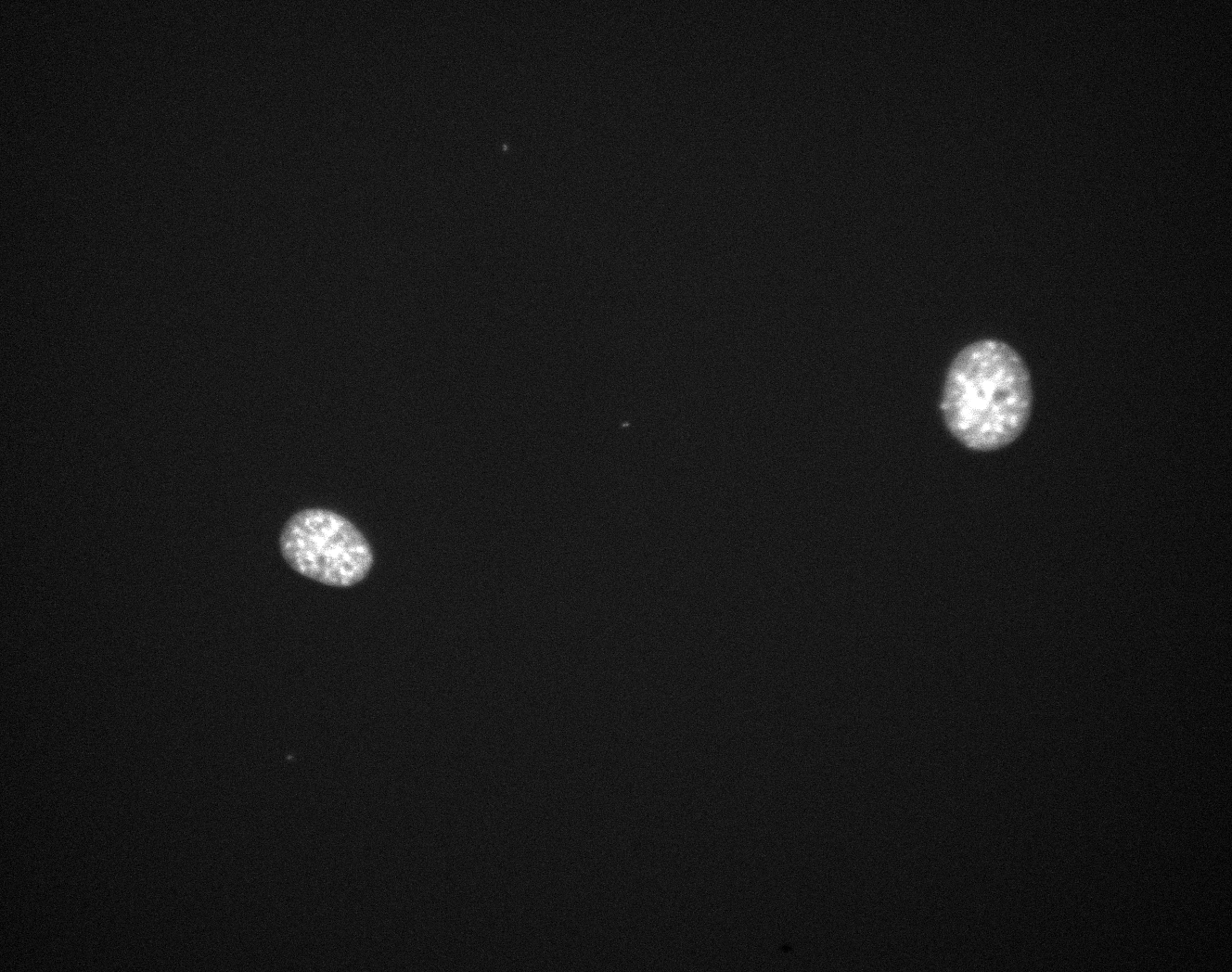

### C_Plate_R_p00_0_A01f05d1.TIF

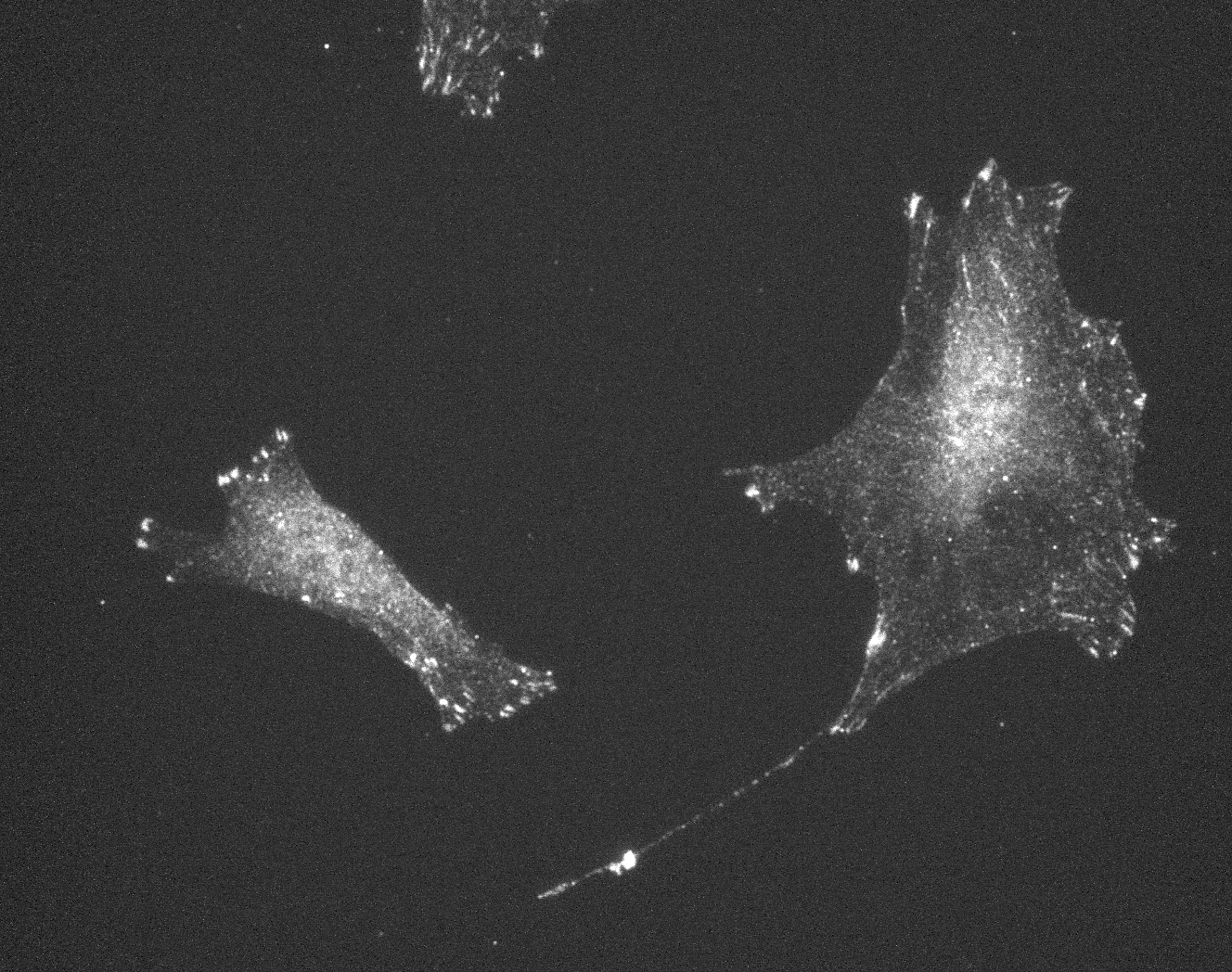

### C_Plate_R_p00_0_A01f05d2.TIF

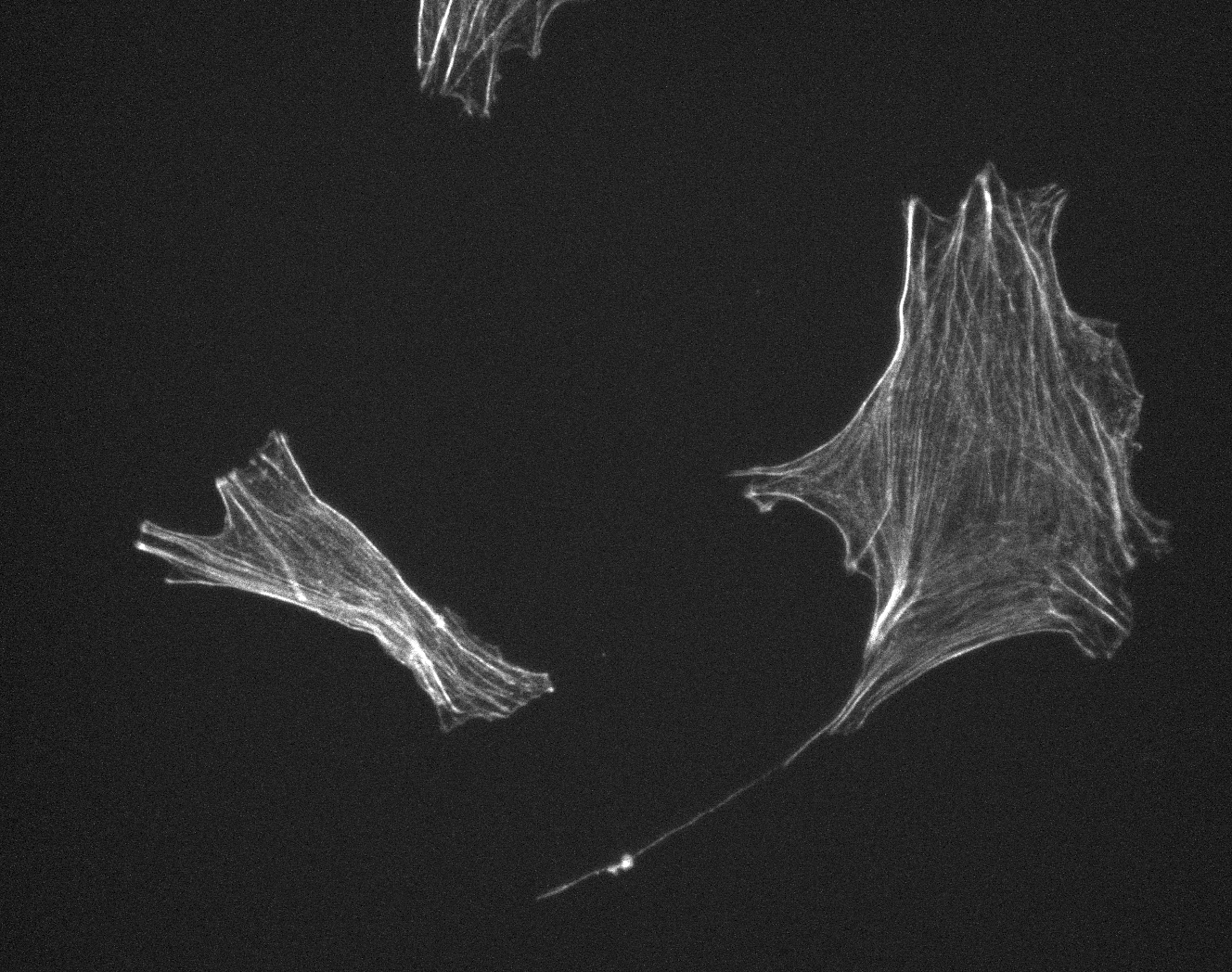

### C_Plate_R_p00_0_A01f05d2_CytoplasmOutline.tiff

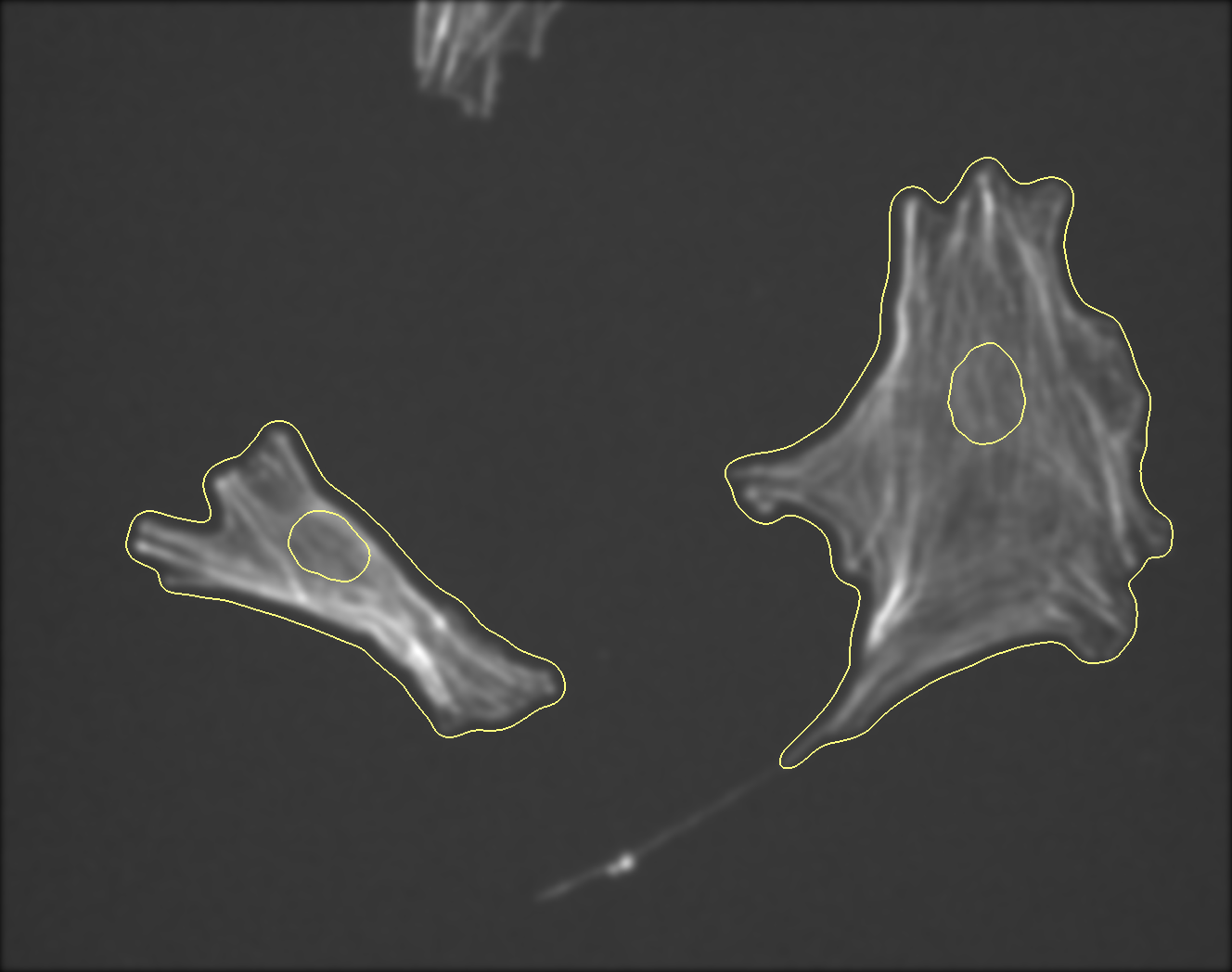

### C_Plate_R_p00_0_A01f05d3.TIF

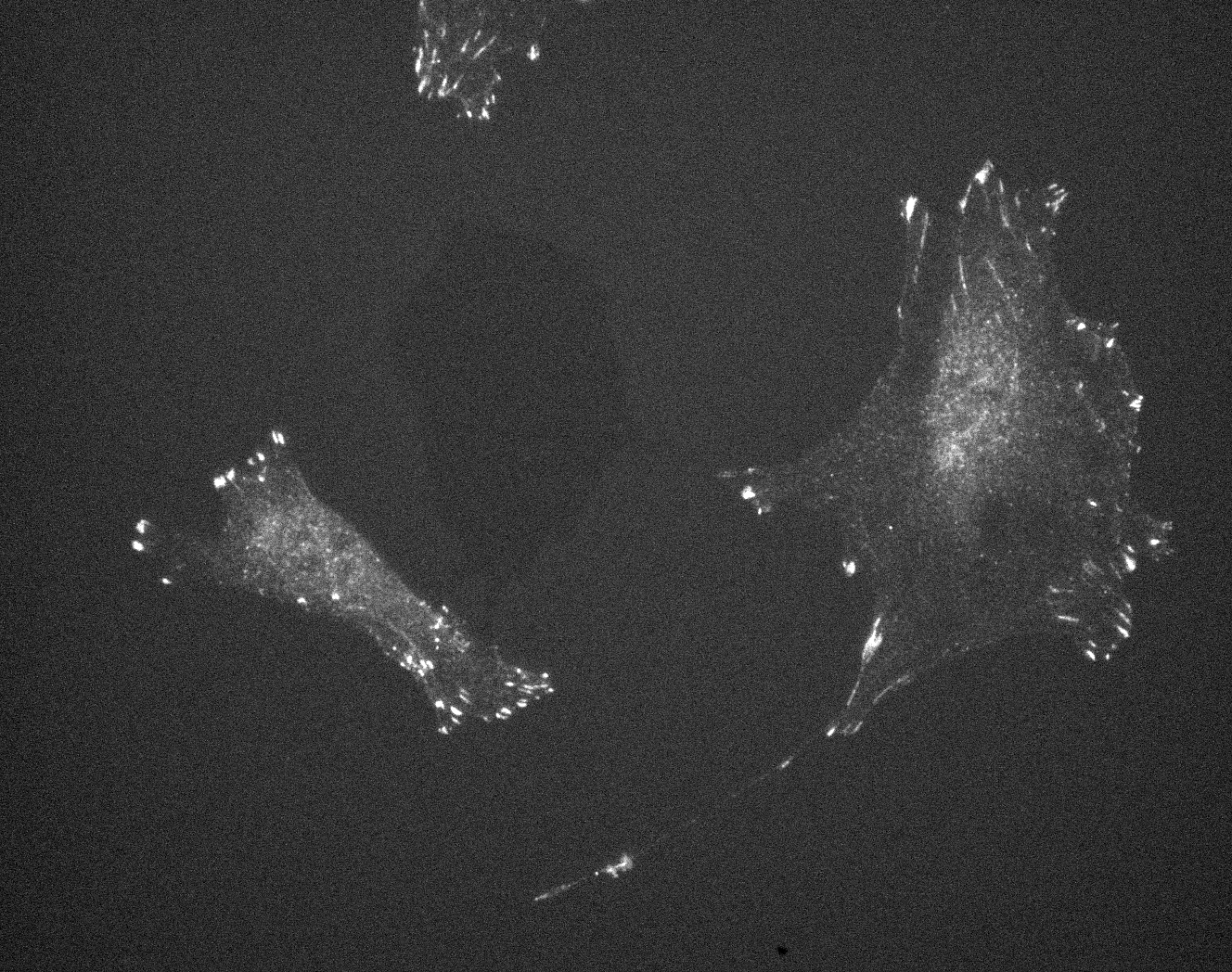

### C_Plate_R_p00_0_A01f17d3_FAOutline.tiff

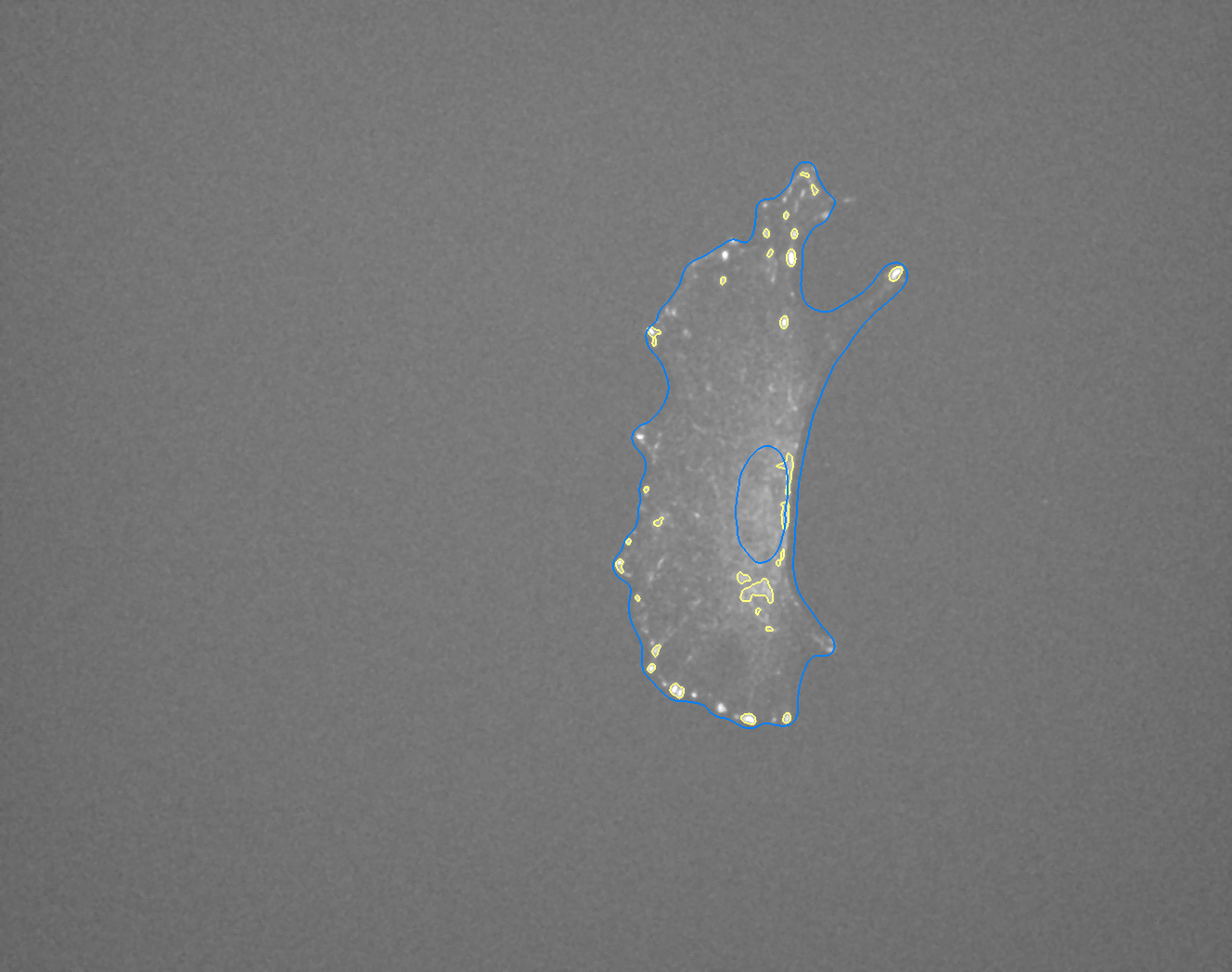

### C_Plate_R_p00_0_A01f30d0.TIF

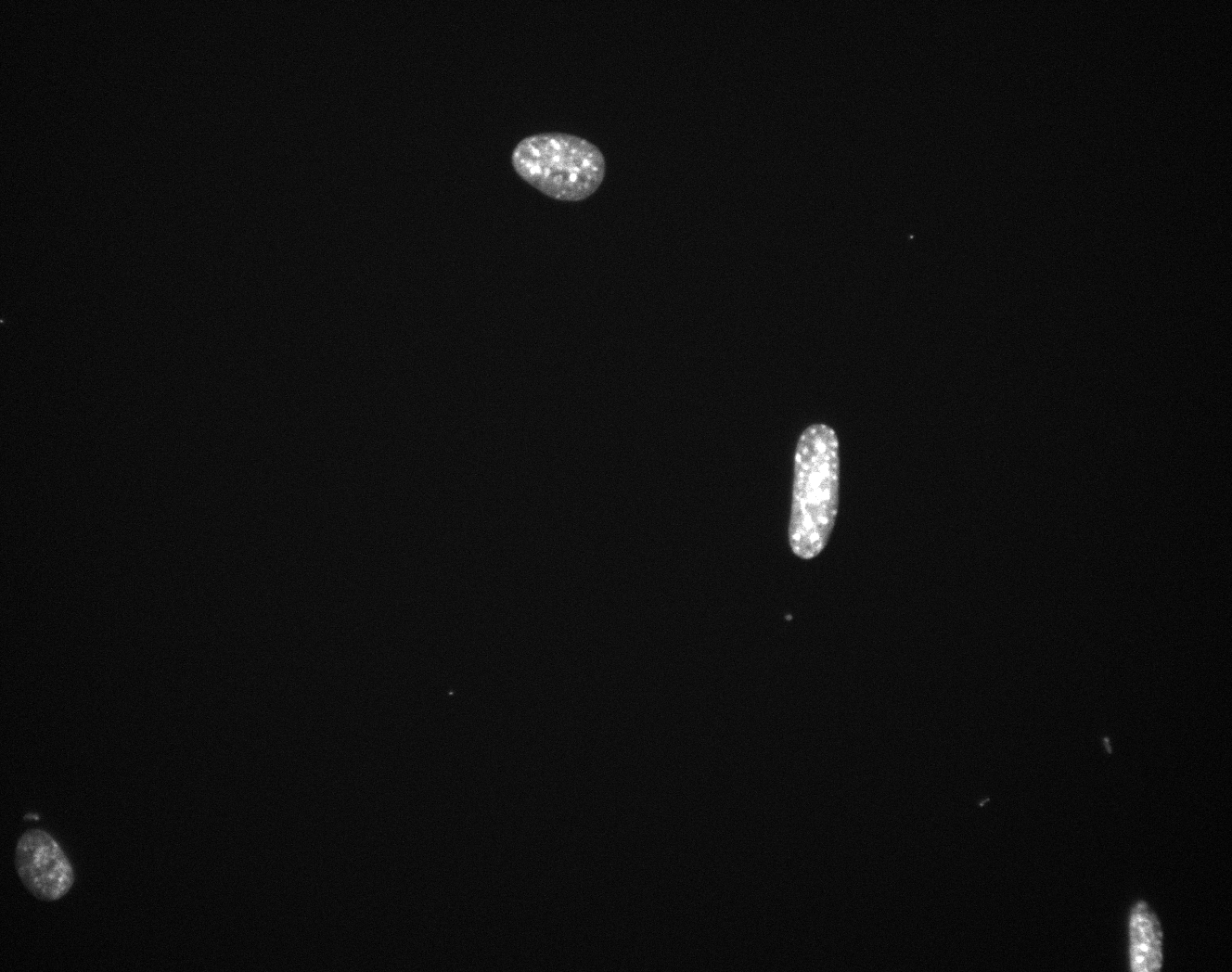

### C_Plate_R_p00_0_A01f30d1.TIF

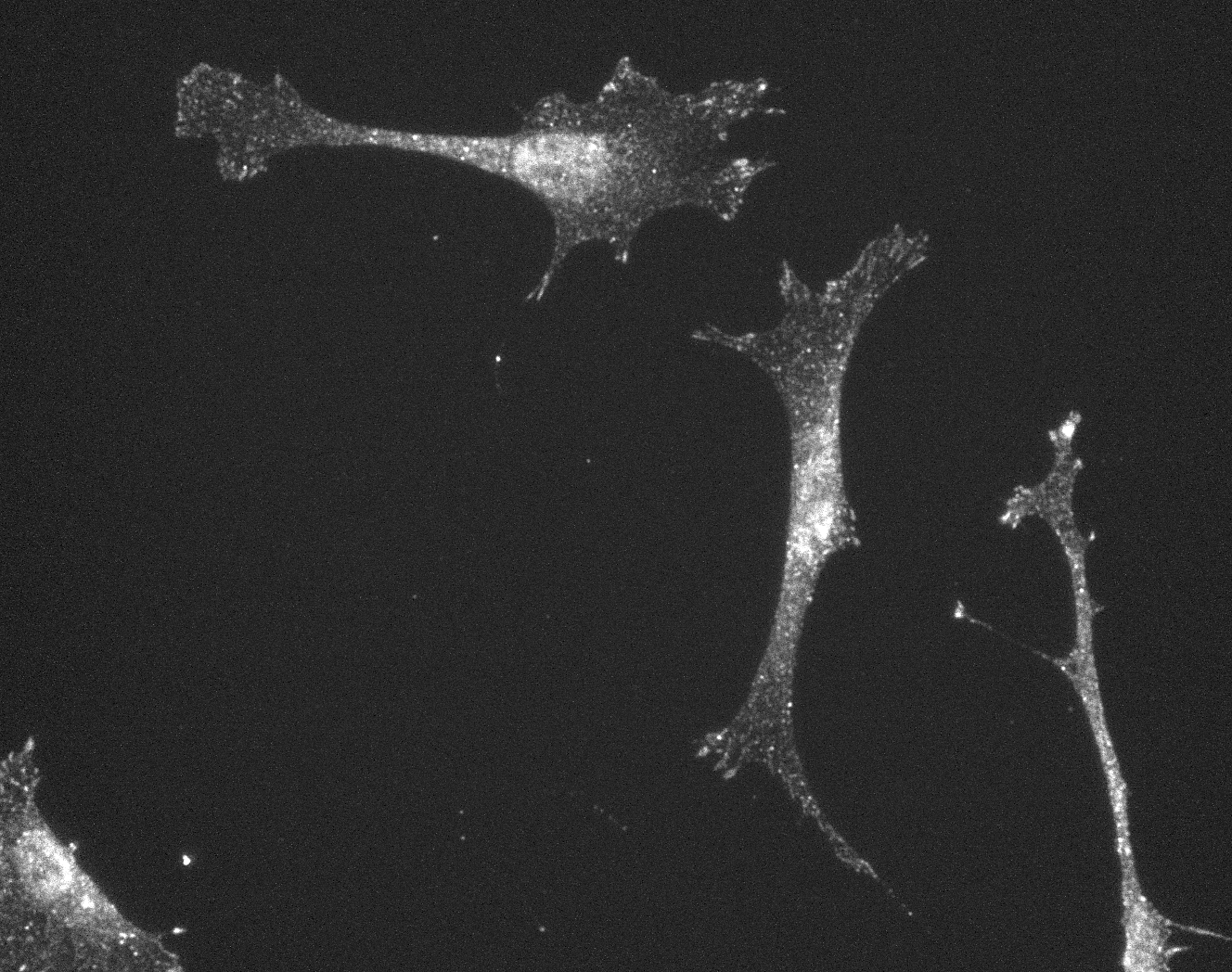

### C_Plate_R_p00_0_A01f30d2.TIF

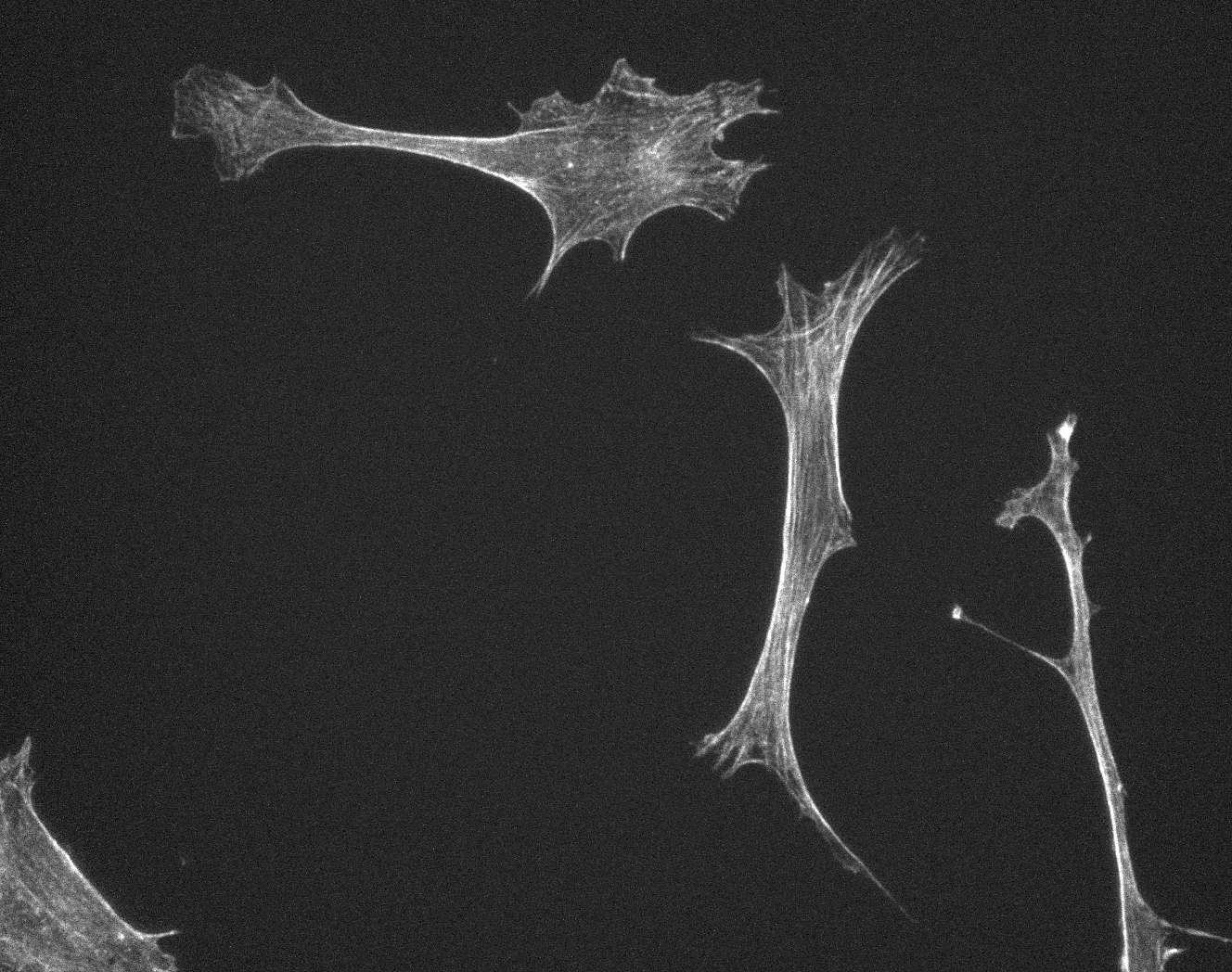

### C_Plate_R_p00_0_A01f30d3.TIF

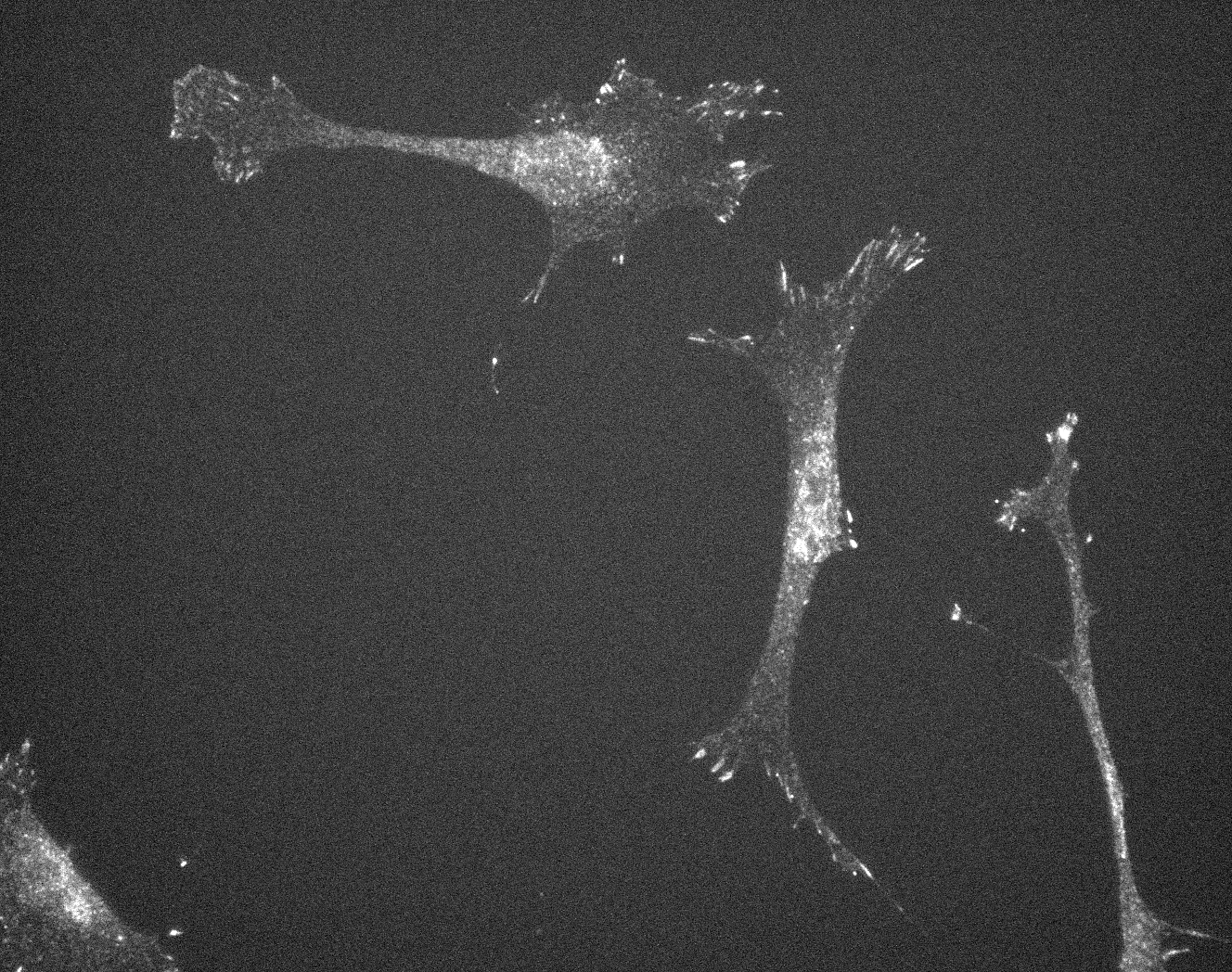

### D_Plate_R_p00_0_A01f00d0.TIF

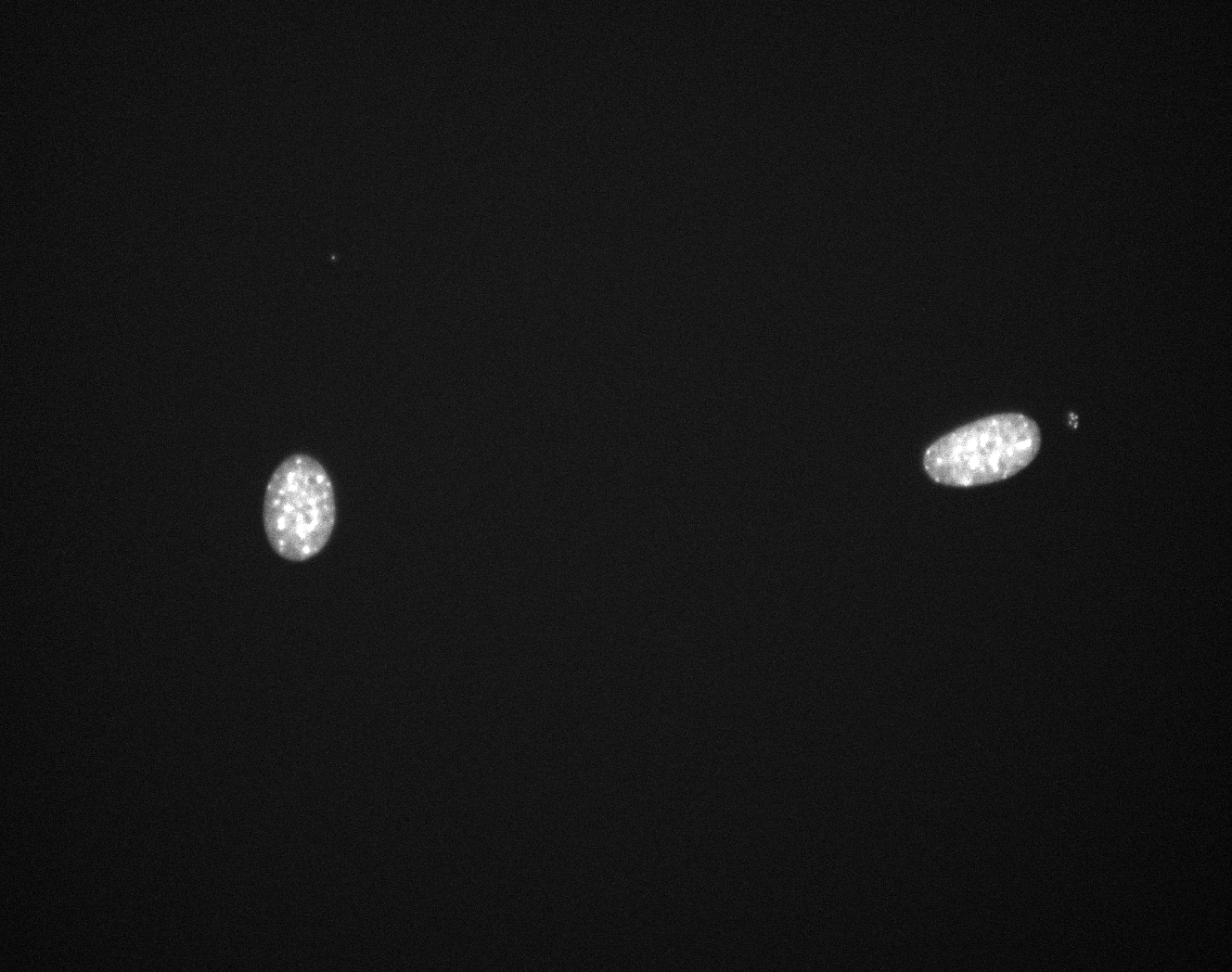

### D_Plate_R_p00_0_A01f00d1.TIF

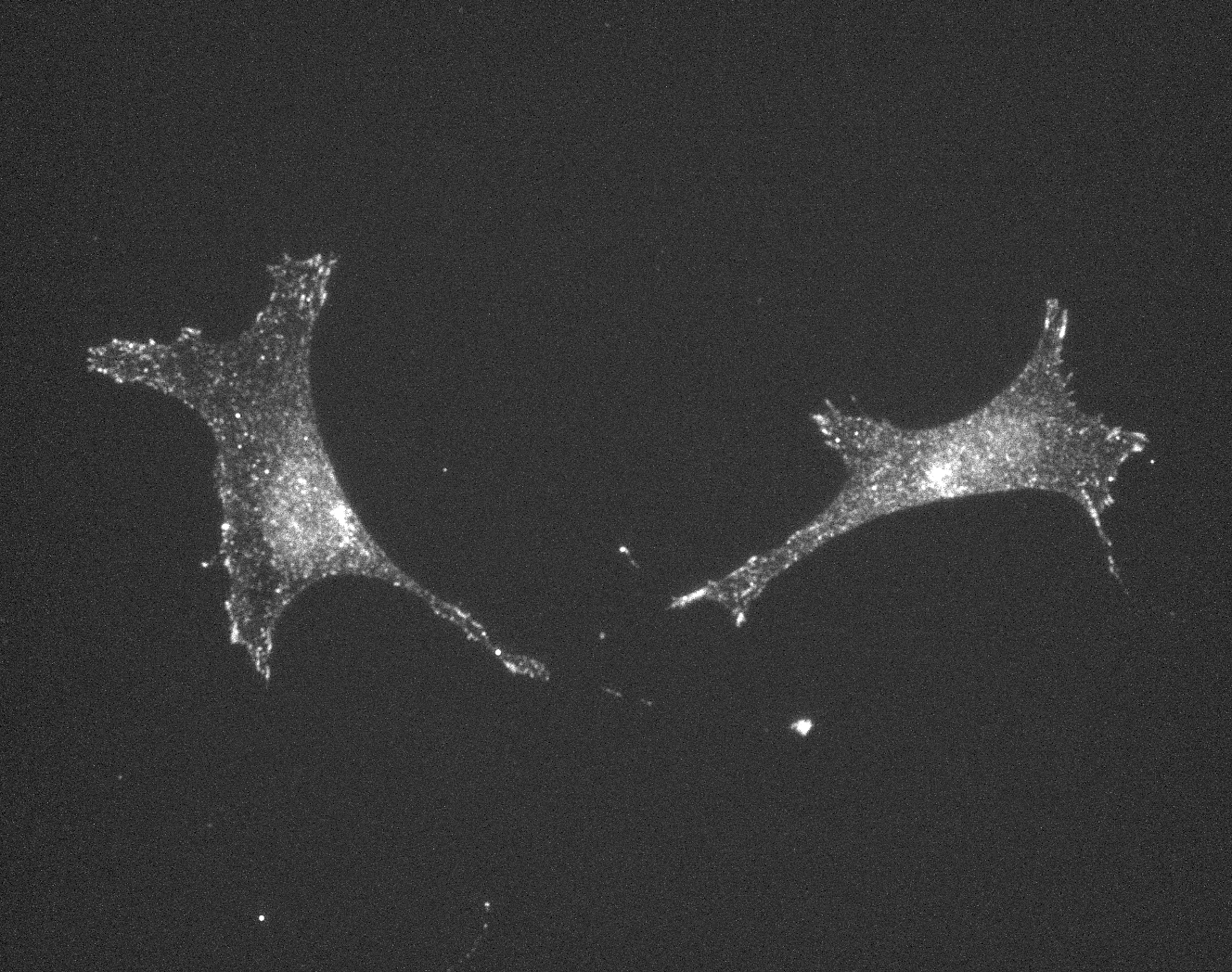

### D_Plate_R_p00_0_A01f00d2.TIF

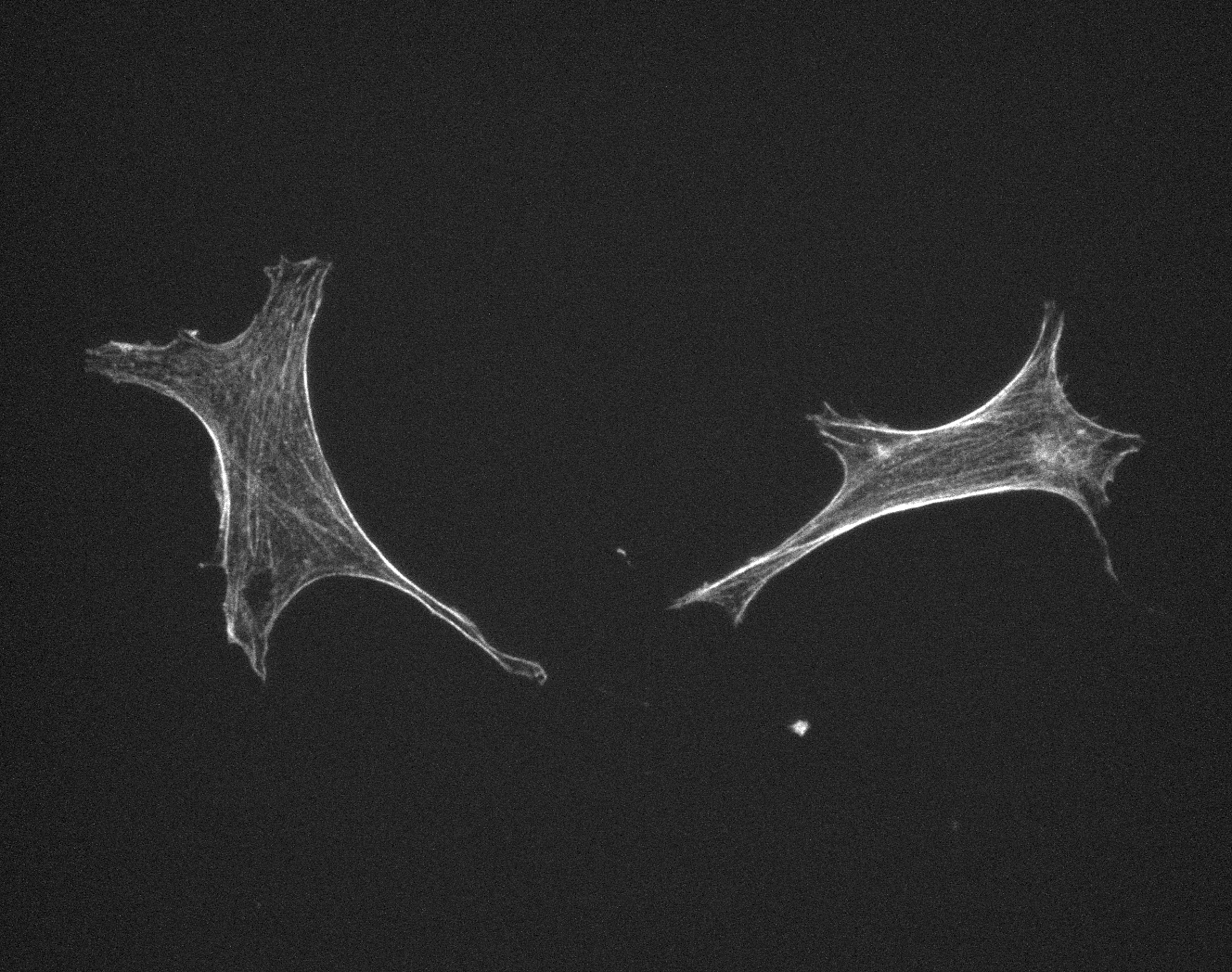

### D_Plate_R_p00_0_A01f00d2_CytoplasmOutline.tiff

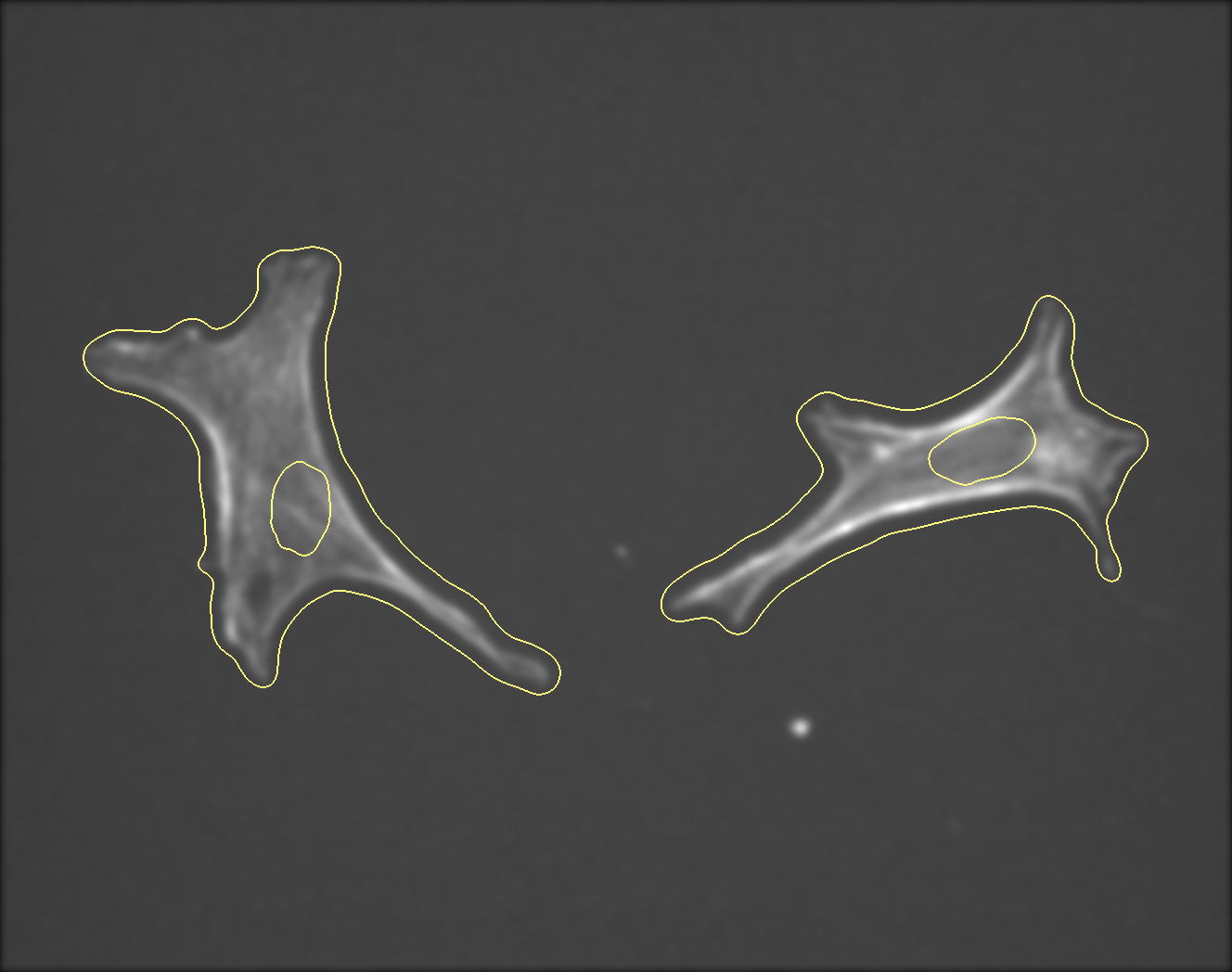

### D_Plate_R_p00_0_A01f00d3.TIF

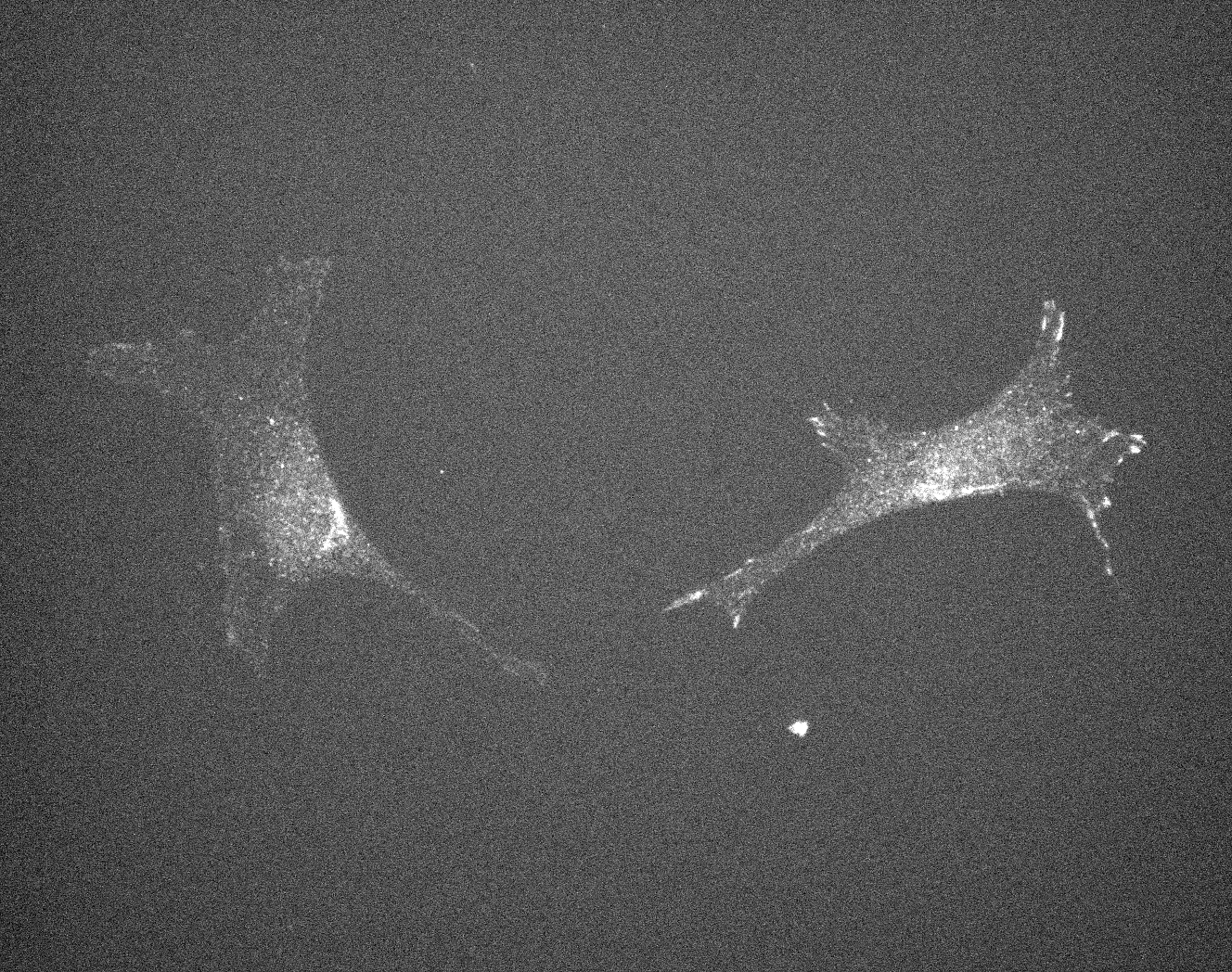

### D_Plate_R_p00_0_A01f48d0.TIF

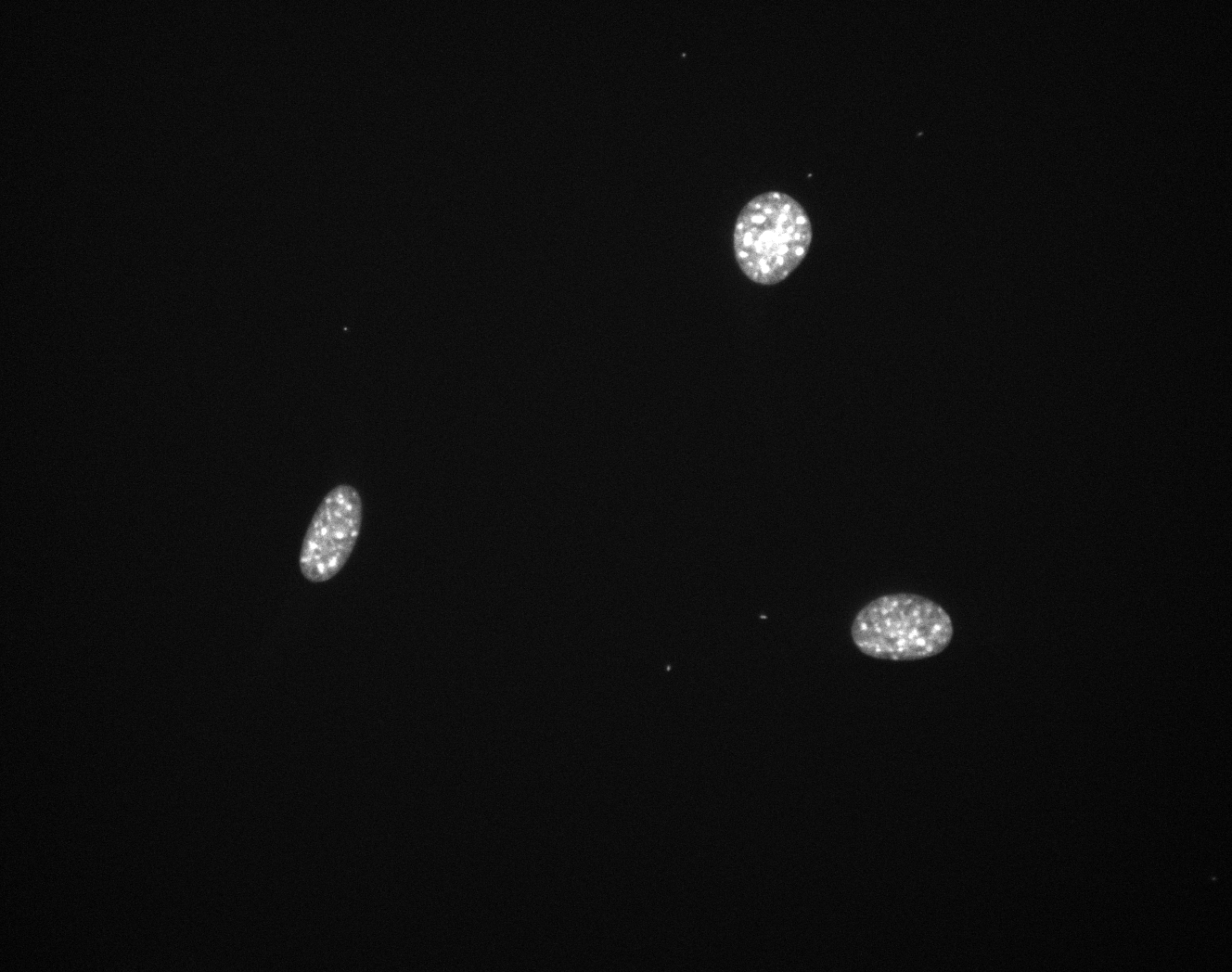

### D_Plate_R_p00_0_A01f48d1.TIF

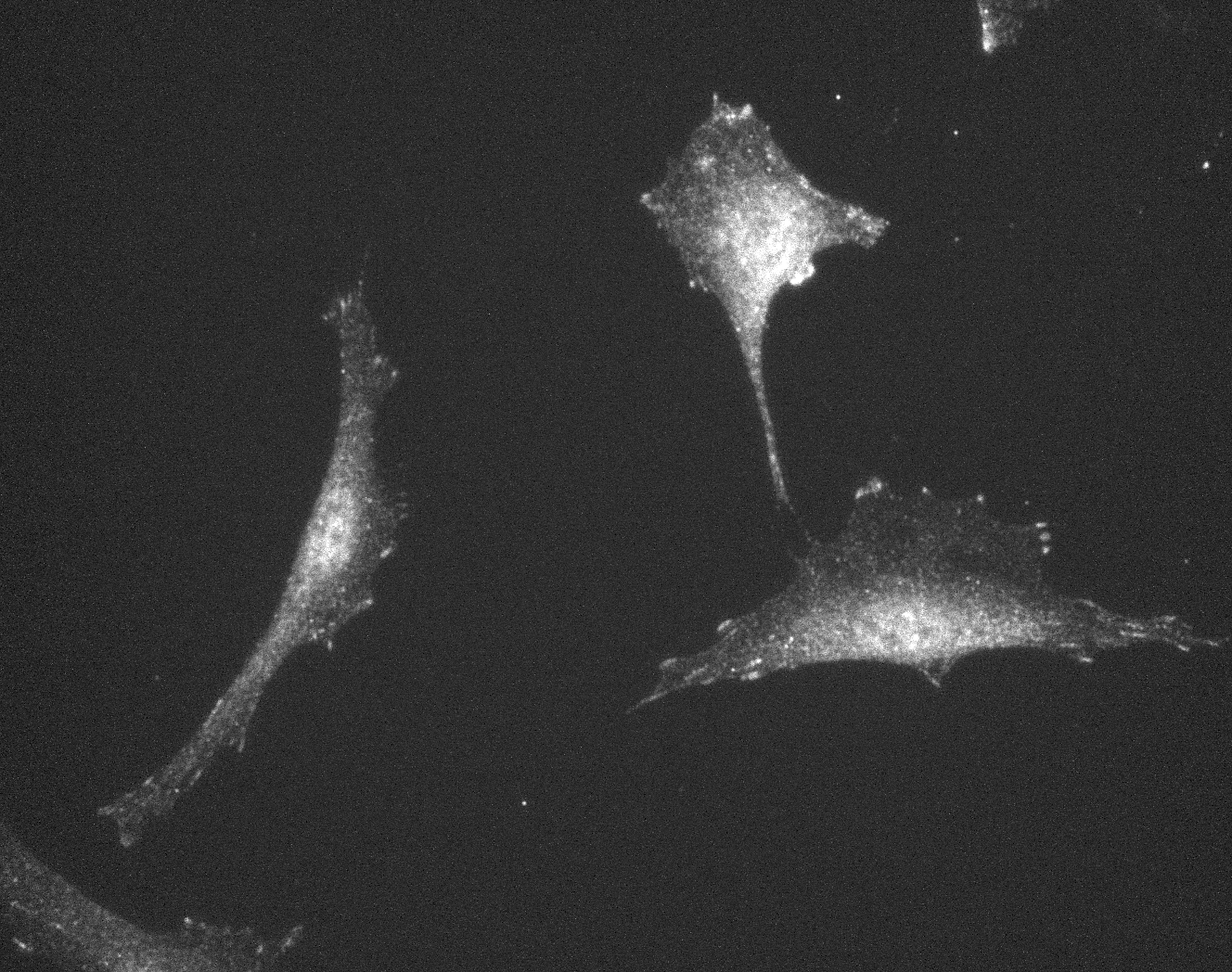

### D_Plate_R_p00_0_A01f48d2.TIF

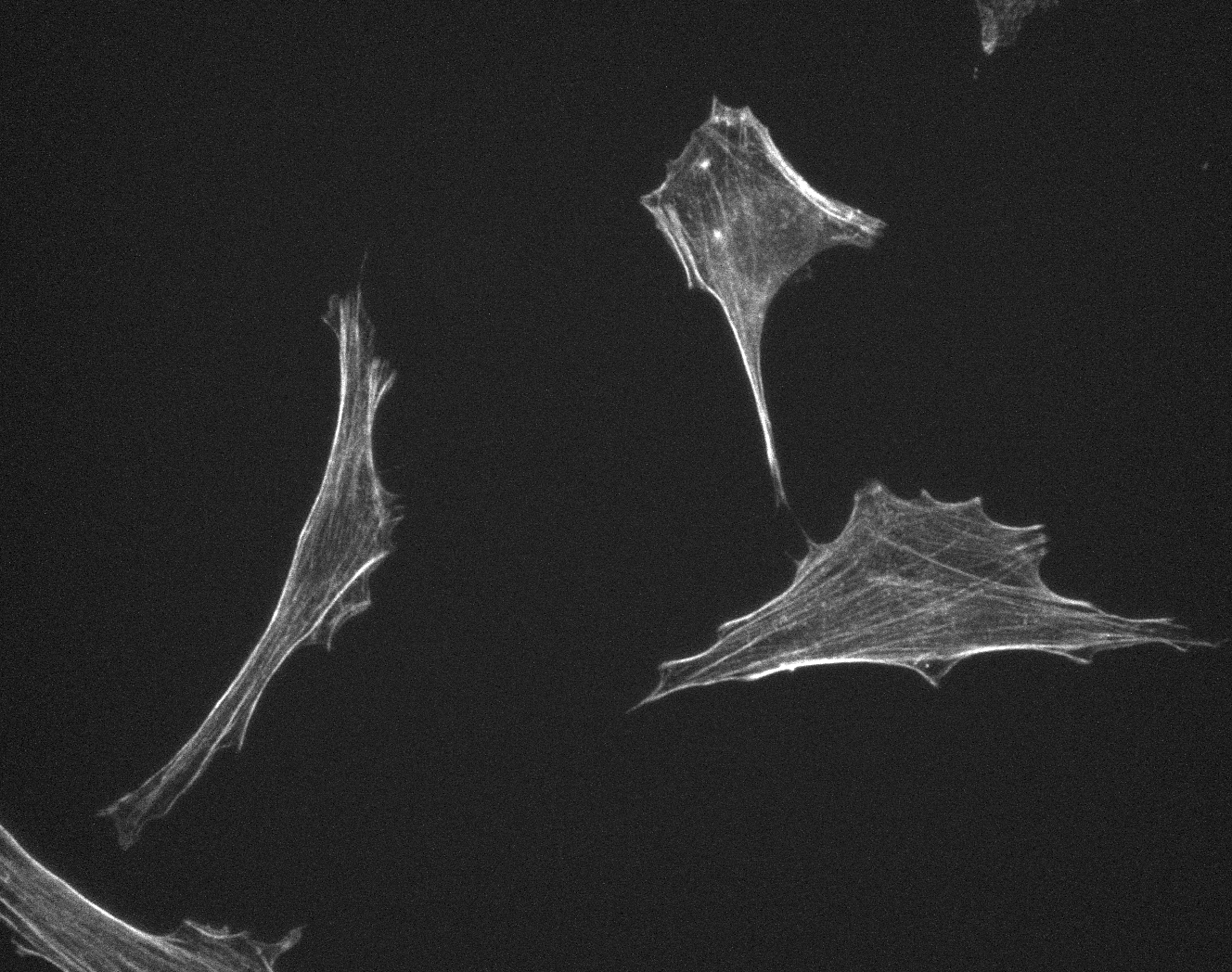

### D_Plate_R_p00_0_A01f48d3.TIF

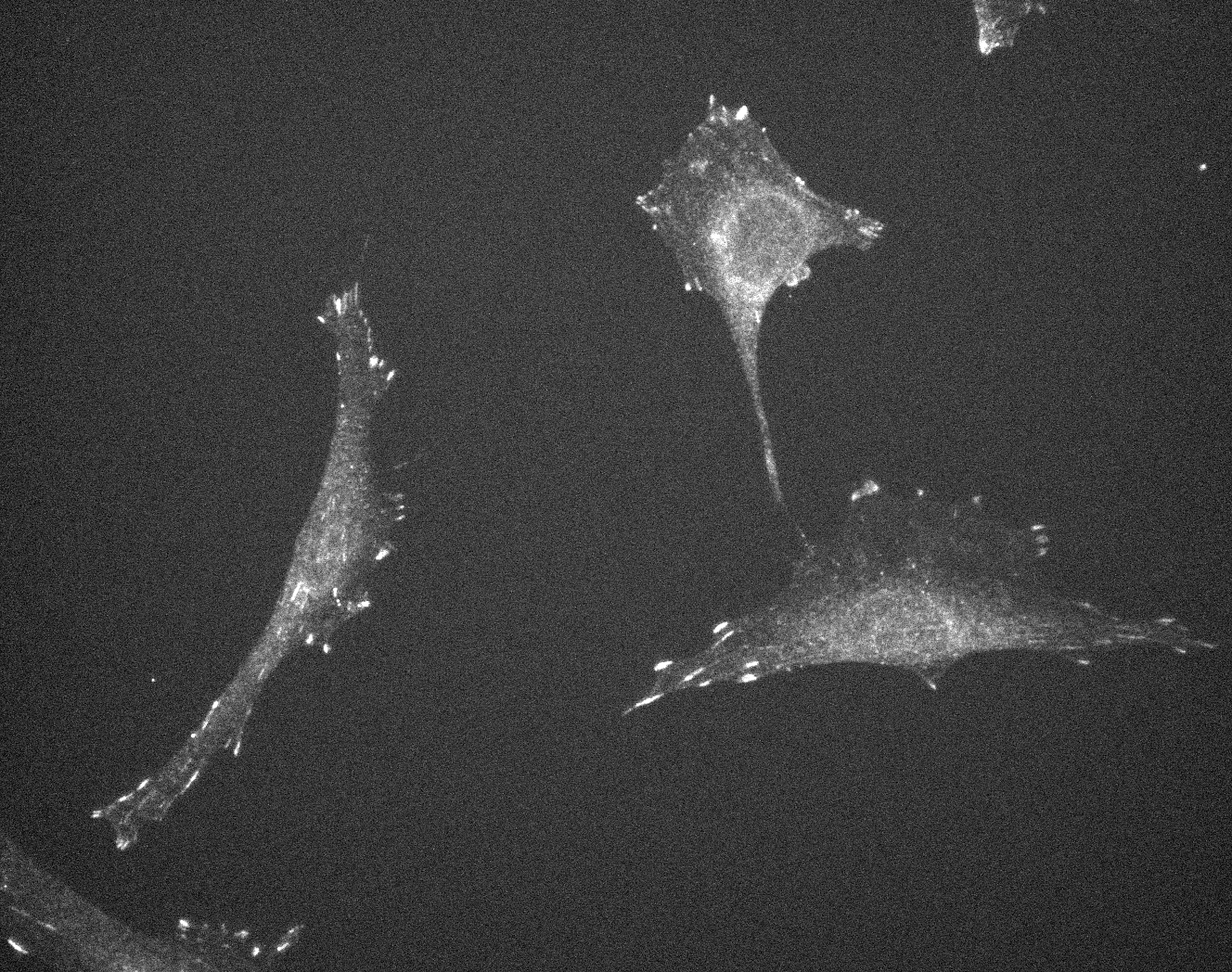

### D_Plate_R_p00_0_A01f59d3_FAOutline.tiff

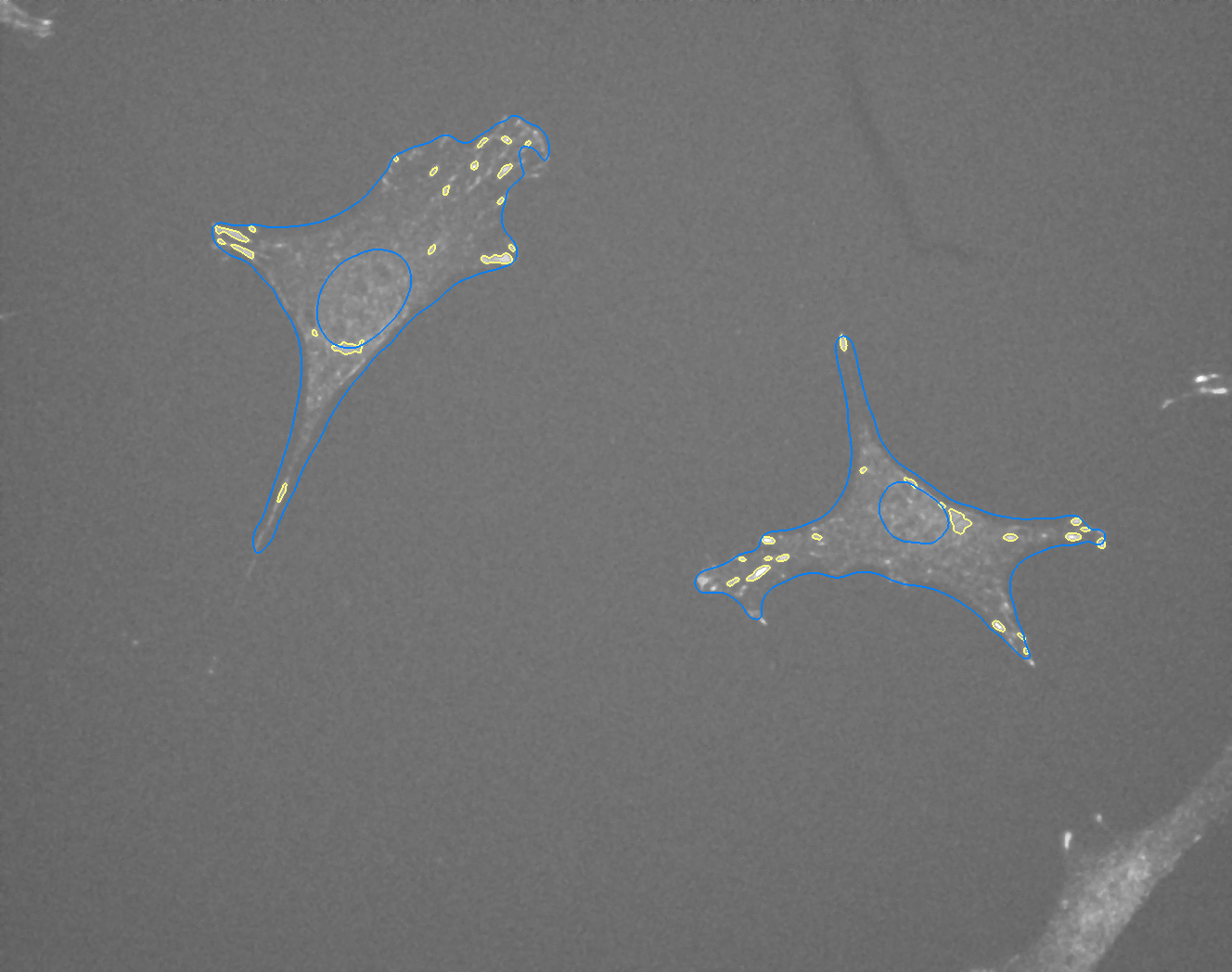

### E_Plate_R_p00_0_A01f12d0.TIF

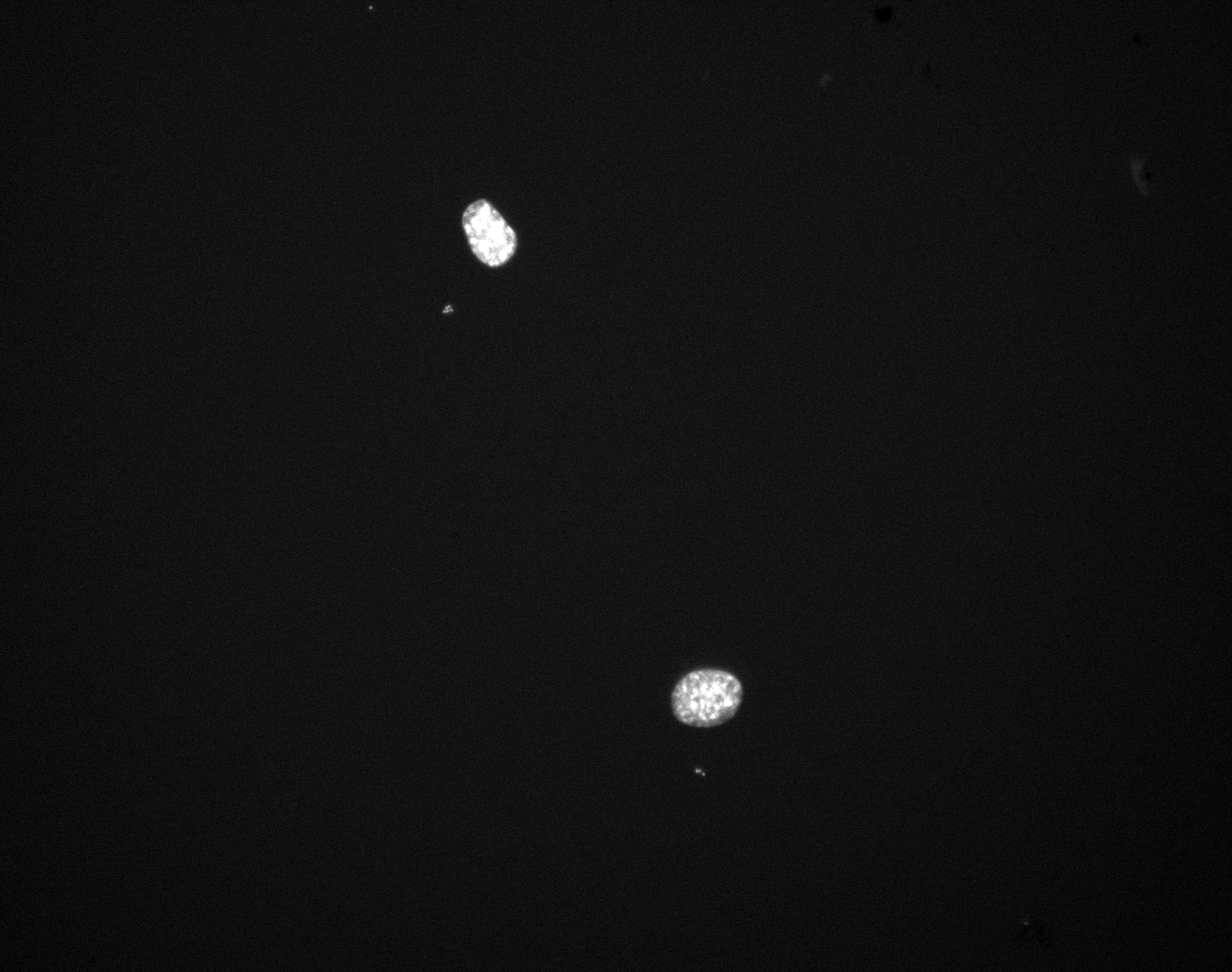

### E_Plate_R_p00_0_A01f12d1.TIF

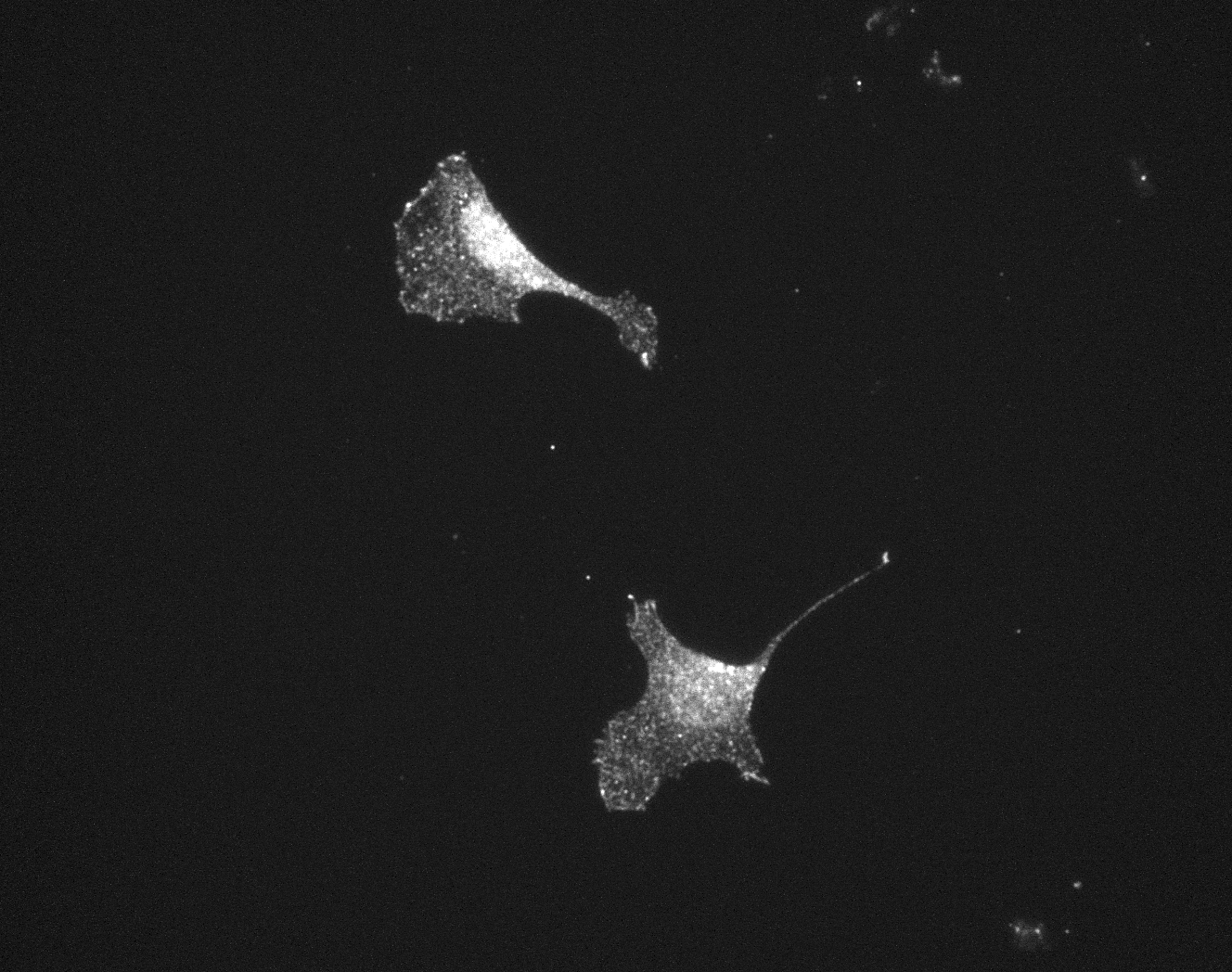

### E_Plate_R_p00_0_A01f12d2.TIF

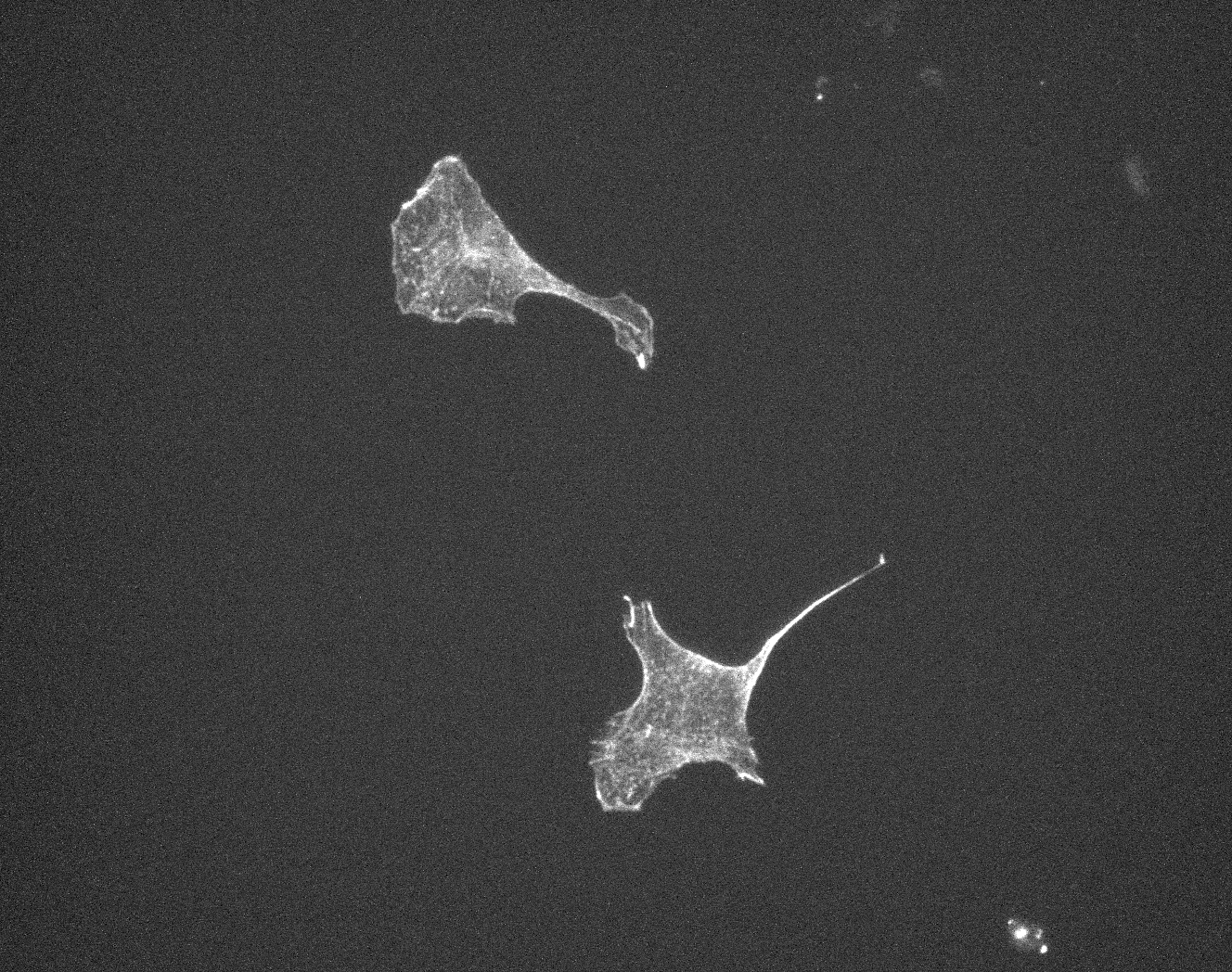

### E_Plate_R_p00_0_A01f12d3.TIF

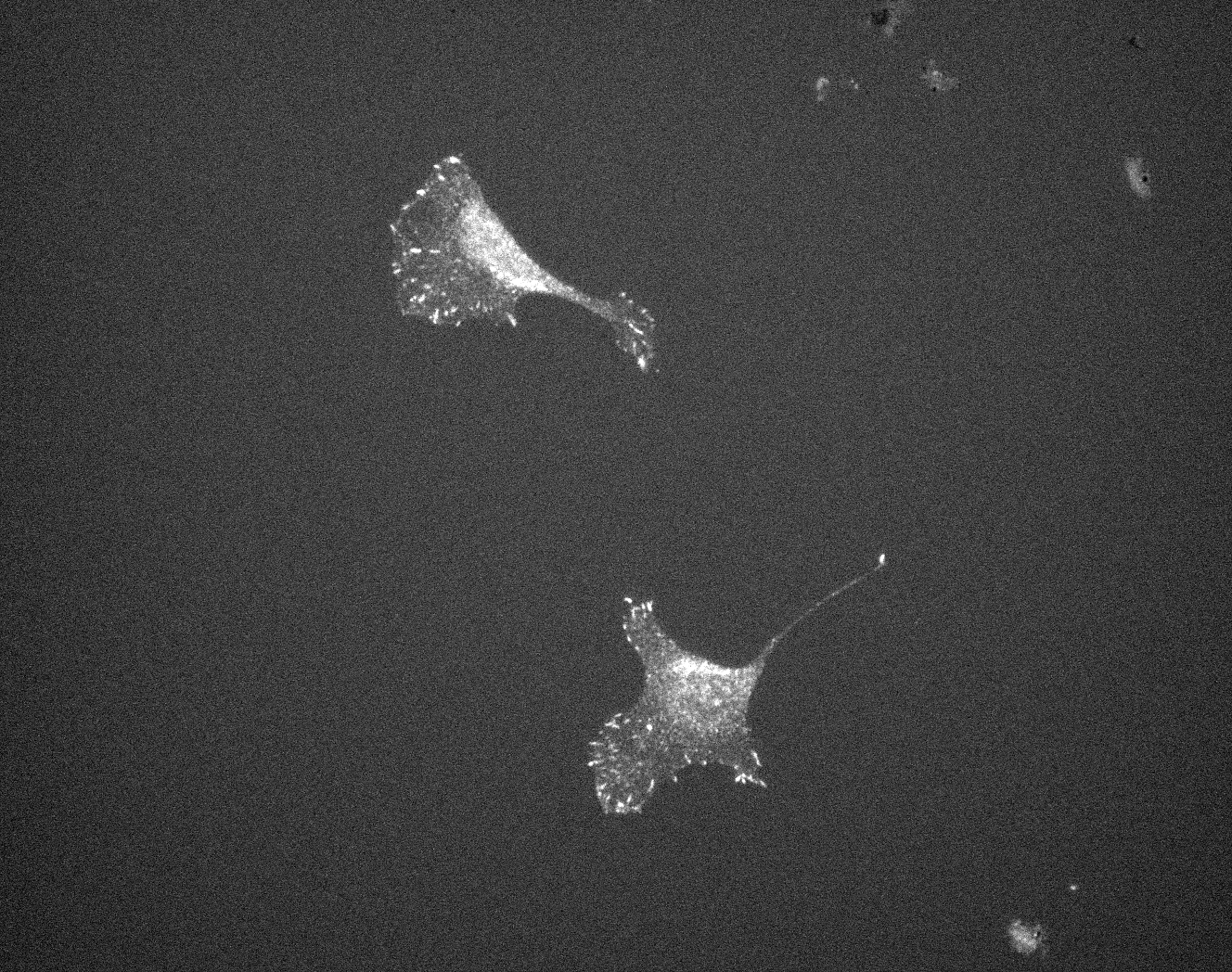

### E_Plate_R_p00_0_A01f16d3_FAOutline.tiff

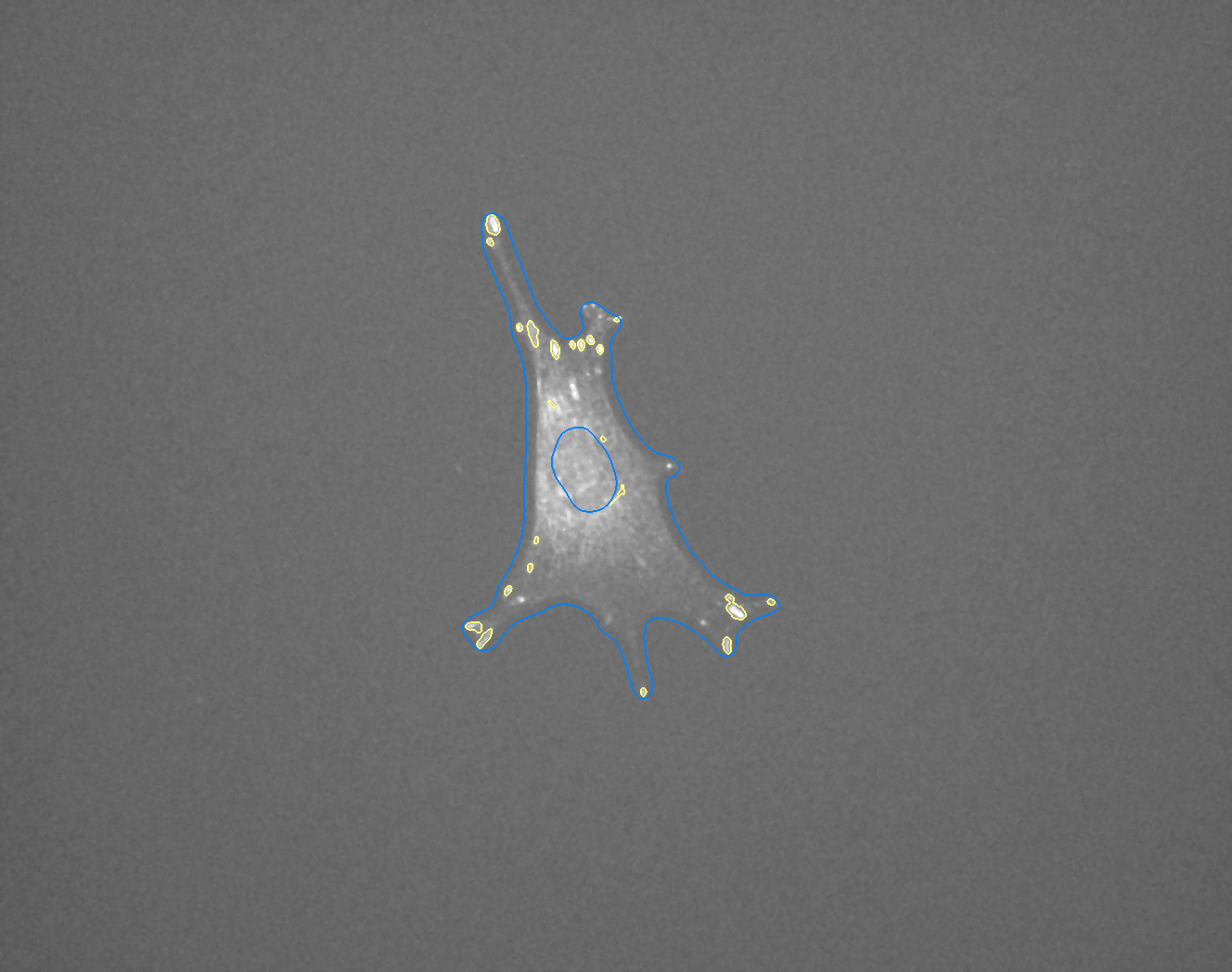

### E_Plate_R_p00_0_A01f54d0.TIF

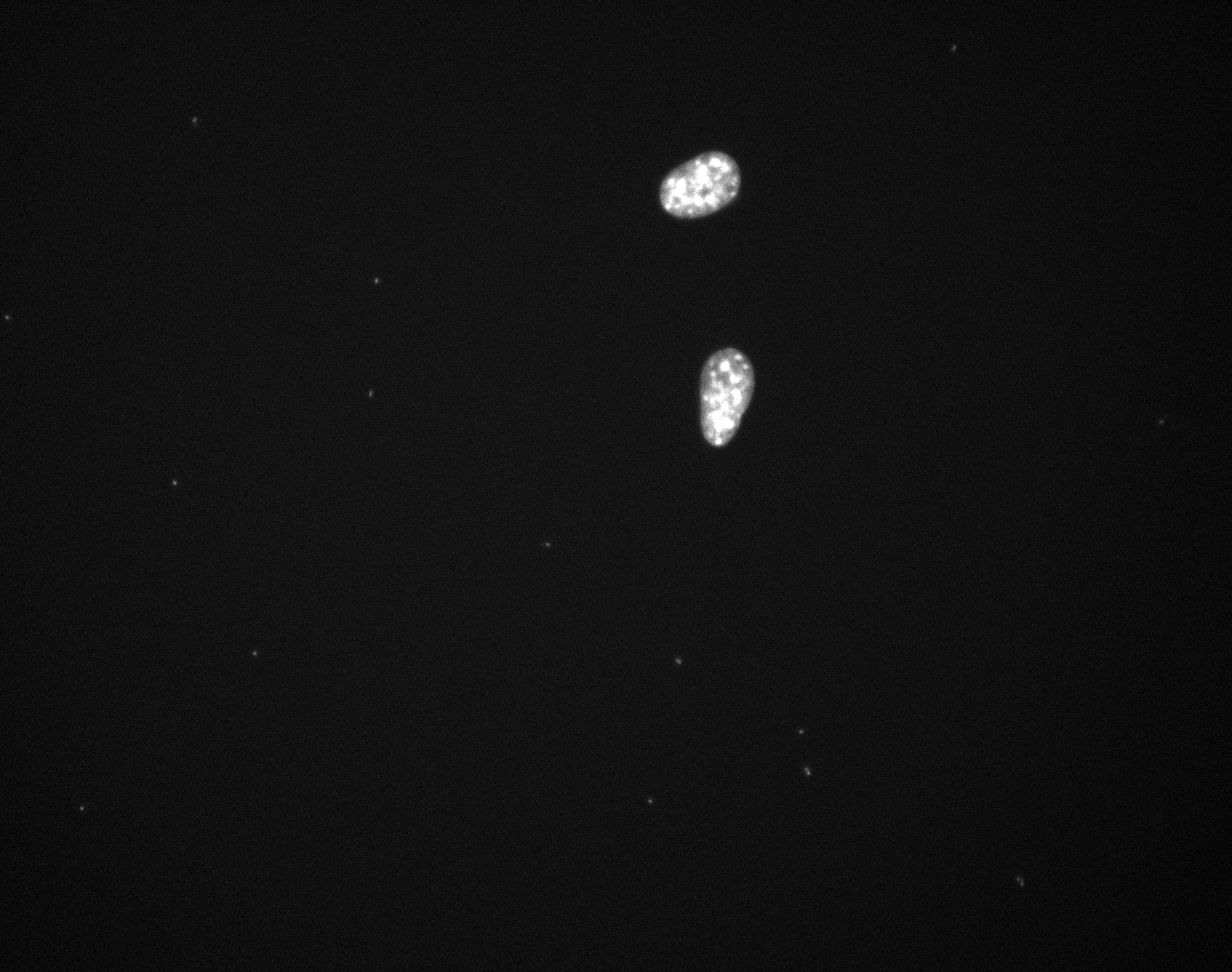

### E_Plate_R_p00_0_A01f54d1.TIF

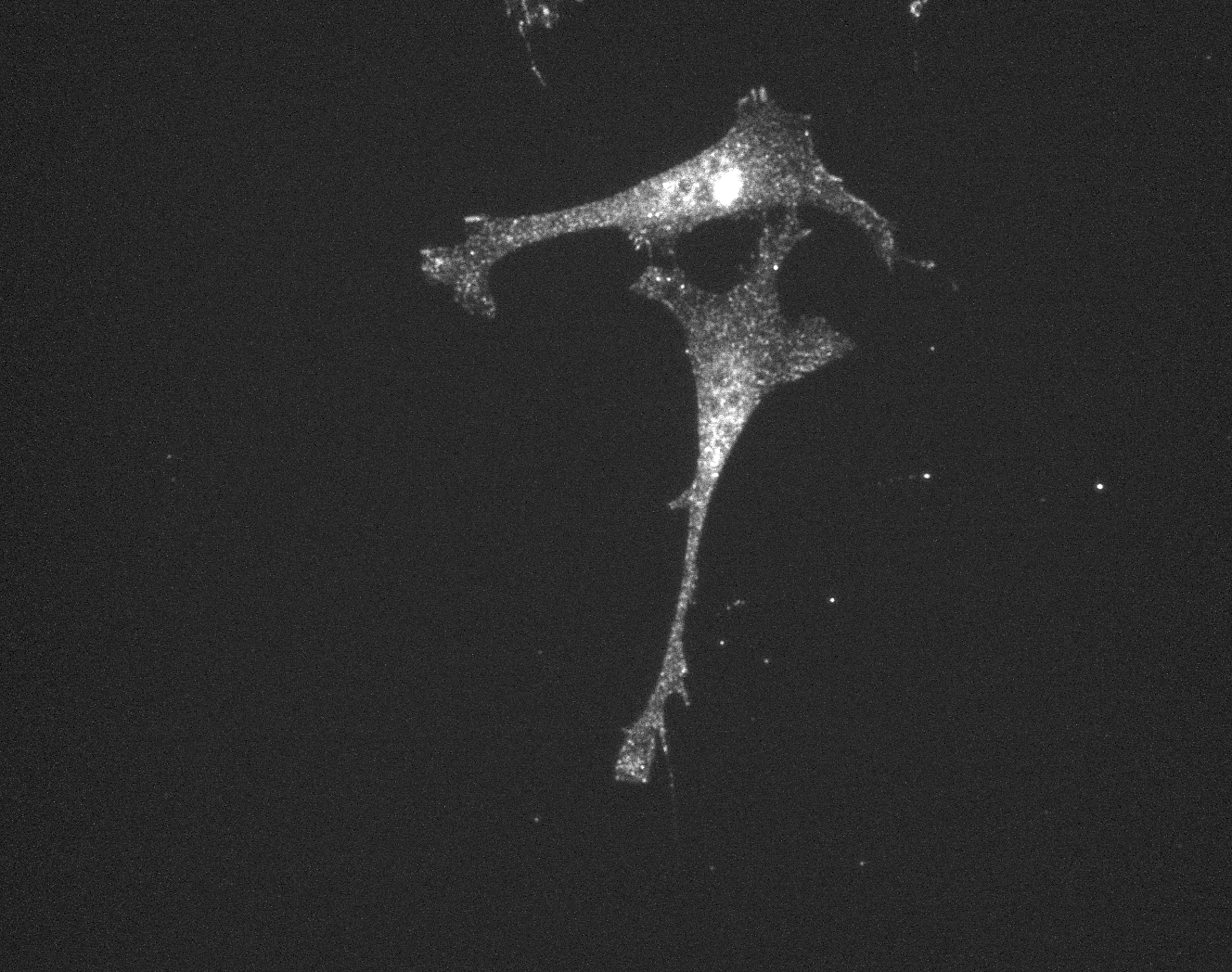

### E_Plate_R_p00_0_A01f54d2.TIF

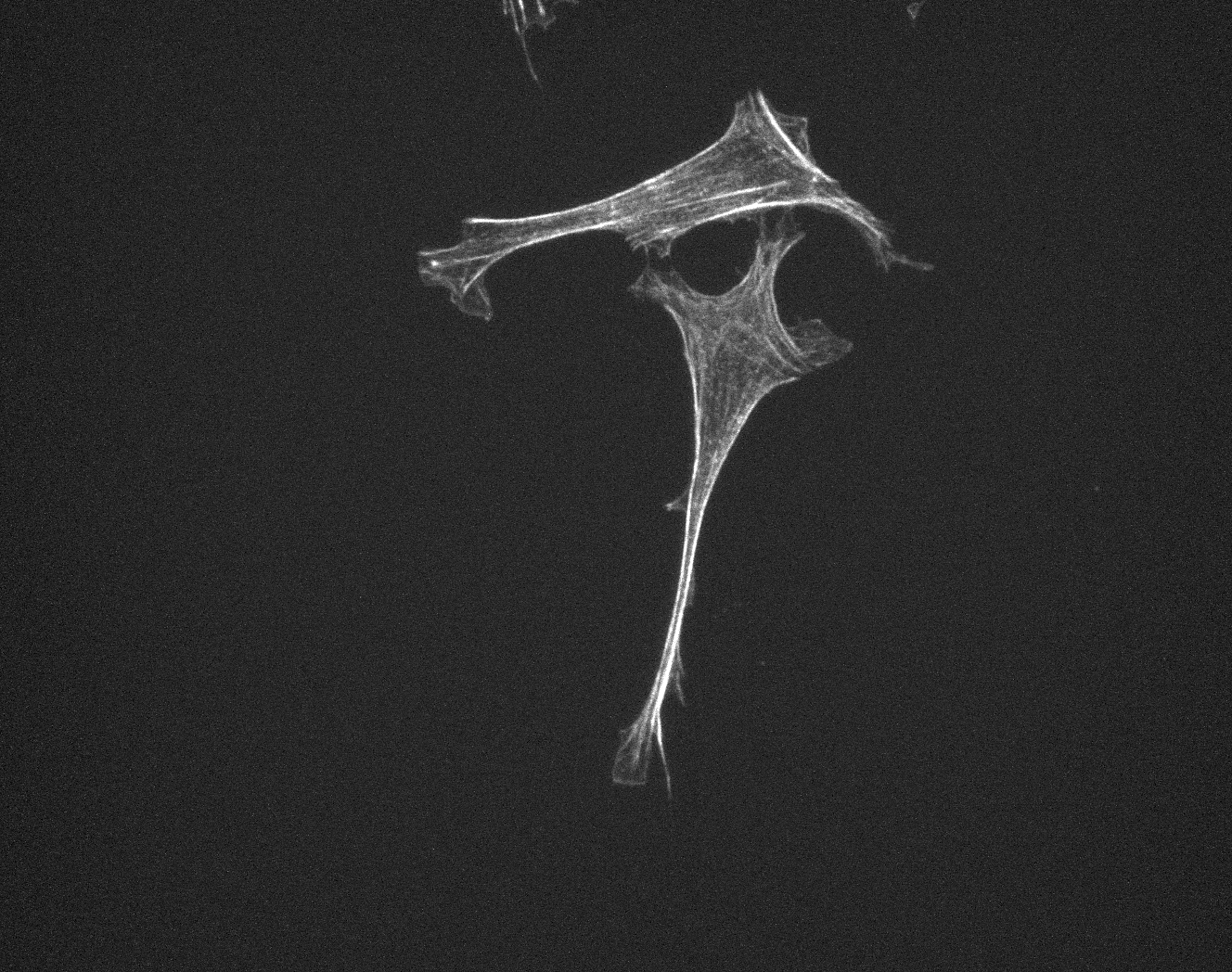

### E_Plate_R_p00_0_A01f54d2_CytoplasmOutline.tiff

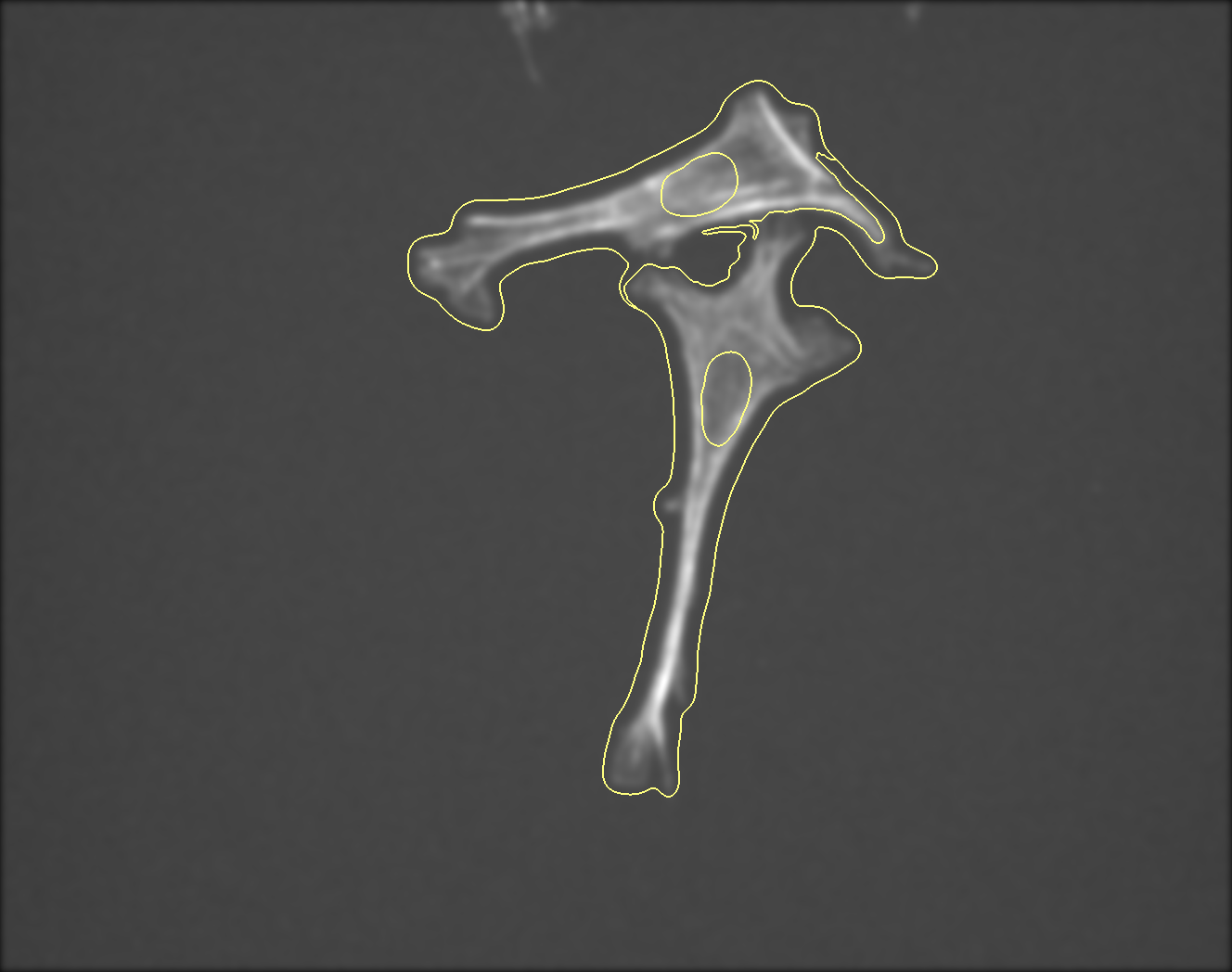

### E_Plate_R_p00_0_A01f54d3.TIF

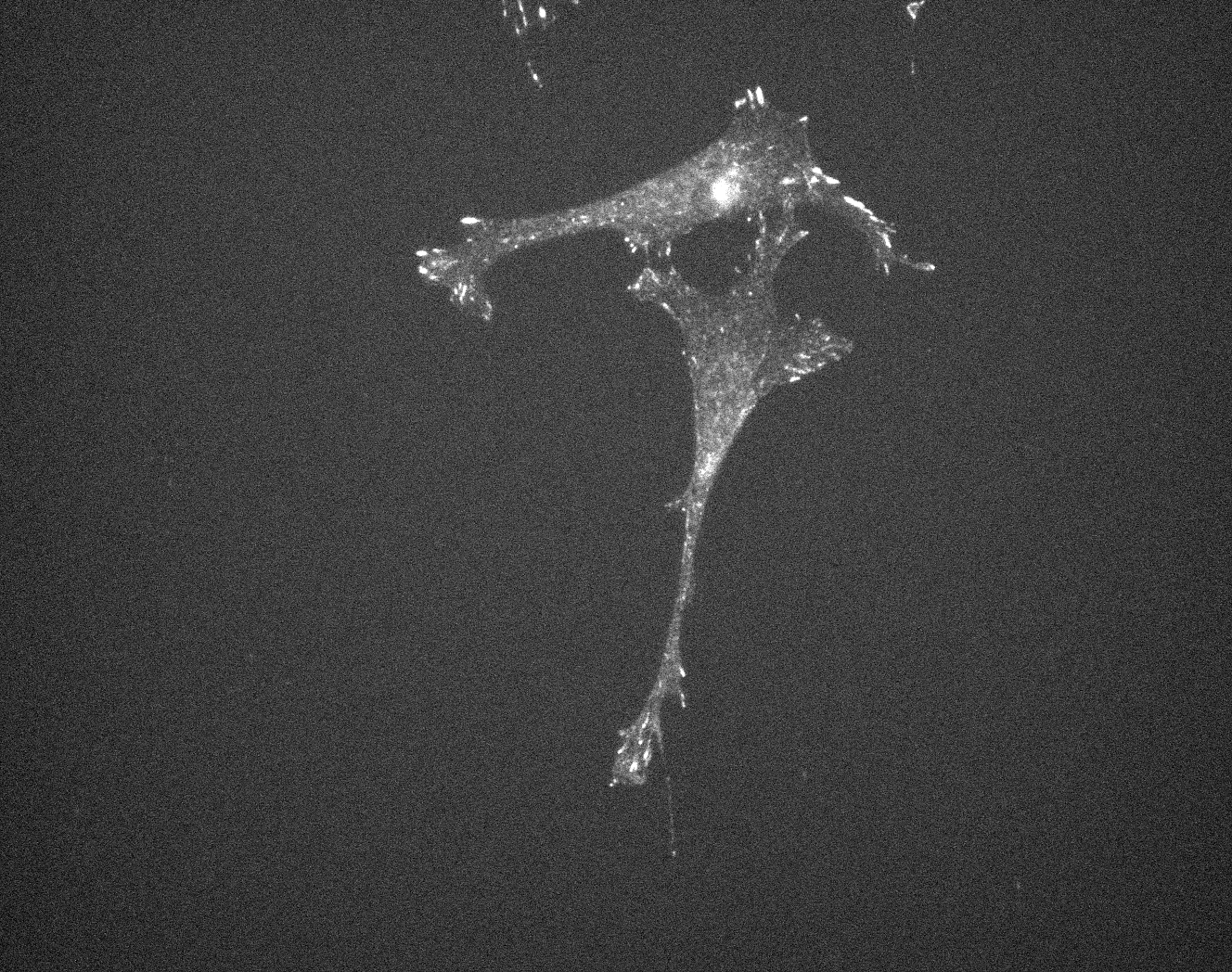

### F_Plate_R_p00_0_A01f25d0.TIF

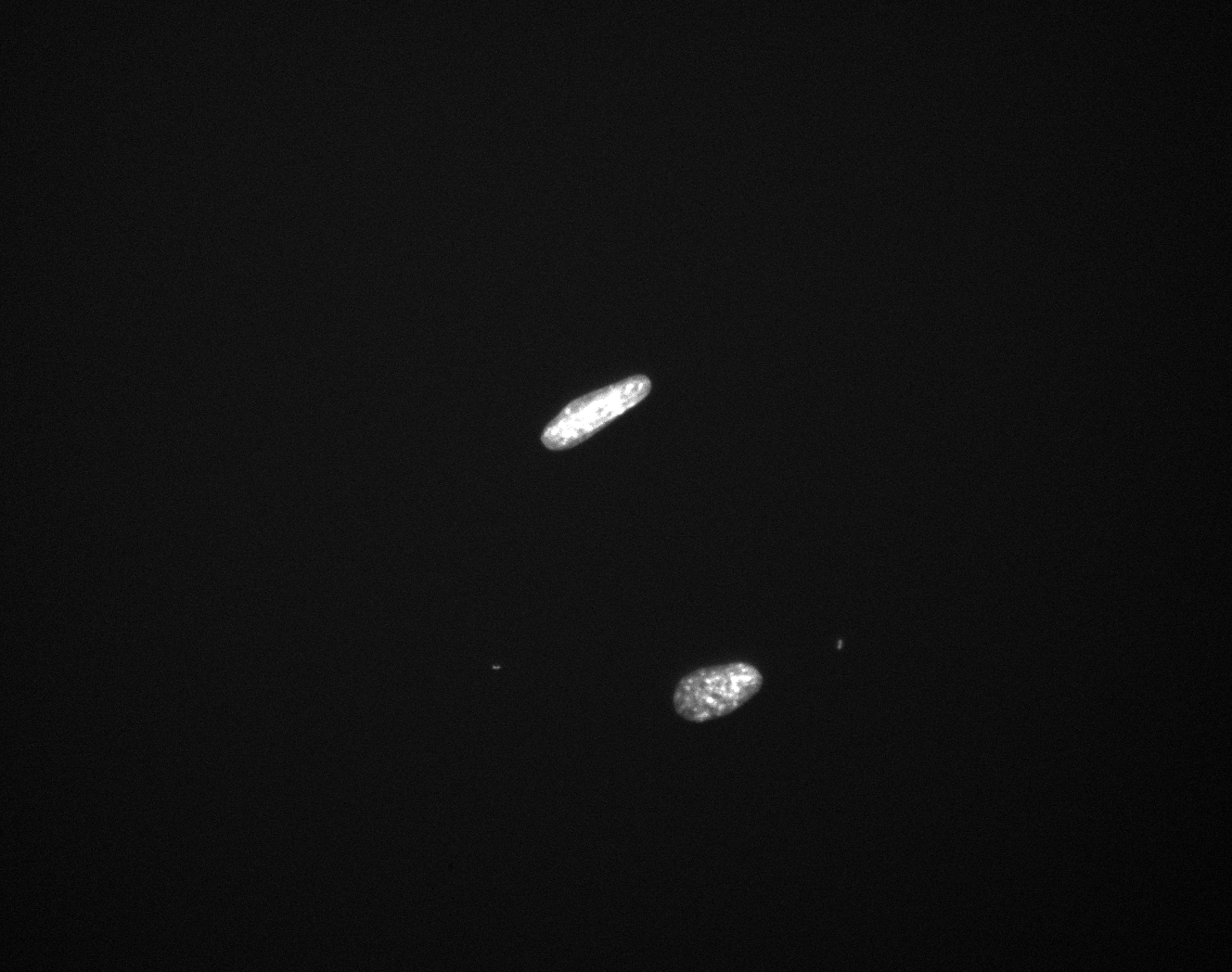

### F_Plate_R_p00_0_A01f25d1.TIF

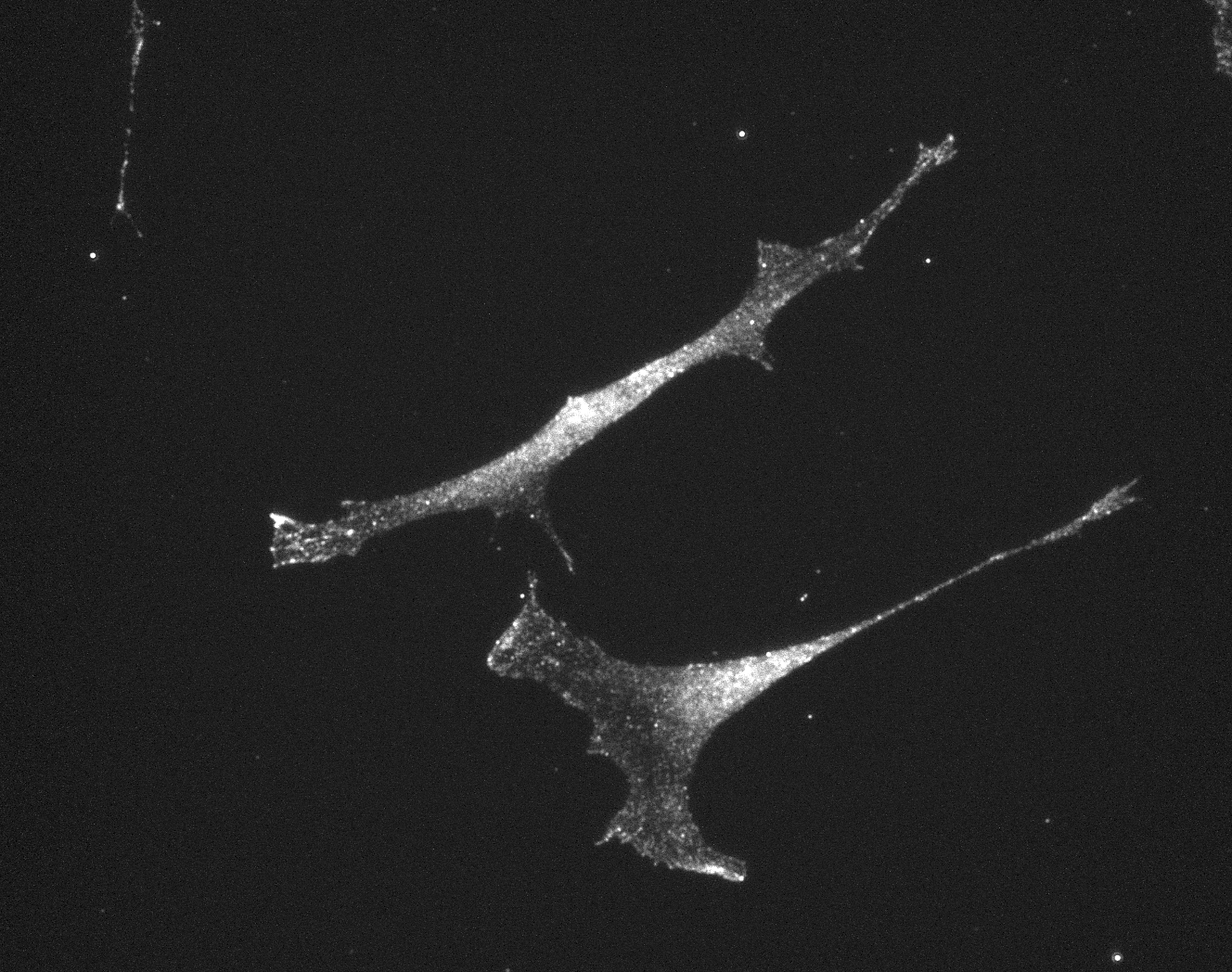

### F_Plate_R_p00_0_A01f25d2.TIF

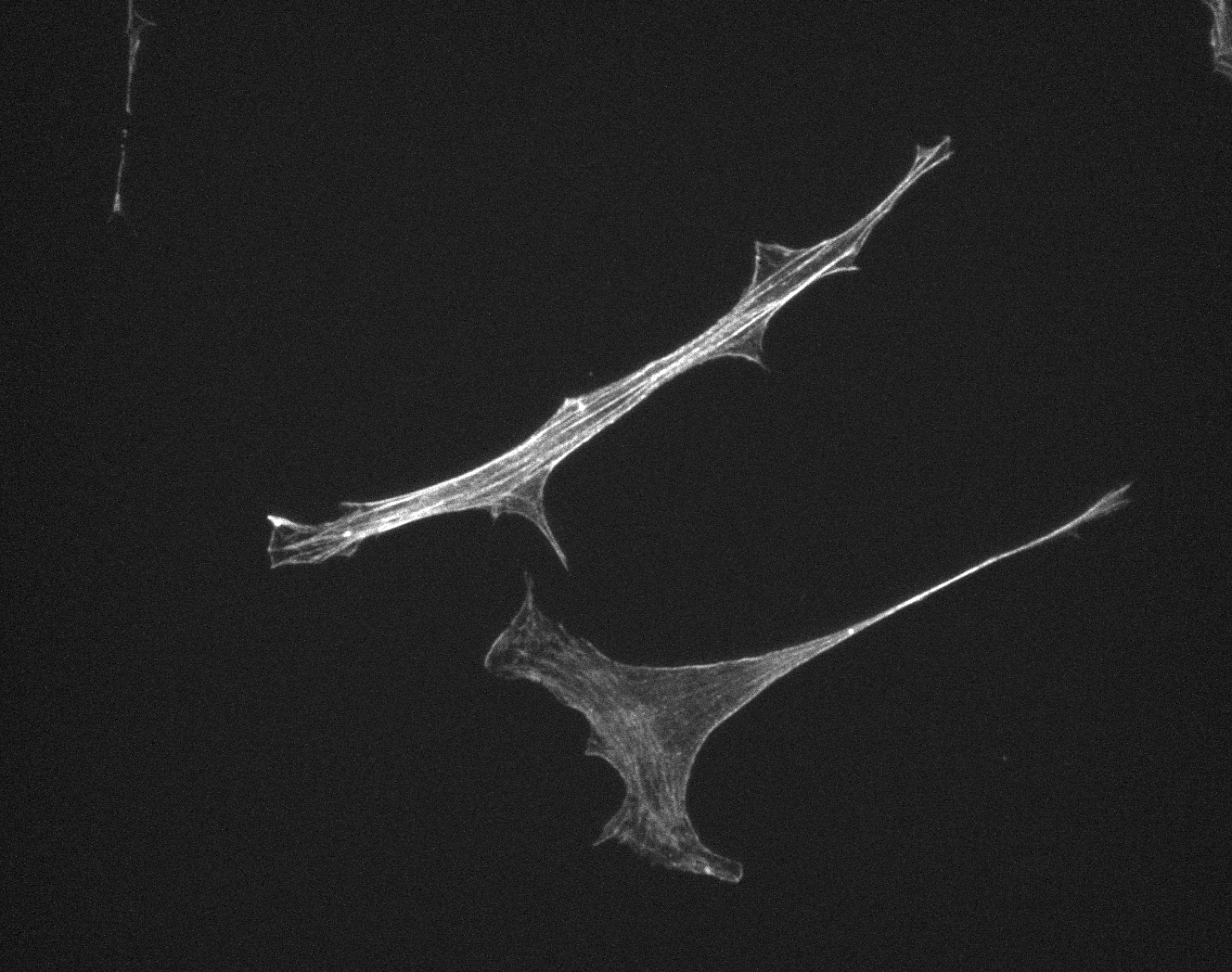

### F_Plate_R_p00_0_A01f25d3.TIF

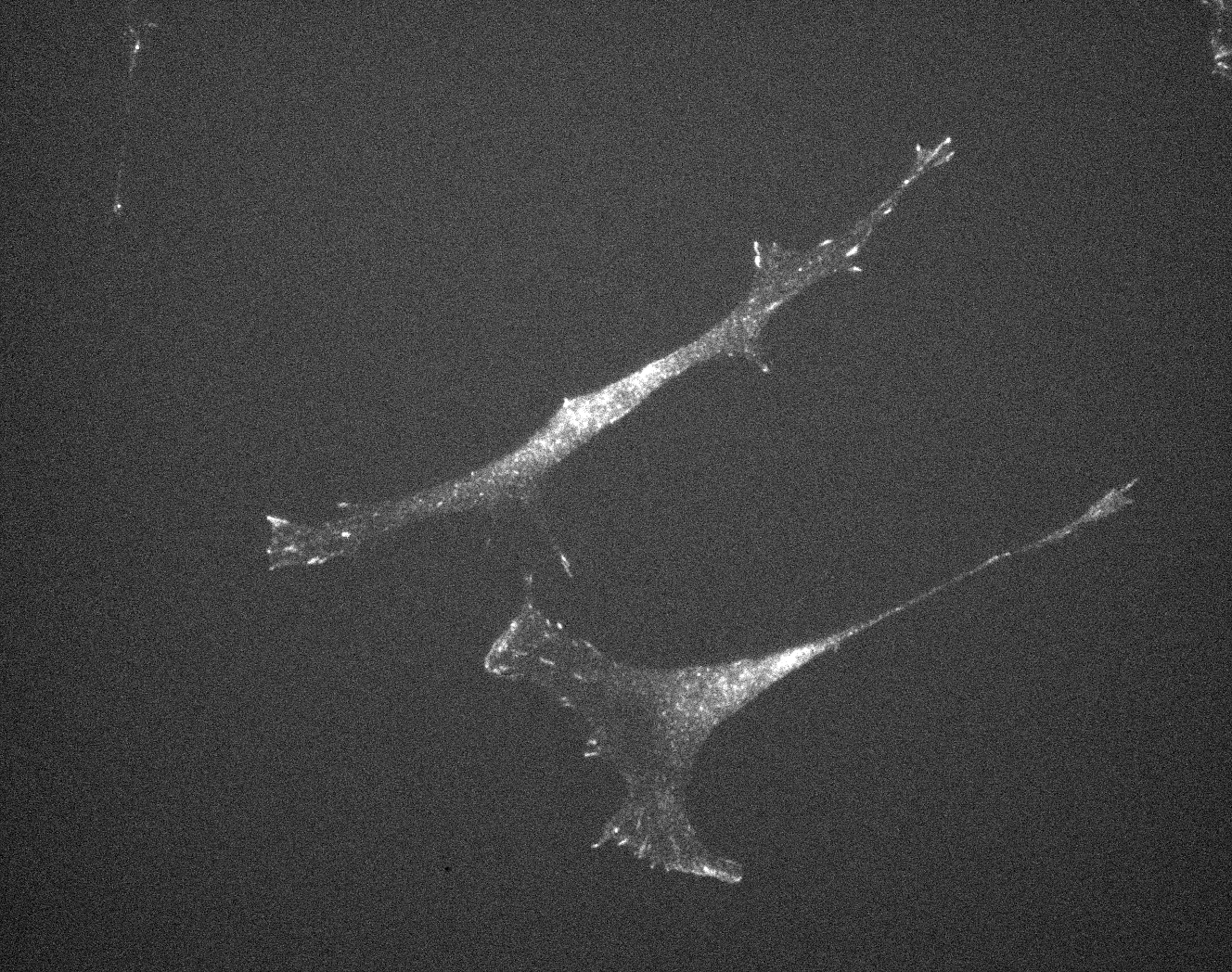

### F_Plate_R_p00_0_A01f37d3_FAOutline.tiff

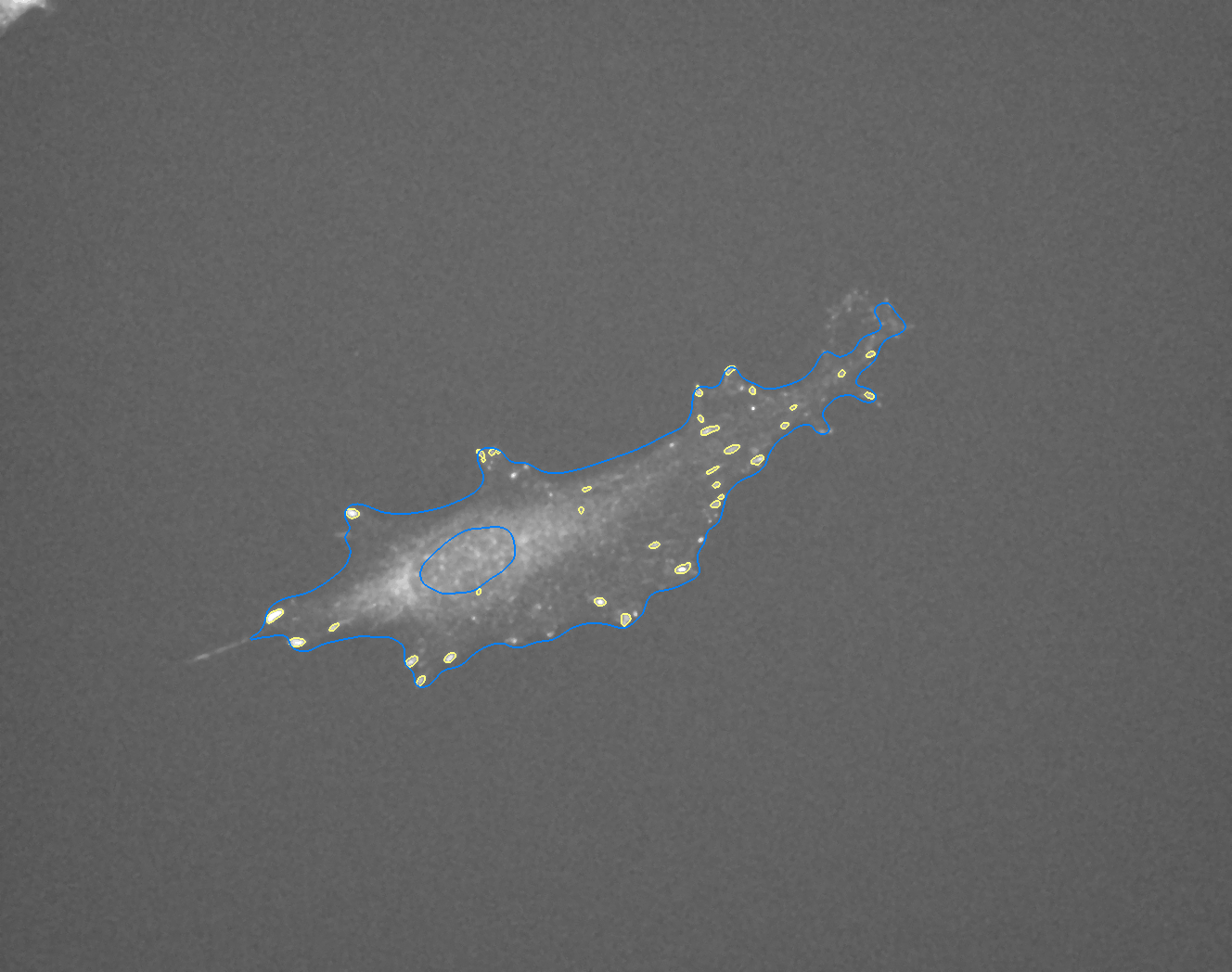

### F_Plate_R_p00_0_A01f47d0.TIF

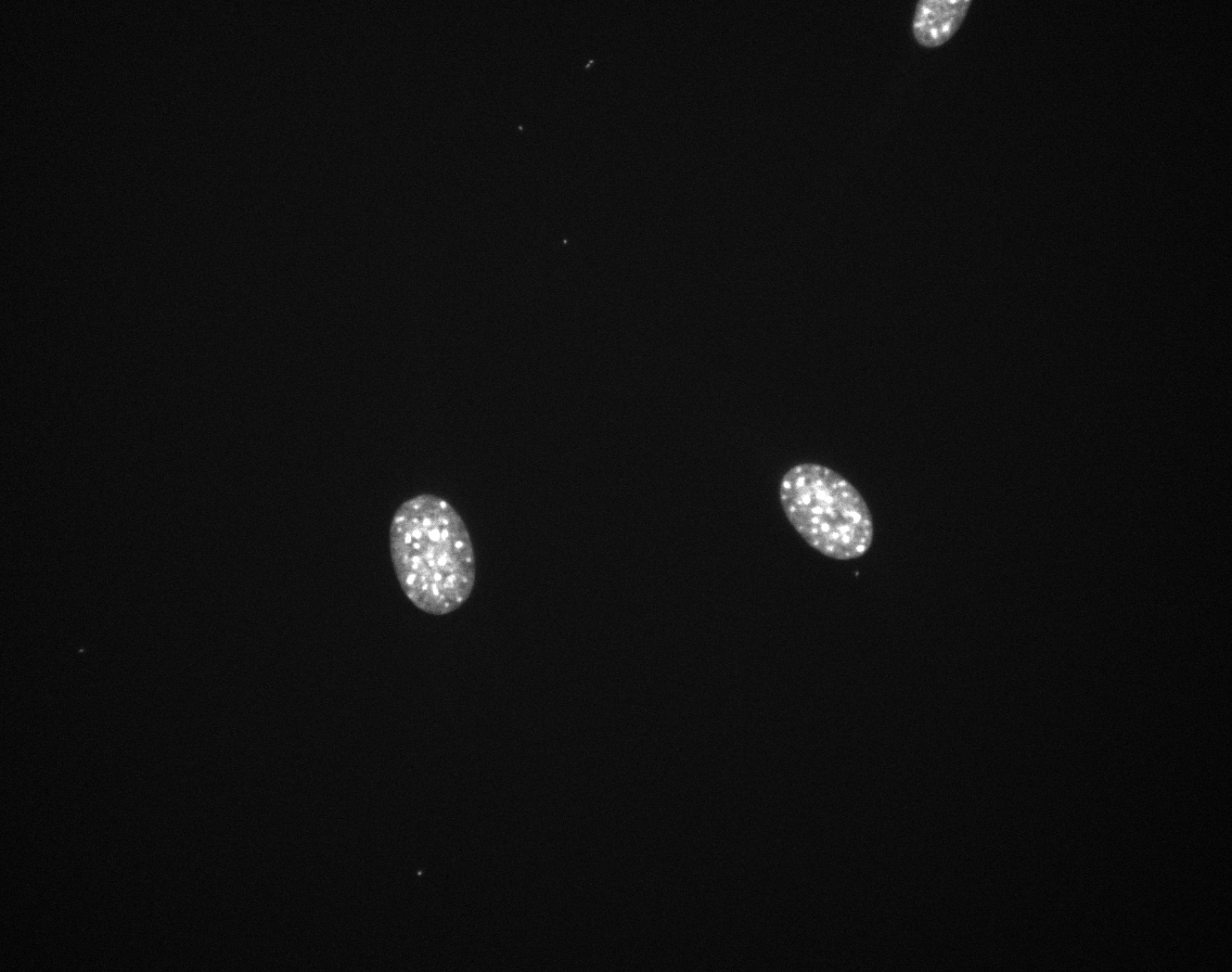

### F_Plate_R_p00_0_A01f47d1.TIF

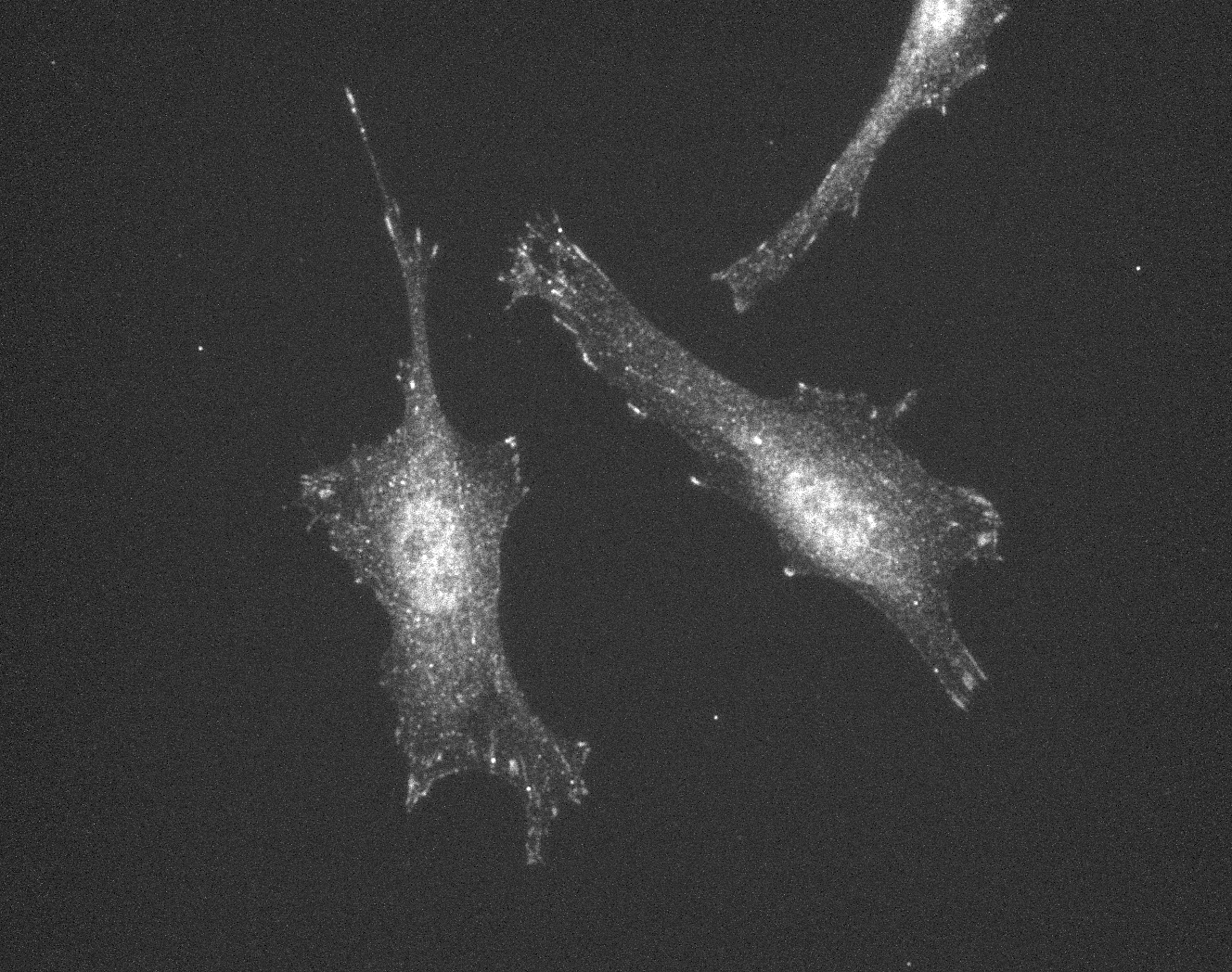

### F_Plate_R_p00_0_A01f47d2.TIF

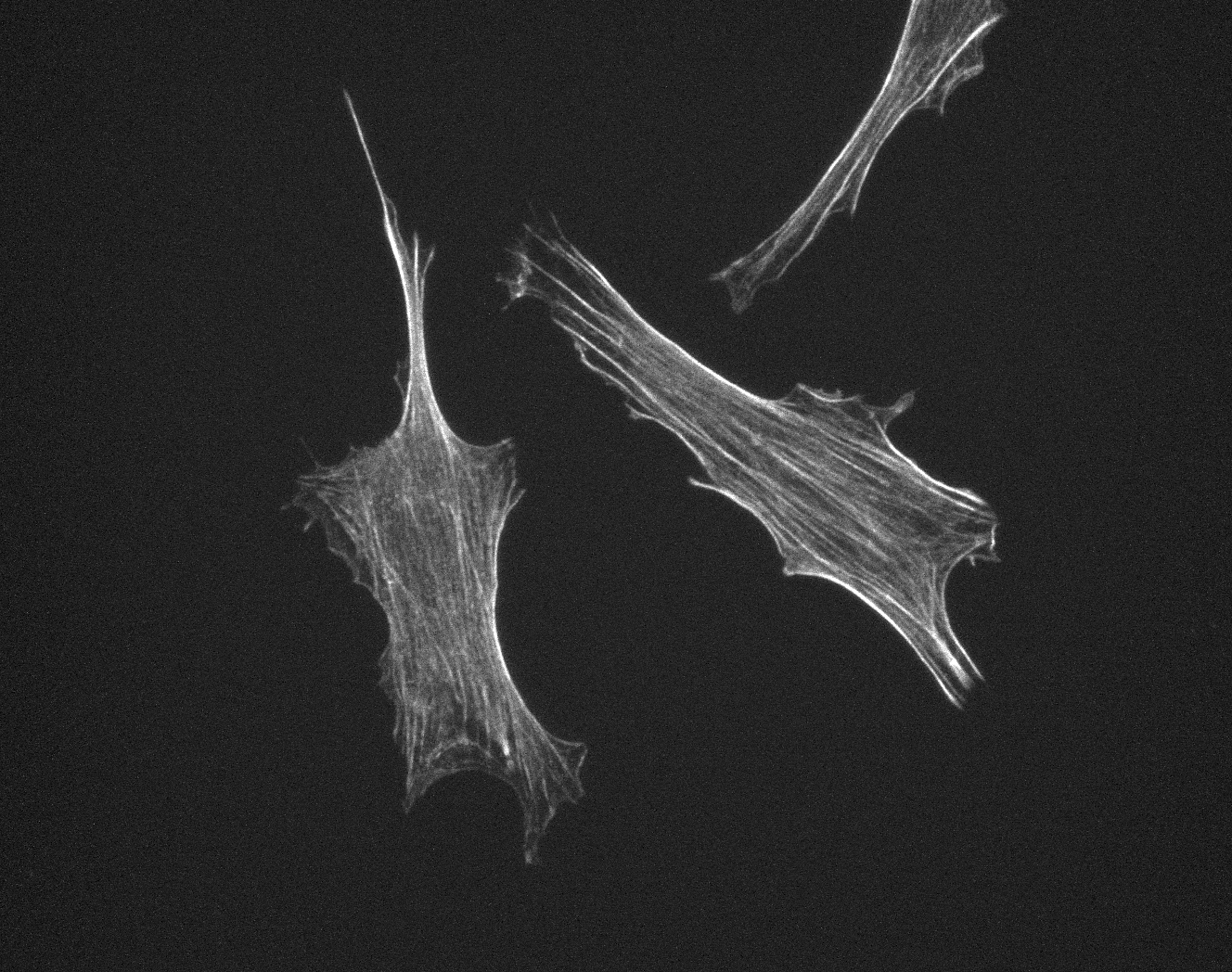

### F_Plate_R_p00_0_A01f47d2_CytoplasmOutline.tiff

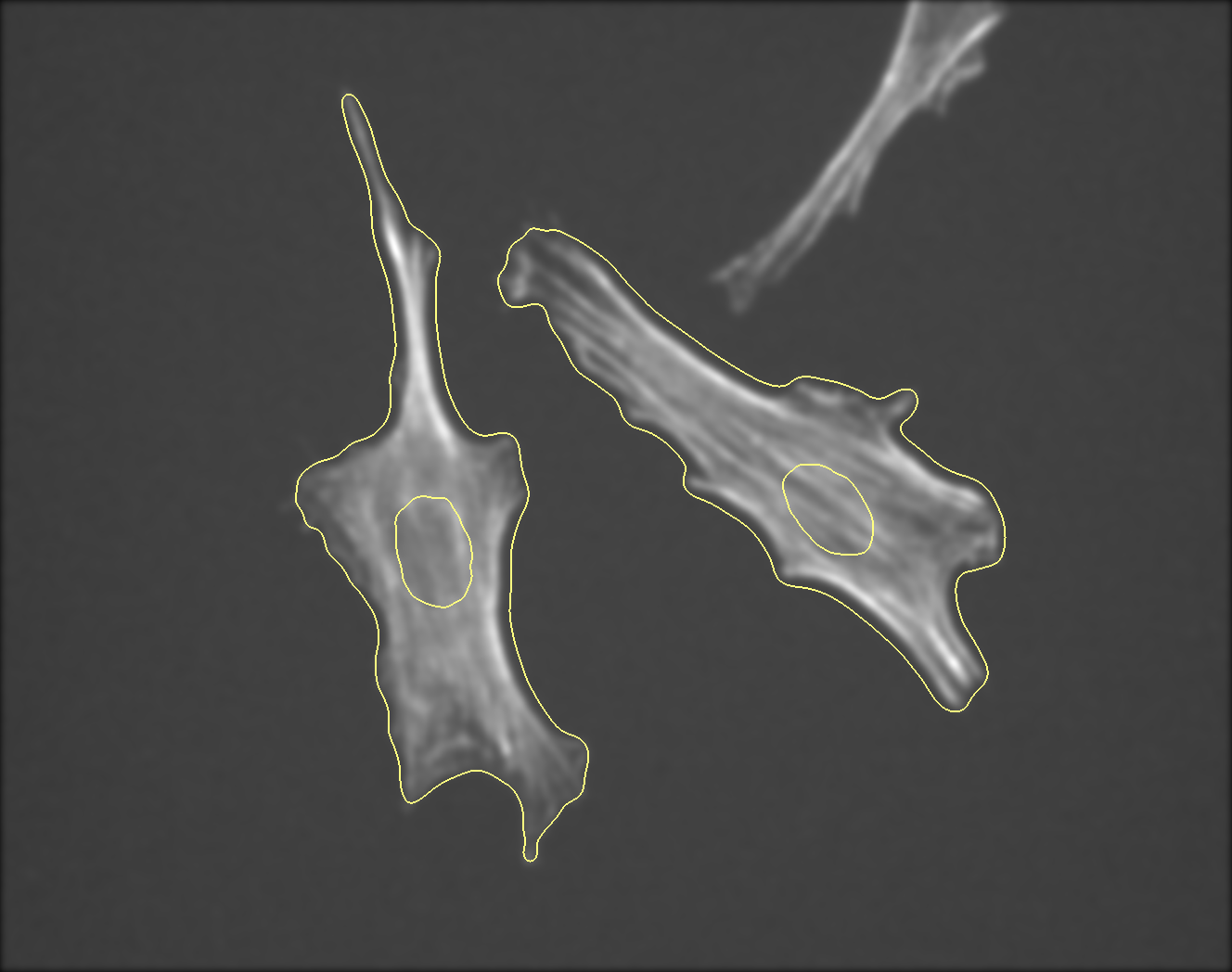

### F_Plate_R_p00_0_A01f47d3.TIF

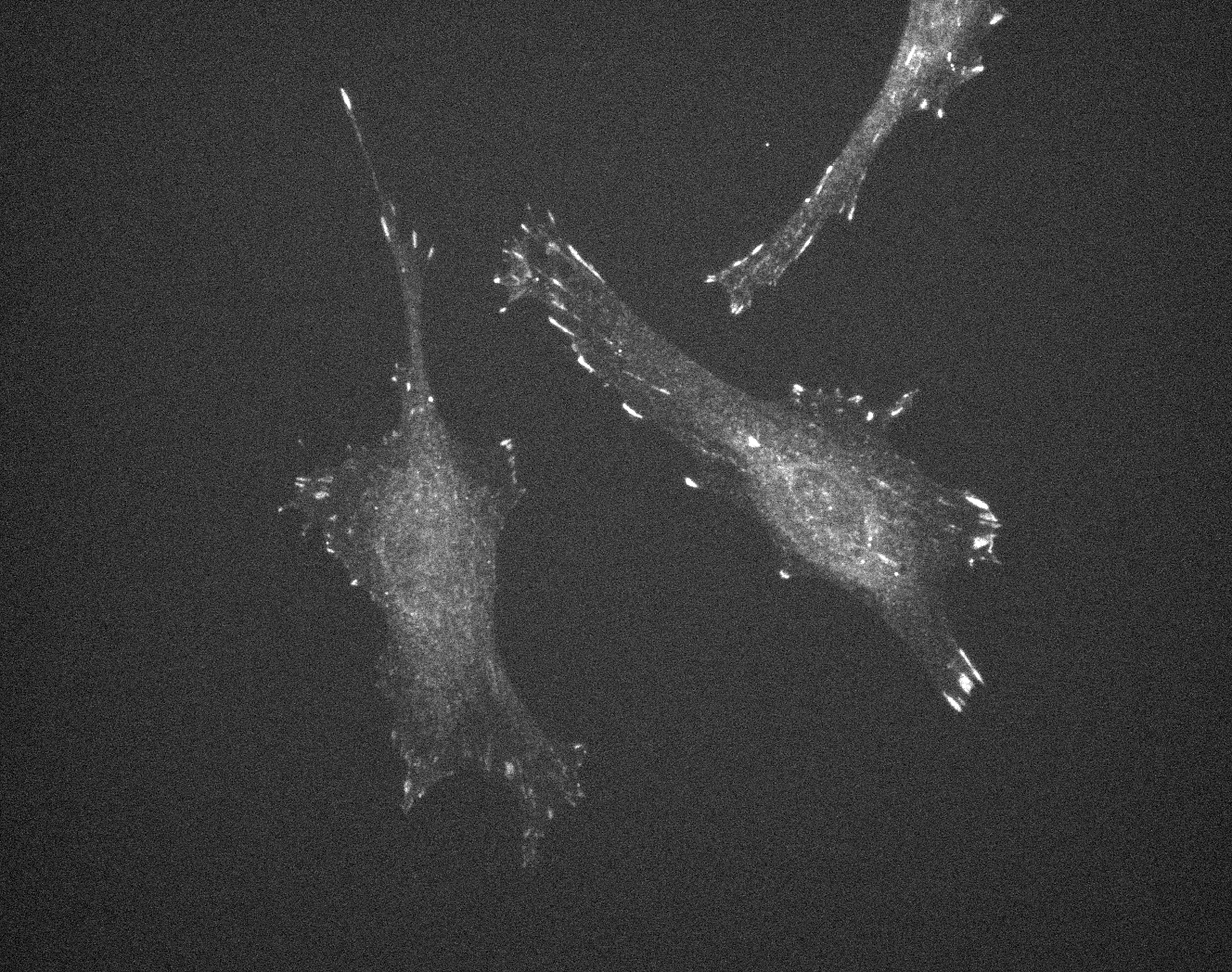

### G_Plate_R_p00_0_A01f02d0.TIF

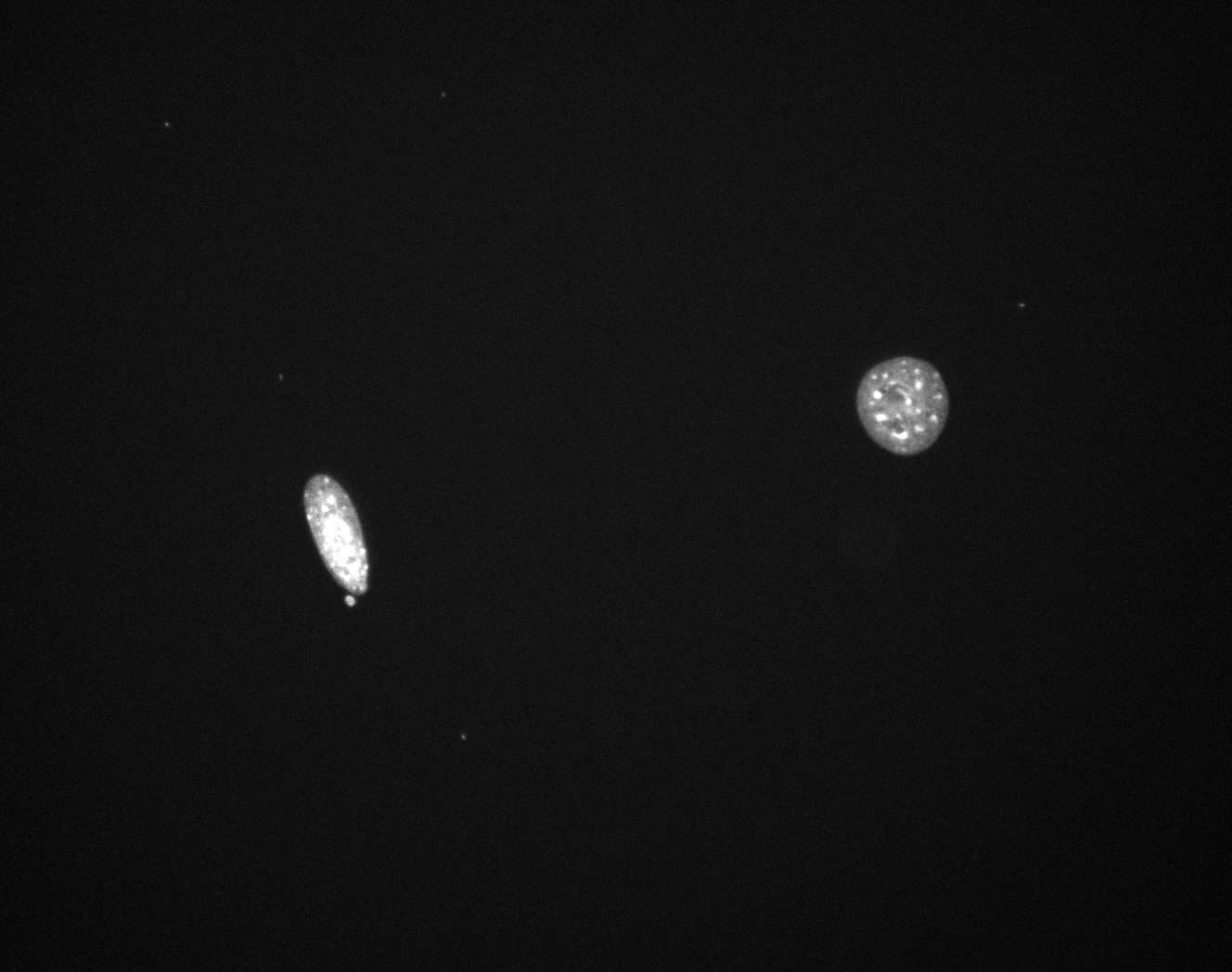

### G_Plate_R_p00_0_A01f02d1.TIF

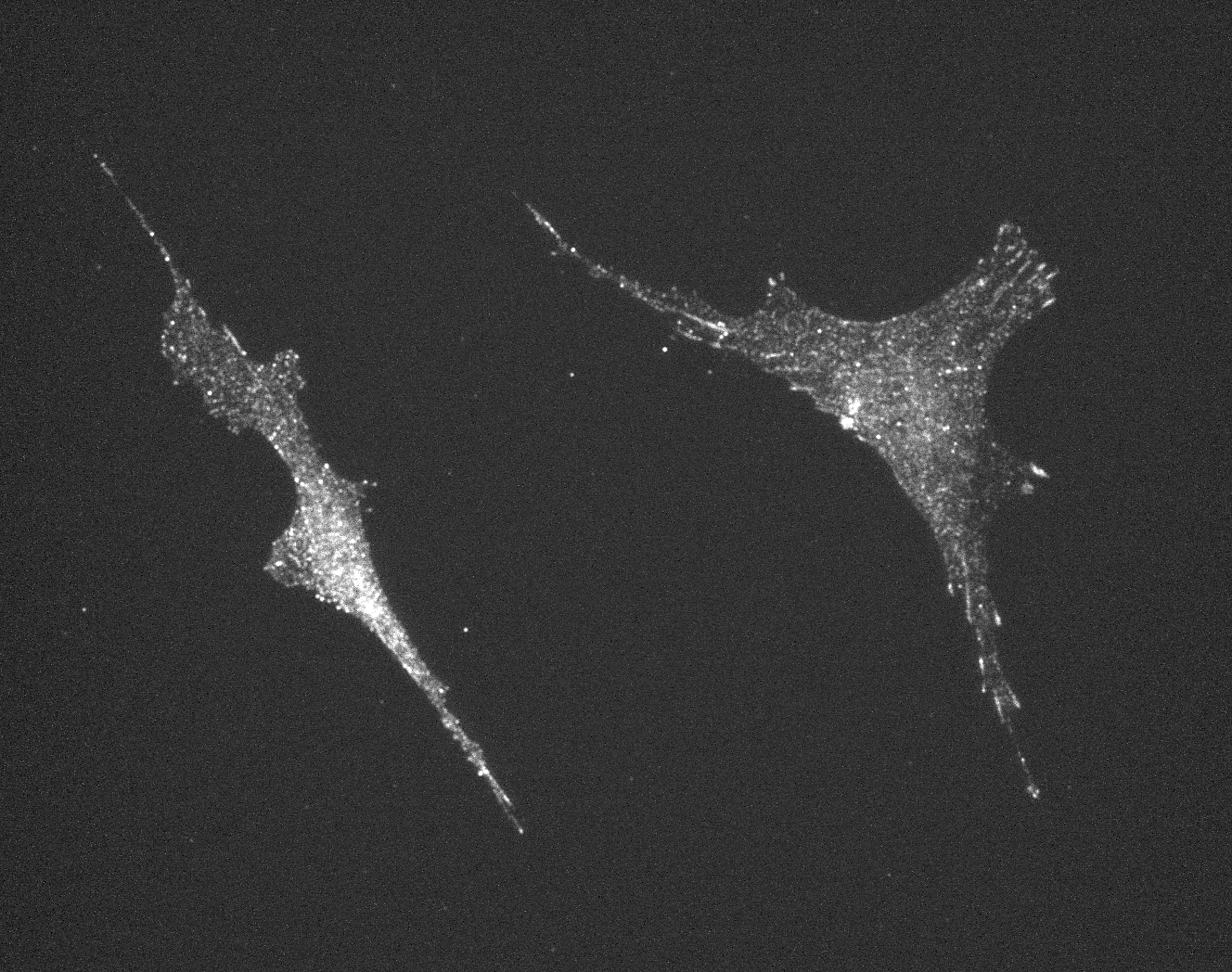

### G_Plate_R_p00_0_A01f02d2.TIF

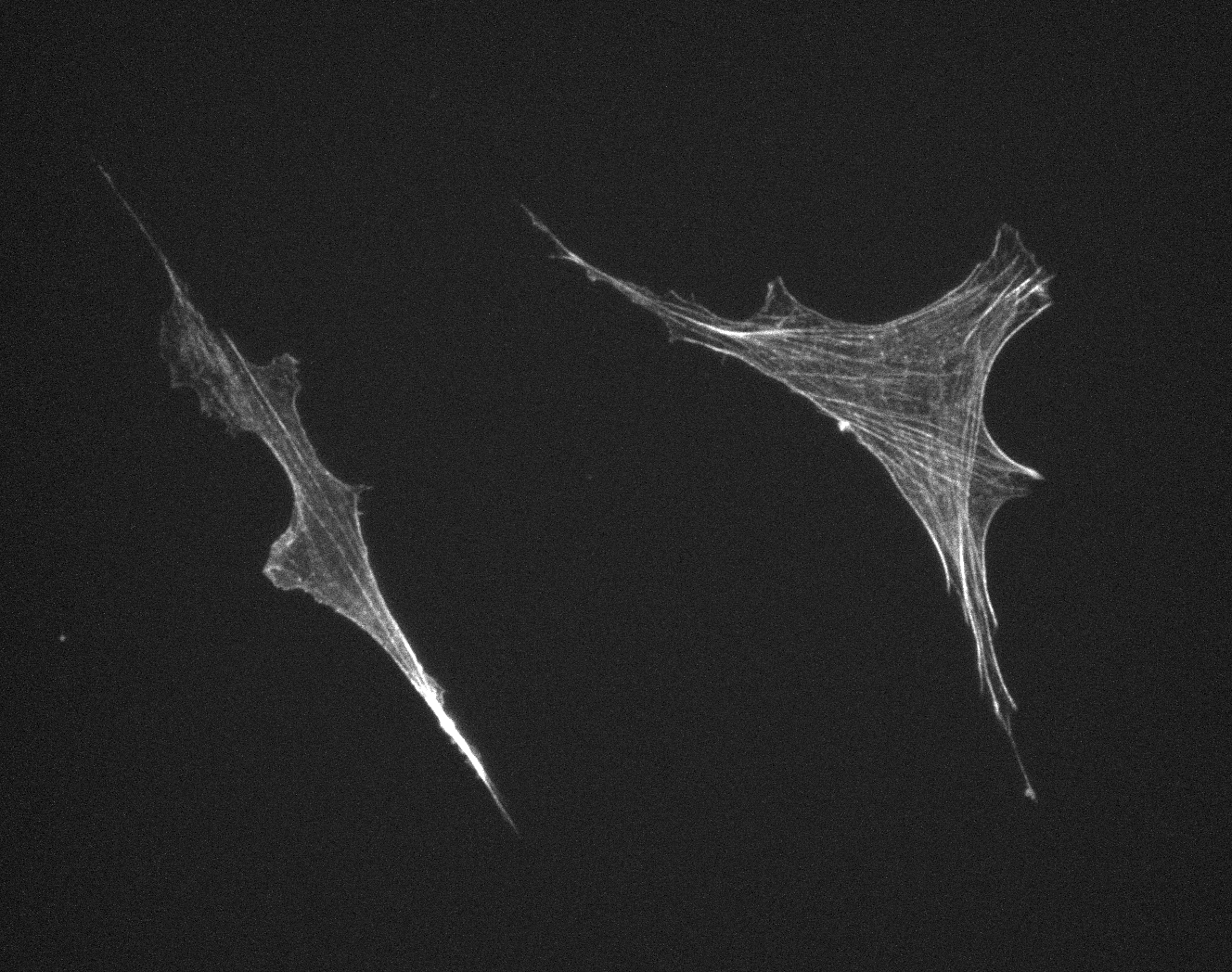

### G_Plate_R_p00_0_A01f02d3.TIF

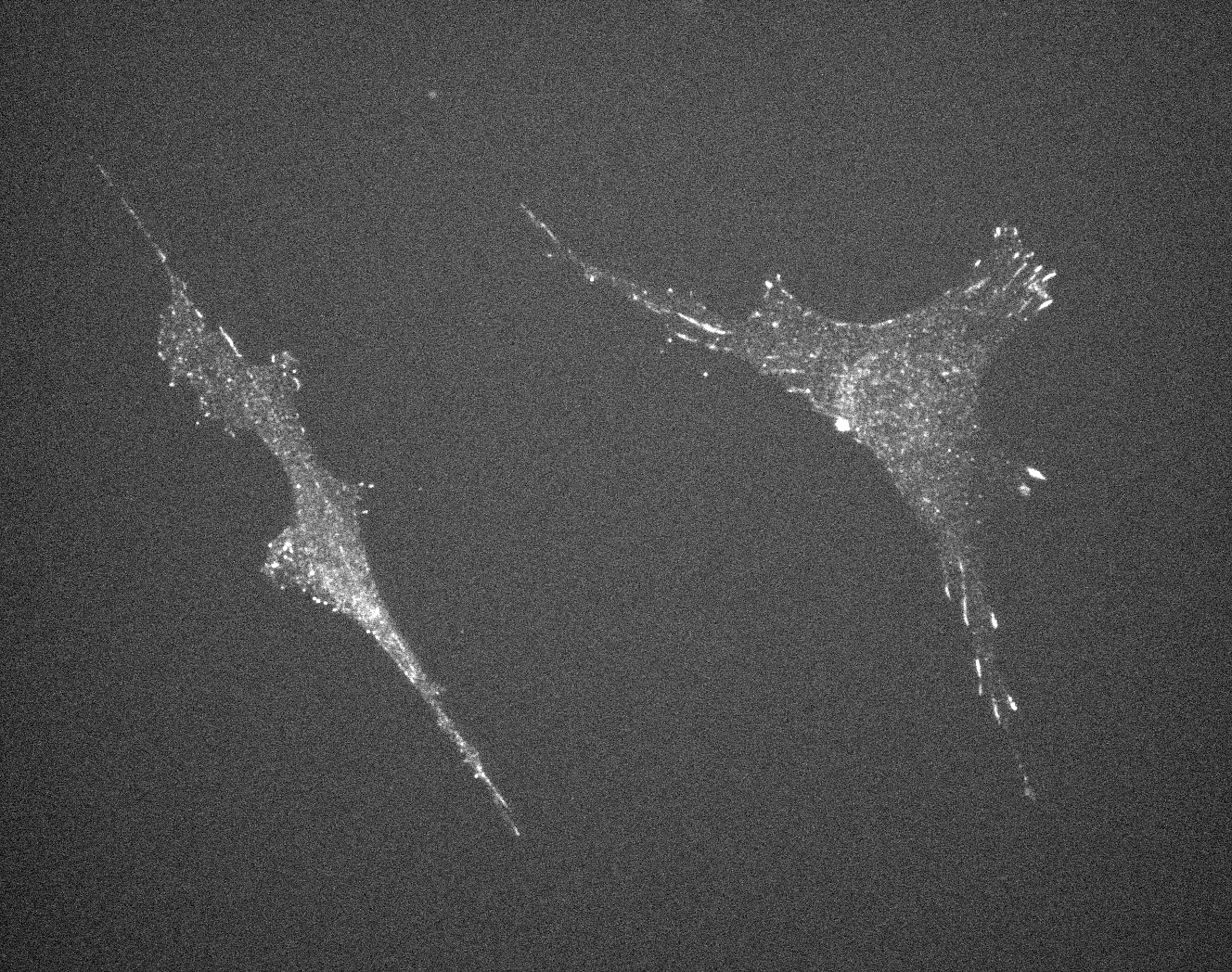

### G_Plate_R_p00_0_A01f03d0.TIF

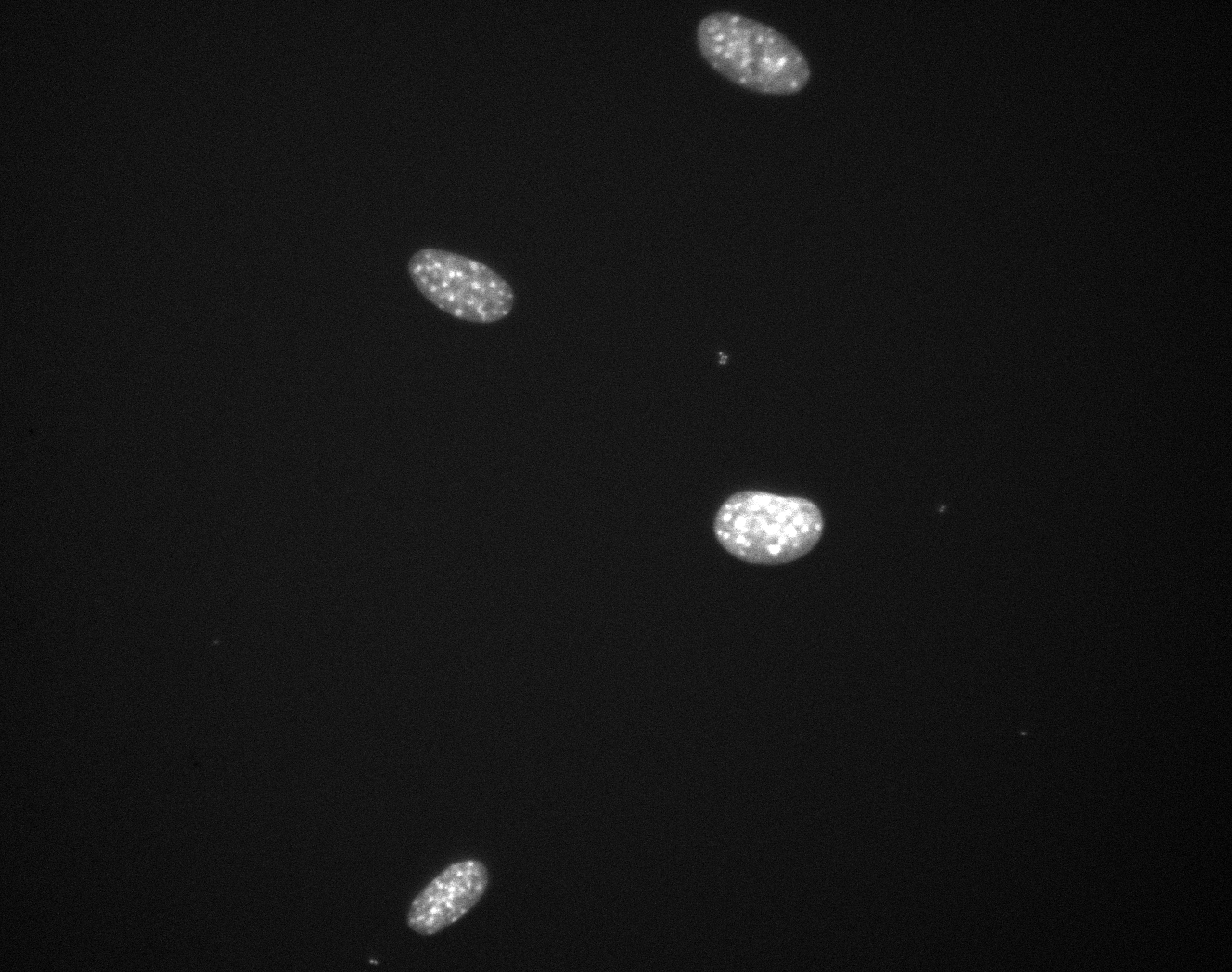

### G_Plate_R_p00_0_A01f03d1.TIF

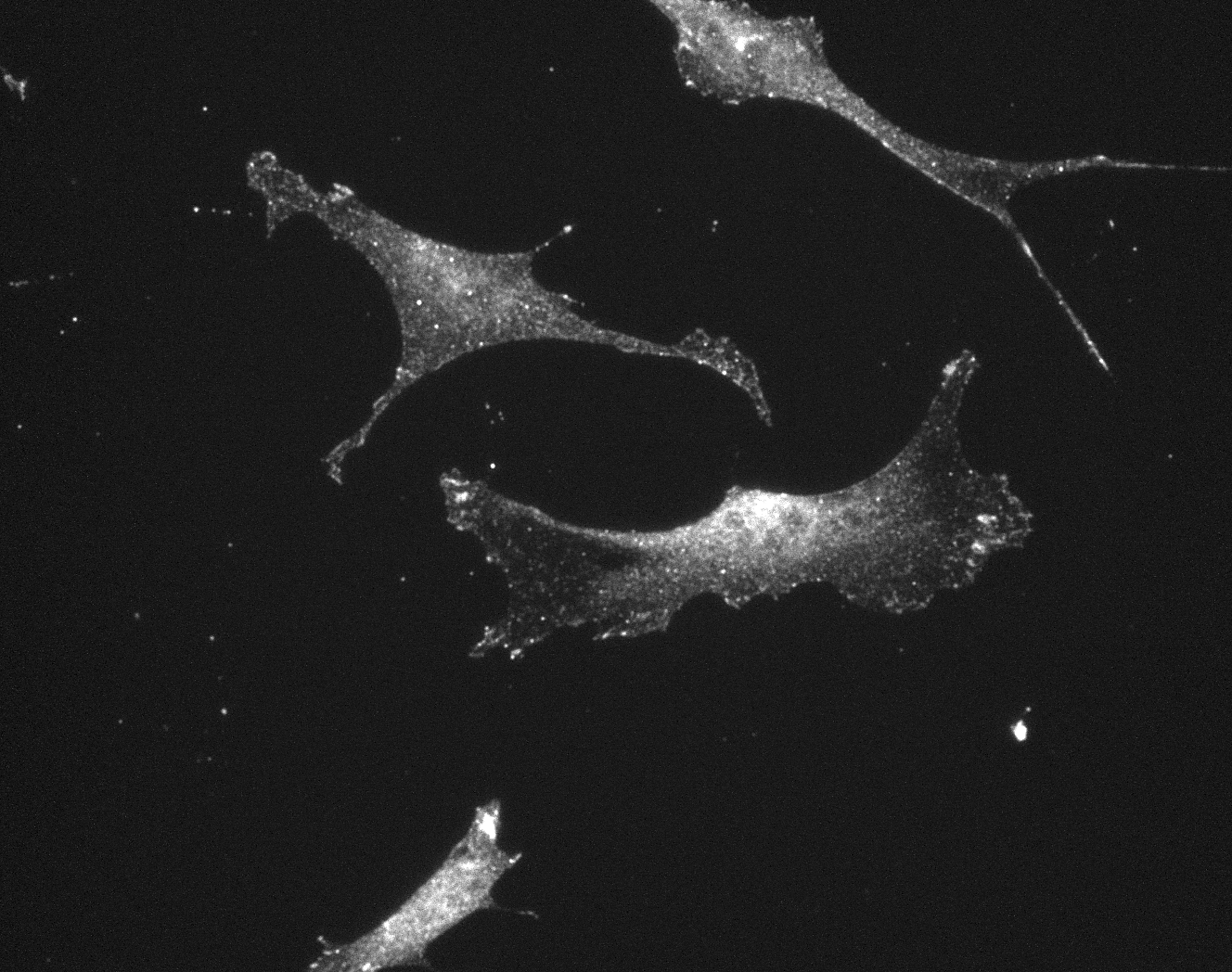

### G_Plate_R_p00_0_A01f03d2.TIF

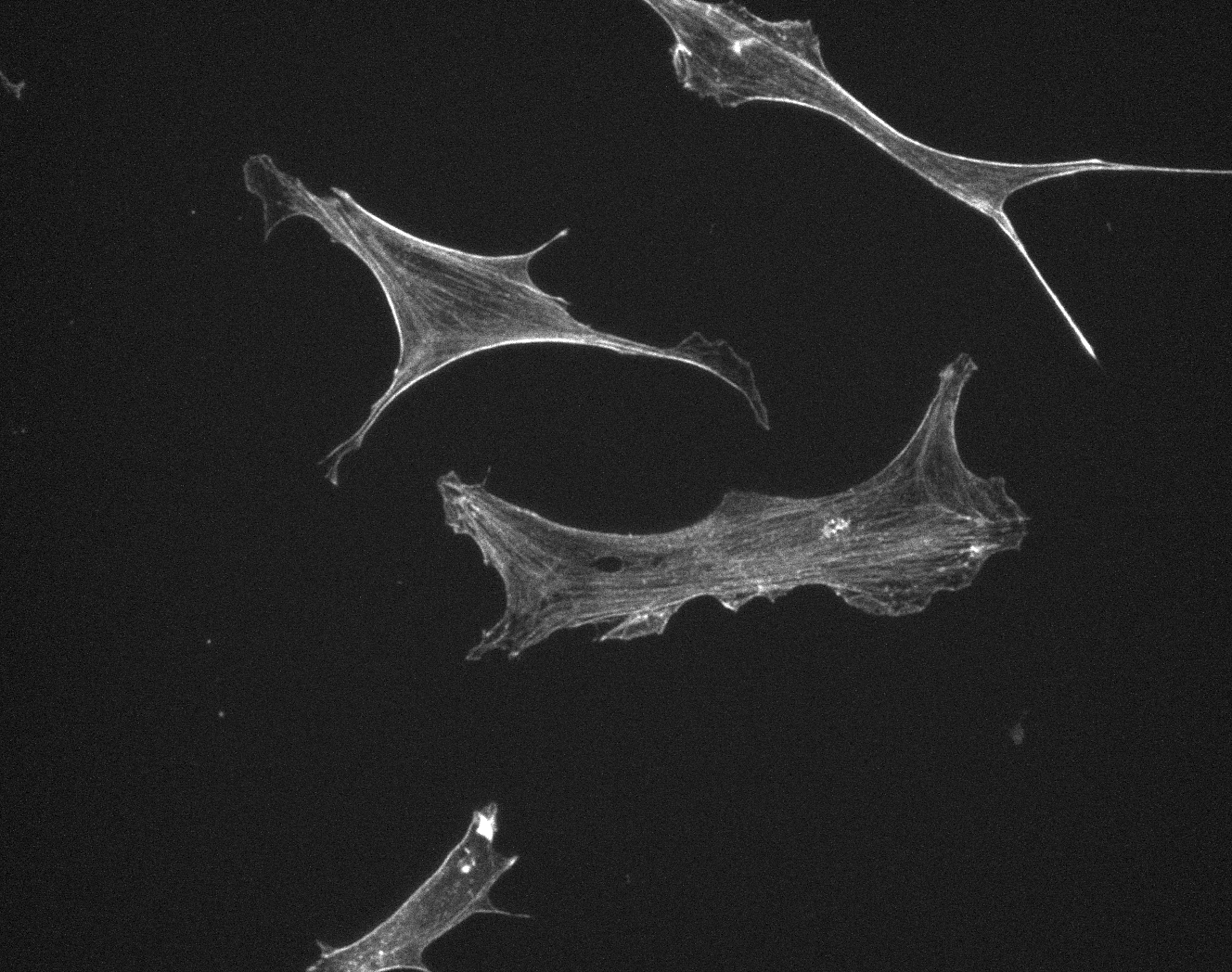

### G_Plate_R_p00_0_A01f03d2_CytoplasmOutline.tiff

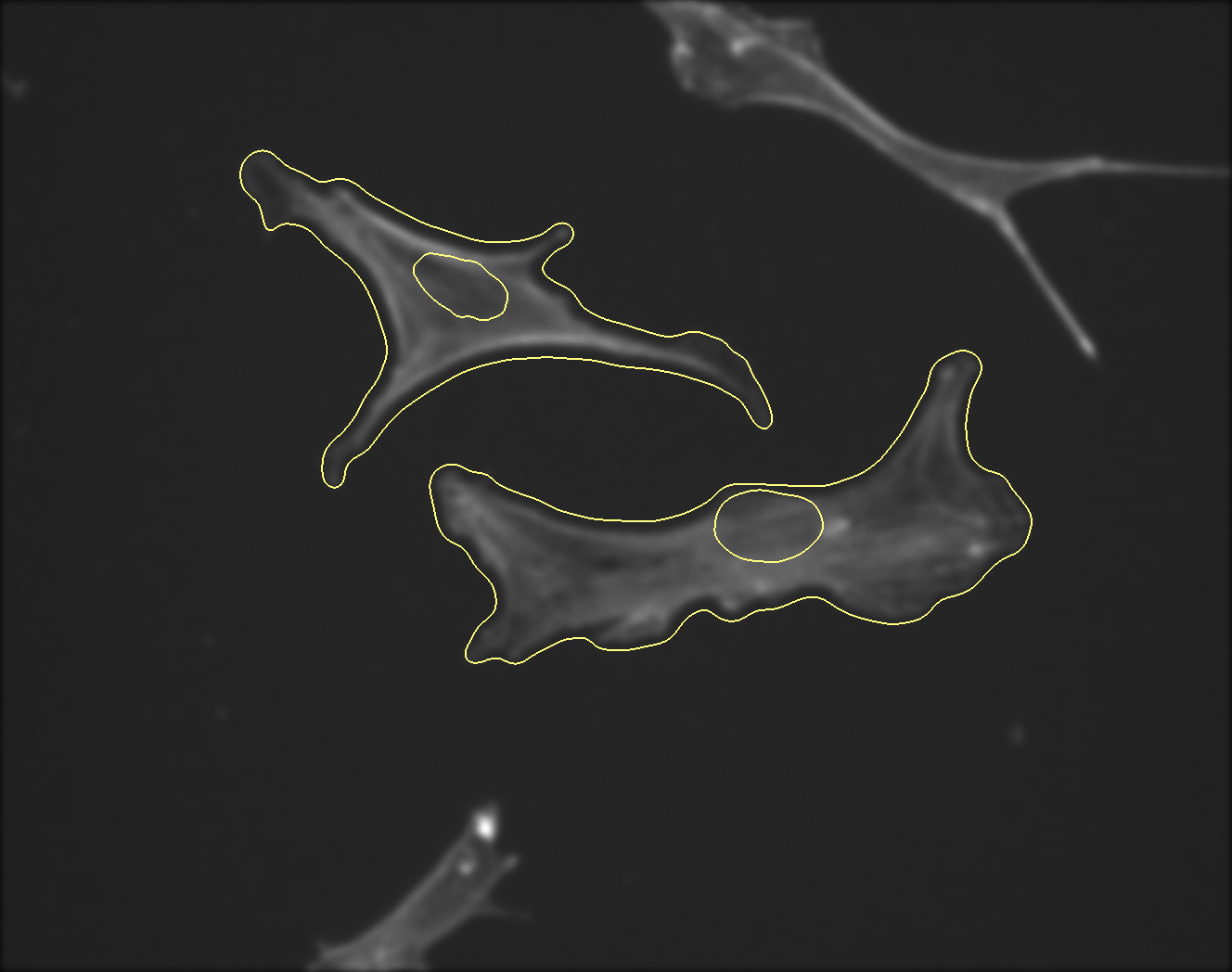

### G_Plate_R_p00_0_A01f03d3.TIF

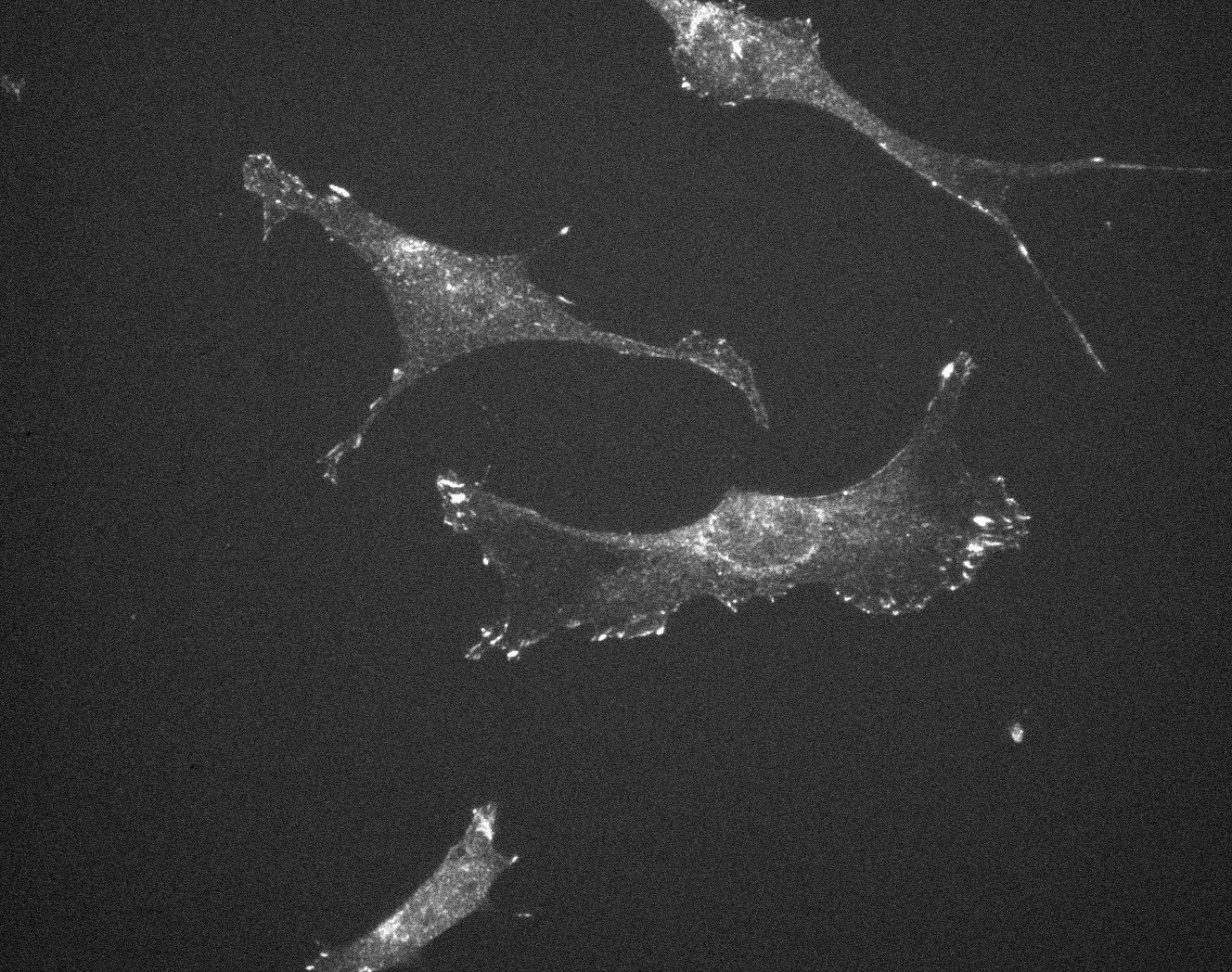

### G_Plate_R_p00_0_A01f26d3_FAOutline.tiff

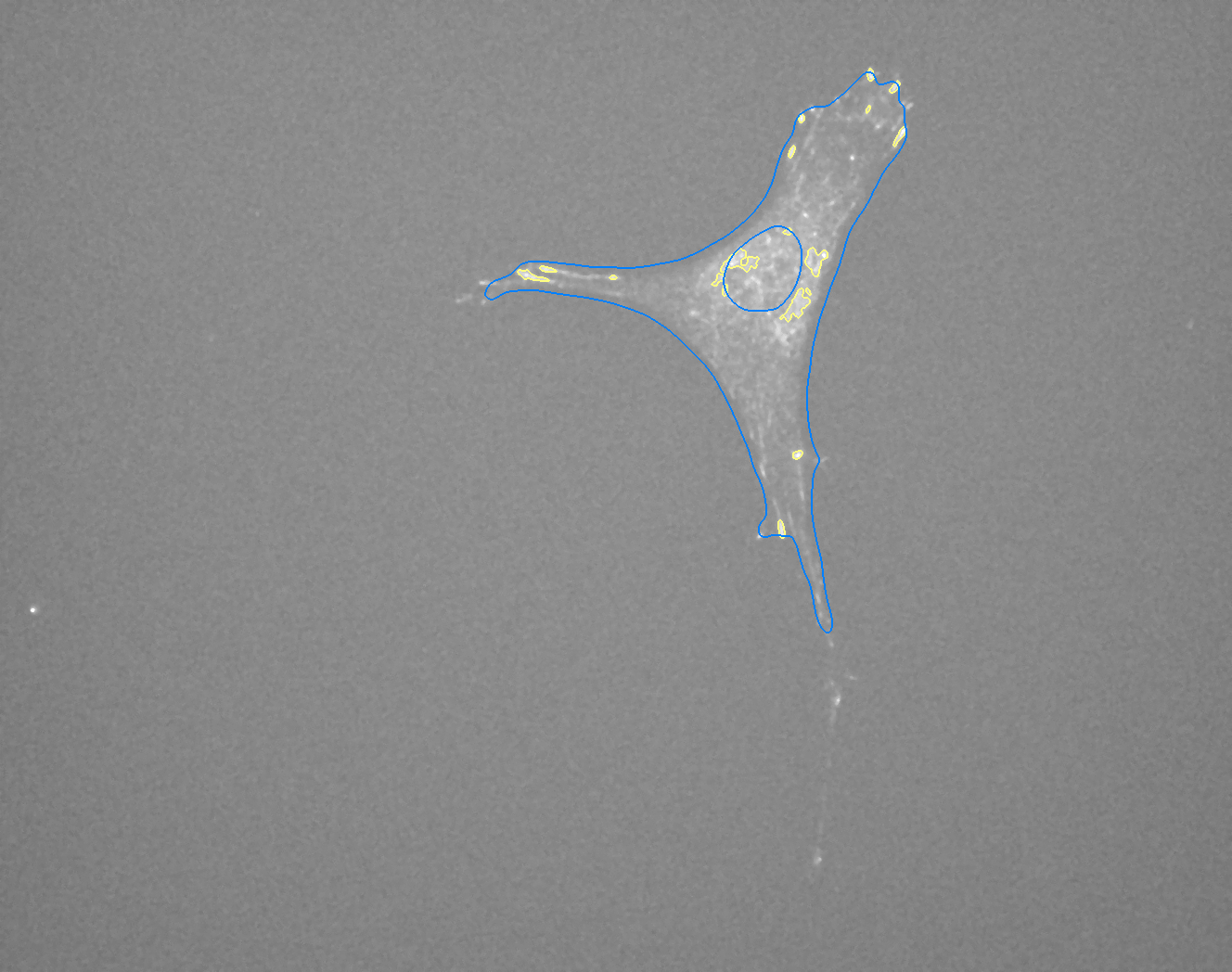

### H_Plate_R_p00_0_A01f11d0.TIF

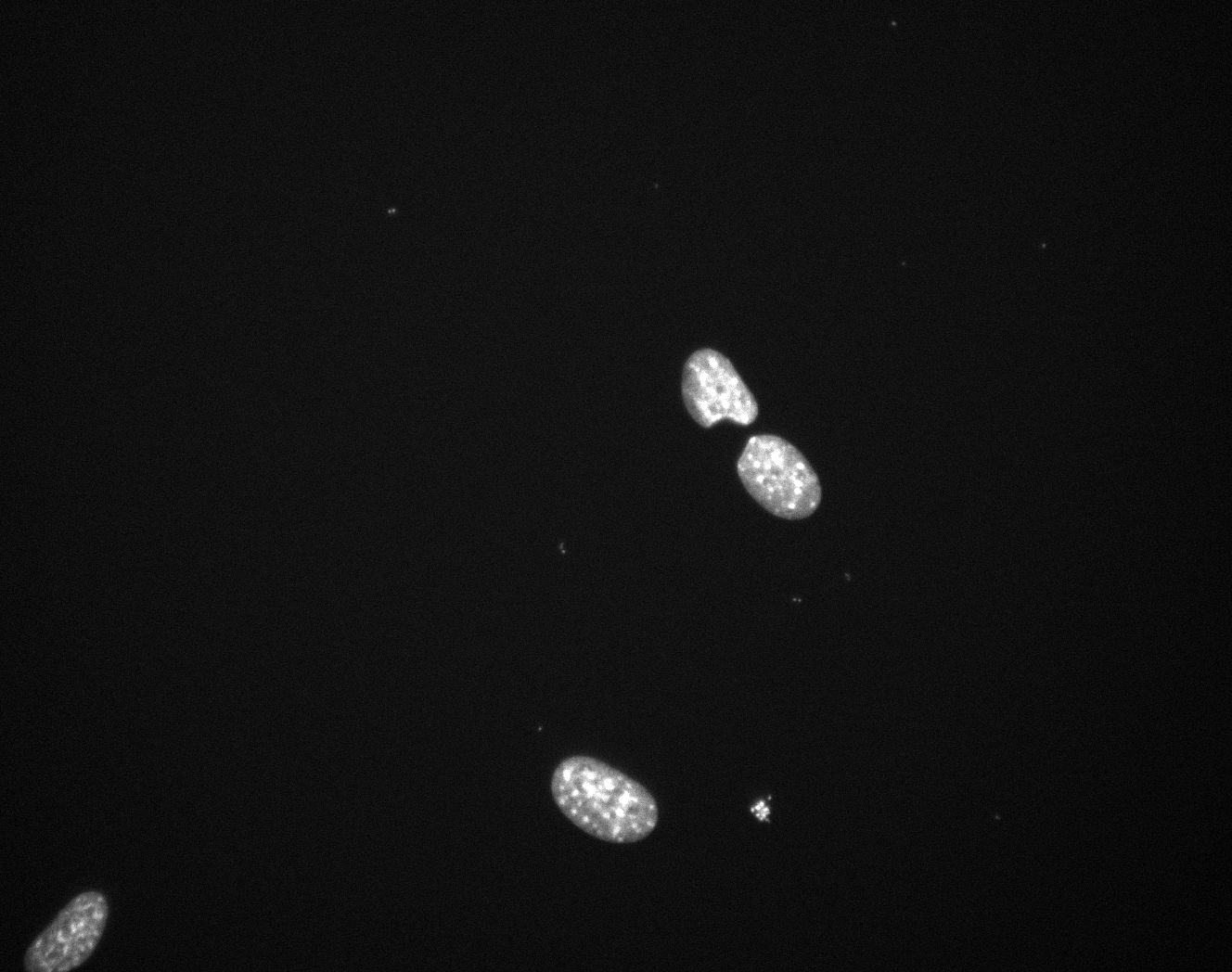

### H_Plate_R_p00_0_A01f11d1.TIF

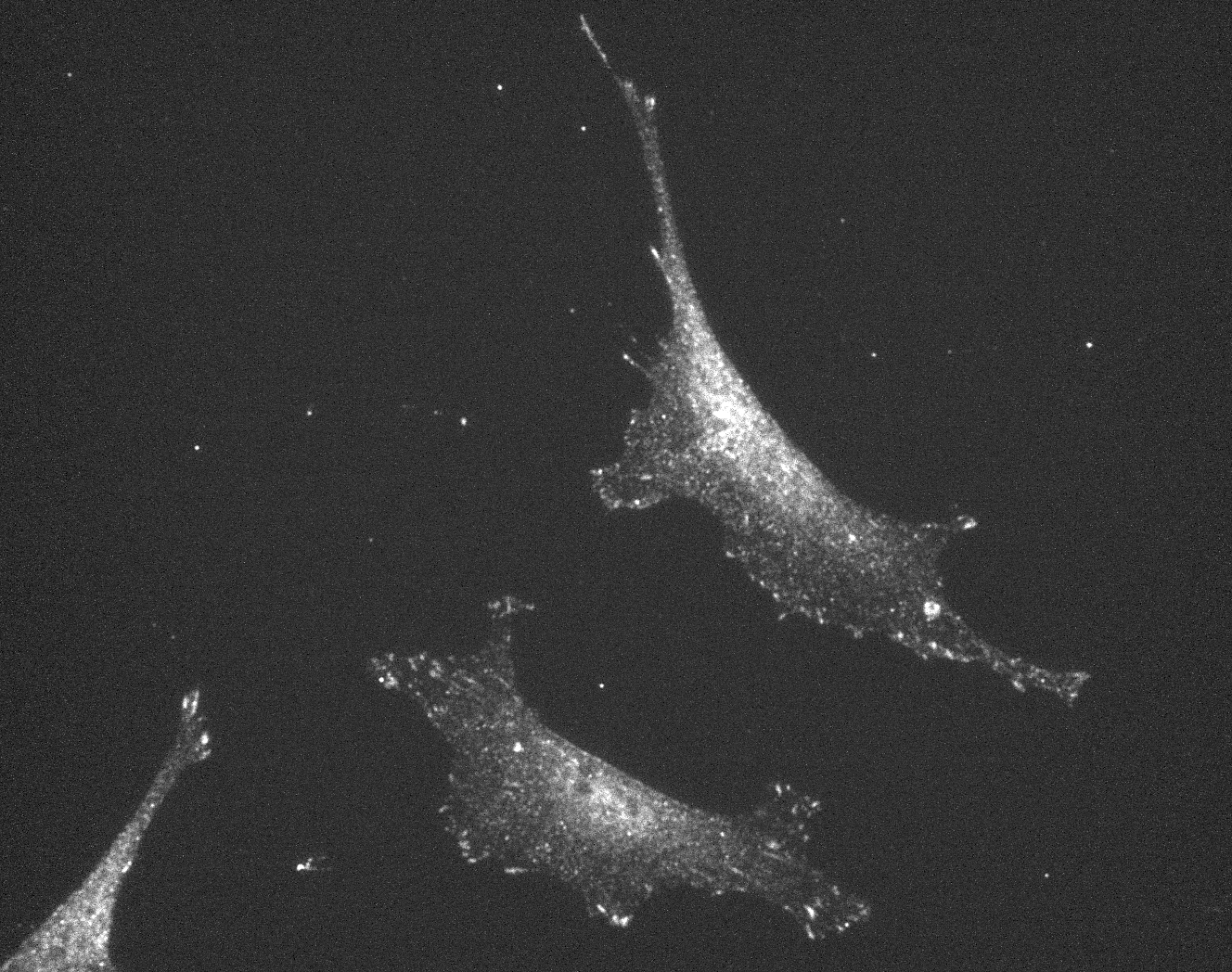

### H_Plate_R_p00_0_A01f11d2.TIF

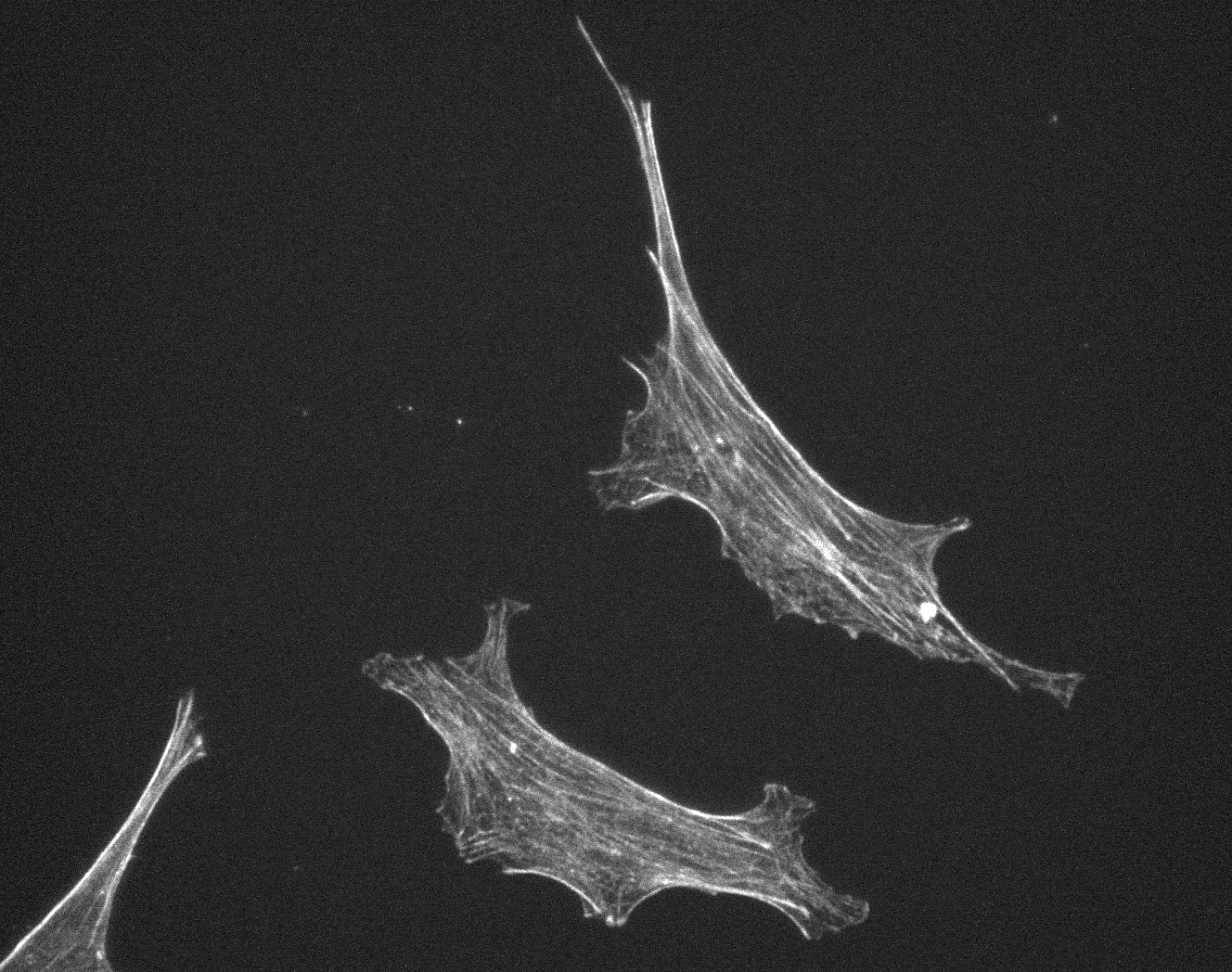

### H_Plate_R_p00_0_A01f11d3.TIF

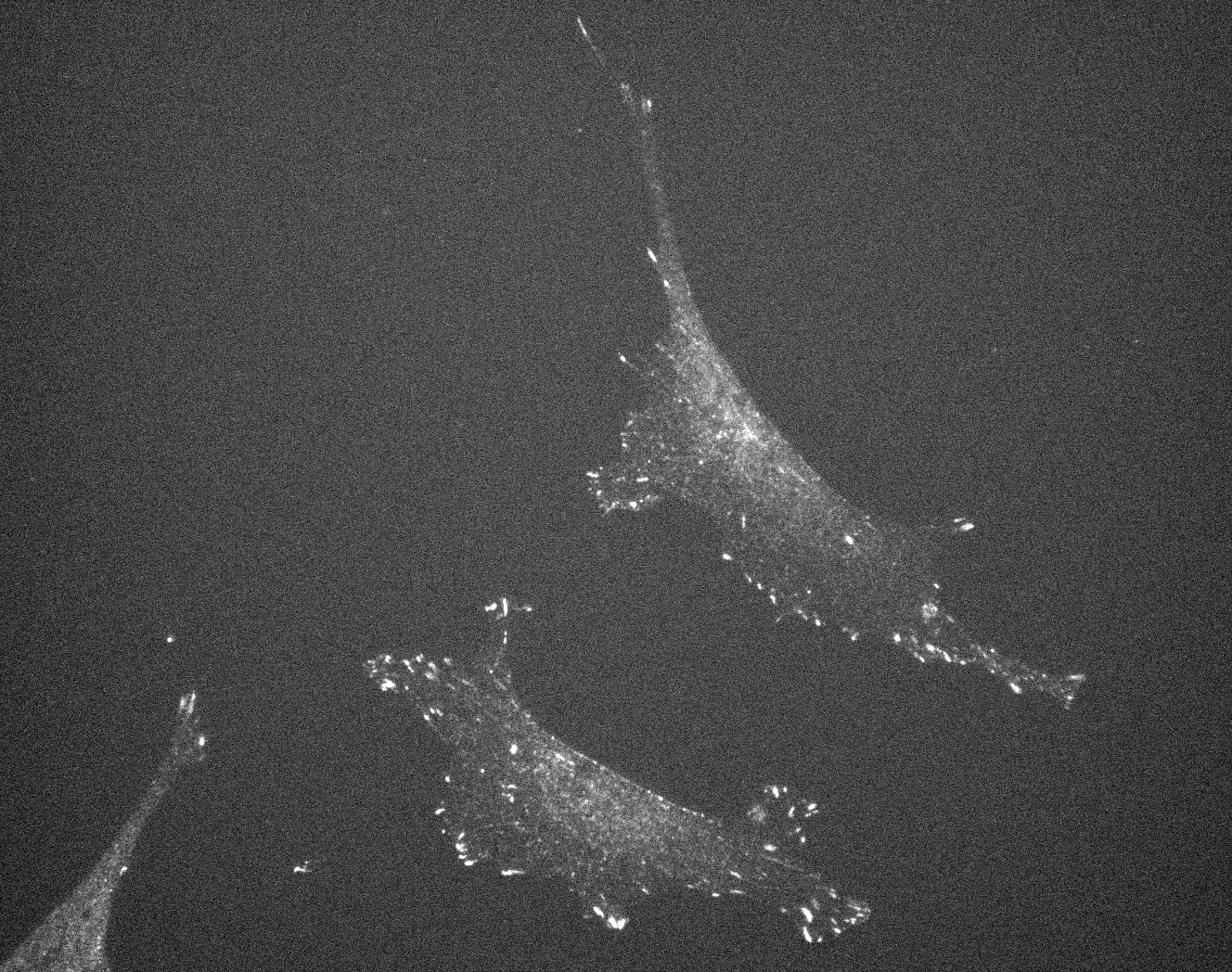

### H_Plate_R_p00_0_A01f19d0.TIF

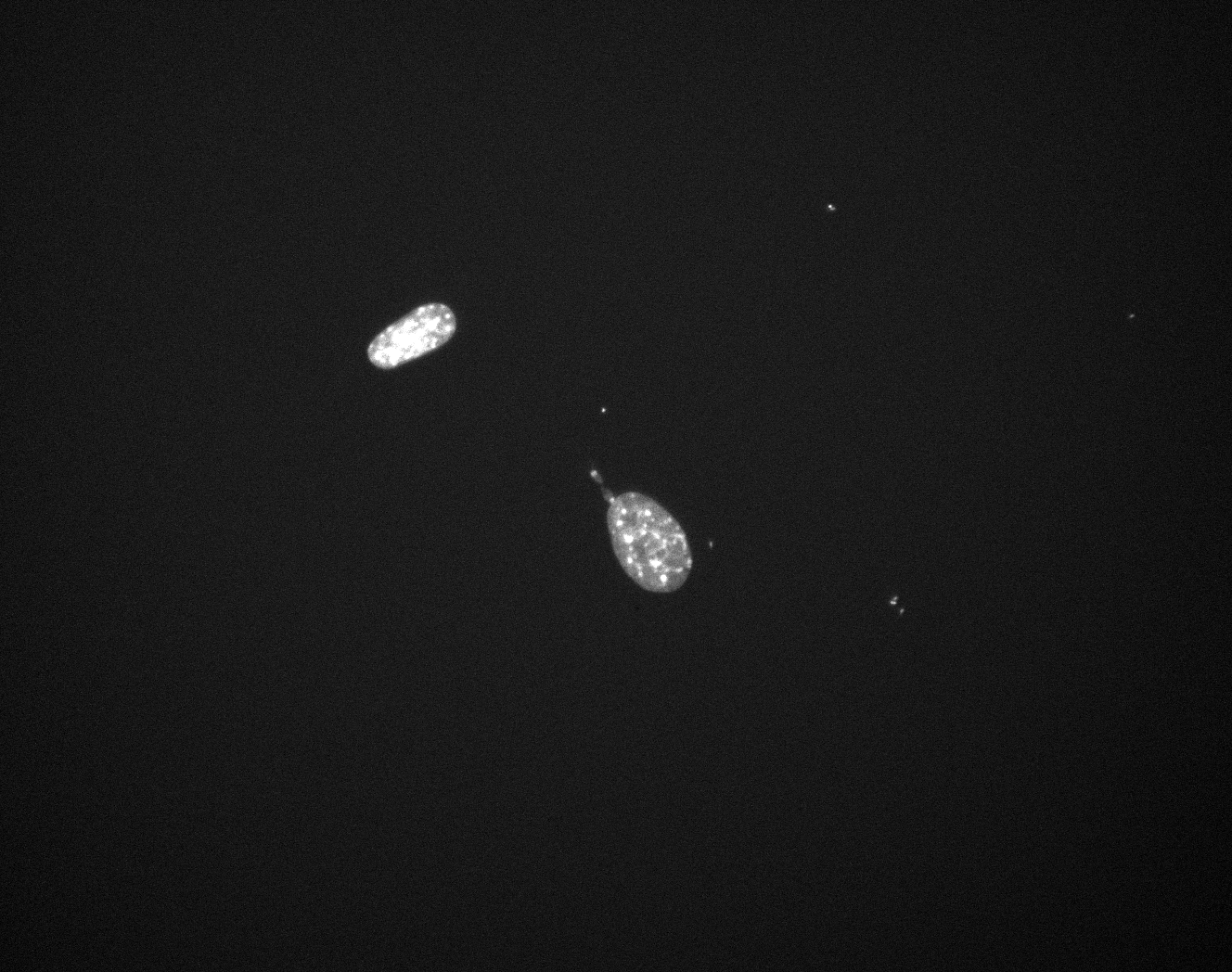

### H_Plate_R_p00_0_A01f19d1.TIF

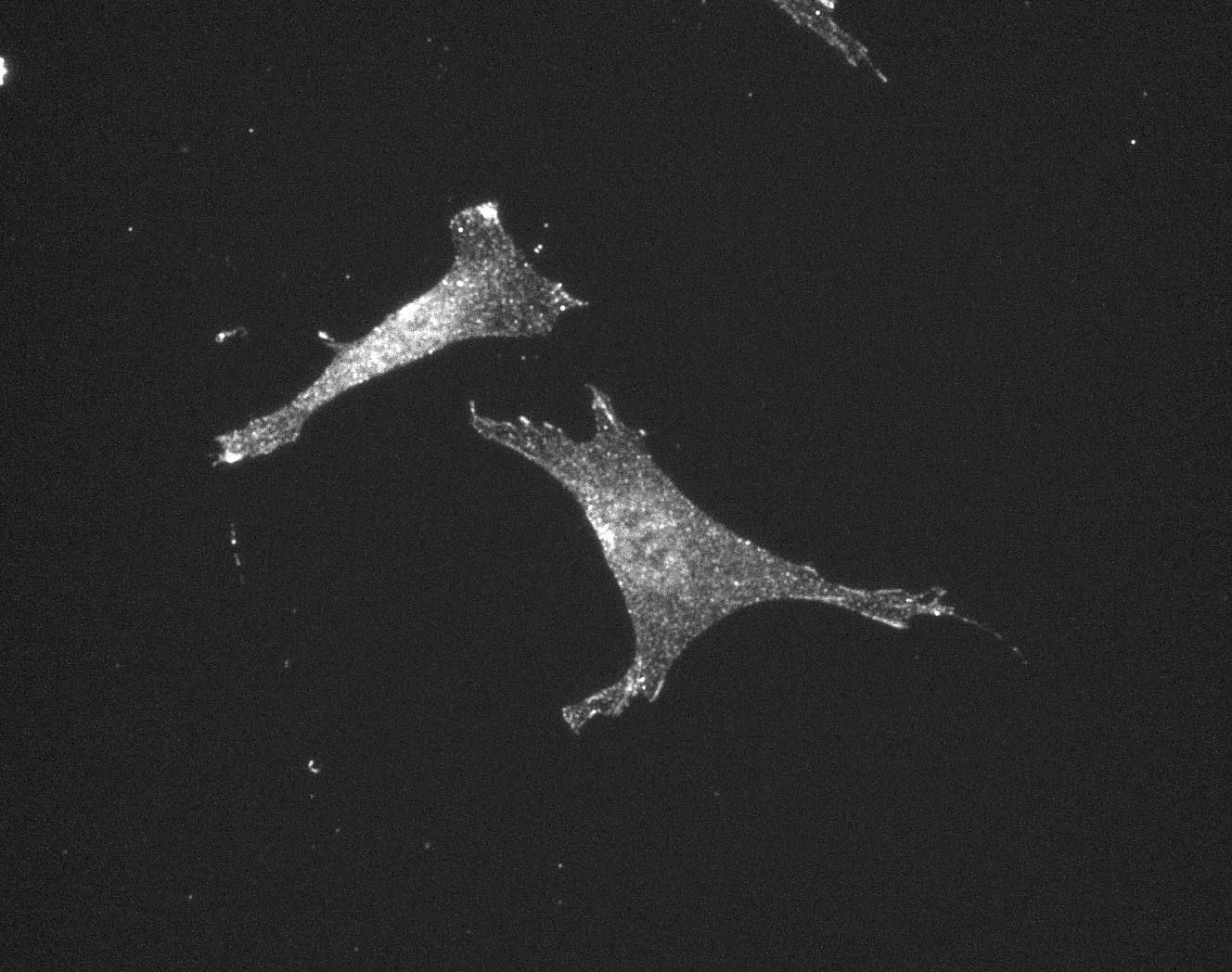

### H_Plate_R_p00_0_A01f19d2.TIF

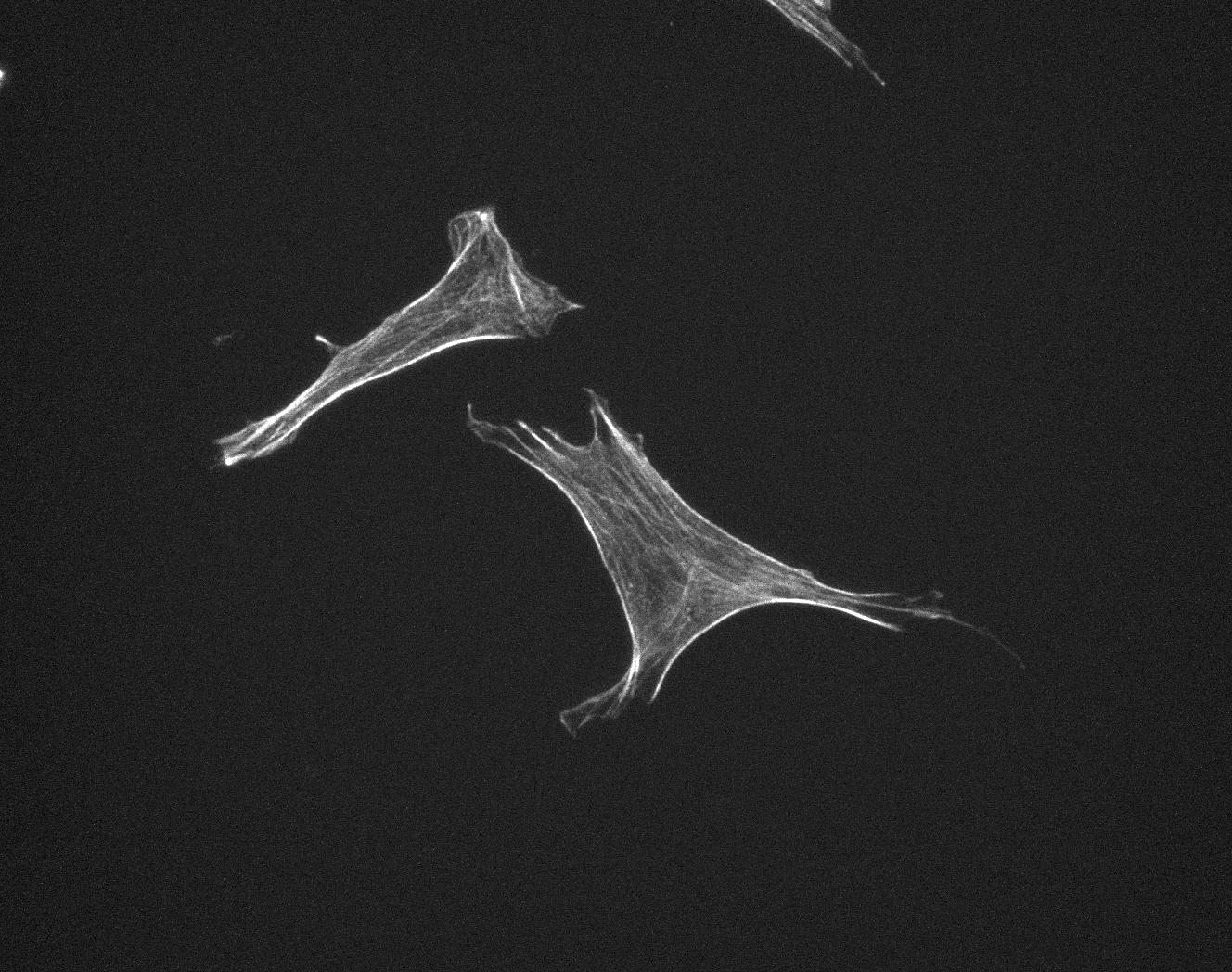

### H_Plate_R_p00_0_A01f19d2_CytoplasmOutline.tiff

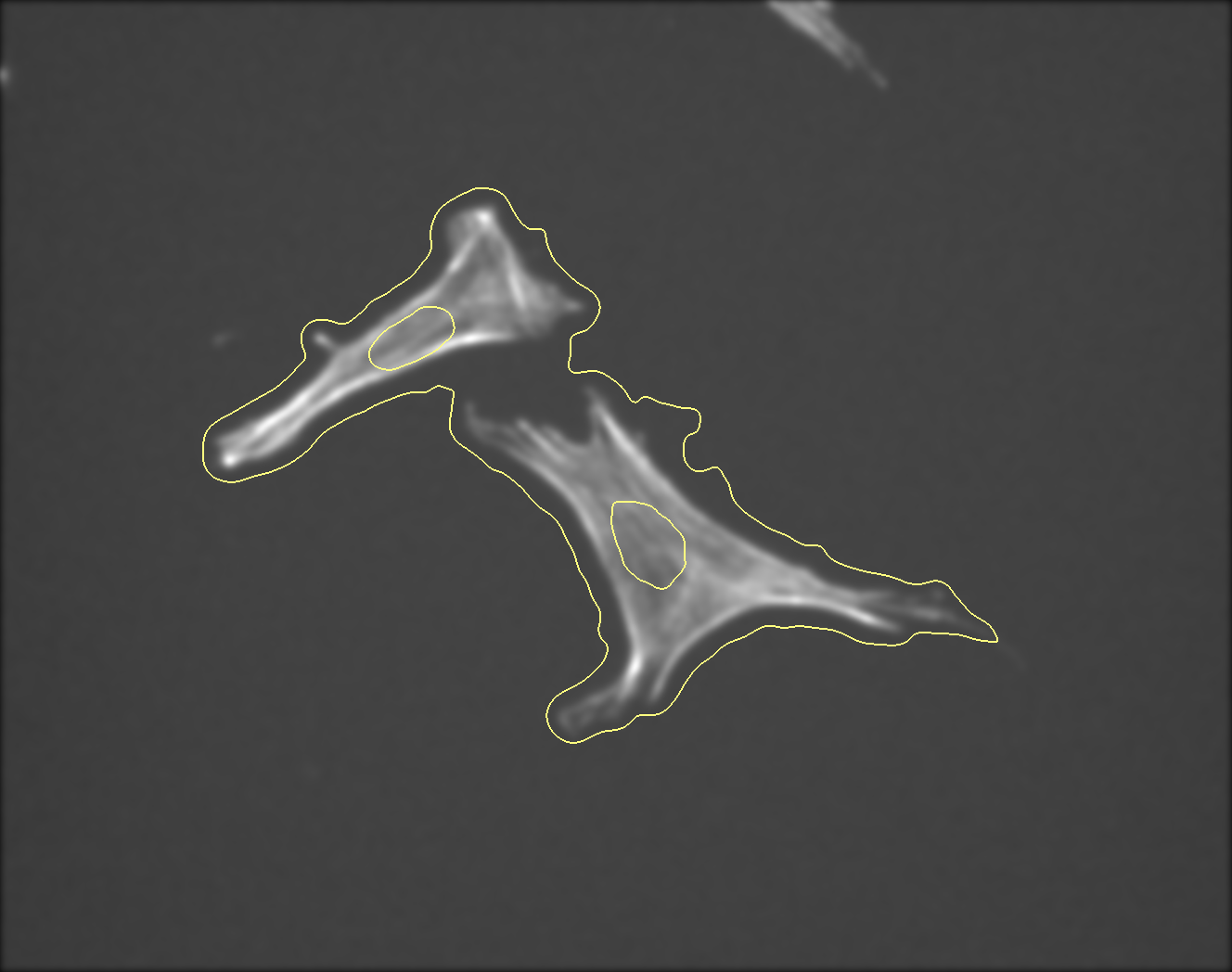

### H_Plate_R_p00_0_A01f19d3.TIF

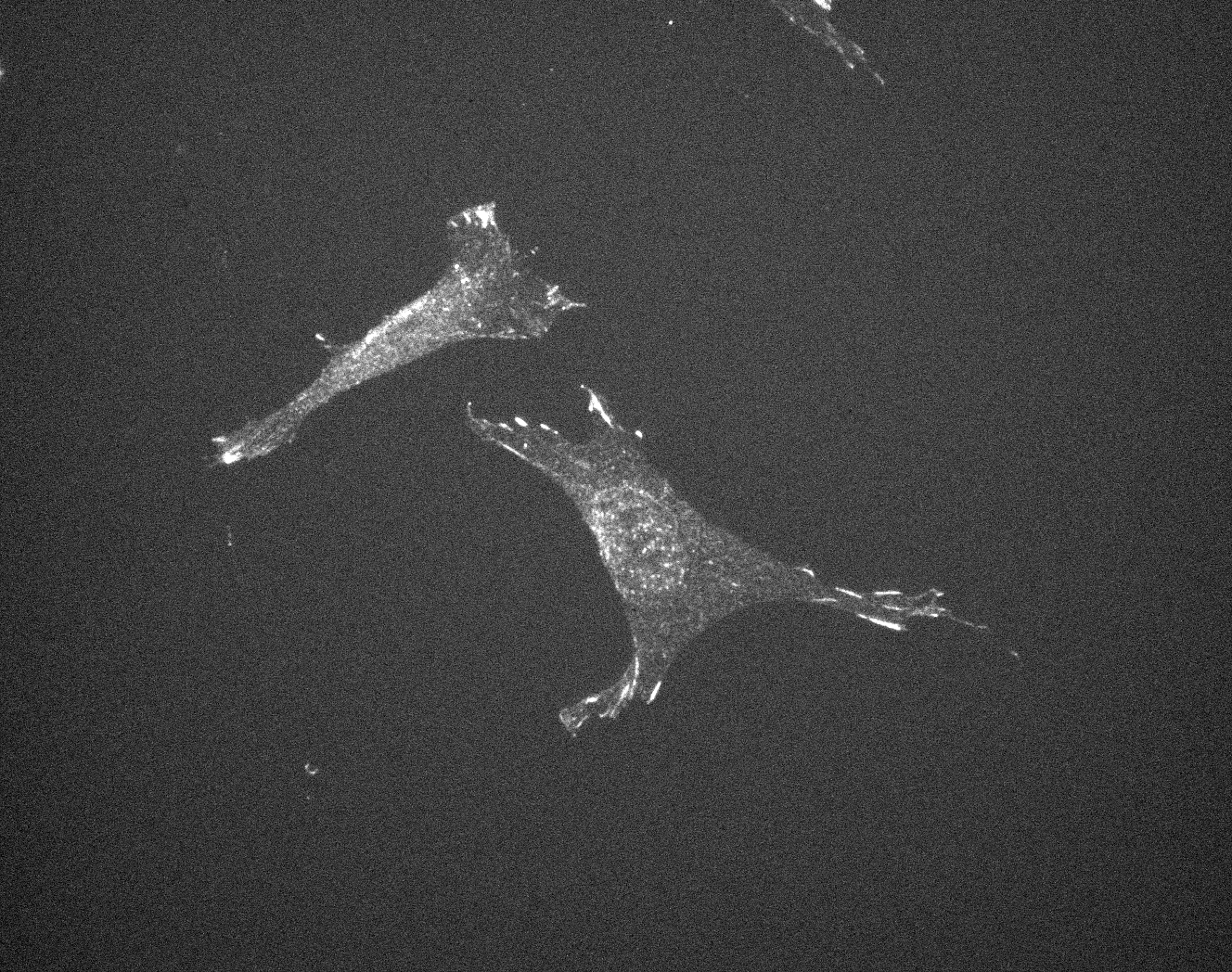

### H_Plate_R_p00_0_A01f41d3_FAOutline.tiff

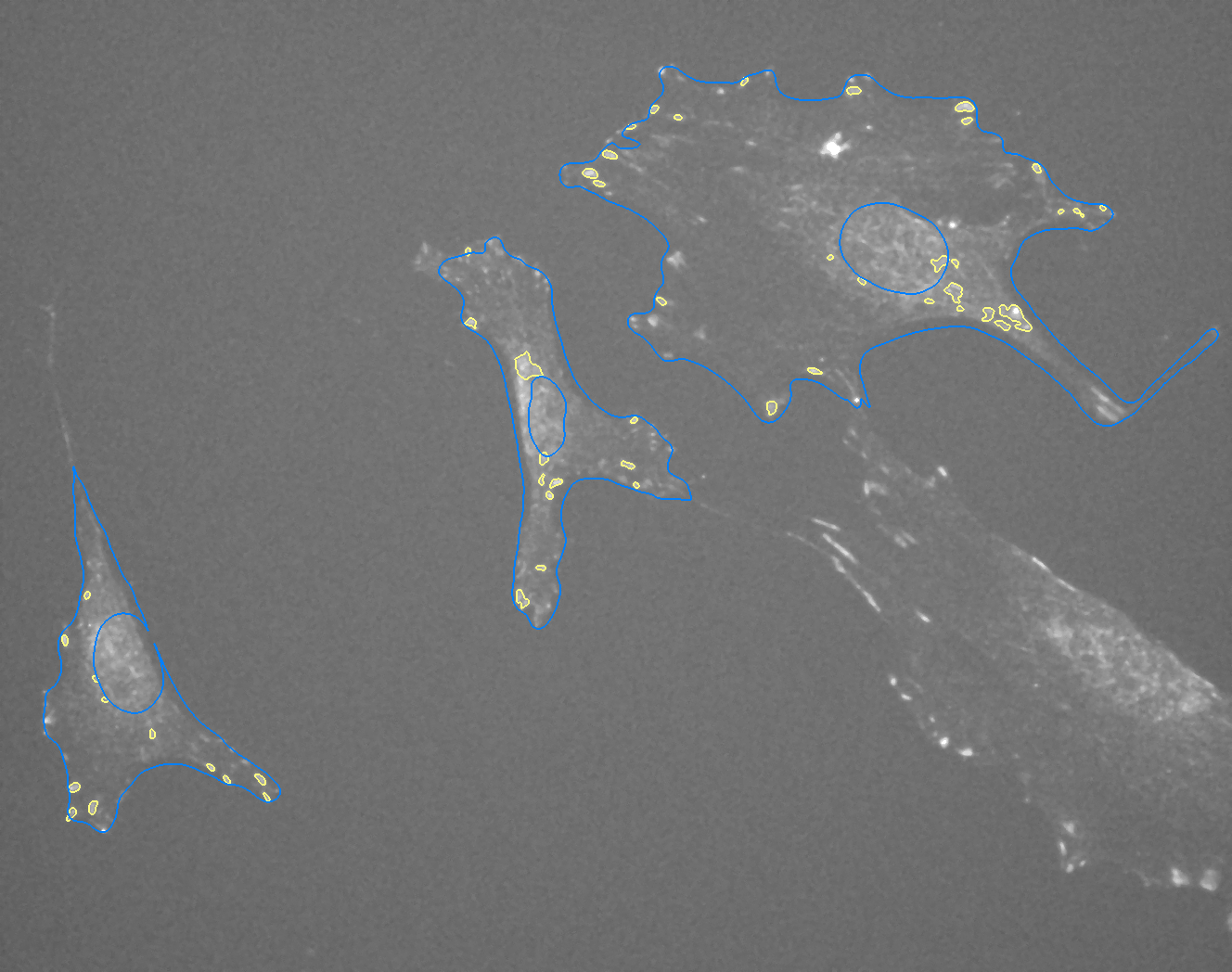

### I_Plate_R_p00_0_A01f39d3_FAOutline.tiff

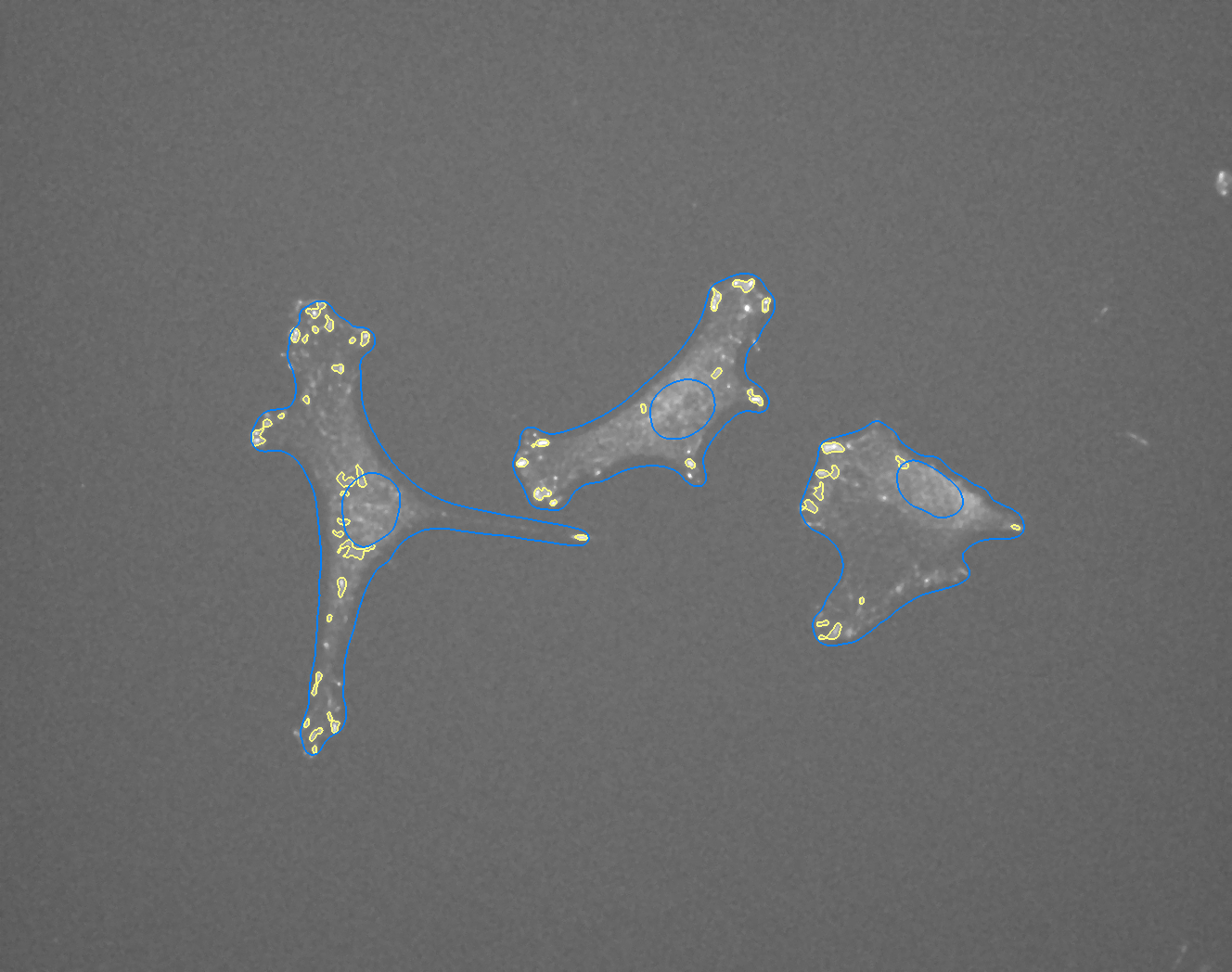

### I_Plate_R_p00_0_A01f44d0.TIF

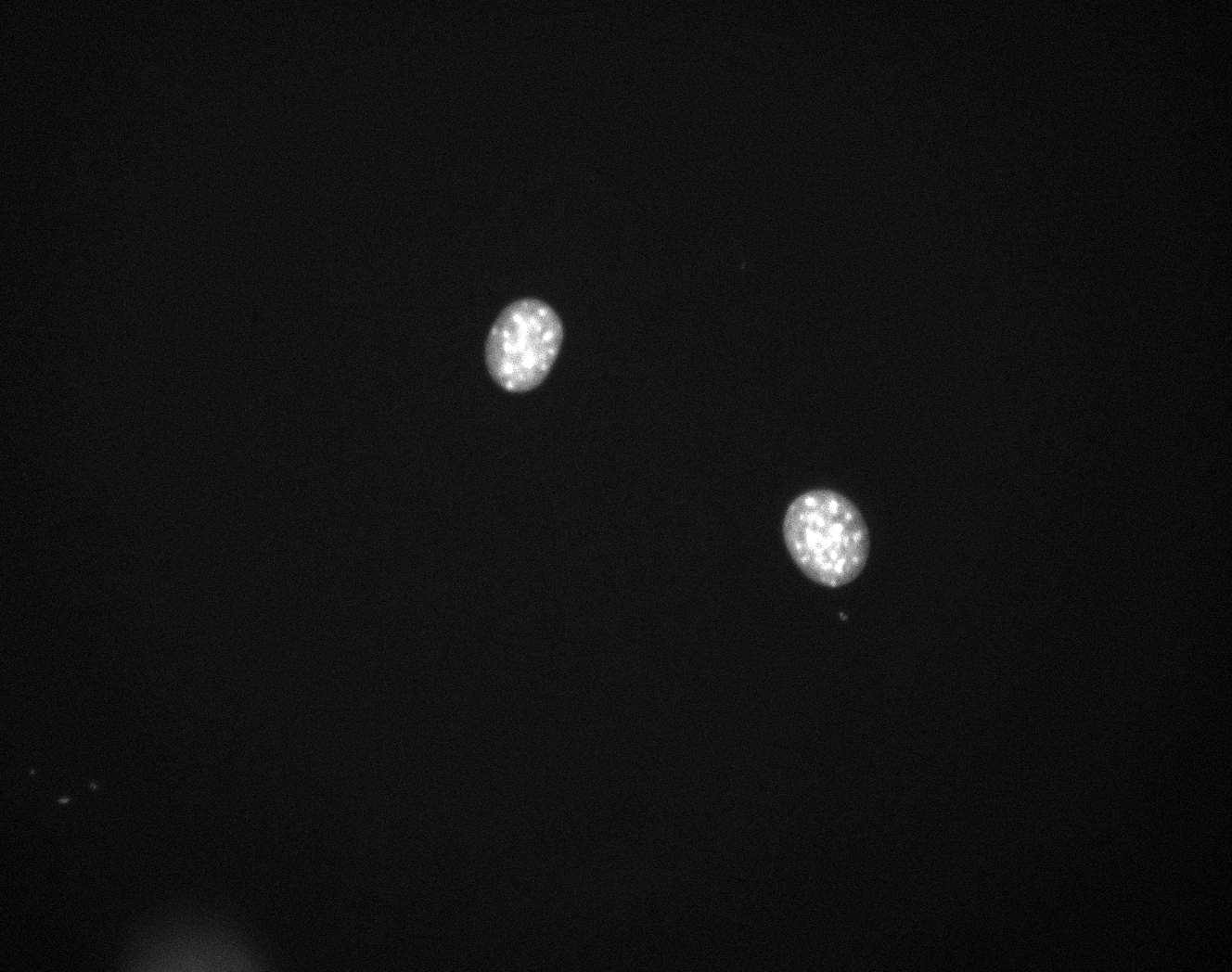

### I_Plate_R_p00_0_A01f44d1.TIF

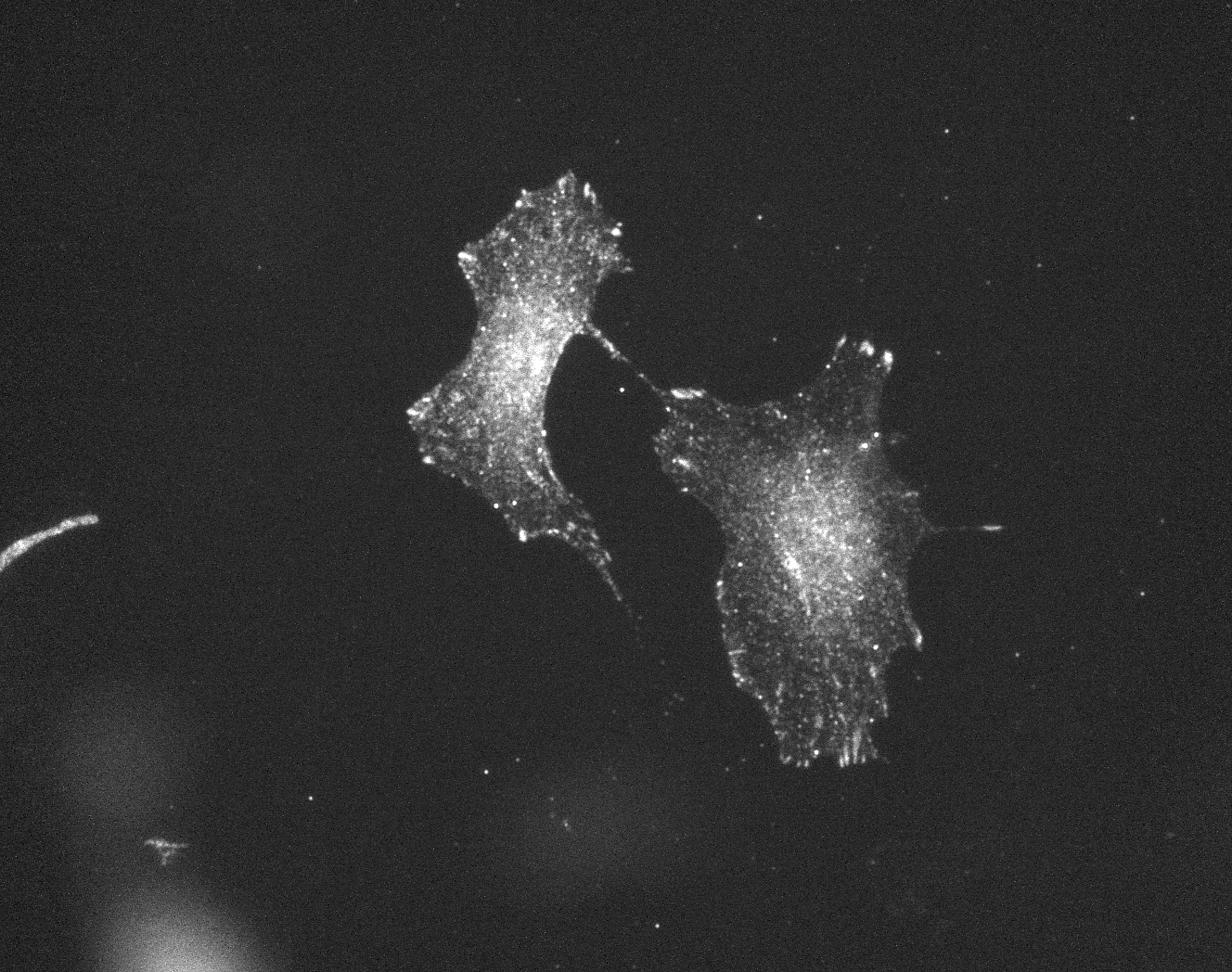

### I_Plate_R_p00_0_A01f44d2.TIF

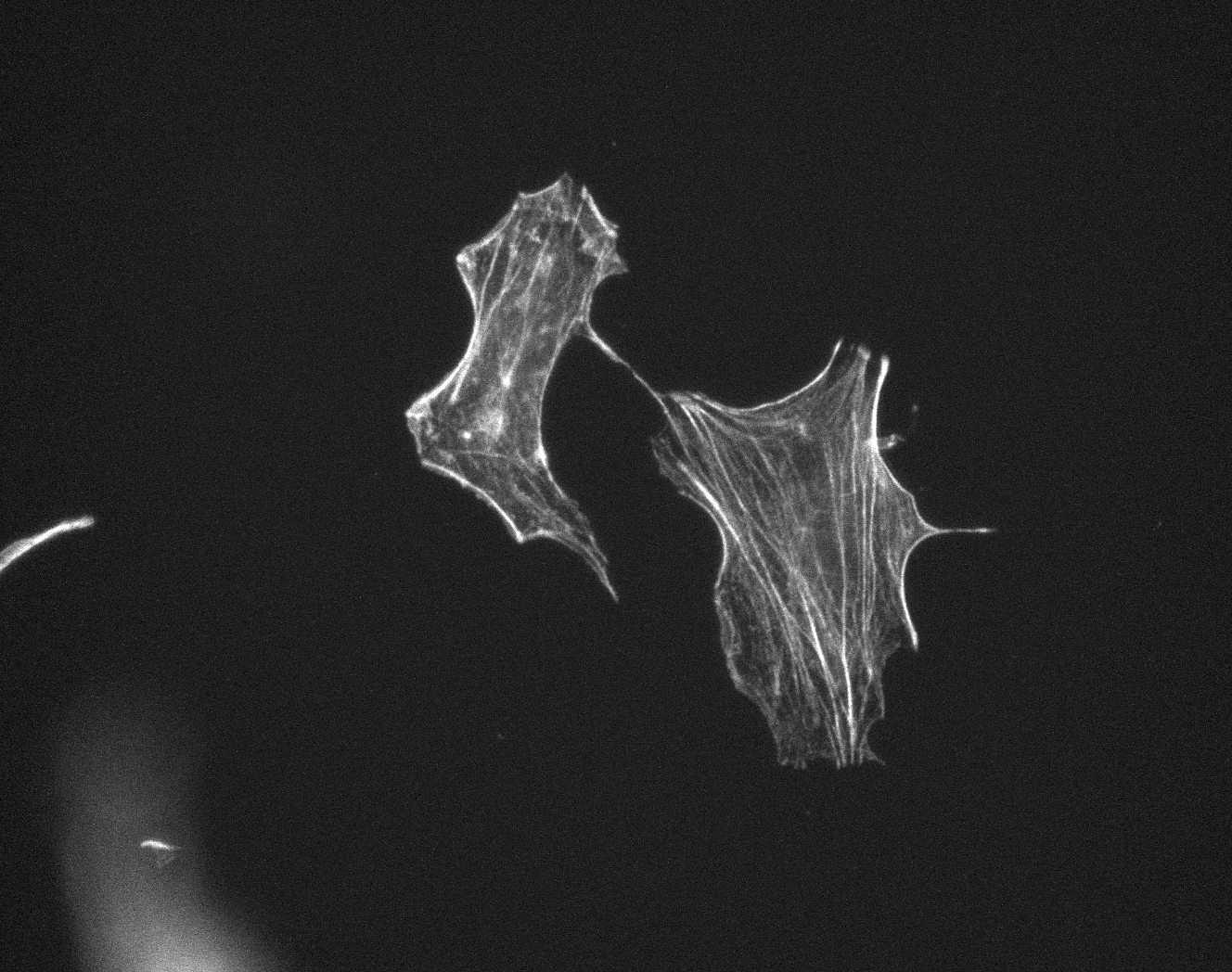

### I_Plate_R_p00_0_A01f44d2_CytoplasmOutline.tiff

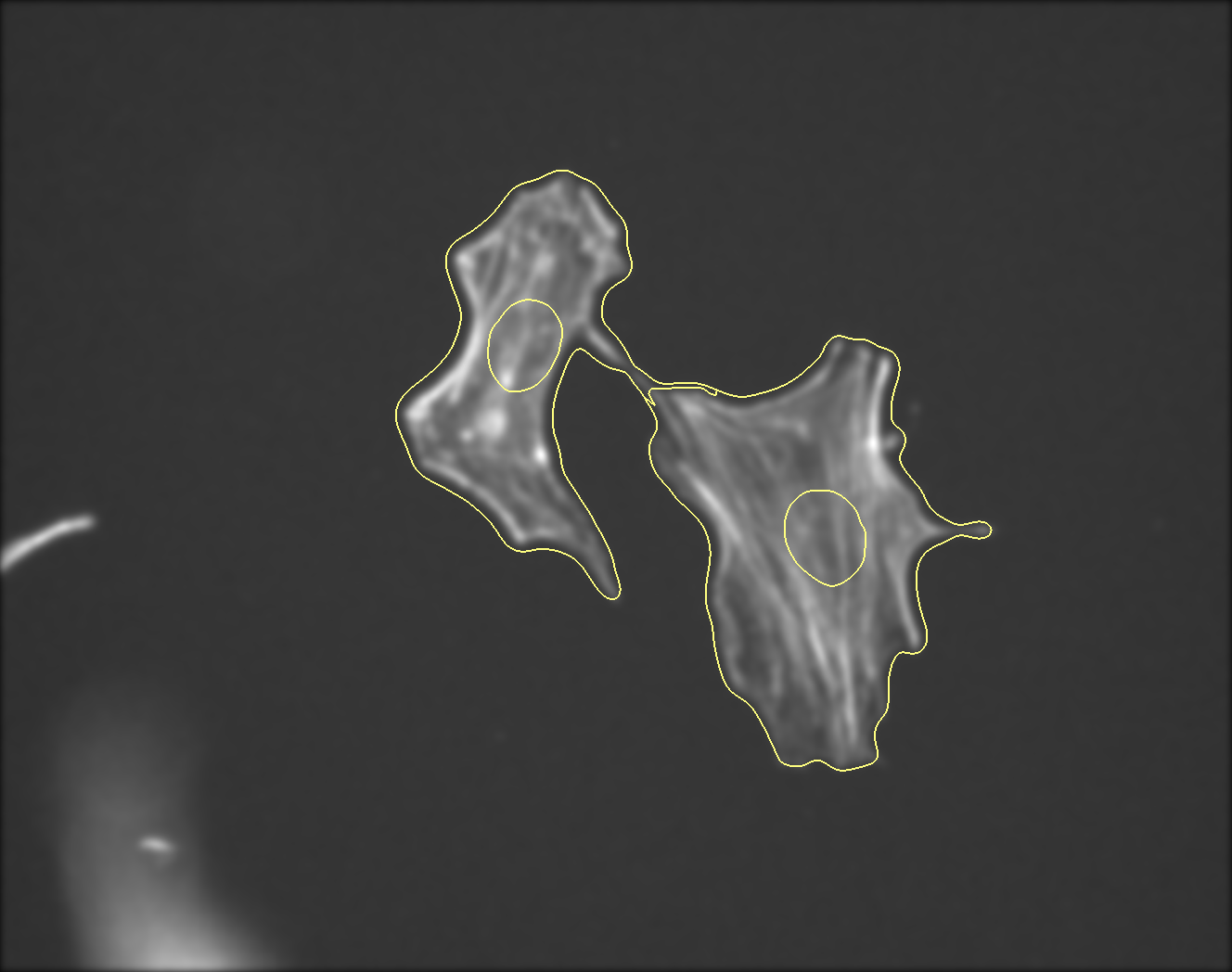

### I_Plate_R_p00_0_A01f44d3.TIF

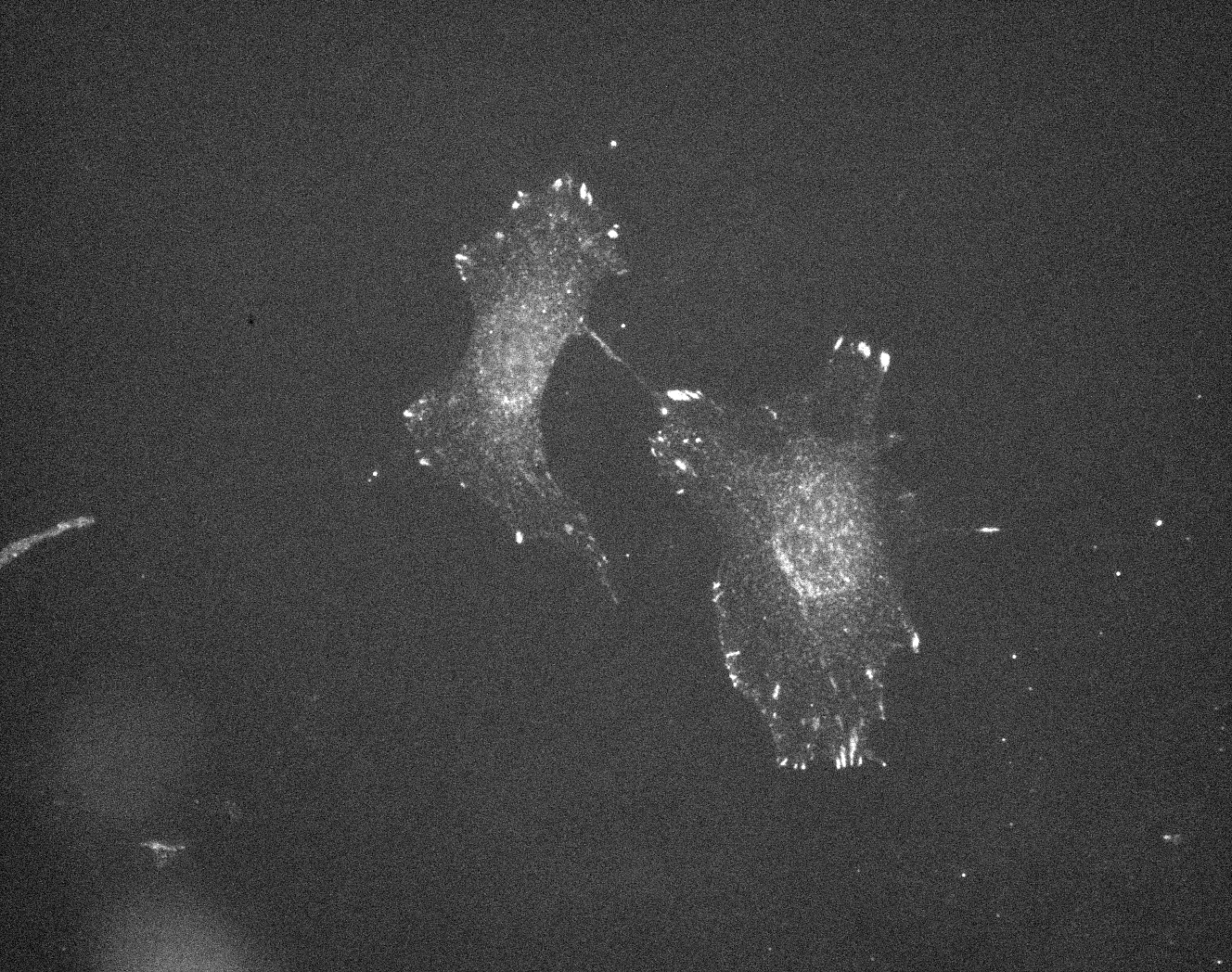

### J_Plate_R_p00_0_A01f49d3_FAOutline.tiff

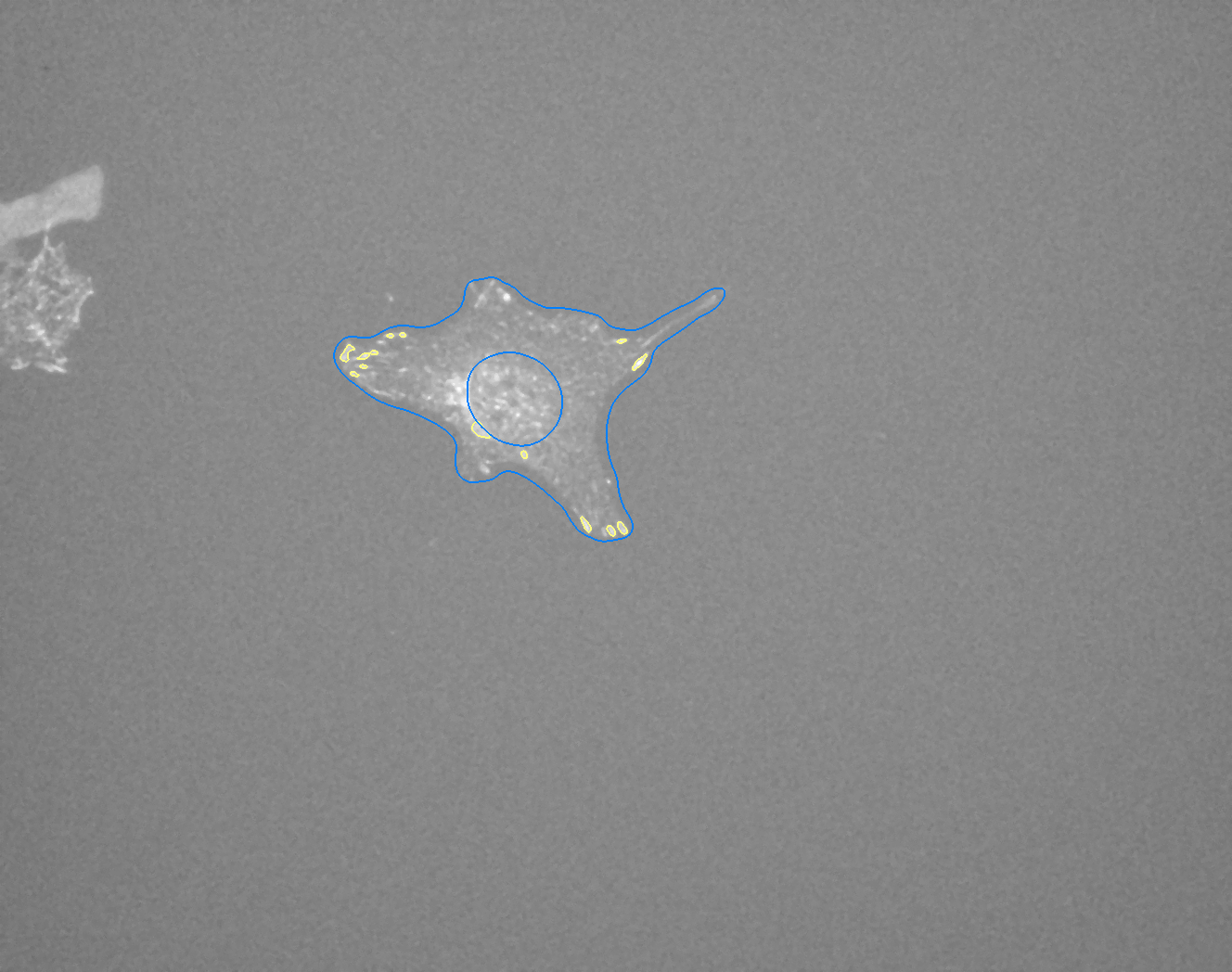

### J_Plate_R_p00_0_A01f51d0.TIF

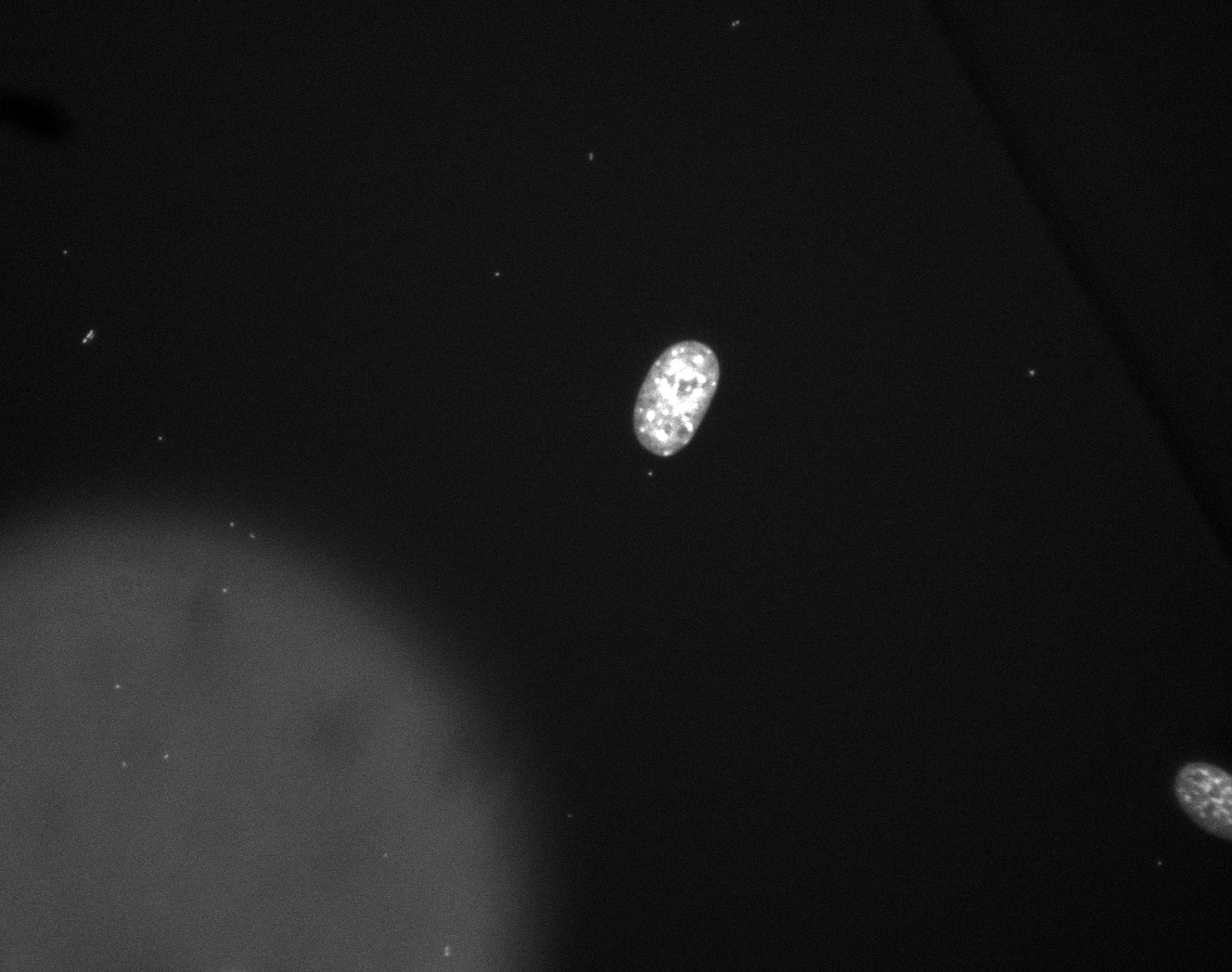

### J_Plate_R_p00_0_A01f51d1.TIF

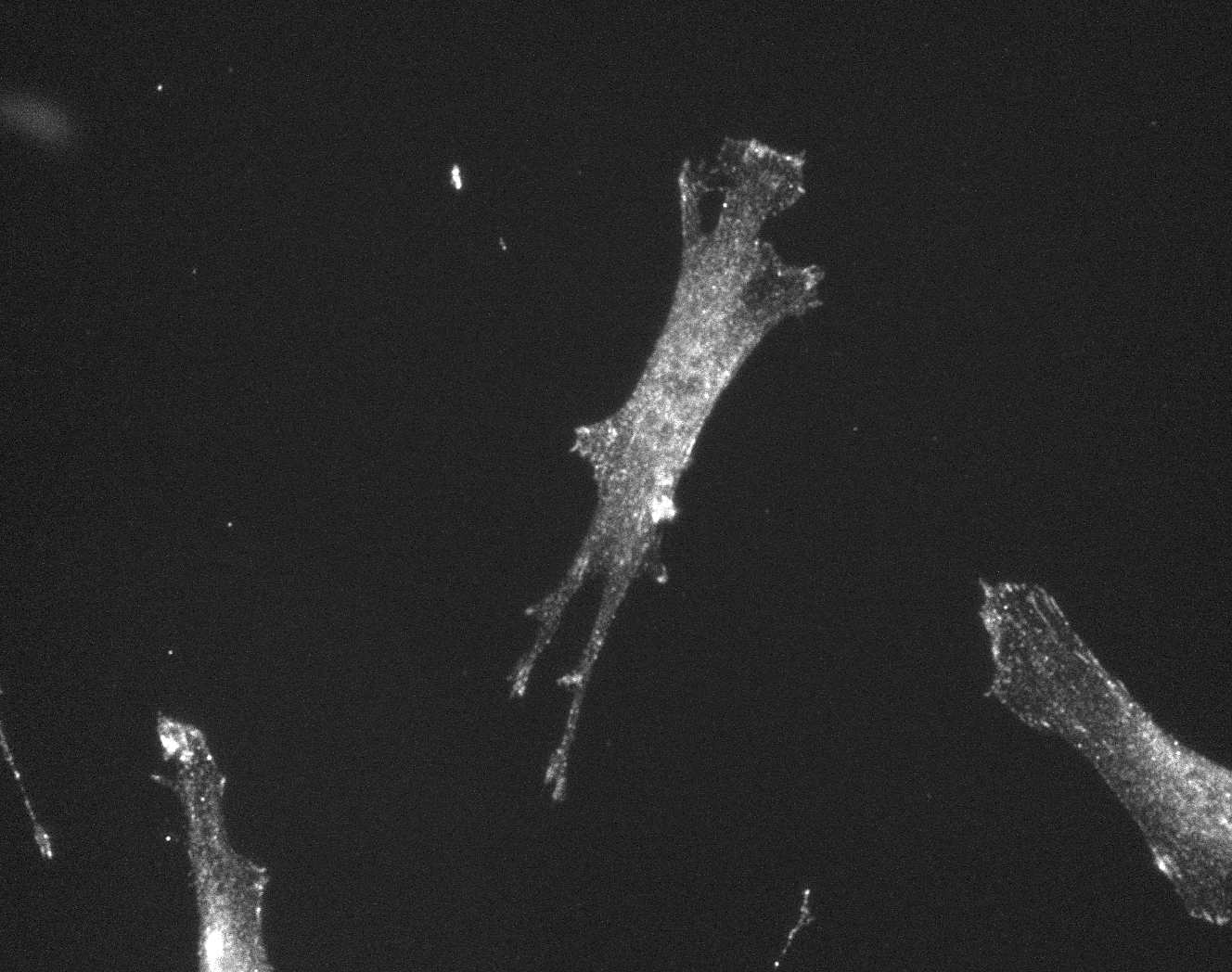

### J_Plate_R_p00_0_A01f51d2.TIF

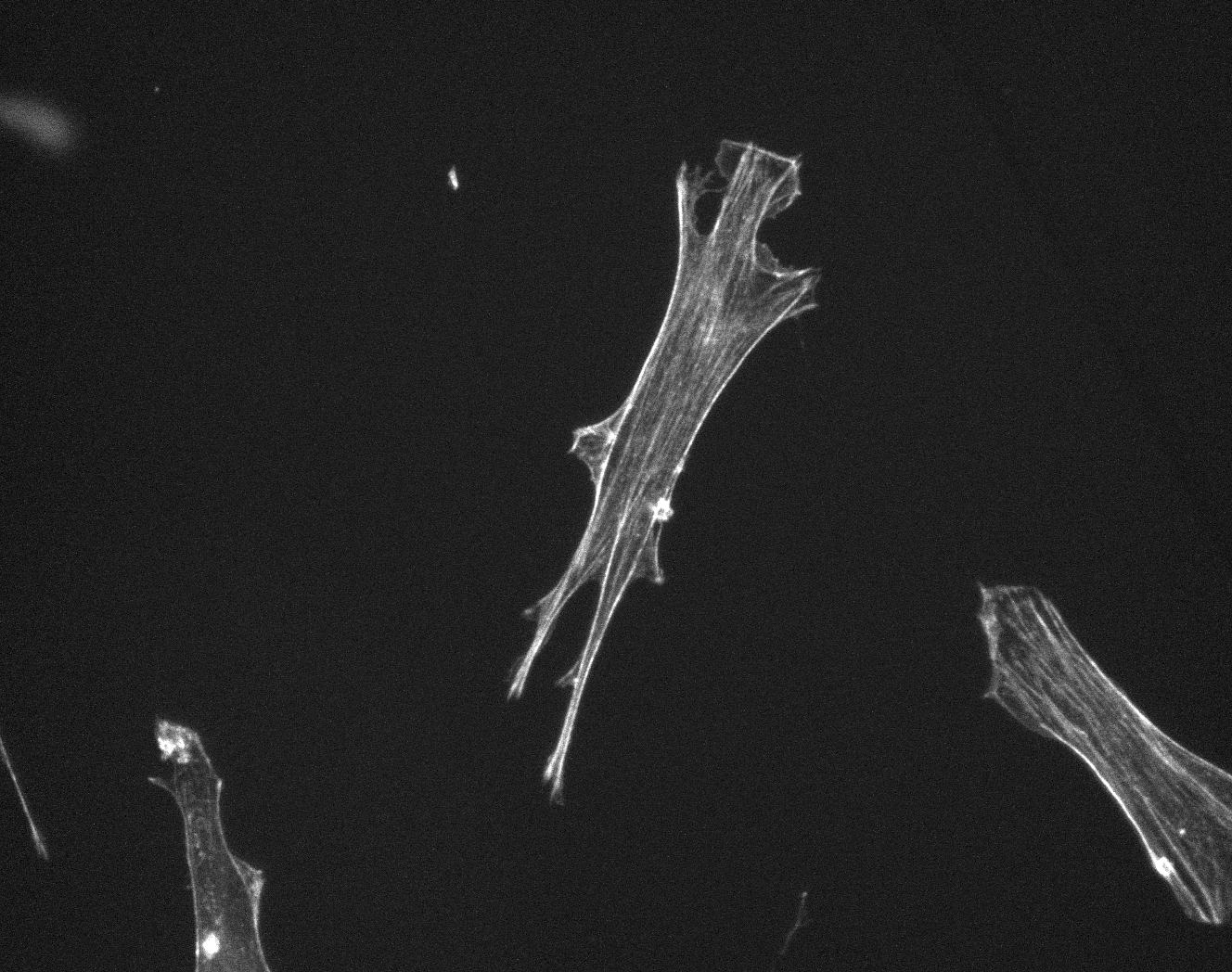

### J_Plate_R_p00_0_A01f51d2_CytoplasmOutline.tiff

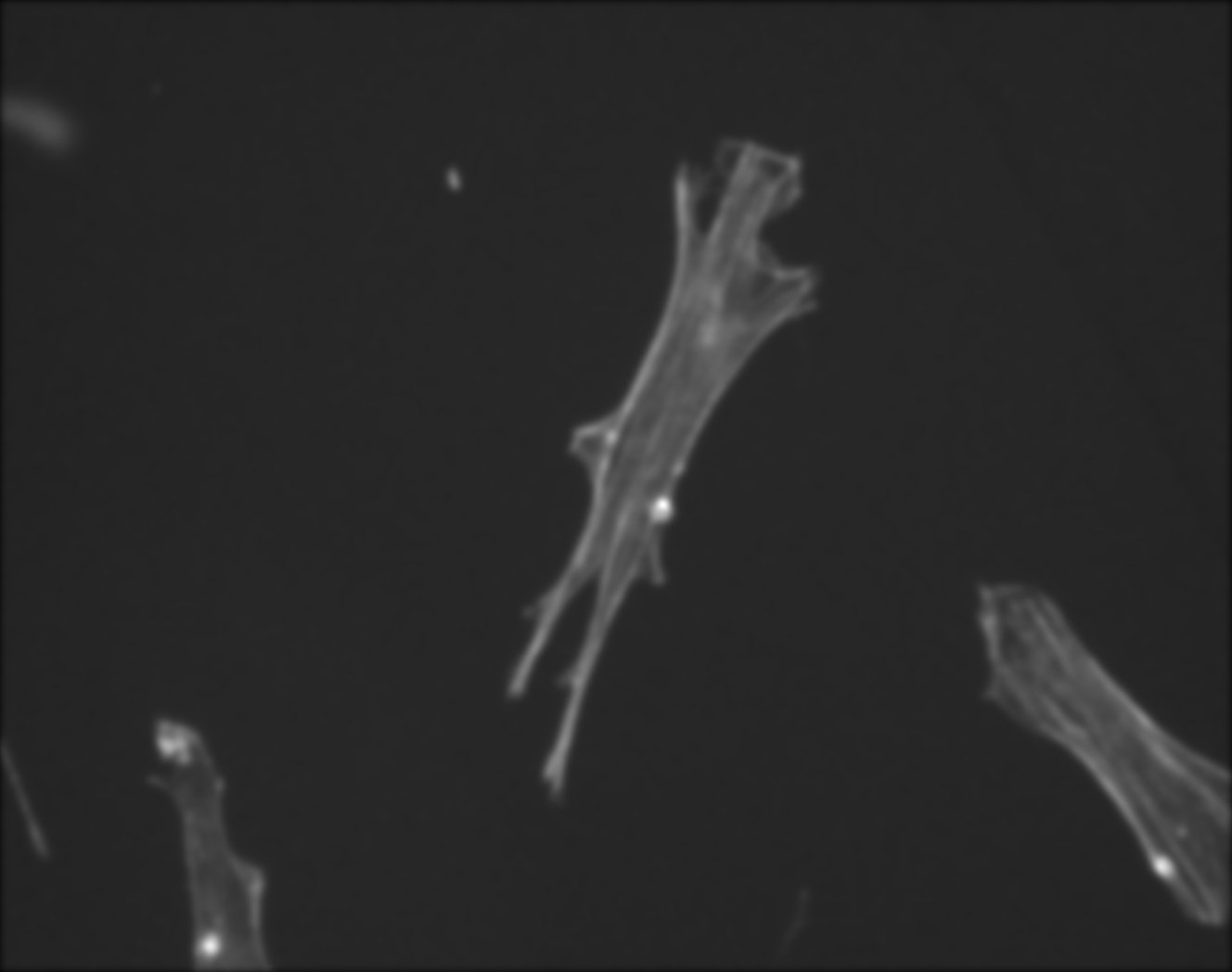

### J_Plate_R_p00_0_A01f51d3.TIF

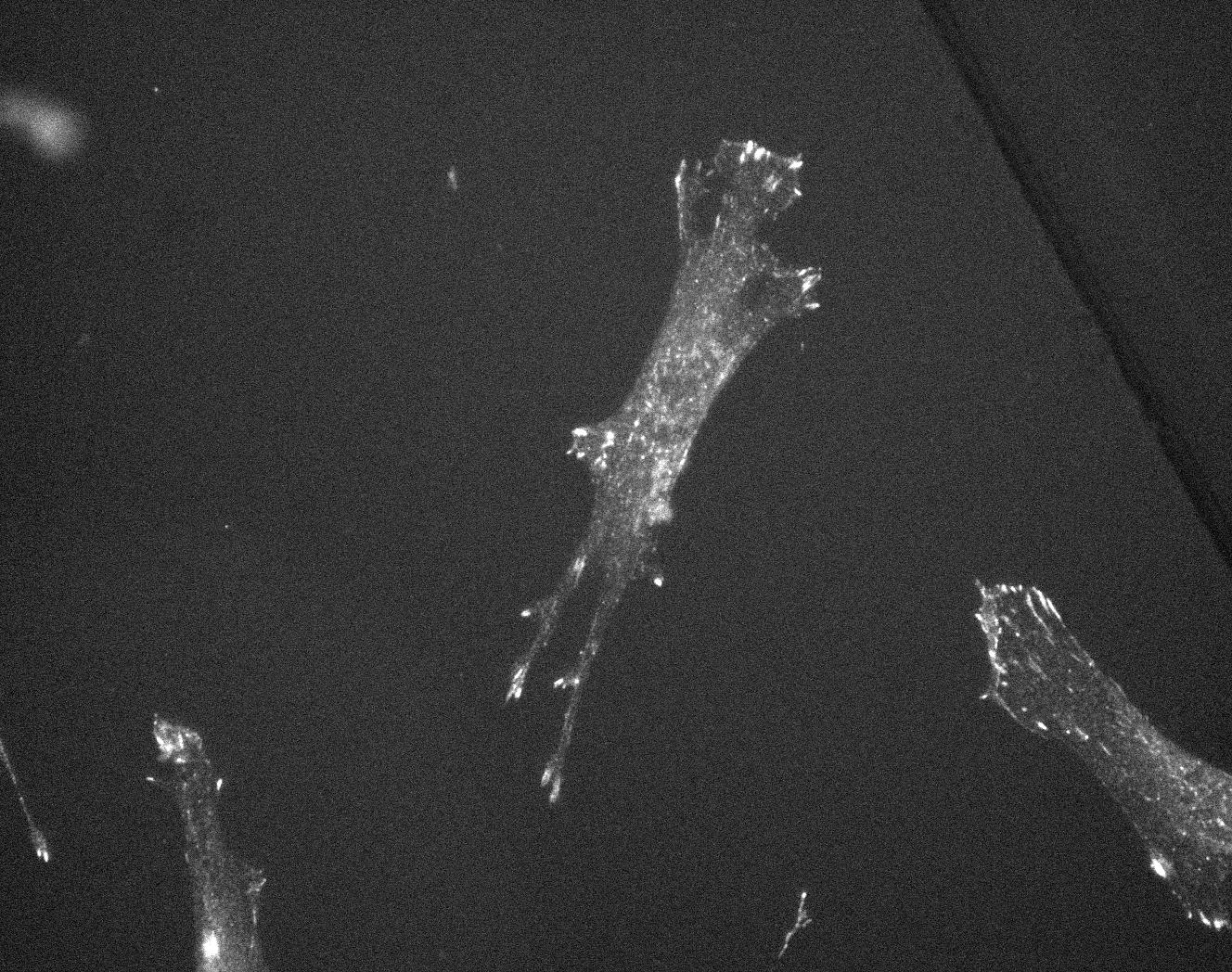
